# Insights into RAG evolution from the identification of “missing link” family A *RAGL* transposons

**DOI:** 10.1101/2023.08.20.553239

**Authors:** Eliza C. Martin, Lorlane Le Targa, Louis Tsakou-Ngouafo, Tzu-Pei Fan, Che-Yi Lin, Jianxiong Xiao, Yi Hsien Su, Andrei-Jose Petrescu, Pierre Pontarotti, David G. Schatz

## Abstract

A series of “molecular domestication” events are thought to have converted an invertebrate RAG-like (RAGL) transposase into the RAG1-RAG2 (RAG) recombinase, a critical enzyme for adaptive immunity in jawed vertebrates. The timing and order of these events is not well understood, in part because of a dearth of information regarding the invertebrate *RAGL-A* transposon family. In contrast to the abundant and divergent *RAGL-B* transposon family, *RAGL-A* most closely resembles *RAG* and is represented by a single orphan *RAG1-like* (*RAG1L*) gene in the genome of the hemichordate *Ptychodera flava* (*PflRAG1L-A*). Here, we provide evidence for the existence of complete *RAGL-A* transposons in the genomes of *P. flava* and several echinoderms. The predicted RAG1L-A and RAG2L-A proteins encoded by these transposons intermingle sequence features of jawed vertebrate RAG and RAGL-B transposases, leading to a prediction of DNA binding, catalytic, and transposition activities that are a hybrid of RAG and RAGL-B. Similarly, the terminal inverted repeats (TIRs) of the *RAGL-A* transposons combine features of both *RAGL-B* transposon TIRs and RAG recombination signal sequences. Unlike all previously described RAG2L proteins, PflRAG2L-A and echinoderm RAG2L-A contain an acidic hinge region, which we demonstrate is capable of efficiently inhibiting RAG-mediated transposition. Our findings provide evidence for a critical intermediate in RAG evolution and argue that certain adaptations thought to be specific to jawed vertebrates (e.g., the RAG2 acidic hinge) actually arose in invertebrates, thereby focusing attention on other adaptations as the pivotal steps in the completion of RAG domestication in jawed vertebrates.

## INTRODUCTION

V(D)J recombination is essential for adaptive immunity in jawed vertebrates and in many species is responsible for generating the vast repertoire of antigen receptors expressed by developing lymphocytes (Flajnik 2014; Gellert 2002). A heterotetramer composed of RAG1 and RAG2 (hereafter, RAG) initiates V(D)J recombination by cleaving DNA at specific recombination signal sequences (RSSs) that flank each V, D, and J gene segment that participates in the reaction (Figure 1A) (Kim et al. 2015; Schatz and Swanson 2011). RSSs consist of conserved heptamer and nonamer components separated by either a 12- or 23-bp spacer and cleavage occurs efficiently only in a synaptic complex containing a 12RSS/23RSS pair (the 12/23 rule) (Figure 1A) (Gellert 2002). RAG1 is composed of a core region essential for DNA cleavage (aa 384-1008; mouse RAG aa numbers are used unless otherwise specified) flanked by a long N-terminal region (NTR; aa 1-383) and a short C-terminal tail (CTT; aa 1009-1040). RAG2 consists of a core region required for cleavage activity (aa 1-350), an acidic ‘hinge’ (AH) region of approx. 60 aa, and a C-terminal plant homeodomain (PHD) (Kim et al. 2015; Matthews et al. 2007; Schatz and Swanson 2011).

**Figure 1.**
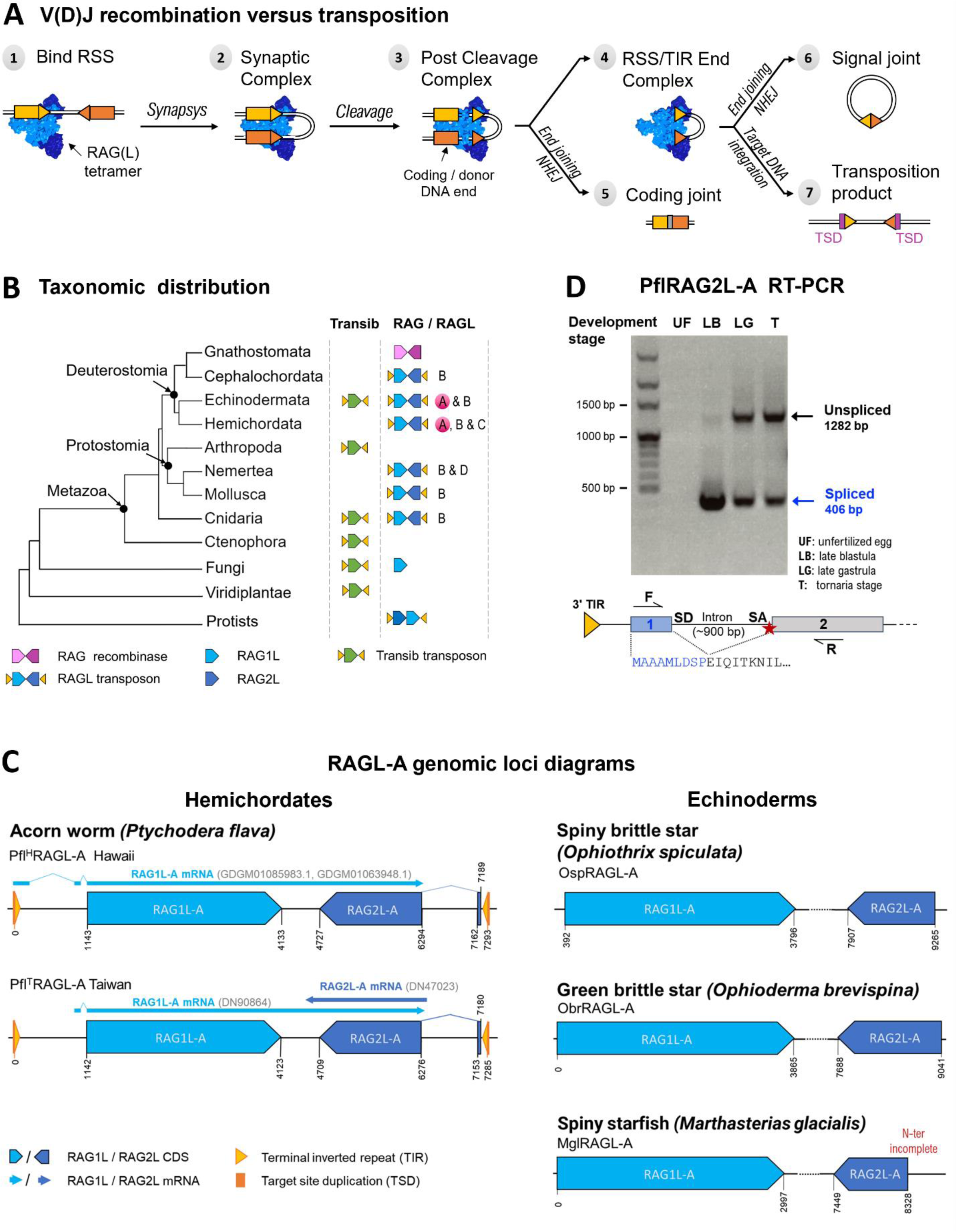
Overview of *RAGL-A* transposons in hemichordates and echinoderms. **A) Schematic of V(D)J recombination and transposition.** The RAG tetramer (blue) binds two RSSs (triangles) flanking the gene segments (rectangles) to form a synaptic complex (complex 2), within which cleavage takes place (complex 3). Subsequently, the RSS flanking regions (coding ends) are processed by non-homologous end joining (NHEJ) to yield a coding joint (product 5). RAG remains bound to the RSS ends (complex 4) with two possible outcomes: in V(D)J recombination, processing by NHEJ enzymes to yield a signal joint (product 6), or in transposition, the TIR end complex Inserts the mobile element into a new locus (product 7), generating a 5 bp target site duplication (TSD). **B) Taxonomic distribution of *RAG*/*RAGL* and *Transib* in eukaryotes**. The presence of *Transib* and/or *RAG*/*RAGL* families (A-D) in clades of eukaryotes is indicated (clades lacking *RAG/RAGL/Transib* elements are omitted). Newly identified *RAGL-A* elements in hemichordates and echinoderms are highlighted in magenta. **C) Genomic loci diagrams** of the most conserved copies of *RAGL-A* identified in hemichordates (left) and echinoderms (right). Transcriptomic support, whenever present, is mapped above gene diagrams with arrows. **D) PflRAG2L-A RT-PCR** illustrating PCR products of the size expected from spliced (blue) and unspliced mRNA, with mRNA samples from Taiwan *P. flava* of four developmental stages: unfertilized egg (UF), late blastula (LB), late gastrula (LG), and tornaria (T). Replicates and controls are shown in Figure S2. Diagram below illustrates the location of the PCR primers, the 9 aa contributed by exon 1 (blue), and an in frame stop codon upstream of exon 2 (red star).

Structures of the RAG1 core/RAG2 core tetramer bound to DNA or in apo form have provided numerous insights into the mechanism of DNA binding and cleavage (Chen et al. 2020a; Kim et al. 2018; Kim et al. 2015; Ru et al. 2015). DNA cleavage is performed by an RNaseH-fold DDE catalytic domain in the RAG1 core that shares structural similarity with the catalytic domains of cut and paste transposases and retroviral integrases (Kim et al. 2015; Montano and Rice 2011). The RAG1 core also contains the nonamer binding domain (NBD), which forms a dimer that binds the nonamers of the 12RSS and 23RSS in the synaptic complex. The NBD dimer is able to accommodate the length asymmetry of the 12/23RSS pair by pivoting on a flexible linker and is responsible for enforcing the 12/23 rule (Kim et al. 2018; Kim et al. 2015; Lapkouski et al. 2015; Ru et al. 2015). The RAG2 core is a 6-bladed kelch domain (Kim et al. 2015).

Many findings support the model that RAG evolved from a cut and paste transposon, including biochemical and structural similarities with transposases (Carmona and Schatz 2017; Liu et al. 2022) and the ability of RAG to perform transposition *in vitro* (Agrawal et al. 1998; Hiom et al. 1998). Transib mobile elements were the first to be suggested to share a common ancestor with RAG due to sequence similarities between Transib transposases and the catalytic core of RAG1 and between Transib terminal inverted repeats (TIRs) and RSSs (Kapitonov and Jurka 2005). Subsequent analyses revealed functional and structural similarities between RAG1 and the Transib transposase from the insect *Helicoverpa zea* (HzTransib) (Carmona and Schatz 2017; Hencken et al. 2012; Liu et al. 2019). Transib transposons do not, however, encode a protein resembling RAG2. The discovery of a transposon encoding RAG1-like (RAG1L) and RAG2-like (RAG2L) proteins in *Branchiostoma belcheri* (*Bbe*) provided a closer intermediate in evolution to jawed vertebrate RAG (Huang et al. 2016). The *BbeRAG1L/2L* gene pair preserves the convergent transcriptional orientation of jawed vertebrate *RAG1/2* and is flanked by TIRs with heptamer sequences that resemble those of RSSs and Transib TIRs. The BbeRAG1L/2L transposase cleaves DNA by the same nick/hairpin mechanism as RAG and HzTransib (Huang et al. 2016), closely resembles RAG structurally (Zhang et al. 2019), and generates a 5 bp target site duplication (TSD) during integration, as is the case for RAG and HzTransib (Hencken et al. 2012; Huang et al. 2016). *RAGL* transposons have now been found in numerous invertebrate clades - deuterostomes, protostomes, cnidarians, protists - some of which possess the full complement of expected transposon components: *TSD-TIR5’-RAG1L-RAG2L-TIR3’-TSD* (Martin et al. 2020; Morales Poole et al. 2017; Tao et al. 2022).

Four families of RAG/RAGL protein sequences have been identified (Martin et al. 2020; Morales Poole et al. 2017) (Figure 1B). Family A is represented by jawed vertebrate RAG and orphan RAG1L-A open reading frames in the hemichordate *Ptychodera flava* (acorn worm) while family B encompasses virtually all RAGL elements identified in invertebrates. Families C and D are minor variants restricted to one lineage or species. The finding of a divergent RAGL element with an atypical organization of its *RAG1L* and *RAG2L* genes in the protist *Aureococcus anophagefferens (Aan)* (Tao et al. 2022) suggests that additional families might be identified.

RAG appears to have undergone multiple adaptations during its evolution from transposase to recombinase, resulting in a tightly regulated complex whose enzymatic activities can be tolerated by the host. These adaptations, which include “coupled” cleavage of its two substrates, a requirement for asymmetric substrates (12/23 rule), and strong suppression of transposition activity *in vivo*, involved numerous, seemingly unrelated changes to the RAG1 and RAG2 proteins, arguing that RAG domestication occurred in multiple steps (Liu et al. 2022). The order and timing of these steps and whether they occurred before or after jawed vertebrate speciation is unknown.

A significant impediment to our understanding of RAG evolutionary history has been the large gap that exists between known RAGL transposases and RAG. Specifically, intact *RAGL-A* transposons have not been described and RAGL-B proteins exhibit multiple differences with RAG (Martin et al. 2020; Morales Poole et al. 2017). This latter point is illustrated by three adaptations that suppress transposition, arginine 848 in RAG1 and the AH and an inhibitory loop in RAG2, all of which have thus far been identified only in jawed vertebrates, leading to the model that they arose specifically in jawed vertebrates to protect against deleterious transposition events (Liu et al. 2022; Zhang et al. 2019; Zhang et al. 2020).

Using iterative search algorithms developed in our previous study (Martin et al. 2020), we have identified previously overlooked *RAG2L* open reading frames, revealing the first complete *RAGL* transposon of the A family in *P. flava* (*PflRAGL-A*) and nearly complete A family elements in several species of echinoderms: *Ophyothrix spiculata* (spiny brittle star) (*OspRAGL-A*), *Ophioderma brevispina* (green brittle star) (*ObrRAGL-A*) and *Marthasterias glacialis* (spiny starfish) (*MglRAGL-A*). The encoded RAGL-A proteins intermingle domains and sequence features of jawed vertebrate RAG and RAGL-B transposases while the *PflRAGL-A* TIRs combine features of RSSs and *RAGL-B* TIRs. Furthermore, unlike all previously described RAG2L proteins, both hemichordate and echinoderm RAG2L-A contain AHs, which we demonstrate are capable of potently suppressing RAG-mediated transposition. These findings demonstrate that the AH did not arise uniquely in jawed vertebrates and suggest that inhibition of transposition activity began to arise prior to jawed vertebrate speciation. The identified invertebrate RAGL-A element*s* help bridge the gap between *RAGL-B* and jawed vertebrate *RAG* and provide insight into the order and timing of events during RAG domestication.

## RESULTS

### Multiple complete RAGL-A transposon copies in two populations of *P. flava*

Using a sensitive iterative search strategy described previously (Martin et al. 2020), we performed searches for *RAG2L-A* sequences in two publicly available sequence scaffolds previously reported to contain *PflRAG1L-A* (Morales Poole et al. 2017) (see Materials and Methods). This led to the identification of two *RAG2L-A* sequences, 97% identical at the nucleotide level, encoding a 6-bladed kelch-like domain, an AH, and a PHD-like domain approximately 600 bp downstream of and in convergent orientation with *RAG1L_A*. Using these *RAG2L-A* sequences for further searches of the *P. flava* genome revealed two additional *RAG2L-A* loci (97-98% identity) for which no linked *RAG1L-A* counterpart could be found, although their location near scaffold boundaries precludes firm conclusions in this regard.

While the *RAG1L/2L-A* genomic loci thus identified all appeared to be pseudogenes containing frameshifts, searches of *P. flava* transcriptomic (TSA) data revealed mRNA sequences containing intact *RAG1L-A* open reading frames, several of which contained partial but intact *RAG2L-A* sequences on the reverse non-coding strand. This suggested the existence of at least one additional transposon copy, not covered by the public whole genome sequence (WGS) data, in the genome of *P. flava* (Hawaiian population; *P. flava^H^*), from which both the WGS and TSA data were derived. Indeed, targeted sequencing of *P. flava^H^* bacterial artificial chromosome (BAC) clones resulted in the identification of a *RAGL-A* transposon with the expected complete *TSD-TIR5’-RAG1L-RAG2L-TIR3’-TSD* configuration (Figure 1C, Supplemental File S1). We then searched for RAGL-A sequences in WGS data generated from *P. flava* from Taiwan (*P. flava^T^*), which identified four *RAGL-A* loci, one of which had a *TSD-TIR5’-RAG1L-RAG2L-TIR3’-TSD* configuration with intact *RAG1L-A* and *RAG2L-A* genes (Figure 1C). Notably, in both *P. flava^H^* and *P. flava^T^*, the various RAGL-A cassettes reside in different genomic locations and contain numerous unique mutations (Figure S1), arguing that *PflRAGL-A* transposition events continued to occur after the Hawaii and Taiwan populations split.

### A 5’ coding exon provides the *PflRAG2L-A* start codon

The *PflRAG2L-A* genomic loci and mRNA sequences from Hawaii and Taiwan populations encode kelch-AH-PHD configurations but all lacked an ATG start codon at the beginning of the kelch domain. The first in-frame ATG codon was within the second blade of the kelch domain and initiating protein synthesis at this site would almost certainly undermine the structural stability and functionality of the domain and would omit numerous upstream in-frame codons. Analysis of the region between the large RAG2L-A exon and the 3’ TIR revealed the presence of several potential mRNA splice donor/acceptor motifs, with one pair displaying good agreement with the canonical motifs. The possibility of mRNA splicing in this region was investigated by reverse transcription combined with PCR (RT-PCR) using *P. flava^T^* RNA samples purified from four development stages: unfertilized eggs, late blastula, late gastrula, and tornaria (a larval stage). A PCR product consistent with the predicted spliced mRNA was consistently detected in late blastula, late gastrula, and tornaria stages and was undetectable in unfertilized eggs (Figure 1D; all three biological replicates shown in Figure S2A). Control reactions showed that detection of the splice product was dependent on reverse transcription and that genomic DNA contamination was present in some samples, explaining strong signals seen for the unspliced product in some reactions (Figure S2B). mRNA quality was verified by amplification of *Vasa* using intron-spanning primers, confirming the presence of intact mRNA in unfertilized egg samples (Figure S2C) and supporting the conclusions that maternal *PflRAG2L-A* transcripts are absent in the unfertilized egg and that expression is induced during early development. Sequencing of the *PflRAG2L-A^T^* spliced PCR product revealed the predicted splice junction and confirmed the presence of an upstream exon capable of adding 9 aa to the N-terminus of the protein (Figure S2D). This upstream exon and the splice donor and acceptor sites are conserved in the intact *RAGL-A* transposon from *P. flava^H^* (Supplemental File S1). We conclude that *PflRAG2L-A* is induced and undergoes mRNA splicing during early development and that an upstream exon encodes the N-terminal residues of the PflRAG2L-A protein.

### The *RAGL-A* family is also present in two echinoderm classes

Public WGS and TSA database screening led to the identification of additional *RAG1L/2L-A* gene pairs in another invertebrate phylum – echinoderms – in both the Ophiuroidea class (spiny brittle star *Ophiothrix spiculata* (*Osp*) and green brittle star *Ophioderma brevispina* (*Obr*)) and the Asteroidea class (spiny starfish *Marthasterias glacialis* (*Mgl*)) (Figure 1C). The three echinoderm *RAG1L-A* genes encode complete and potentially functional endonucleases while a complete RAG2L-A counterpart was found only in *O. spiculata* and *O. brevispina* (the identified *MglRAG2L-A* locus lacks coding information for a portion of the RAG2L N-terminus). Multiple putative *RAGL-A* loci were identified in another Asteroidea member (spiny starfish *Zoroaster sp*.), but all appear to be degraded, non-functional loci and were not analyzed further (Supplemental File S1).

All identified echinoderm *RAG2L-A* genes are found downstream of *RAG1L-A* in the expected reverse orientation (Figure 1C) and encode RAG2L-A proteins containing an AH between kelch and PHD domains, as in PflRAG2L-A. *OspRAG1L-A* (but not *OspRAG2L-A*) was also recently reported by Tao et al. (Tao et al. 2022). The WGS database was also found to contain two additional *OspRAG2L-A* loci (OspRAG2L-A.2, OspRAG2L-A.3) that are apparently unlinked to a *RAGL1_A* counterpart and that encode proteins with a complete kelch-AH-PHD configuration (Supplemental File S1). An additional *RAGL* locus was identified in *O. spiculata* in which *RAG2L* contains an AH but both RAG1L and RAG2L appear to have been pseudogenized (data not shown).

### Phylogenetic analysis: RAGL-A clusters with jawed vertebrate RAG

Phylogenetic analyses based on RAG1(L) catalytic core sequences found that RAG1L-A clusters with jawed vertebrate RAG1 rather than with RAG1L-B (Figure 2A, Figure S3A, B), consistent with previous studies (Martin et al. 2020; Morales Poole et al. 2017; Tao et al. 2022). Bootstrap support of the RAG1L phylogenetic tree indicates a statistically significant separation between the RAG1L-A and -B families and RAG1L-A catalytic core sequences show greater sequence identity with RAG1 (35-39%) than with RAG1L-B (26-33%) (Figure S4A). Notably, echinoderm RAG1L-A sequences consistently form an outgroup to hemichordate and jawed vertebrate RAG1L-A (Figure 2A, Figure S3A, B), with potential implications for the origin of jawed vertebrate RAG (see Discussion).

**Figure 2.**
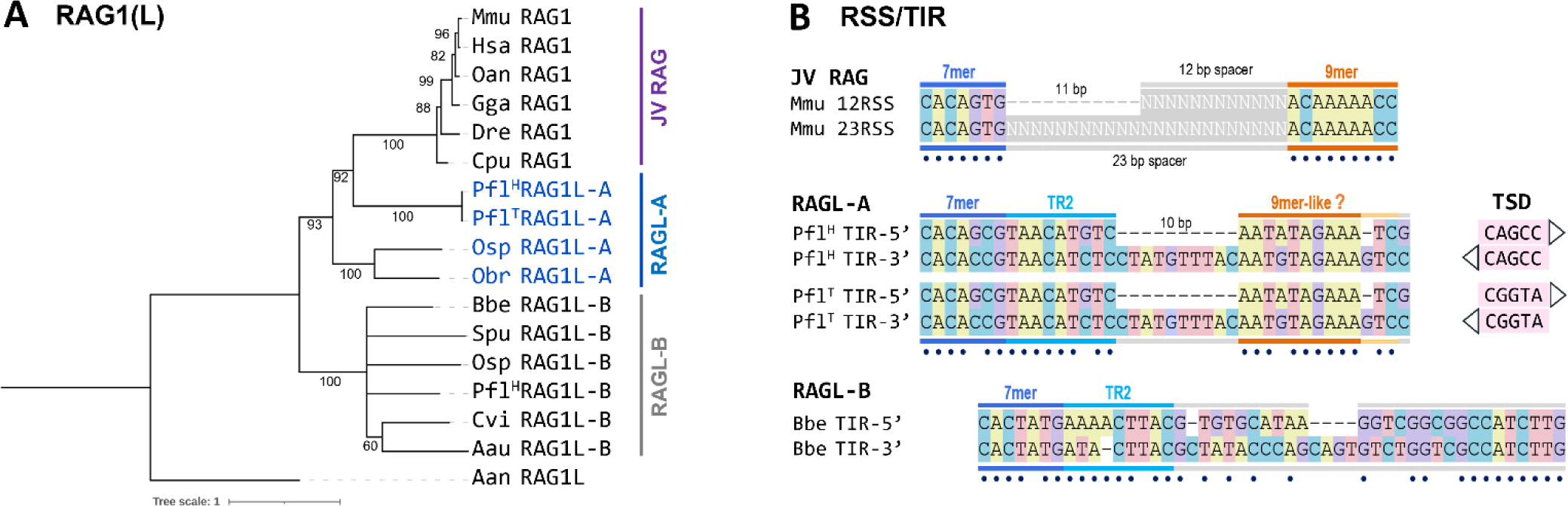
Phylogeny and TIRs of *RAGL-A* elements. **A)** Phylogeny trees of RAG1/RAG1L computed using Maximum Likelihood method using Iqtree as described in Methods. Branch support (1000 UFBoot replicates) is indicated next to each branch. **B)** Comparison of selected RSSs and TIRs. Heptamer, TR2, spacer, and nonamer regions are indicated. Matching nucleotide pairs are depicted with dots. Target site duplication (TSD) pairs are shown next to the *PflRAGL-A* TIR sequences.

RAG2(L) phylogenetic trees (computed using blades 2-5 of the RAG2(L) kelch domain) are less reliable than those for RAG1L with lower bootstrap support at many branches (Figure S3C, D), as expected given the weaker conservation of RAG2(L) compared to RAG1(L) (Figure S4) (Martin et al. 2020; Morales Poole et al. 2017). RAG2L trees maintain the same overall structure as for RAG1(L) (Figure S3C, D). The orphan OspRAG2L-A.2 and .3 sequences group well with OspRAG2L-A.1 (which is paired with OspRAG1L-A) (Figure S3) but the three sequences share only 42-52% identity in their most conserved region (Kelch domain blades 2-5). This high divergence might be due to either the existence of different A subfamilies or an increased rate of change in the loci hosting the isolated *RAG2L-A* genes.

Notably, both the *P. flava* and *O. spiculata* genomes also contain *RAGL-B* transposons (Supplemental File S1), which encode RAG1L-B proteins with 42-45% identity in the catalytic core region with RAGL-B in other species, but lower identity (30-31%) with their intraspecies RAGL-A counterparts (Figure S4). This argues that the *RAGL-A* and *RAGL-B* transposon lineages evolved independently and in parallel in both the *P. flava* and *O. spiculata* genomes.

### *PflRAGL-A* TIRs are chimeras of RSSs and *RAGL-B* TIRs

The TIRs elements of *PflRAGL-A^H^* and *PflRAGL-A^T^*are identical, indicative of the strong conservation of *RAGL-A* in the two populations, and exhibit a mixture of features of jawed vertebrate RSSs and invertebrate RAGL-B TIRs (Figure 2B and Supplemental File S1). Like 12/23RSSs, the 5’ and 3’ TIRs contain a conserved heptamer closely resembling that of the consensus RSS separated from an AT-rich nonamer-like sequence by a “spacer” region with a 10 bp length asymmetry. Such asymmetry is not observed in BbeRAGL-B TIRs, which also lack an AT-rich nonamer-like sequence (Figure 2B). Unlike 12/23RSSs, however, the “spacer” regions of the *PflRAGL-A* 5’ and 3’ TIRs are highly conserved for their first 9 bp, and in addition, the conserved region begins with a sequence rich in adenines. This conserved 9-10 bp region immediately adjacent to the heptamer, referred to as TIR region 2 (TR2), is observed in many deuterostome and protostome RAGL-B TIRs (Martin et al. 2020) and is important for DNA cleavage by BbeRAGL-B (Zhang et al. 2019). *PflRAGL-A* TIRs therefore display a hybrid heptamer-TR2-asymmetric spacer-nonamer organization.

The TIRs of several PflRAGL-A cassettes are flanked by 5 bp TSDs and these TSDs differ in sequence between *P. flava^H^*and *P. flava^T^* (Figure 2B, S1), further supporting ongoing *RAGL-A* transposition activity in the *P. flava* genome after the divergence of the Hawaiian and Taiwanese populations.

In *O. spiculata,* the single *RAG1/2L-A* gene pair identified was located close (within 400 bp) to one end of the sequence scaffold, preventing identification and proper validation of TIR elements, in contrast to *P. flava* where multiple transposon copies could be used to validate the transposon cassette margins. Similarly, no TIR elements were identified in the other echinoderm species despite the fact that for several loci, substantial amounts of genomic DNA flanking the *RAG1L* and *RAG2L* genes is available.

### RAG1L-A proteins display a mixture of invertebrate RAG1L-B and jawed vertebrate RAG1 traits

In *P. flava*, *O. spiculata*, *O. brevispina*, and *M. glacialis*, RAG1L-A protein sequences display all known essential catalytic core domain components, including the catalytic DEDE tetrad and four Zn-coordinating residues that orchestrate folding of a zinc binding domain (ZnB) that makes up much of the C-terminal portion of the catalytic core (Figure 3A, B, and Supplemental File S2). The proteins also contain three conserved cysteine-rich motifs (C1, C2, and C3) found in the RAG1 N-terminal region and all except OspRAG1L-A possess a RING-ZnF domain. Loss of the RING-ZnF domain has previously been reported in *S. purpuratus* RAG1L-B and in several protostome RAGL-B proteins (Fugmann et al. 2006; Martin et al. 2020).

**Figure 3.**
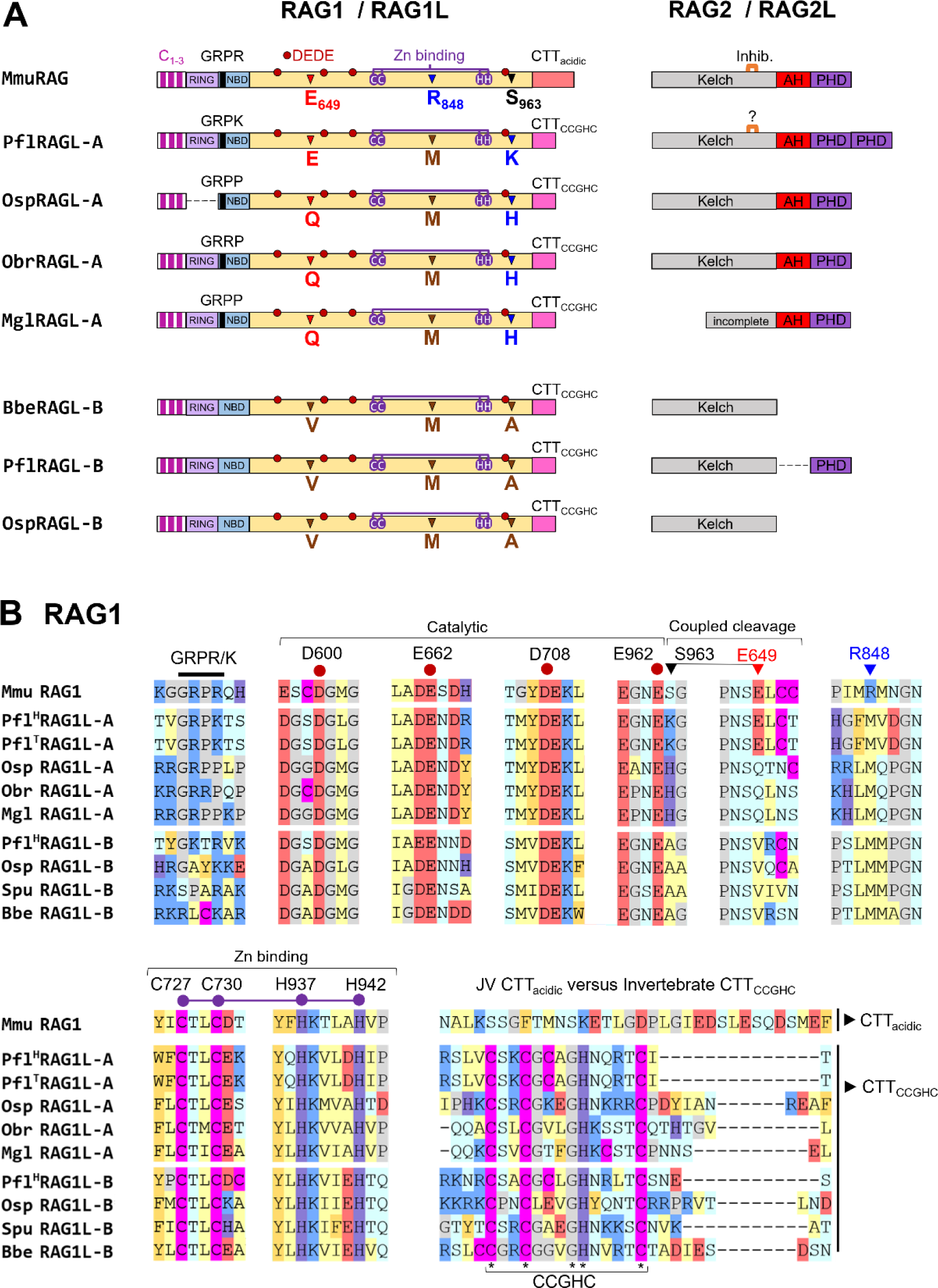
RAG1L-A and RAG2L-A proteins. **A)** Domain organization of RAG1(L) – RAG2(L) pairs from the most preserved copies of *RAGL-A* and *RAGL-B* from *P. flava, O. spiculata*, *O. brevispina,* and *M.* glacialis, mouse (Mmu) RAG, *B. belcheri*. The RAG1(L) conserved catalytic DEDE tetrad and Zinc binding residues are depicted with red and purple circles. Regulatory adaptations E649, R848, and S963 identified in mouse RAG1 are indicated with triangles. Kelch, RAG2 kelch domain encoding blades 1-6; AH, acidic hinge; PHD, plant homeodomain; Inhib, B6-C6 transposition inhibitory loop in RAG2. **B)** Sequence conservation and variation at key positions and motifs in RAG1(L) proteins.

In contrast to RAG1L-B proteins, *P. flava* RAG1L-A exhibits two features characteristic of jawed vertebrate RAG1. The first is an NBD containing a GRPR/K motif (hereafter, NBD_GRPR/K_) (Figure 3A, B). In RAG1, this motif makes direct contact with the A/T-rich portion of the nonamer and is required for RAG cleavage activity (Difilippantonio et al. 1996; Schatz and Swanson 2011; Spanopoulou et al. 1996; Yin et al. 2009). The presence of NBD_GRPR/K_ in PflRAG1L-A together with asymmetric TIRs containing an A/T-rich nonamer-like sequence is consistent with the possibility that PflRAGL-A mediates nonamer recognition by a mechanism similar to that of RAG. In echinoderm RAG1L-A, the corresponding sequence is GRP<u>P</u> or GRR<u>P</u>, whose effect on DNA binding activity is difficult to predict and, in the case of GRPP, might compromise binding due to loss of electrostatic interactions between the R/K residue and the DNA backbone (Yin et al. 2009). *BbeRAGL* TIR elements are almost symmetrical in length and lack an adenine-rich nonamer region, which is mirrored by the fact that the BbeRAG1L NBD-equivalent domain lacks the GRPR/K motif and makes only a modest contribution to cleavage activity (Zhang et al. 2019).

The second feature shared uniquely between PflRAG1L-A and RAG1 is glutamate at the position corresponding to mouse RAG1 E649 (Figure 3A, B). DNA cleavage by RAG occurs in a synchronous, or “coupled” fashion and only when both of its substrates are bound in the synaptic complex. The enforcement of coupled cleavage is dependent on residue E649, which is thought to exert its influence in part through hydrogen bond formation with RAG1 S963 (Kriatchko et al. 2006; Zhang et al. 2019). The E649/S963 pair, highly conserved in jawed vertebrate RAG1, is E/K in PflRAG1L-A and Q/H in Osp, Obr, and Mgl RAG1L-A proteins (Figure 3B). These residues possess bulky side chains that could engage in hydrogen bonds and interact electrostatically, raising the possibility that PflRAGL-A and echinoderm RAGL-A possess some degree of coupled cleavage.

RAG1 E649 also suppresses RAG-mediated transposition modestly (approx. 2-fold) in a manner that is independent of S963 (Zhang et al. 2019), and transposition by PflRAGL-A and echinoderm RAGL-A might similarly be modestly downregulated by the E and Q residues, respectively, they possess at this position (Q shares many physiochemical properties with E). A second highly conserved residue in RAG1, arginine 848, also contributes to transposition suppression, but in this case, almost completely eliminates transposition activity (Zhang et al. 2019). This residue is a hydrophobic aa, most often methionine, in invertebrate RAG1L proteins, and all identified RAG1L-A proteins retain the transposition-permissive M at this position (Figure 3A, B).

Interestingly, the C-terminal tails (CTTs) of the hemichordate and echinoderm PflRAG1L-A proteins possess a conserved <u>C</u>X_2_<u>C</u>X_3_<u>GH</u>X_4_<u>C</u> motif (CCGHC motif hereafter) (Figure 3A, B), found in nucleic acid-binding “zinc-knuckle” domains (De Guzman et al. 1998; Klein et al. 2000). The CCGHC motif is present in the CTTs of virtually all invertebrate RAG1L proteins but not jawed vertebrate RAG1 (where CTT is acidic). In BbeRAG1L-B, CTT_CCGHC_ is a DNA binding “clamp” that interacts with TR2 and is critical for cleavage activity (Zhang et al. 2019). The presence of a GRPR/K-containing NBD, asymmetric TIRs with an AT-rich nonamer-like region, and CTT_CCGHC_ in PflRAG1L-A suggests competing, and potentially redundant, modes of DNA binding (see Discussion).

We used artificial intelligence based methods (AlphaFold, OmegaFold) and homology 3D modelling pipelines to compute predictive structural models of the core regions of PflRAG1L-A and OspRAG1L-A, revealing that they can adopt structures similar to those of RAG1 and BbeRAG1L-B and that their predicted DNA binding surfaces exhibit strikingly similar charge distributions to those of RAG1/BbeRAG1L-B (Figure S5). This observation suggests substantial parallels between the mechanisms of DNA engagement by RAG1L-A proteins and that by RAG1 and BbeRAG1L-B.

In summary, PflRAG1L-A and Osp/Obr/MglRAG1L-A are potentially catalytically active proteins possessing features distinctive of jawed vertebrate RAG1 (GRPR/K or related motif, E/Q649) as well as features found previously only in invertebrate RAG1L-B (M848, CTT_CCGHC_).

### RAG2L-A proteins contain an acidic hinge capable of suppressing transposition

We previously reported that sequence conservation is unevenly distributed in RAG2L-B Kelch domains, with higher levels (∼25% identity) in blades 2-5 and lower levels in blades 1 and 6, with blades 1 and 6 often exhibiting an atypical GG motif (a glycine doublet that is characteristic of beta strand B) (Martin et al. 2020). The same trend is observed in RAG2L-A sequences where only predicted blades 2-4/5 display the canonical A-B-C-D beta sheet structure (Supplementary File S2B). However, RAG2L-A proteins diverge from RAG2L-B and resemble jawed vertebrate RAG2 proteins in that they lack the basic amino acid patch on the large loop connecting beta strands D6 and A1, seen clearly in BbeRAG2L-B (Supplementary File S2B).

The structural effects of the sequence differences in blades 1 and 6 are particularly clear when examining blade 6 in the cryo-electron microscopy (cryo-EM) structures of BbeRAG2L-B and RAG2. In jawed vertebrate RAG2, the loop between β-strands B6 and C6 is longer than in RAG2L-B proteins (Figure 4A) and the B6 GG motif and B6-C6 loop are shifted out of the plane defined by the other 5 blades; this is not the case in BbeRAGL2-B (Figure 4B). The protruding B6-C6 loop of RAG2 is able to extend into a deep pocket and contact target site DNA in the RAG target capture and strand transfer complexes (Figure 4C) (Chen et al. 2020b; Zhang et al. 2020). Deletion of four amino acids at the loop tip increases RAG-mediated transposition 2-3 fold and it has been suggested that the loop sterically interferes with target DNA capture (Zhang et al. 2020). Notably, while this inhibitory loop is only 5 aa long in BbeRAG2L-B and other invertebrate RAG2L-B proteins, it is 8 aa long in PflRAG2L-A, nearly as long as the 10 aa loop in mouse RAG2 (Figure 4A). In contrast, the echinoderm Osp/ObrRAG2L-A sequences exhibit a short 5 aa B6-C6 loop (Figure 4A).

**Figure 4.**
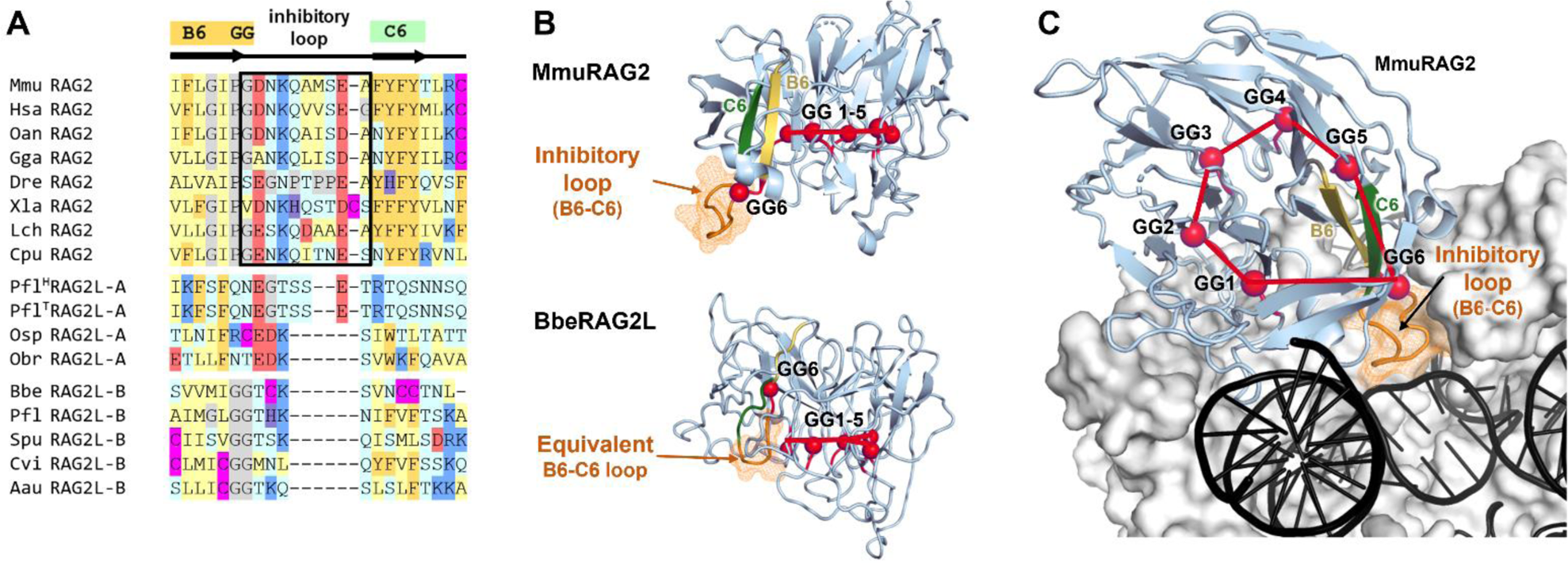
RAG2 kelch blade 6 transposition inhibitory loop. **A)** Sequence of jawed vertebrate RAG2 loop between β-strands B6 (yellow) and C6 (green) compared to the equivalent region in RAG2L-A and RAG2L-B proteins. GG, a double glycine motif (sometimes PG) frequently present at the end of β-strand B in kelch domain blades. **B)** Structural differences in blade 6 as observed in the cryo-EM structures of mouse RAG2 and BbeRAG2L. The GG motif from blade 6 (GG6) shifts in opposite directions in the two proteins with respect to the plane generated by the rest of the GG motifs (GG1-5) (PDB: 6XNY, 6B40). **C)** Top view of mouse RAG strand transfer complex illustrating mouse RAG2 structure (blue ribbon model) and the downward projection of the sixth blade and the B6-C6 loop (orange mesh representation) to make contact with target DNA (black). RAG1 in gray. (PDB: 6XNY).

A unique feature of RAG2L-A sequences, not observed in any other invertebrate RAG2L protein reported thus far, is an acidic region between the kelch and PHD domains. The RAG2L-A AH is somewhat longer (66-78 aa) than that of jawed vertebrate RAG2 (52-62 aa) and has nearly as high a density of acidic residues (34-40%) as in jawed vertebrate RAG2 AHs (39-44%) (Figure 5A). The distribution of D/E residues shows some similarities between the different AHs, including a region of high D/E density near the AH C-terminus, but other sequence similarities are not apparent.

**Figure 5.**
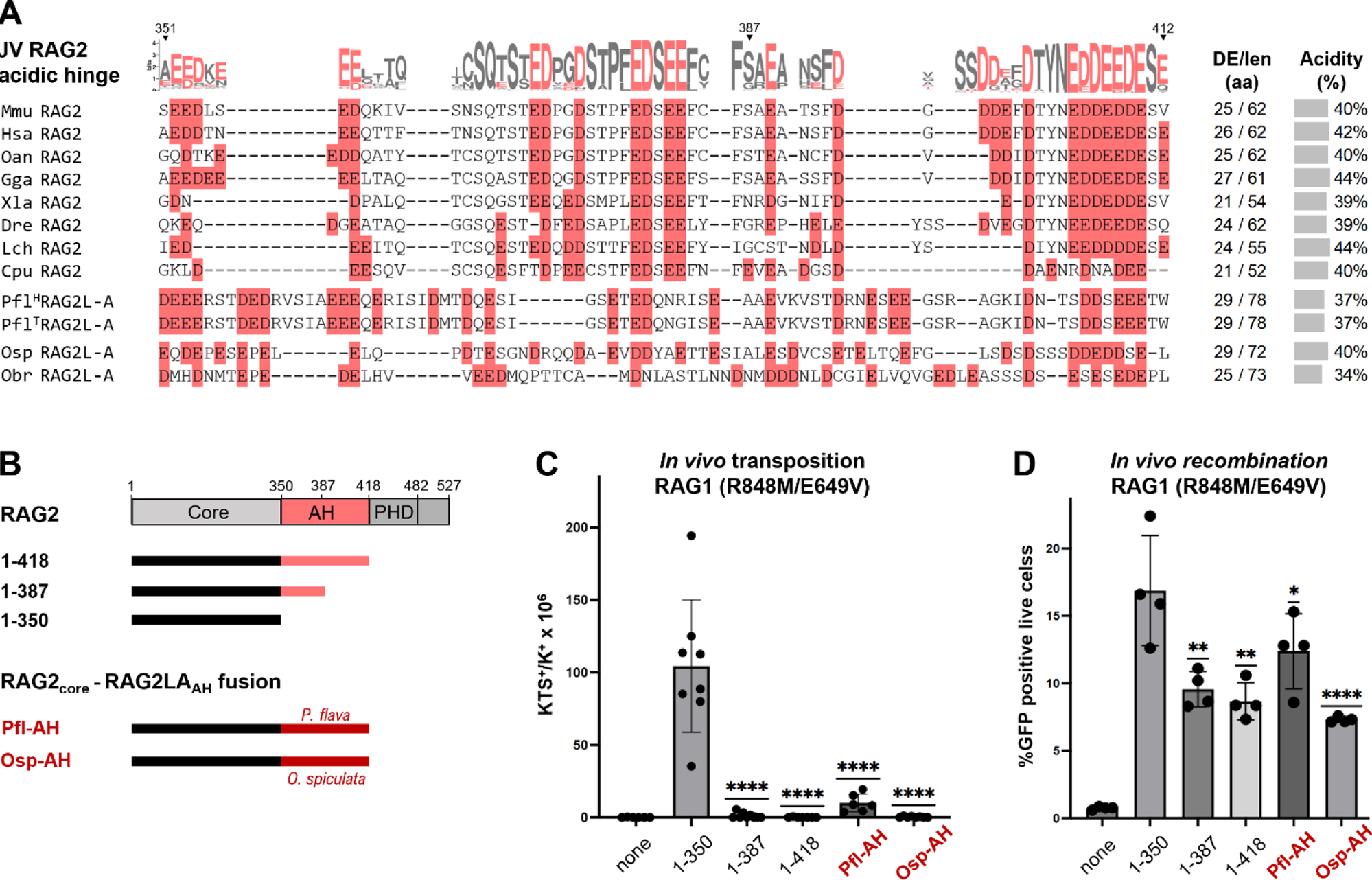
RAG2L-A acidic hinge sequence and transposition inhibitory activity. **A) Comparison of the acidic hinge (AH) of jawed vertebrate (JV) RAG2 and RAG2L-A sequences** in hemichordates (*P. flava*) and echinoderms (*O. spiculata* and *O. brevispina*), with acidic residues Asp and Glu highlighted in red. Overall length, number of acidic residues, and percent acidic residues shown at right. Conservation profile of jawed vertebrate RAG2 was computed as KL divergence as described in Methods (letter height is proportional to conservation). **B) Schematic diagrams of RAG2 proteins tested for activity.** Mouse RAG2 is shown at top. Mouse RAG2 core (black line) alone or fused to either its own AH or that of PflRAG2L-A or OspRAG2L-A were tested. **C, D) *In vivo* transposition (C) and recombination (D)** assays performed in human 293 cells upon expression of full length mouse RAG1 containing transposition activating mutations R848M and E649V and the indicated RAG2 fusion proteins (diagrammed in panel B). Data points are biological replicates derived from independent experiments. Statistically significance calculated compared to RAG2 1-350 using two-tailed T test (P<0.05 (*), <0.01 (**), <0.0001 (****).

The mouse and human RAG2 AHs suppress transposition potently (>50 fold) (Zhang et al. 2019). To test whether invertebrate RAG2L-A AH sequences also possess transposition inhibitory activity, we appended the PflRAG2L-A or OspRAG2L-A.1 AH to the mouse RAG2 core region (Figure 5B) and assayed the resulting chimeric proteins for transposition activity together with mouse RAG1 bearing transposition activating mutations R848M and E649V. The results demonstrate that the *P. flava* and *O. spiculata* AH regions potently suppress transposition activity, to an extent equivalent or nearly equivalent to that observed with a partial (aa 351-387) or full (aa 351-418) mouse AH (Figure 5C). An assay for recombination confirmed that all of the RAG2 fusion proteins tested support DNA cleavage, though as expected (Lu et al. 2015; Zhang et al. 2019), appending any AH region reduced recombination somewhat (Figure 5D). We conclude that AH domains with transposition suppressive potential exist in RAG2L-A proteins in hemichordates and echinoderms and that the AH is not a jawed vertebrate-specific adaptation.

### PflRAG2L-A encodes a double PHD

The jawed vertebrate RAG2 PHD contains two zinc fingers that create a pocket that binds the N-terminal tail of histone 3 when lysine 4 is trimethylated (H3K4me3) (Liu et al. 2007; Matthews et al. 2007). Pfl, Osp, and ObrRAG2L-A sequences contain a PHD abutting the AH (hereafter PHD_1_) in which the zinc coordinating cysteine and histidine residues are readily identifiable, with the exception of OspRAG2L-A.3 which lacks the final two cysteine residues. The proteins also contain a highly conserved tryptophan residue (W453 in mouse RAG2) required for methylated lysine binding (Matthews et al. 2007) (Figure 6A, Supplemental File S2b). Despite this similarity with jawed vertebrate PHD_1_, RAG2L-A PHD_1_ more closely resembles PHD_1_ of RAG2L-B in possessing an additional conserved cysteine residue (* in Figure 6A) and aa changes at conserved methyl-lysine-binding residues of jawed vertebrate PHD_1_ (Y415→W or L and M443→W (Ramon-Maiques et al. 2007) (Figure 6A). Hence, the histone recognition profile of RAG2L-A PHD_1_ might differ from that of vertebrate RAG2 PHD_1_, consistent with the finding that PHD_1_ of SpuRAG2L-B preferentially binds H3K4me2 instead of H3K4me3 (Wilson et al. 2008).

**Figure 6.**
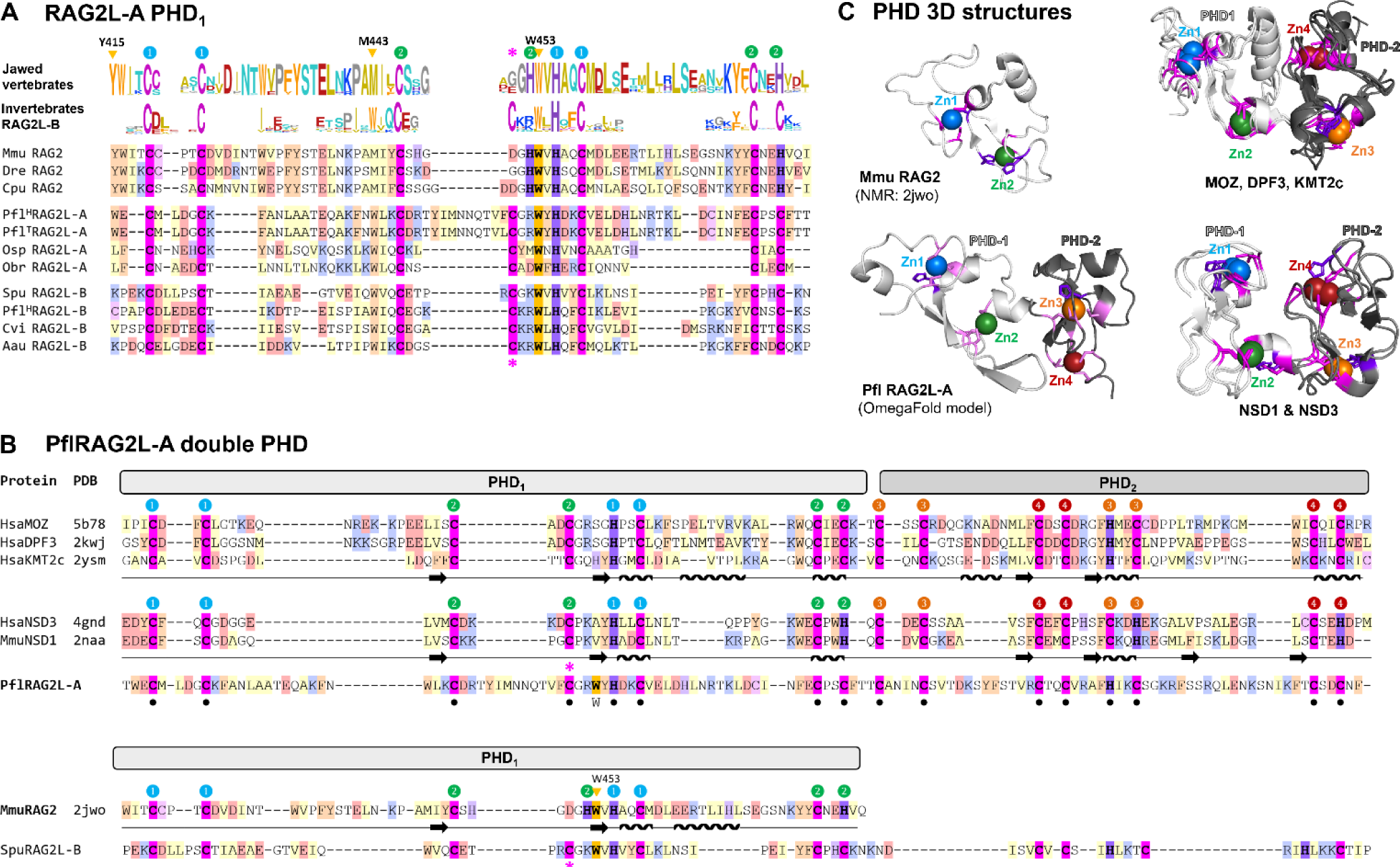
RAG2L-A PHD region. A) Comparison of PHD finger 1 (PHD_1_) of RAG2L-A with other invertebrate RAG2L and jawed vertebrate RAG2 proteins. The Cys/His pattern is highlighted in magenta and purple, respectively with numbers indicating residues that coordinate zinc atoms 1 and 2. Orange triangles, conserved residues in jawed vertebrate RAG2 important for binding of H3K4me3; pink asterisk, Cys residue conserved in RAG2L but not RAG2 proteins. Sequence conservation profile displayed above the alignment was computed as described in Figure 5. B) Sequence comparison of the double PHD region (PHD_1_ and PHD_2_) of PflRAG2-A with two groups of double PHD domain proteins for which experimental structures are available and which display similar Cys/His patterns. Colored labels 1-4 indicate the Zn binding topology. The mouse single PHD and the incomplete extended PHD pattern of *S. purpuratus* RAG2L-B are shown below for comparison. Secondary structure elements indicated by the experimental 3D structures are shown below the amino acid sequence (arrow, β-strand, wavy line, α-helix). C) **Structural comparison of double PHD domains.** 3D structures of mouse RAG2 single PHD (upper left, NMR structure), *P. flava* RAG2L-A double PHD (lower left, predicted model using OmegaFold), superimposition of double PHD domains of MOZ, DPF3 and KMT2c (upper right; PDB: 5B78, 2KWJ, 2YSM, respectively) and NSD1 and NSD3 (lower right; PDB: 4GDN, 2NAA, respectively). Zinc-binding Cys/His residues are displayed in magenta/purple, while Zn ions are colored as in labels in panels A and B.

Unlike any RAG2(L) protein described to date, PflRAG2L-A contains a second complete PHD (hereafter, PHD_2_) immediately following PHD_1_, that, like PHD_1_, can encode CCHC and CCCC zinc fingers (fingers 3 and 4 in Figure 6B). The double PHD_1_-PHD_2_ domain of PflRAG2L-A has limited sequence identity with two groups of structurally related double PHDs: 1) the double PHDs of histone acetyltransferase MOZ, chromatin remodeling complex subunit DPF3, and histone-lysine N-methyltransferase KMT2C (Xiong et al. 2016; Zeng et al. 2010), and 2) the double PHDs of the histone-lysine N-methyltransferases NSD1 and NSD3 (Berardi et al. 2016; He et al. 2013) (Figure 6B, C). While sequence homology between the two double PHD groups is largely limited to their zinc-coordinating C/H residues, their 3D architectures bear striking similarities, including their 4 zinc-binding pockets with similar C/H patterns (Figure 6B, C). The C/H pattern in the double PHD of PflRAG2L-A precisely matches that of the MOZ/DPF3/KMT2C group, and it is plausible that it adopts a similar structure. We note that SpuRAG2L-B might contain a degenerate PHD_2_ (Figure 6B).

## DISCUSSION

The absence of identified *RAGL-A* transposons had left a substantial gap in our understanding of the evolutionary history of jawed vertebrate RAG, in particular, numerous uncertainties as to the order in which various domains/residues were gained and lost and whether key functional adaptations occurred prior to or after jawed vertebrate speciation. Our identification of complete *RAGL-A* elements in hemichordates and echinoderms provides for clarification of these issues and allows for a more nuanced description of the evolution of regulated DNA binding, DNA cleavage, and transposition activities of the RAG recombinase.

The *RAGL-A* open reading frames in *P. flava*, *O. spiculata*, and *O. brevispina* are intact and encode predicted RAG1L-A and RAG2L-A proteins that contain all of the domains and catalytic and structural elements predicted to be required for catalytic activity. It is therefore plausible that these elements encode active DNA endonucleases, and in the case of *P. flava*, that the elements flanked by TIRs remain active transposons. Recent transposition activity of *PflRAGL-A* is supported by the presence of multiple *RAGL-A* copies (one fully intact, several exhibiting pseudogenization) in both the Hawaiian and Taiwanese populations of *P. flava* and by the different TSDs and chromosome regions that flank them.

While additional analyses might reveal TIRs flanking echinoderm *RAGL-A* elements, their apparent absence parallels the structure of the first *RAG1L*-*RAG2L* gene pair to be identified in invertebrates (in the purple sea urchin *S. purpuratus*) (Fugmann et al. 2006). Indeed, to our knowledge, no intact *RAGL* element predicted to be capable of transposition has been identified in echinoderms despite the identification of *RAG1L*-*RAG2L* loci in multiple echinoderm species (this report and (Fugmann 2010; Kapitonov and Koonin 2015; Martin et al. 2020; Morales Poole et al. 2017; Tao et al. 2022; Yakovenko et al. 2022)), and despite the identification of potentially active *RAGL* transposons in multiple lineages of deuterostomes, protostomes, and cnidarians, and even in a protist (Huang et al. 2016; Martin et al. 2020; Morales Poole et al. 2017; Tao et al. 2022). In contrast, *Transib* transposons with apparently intact TIRs are present in numerous species of echinoderms (our unpublished data and (Kapitonov and Jurka 2005; Kapitonov and Koonin 2015; Tao et al. 2022)), suggestive of different selective pressures acting on *Transib* and *RAGL* transposons in this clade. Our findings indicate that *RAG1L/2L-A* loci might represent domesticated transposases performing novel functions for their echinoderm hosts, as has been proposed for *RAGL-B* loci in sea urchins (Fugmann et al. 2006; Yakovenko et al. 2022). Our findings also argue that *RAGL-A* and *RAGL-B* transposons evolved side-by-side in both hemichordates and echinoderms over extended evolutionary periods.

A particularly striking feature of the *RAGL-A* elements reported here is their chimeric nature, with the RAGL-A proteins and their TIRs exhibiting features distinctive of jawed vertebrate RAG/RSSs and of RAGL-B proteins/TIRs (summarized in Figure 7A). Phylogenetic analyses demonstrate that invertebrate RAGL-A proteins are more closely related to jawed vertebrate RAG than are RAGL-B proteins (Figure 2A and Figure S4). Our findings have implications for the evolution of RAG’s RSS substrates and its DNA binding, DNA cleavage, and transposition activities.

**Figure 7.**
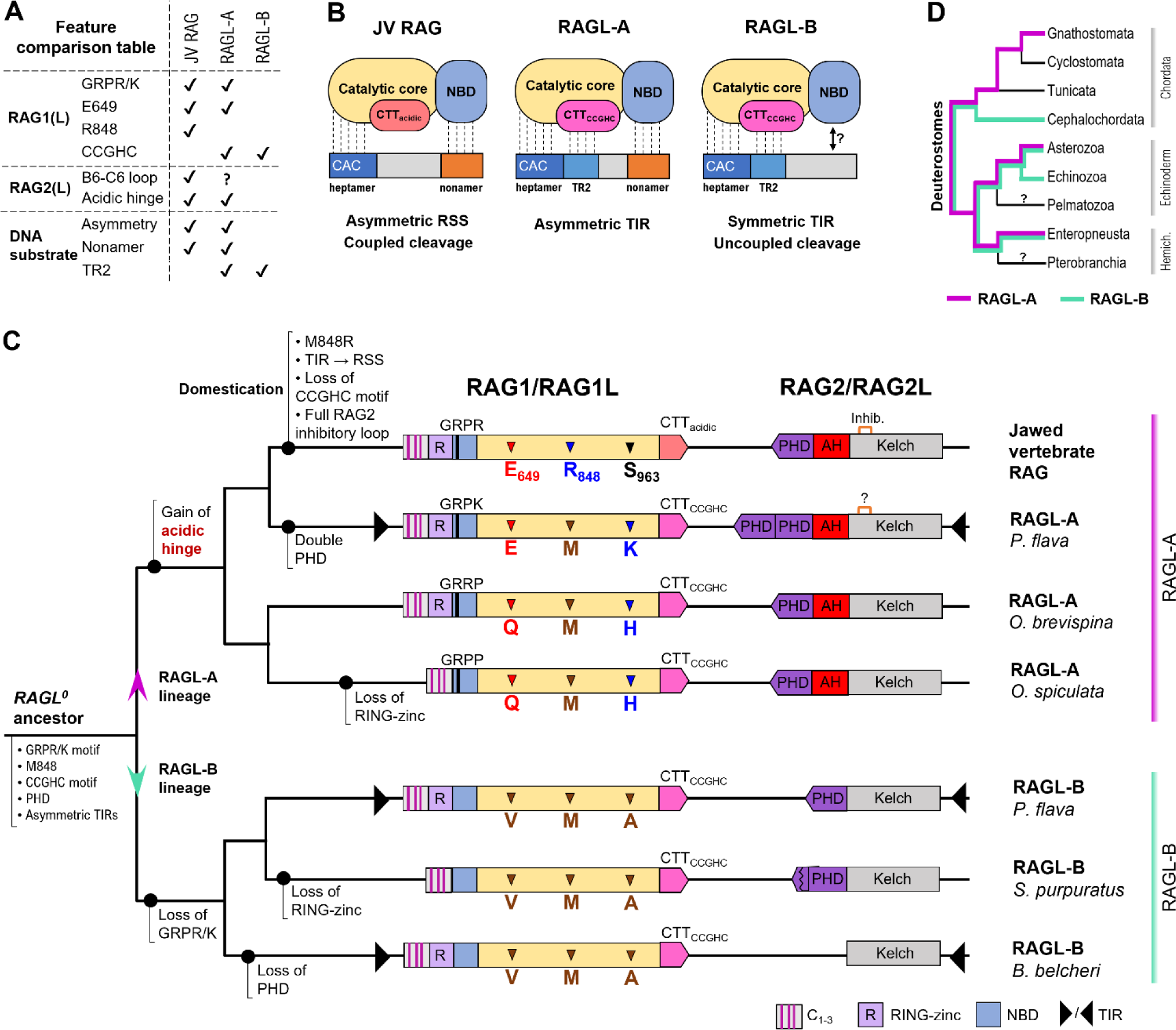
Features, DNA binding modes, and model for evolution of RAG(L) proteins. **A) Feature comparison table** of functionally important elements of the different RAG(L) lineage proteins and DNA substrates. **B) Differences in DNA binding modes** between jawed vertebrate RAG and BbeRAGL-B and hypothesized DNA binding modes available to RAGL-A proteins. **C) Model for the evolution of RAGL-A, RAGL-B, and RAG.** Beginning with *RAGL^0^*, the presumed ancestral *RAGL* transposon, evolutionary events leading to the gain or loss of traits and changes in domain architecture are depicted. See text for additional details. RAG1(L) and RAG2(L) proteins for the indicated species are diagrammed in the tail-to-tail configuration observed for their respective genes. C_1-3_, domain containing three cysteine pairs; R, RING-zinc finger domain; GRPR/K, motif involved in DNA binding at the N-terminus of the NBD; CTT_acidic_, acidic C-terminal tail; CTT_CCGHC_, C-terminal tail containing DNA binding domain with conserved CCGHC residues; Kelch, kelch domain constituting RAG2 core; AH, acidic hinge; PHD, plant homeodomain; black triangles, TIRs. For *ObrRAGL-A* and *OspRAGL-A*, TIRs have not yet been identified and hence are not shown, though for *OspRAGL-A*, the available DNA sequence assembly does not allow a definitive conclusion on this issue. Tree structure reflects the evolution of protein domains and features and is not meant to represent species phylogeny. **D) Schematic evolutionary tree of deuterostomes** depicting the branches where *RAGL-A* (magenta) and *RAGL-B* (green) elements are found. Length of tree branches does not indicate degree of relatedness.

### Evolution of RAG DNA binding and RSS substrates

RAG and BbeRAGL-B bind substrate DNA through distinct modes: while RAG is completely dependent on the RAG1 NBD for activity and CTT_acidic_ is dispensable, BbeRAGL relies heavily on BbeRAG1L CTT_CCGHC_ for activity and its NBD (which lacks a GRPR/K motif) makes only a minor contribution (Zhang et al. 2019) (Figure 7B). To our knowledge, PflRAG1L-A is the first RAG1L protein to be described that contains both NBD_GRPR/K_ and CTT_CCGHC_. As a result, it would be predicted to be capable of both modes of DNA engagement (Figure 7B), a conclusion supported by the presence of conserved TR2 and AT-rich nonamer-like conserved sequence in its TIRs. It is unclear whether one or the other of these two strong DNA binding modes might be dominant. Biochemical experiments with chimeric RAG1-BbeRAG1L proteins provide some insight into these issues (Zhang et al. 2019): i) attaching NBD_GRPR/K_ to BbeRAG1L creates a BbeRAGL enzyme that is no longer dependent on CTT_CCGHC_ and TR2 for cleavage activity but which now requires a nonamer spaced 12 or 23 bp from the heptamer, and reciprocally, ii) attaching CTT_CCGHC_ to RAG1 creates a RAG enzyme that no longer requires NBD_GRPR/K_ and is capable of robustly cleaving TR2-containing substrates lacking a nonamer. Hence, at least in the context of RAG1 and BbeRAG1L, NBD_GRPR/K_ and CTT_CCGHC_ are functionally redundant, with each rendering the other dispensable. We hypothesize that such redundancy would cause a *RAGL* transposon harboring both domains to be prone to loss of either NBD_GRPR/K_ and the nonamer (as in RAGL-B transposons) or CTT_CCGHC_ and TR2 (as in RAG and the RSS), and that such evolutionary instability might explain the dearth of RAGL enzymes, such as PflRAGL-A, that contain both NBD_GRPR/K_ and CTT_CCGHC_. In addition, loss of CTT_CCGHC_/TR2 by RAG/RSS likely facilitated the evolution of RAG’s capacity for nuanced and flexible DNA recognition that enables it to cleave heterogeneous RSS sequences at widely varying efficiencies (Feeney et al. 2004; Gopalakrishnan et al. 2013; Kim et al. 2018; Ramsden et al. 1994; Swanson 2004; Wu et al. 2020; Yu et al. 2002). Overall, our data strengthen the argument for co-evolution between RAG(L) DNA binding domains and their respective DNA recognition sites.

Our findings and those of other recent studies (Martin et al. 2020; Tao et al. 2022) demonstrate that *RAGL* transposons (of both the A and B family) can possess TIRs with a length asymmetry similar to that of 12/23 RSSs. In addition, our findings with *PflRAGL-A* indicate that jawed vertebrate asymmetric RSSs can readily be explained as arising from the TIRs of a *RAGL-A* transposon—an idea that contradicts the “Transib seed” hypothesis that proposed that RSSs arose directly from a *Transib* element and that was based in part on the premise that appropriate asymmetric TIRs are not found in *RAGL* transposons (Yakovenko et al. 2021).

### Evolution of coupled cleavage

Hydrogen bond formation between RAG1 E649 and S963 helps enforce coupled cleavage by mouse RAG, likely by regulating the structure of an α-helix containing active site residue E962, which in turn determines the integrity of the active site (Kriatchko et al. 2006; Zhang et al. 2019). While RAG1L-A proteins possess position 649/963 aa pairs (E/K or Q/H) that could serve a similar function, this aa pair is V/A in BbeRAG1L-B (Figure 3B), which lacks hydrogen bond potential. Consistent with this, cleavage by BbeRAGL-A is uncoupled (Zhang et al. 2019). More generally, invertebrate RAG1L proteins of the B, C and D families display a non-charged, hydrophobic residue (Val, Ile, Thr) at the position equivalent to E649 and a small amino acid (Ala, Gly, Ser, Cys) at the position equivalent to S963 (Martin et al. 2020). We hypothesize that coupled cleavage activity (and the necessary aa 649/963 pair) arose in a *RAGL-A* transposon prior to speciation of jawed vertebrates, thereby helping to ensure that TIR cleavage occurred in a coordinated fashion in a synaptic complex. Regulated cleavage within an organized synaptic complex is a common feature of bacterial and eukaryotic transposases and site-specific recombinases (Craig 2015) and might have provided a selective advantage to the *RAGL-A* transposon and/or its host by reducing the incidence of uncoupled DNA double strand breaks. An alternative scenario is that coupled cleavage was a property of the ancestral *RAGL* transposon and was subsequently lost in the *RAGL-B* lineage and retained in *RAGL-A*.

### Evolution of a RAG recombinase lacking transposase activity

We previously proposed that suppression of RAG transposition activity began in the jawed vertebrate lineage after creation of the initial “split” antigen receptor gene (Liu et al. 2022). Based on the findings reported here, this hypothesis needs to be revised. Hemichordate and echinoderm RAG2L-A proteins contain an AH of approximately the same size and acidic amino acid content as the AH of jawed vertebrate RAG2 (Figure 5A), and the Pfl and Osp RAG2L-A AH regions potently suppress RAG-mediated transposition when attached to the core region of mouse RAG2 (Figure 5B-D). This is a striking finding given that there is little sequence similarity between the AHs of RAG2L-A and RAG2 and is consistent with the hypothesis that inhibition of transposition depends more on acidic amino acid content than on specific sequence motifs. A similar conclusion was reached regarding the sequence features of the RAG2 AH required to influence repair pathway choice in the post-cleavage phase of V(D)J recombination (Coussens et al. 2013). It will now be important to determine the mechanism by which the AH suppresses transposition—a suppressive activity that is manifest in cells but not when transposition is performed with purified RAG proteins *in vitro* (Zhang et al. 2019).

In addition, PflRAG2L-A possesses a B6-C6 connecting loop that is intermediate in size (8 aa) between that in RAG2L-B (5 aa) and that in jawed vertebrate RAG2 (10 aa). It is not known whether 8 aa is sufficient for transposition inhibition, but regardless, the PflRAG2L-A loop might be indicative of an intermediate in the evolution of the 10 aa loop of RAG2, which suppresses transposition 2-3 fold (Zhang et al. 2020). Furthermore, RAG1L-A proteins possess E or Q at the position equivalent to RAG1 E649, and E at this position suppresses transposition about 2 fold as compared to V (Zhang et al. 2019).

Together, these findings indicate that some adaptations with the potential to suppress transposition (strongly in the case of the AH) arose in invertebrate RAGL-A transposases rather than in jawed vertebrates. Such adaptations, while reducing the mobility of the transposon, might have rendered them less damaging and more readily tolerated by their hosts—a common theme for transposons generally (Almeida et al. 2022; Davies et al. 2000; Levin and Moran 2011; Lohe and Hartl 1996; Saha et al. 2015). Only the change from methionine to arginine at RAG1 position 848 now appears to be a transposition-suppressive adaptation specific to jawed vertebrates. The suppressive effect of the M848R mutation is very strong, particularly *in vivo*, and appears to be due to the ability of methionine, but not arginine, to induce bends in target DNA needed for binding in a deep pocket in the RAG enzyme (Zhang et al. 2020). We refer to R848 as the “gatekeeper” residue for the regulation of RAG-mediated transposition to reflect both its potency and the likelihood that acquisition of arginine at this position a pivotal event in RAG evolution that helped usher in the transition from RAGL transposase to RAG recombinase.

### Evolution of RAG chromatin binding

Through its ability to bind H3K4me3, the RAG2 PHD finger plays a dominant role in specifying sites of chromatin binding by RAG (Maman et al. 2016; Teng et al. 2015) and also increases RAG DNA cleavage activity, apparently by inducing allosteric changes in RAG1 (Bettridge et al. 2017; Lu et al. 2015; Shimazaki et al. 2009). Previous studies have established that the PHD was an early feature of RAG2L proteins, was lost in certain lineages (e.g., BbeRAG2L-B and perhaps amphioxus more generally (Braso-Vives et al. 2022)), and retains the ability to bind methylated lysine in the one case examined (H3K4me2 binding by SpuRAG2L-B) (Fugmann et al. 2006; Huang et al. 2016; Martin et al. 2020; Tao et al. 2022; Wilson et al. 2008). Our findings extend these observations by demonstrating that the PHD is a component of RAG2L-A proteins and by revealing an unexpected “double PHD” in PflRAG2L-A. While certain residues important for methylated lysine binding are intact in RAG2L-A PHDs (e.g., tryptophan at the position equivalent to W453 of mouse RAG2), other important residues are not conserved, and it is difficult to predict whether RAG2L-A PHDs possess histone tail binding activity. The PHD_1_-PHD_2_ of PflRAG2L-A might adopt a structure similar to that of the double PHDs of several chromatin associated proteins (Figure 6C) and it is plausible that it is capable of recognition of some form of acylated lysine. RAG1, very likely through the action of its NTR (aa 1-383), is also able to influence RAG binding in the genome (biasing binding to regions enriched in H3K27ac) (Maman et al. 2016). Some RAG1L proteins of both the A and B families contain all of the major elements found in the RAG1 NTR and hence might also contribute to chromatin binding.

### An updated model of RAG/RAGL evolution

Based on our findings, we propose the following refined model for RAG/RAGL evolution (Figure 7C). It is thought that *Transib*, which was recently identified in bacteria (Tao et al. 2022), preceded *RAGL* in evolution and that the first *RAGL* transposon (which we designate *RAGL^0^*) was created early in eukaryotic evolution when a *RAG2L* gene became incorporated into a *Transib* transposon (Carmona and Schatz 2017; Liu et al. 2022; Tao et al. 2022). While the features of *RAGL^0^* are unknown, parsimony argues that it contained asymmetric TIRs, CTT_CCGHC_, and NBD_GRPR/K_, all of which have been identified in *Transib* transposons (Kapitonov and Jurka 2005; Tao et al. 2022; Zhang et al. 2019). *RAGL^0^* likely also contained a transposition-permissive aa (e.g., methionine) at the position corresponding to RAG1 848 and a PHD at the RAG2L C-terminus. In an early metazoan, *RAGL^0^* gave rise to *RAGL-B* transposons, the defining feature of which was loss of the GRPR/K motif in the NBD and a strong reliance on CTT_CCGHC_ for DNA binding. *RAGL-B* was evolutionarily successful, being transmitted, primarily by vertical transmission, into numerous metazoan lineages including cnidarians, protostomes, and deuterostomes (Martin et al. 2020; Tao et al. 2022). *RAGL^0^* also gave rise to *RAGL-A* through acquisition of an AH in RAG2L and E/Q at the RAG1L position equivalent to RAG1 649 (though as noted above, it is possible that E/Q at position 649 and coupled cleavage were a property of *RAGL^0^*). A *RAGL-A* transposon, perhaps closely resembling *PflRAGL-A*, subsequently found its way into the jawed vertebrate lineage where it created the initial split antigen receptor gene and underwent several adaptations to facilitate its domestication: loss of CTT_CCGHC_ (with strong reliance on NBD_GRPR/K_ and the RSS nonamer), the M848R gatekeeper mutation to terminate transposition, and acquisition of the full B6-C6 RAG2 inhibitory loop.

Questions of particular interest raised by our data and this model are the timing of the emergence of *RAGL-A* and whether *RAGL-A* entered the jawed vertebrate lineage by vertical or horizontal transmission. Vertical inheritance would predict that *RAGL-A* arose in an early deuterostome prior to the divergence of hemichordates and echinoderms, about 560 million years ago (dos Reis et al. 2015). However, the “spotty” evolutionary distribution of *RAGL-A* elements—present in hemichordates and echinoderms but absent from tunicates, cephalochordates, and jawless vertebrates (Figure 7D)—is suggestive of horizontal gene transfer (HGT). Phylogenetic clustering of jawed vertebrate RAG1 and hemichordate RAG1L-A (Figure 2A) is consistent with this idea and with HGT of *RAGL-A* between a hemichordate and an early jawed vertebrate. Genome sequence data from additional species, particularly deuterostomes, is likely to help address the possibility of HGT and might provide an explanation for the absence of TIRs flanking *RAGL* elements in echinoderms. Biochemical analyses of additional Transib transposases might provide insight into the DNA cleavage properties of the RAGL^0^ transposase (e.g., coupled versus uncoupled cleavage).

### Limitations of the study

Our findings do not directly address the functionality of the identified RAGL-A proteins, preventing firm conclusions regarding their ability to perform DNA cleavage or other enzymatic functions. While evidence exists and is presented in our study for mRNA expression for some *RAGL-A* genes, RAGL-A protein expression has not been assessed. Aspects of our model for RAG(L) evolution might need to be revised as the genome sequences of more invertebrate organisms are reported.

## MATERIALS AND METHODS

### Genomic analysis

The genomic and transcriptomic public repositories (WGS/TSA) of all metazoan species were screened using tblastn (Gertz et al. 2006; Johnson et al. 2008) starting from the RAG1L-A sequence identified in *P. flava* (Martin et al. 2020; Morales Poole et al. 2017). The flanking regions of *RAG1L-A* loci were inspected for RAG2-like signatures and positive results were further added to the screening process. Putative *RAG1L/2L-A* loci identified in this way were then subjected to predictions of protein translation using Augustus (Stanke et al. 2008) and Softberry FGENESH+ (Solovyev et al. 2006) and classified as either complete or pseudogenized RAGL sequences depending on the completeness and compliance with canonical RAG1/2 domain organization.

The presence of TIRs, defining the margins of the transposon cassette, was investigated using a homology variation procedure described in (Martin et al. 2020). Briefly, the regions flanking the *RAGL* loci were aligned to identify the transposon ends, based on a higher expectancy of sequence homology within the cassettes as compared to that of the environment at the insertion loci. The cassette end predictions were further scrutinized based on compliance with the expected TIR consensus within the heptamer region and the presence of 5 bp TSDs.

### BAC screening, selection and sequencing

Putative *RAGL1_A* containing *P. flava* clones were identified by screening a *P. flava* Bacterial Artificial Chromosome (BAC) library made from the Hawaiian population (Arshinoff et al. 2022). The presence of *RAG1L-A* was confirmed via colony-PCR followed by Sanger sequencing. The selected BACs were recovered and sequenced on an Illumina MiSeq using reagent kit Index Nextera XT kit V2-500 cycles and assembled with the CLC Genomics Workbench 20.0 v7.5 (QIAGEN). *P. flava* RAGL sequences of the Taiwanese population were retrieved from genome sequence deposited in GenBank (BioProject PRJNA747109).

### N-terminal splicing PCR and sequencing

Adult *P. flava* were collected and embryo cultures were carried out as described previously (Lin et al. 2016). Total RNA was extracted from various developmental stages using the RNeasy Micro kit (Qiagen) and was reverse transcribed using the GoScript Reverse Transcription System (Promega) with oligo dT primers. Primers used to amplify *RAG2L-A* were forward primers (F: 5’- GCAGCCGCCATGCTCGATAGT-3’) and reverse primers (R: 5’- GACCAAAAGCATGAATTTCCTCACCAC-3’). PCR was carried out using the 2xKAPA LongRange HotStart ReadyMix (Kapa Biosystems) for 35 cycles. In some cases, products of the first PCR reaction were used as template for a second PCR. The amplicons were cloned and sequenced to confirm their identities. To examine whether the RNA samples were contaminated with genomic DNA, PCR was performed using equal amount of RNA as template (without reverse transcription). To examine the integrity of the RNA isolated from unfertilized eggs, RT-PCR was conducted to amplify a fragment of *vasa*, a known maternal transcript (Lin et al. 2021), using forward (5’-CGTCAAGCACGTCATCAACT-3’) and reverse (5’-CGTTTCTAATCCCAGGACTC-3’) primers that match to sequences located on different exons.

The *OspRAG1L-A* locus on WGS scaffold JXSR01S003992.1 is interrupted by 2 contig merge areas, each flanked by duplicated segments of <100 bp. To recover the complete sequence, genomic DNA was prepared from a sample of *O. spiculata* (generously provided by T. Arehart, Crystal Cove Conservancy) using SDS/proteinase K digestion and phenol/chloroform extraction followed by isopropanol precipitation. The *OspRAG1L-A* locus was amplified by PCR using 5’- TCGTTCCTGTTTTAGGGACAAAGC and 5’-GTTGTGACCCTCCTTGCCGCATCT as primers. The PCR reaction was carried out using GoTaq Long PCR 2x Master Mix (Promega) for 35 cycles. Amplicons were cloned and the plasmid inserts were initially sequenced using an Applied Biosystems 3730xL DNA Analyzer and then confirmed by the Whole Plasmid Sequencing service from Plasmidsaurus (https://www.plasmidsaurus.com/).

### In vivo recombination and plasmid-to-plasmid transposition assays

The recombination assay was performed in Expi293 cells transfected with 1 μg of pTT5M-RAG1 R848M/E649V and pTT5M-RAG2 variants, and 2 μg of p290G using lipofectamine 2000 as described (Huang et al. 2016; Zhang et al. 2019). Cells were collected 72 hours post-transfection and washed twice with PBS containing 2% FBS. The percentage of live cells expressing GFP was analyzed by flow cytometry as described (Zhang et al. 2019).

Transposition activity was measured using a plasmid-to-plasmid transposition assay as described previously (Zhang et al. 2019). Briefly, 293T cells were transfected with 4 μg each of pTT5M-RAG1 R848M/E649V and pTT5M-RAG2 variants, 6 μg of the donor plasmid (pTetRSS), and 10 μg of the target plasmid (pECFP-1) using polyethyleneimine. The cell medium was changed 24 hours post-transfection and cells were collected after 48 hours. Purified extrachromosomal DNA (300 ng) was used to transform electrocompetent MC1061 bacterial cells, which were plated onto kanamycin or kanamycin-tetracycline-streptomycin (KTS) plates. Transposition efficiency was calculated by dividing the number of colonies on KTS plates by the number of colonies on K plates, correcting for dilution factors. Plasmids from 30 colonies from KTS plates were sequenced to determine whether they contained a bone-fide transposition event (3–7 bp target site duplication (TSD)) and the results used to calculate a corrected transposition efficiency value by counting only the plasmids that contained a transposition event.

### Phylogenetic analyses

The phylogenetic analysis was performed using the Maximum Likelihood method implemented in IQtree (Minh et al. 2020) and PhyML (Guindon et al. 2010) on the most conserved regions of RAG1 (NBD plus catalytic core) and RAG2 (kelch blades 2-5). IQtree phylogeny was performed using automated substitution selection with the FreeRate model for heterogeneity (Soubrier et al. 2012), the ultrafast bootstrap approximation (Minh et al. 2013) and SH-aLRT branch test, both with 1000 replicates, and the approximate Bayes test (Anisimova et al. 2011). In parallel, to assess robustness, phylogeny analyses were performed using PhyML with the SMS model selection (Lefort et al. 2017) using either Akaike or Bayesian Information Criterion (AIC/BIC) and SH-aLRT branch support. Evolutionary relationships between species were retrieved from TimeTree (Kumar et al. 2017) and tree graphics were generated using iTOL (Letunic and Bork 2019).

### Protein sequence analysis

The predicted RAG1L/2L protein sequences were investigated for structural compliance with the expected domain organization and fold characteristics derived from structures of RAG (e.g., (Kim et al. 2015)) and BbeRAGL (Zhang et al. 2019). Secondary structure, relative solvent accessibility and intrinsic disorder predictions were generated using several methods (Buchan and Jones 2019; Cheng et al. 2005; Drozdetskiy et al. 2015; Jones and Cozzetto 2015; Wang et al. 2016) and further merged into a consensus structural profile. Predictive 3D models of the identified *P. flava* and *O. spiculata* sequences were generated using AlphaFold (Jumper et al. 2021), OmegaFold (Wu et al. 2022) and Modeller remote homology modeling (Webb and Sali 2016).

Identity/similarity matrices were computed on the most conserved regions of RAG1 (NBD plus catalytic core) and RAG2 (kelch blades 2-5) using Ugene (Okonechnikov et al. 2012) and in-house scripts. Similarity scores were derived from Blosum62 by considering as similar all amino acid pairs with Blosum62 scores above 0. Protein alignments were performed using MAFFT (Minh et al. 2020) and identity/similarity were computed excluding gap positions.

Sequence variability analysis of the acidic hinge domain and PHD was performed starting from a set of 421 RAG2 sequences retrieved from UniprotKB using Jackhmmer (Johnson et al. 2010) and mouse RAG2 as query. Sequence variability was expressed as KL divergence and computed using Weblogo (Crooks et al. 2004). Analysis of electrostatic 3D surface potential was performed using the Adaptive Poisson-Boltzmann Solver (APBS) and all displayed 3D graphics were generated using the PyMOL Molecular Graphics System, Version 2.2.3 Schrödinger, LLC.

## ACKNOWLEDGEMENTS

The authors thank Tim Arehart for providing *O. spiculata* samples, Ellen Hsu for help in analysis of TIR sequences in the BAC sequence analysis, Katherine Buckley for sending *P. flava* BAC library clones, Yi-Chih Chen for assistance with the RT-PCR analysis, and Shaochun Yuan and Anlong Xu for sharing the sequence of the *OspRAGL-A*_3992 locus. This work was supported by a public grant overseen by the French National Research Agency (ANR) as part of the second “Investissements d’Avenir” program (reference: ANR-17-RHUS-000X) (P.P.), Romanian Academy programs 1 and 3 of IBAR (A.J.P), grant 111-2326-B-001-018 from NSTC, Taiwan (Y.H.S), and NIH grant R01 AI137079 (D.G.S.).

## AUTHOR CONTRIBUTIONS

P.P, D.G.S, A.J.P., and Y.H.S. provided overall direction for the experiments and analyses. E.C.M., L.L.T., and L.T.N. performed database searches, sequence analyses, sequence alignments, and phylogenetic analyses, and contributed to the development of hypotheses and experimental plans. L.T.N., T-P.F., and C-Y.L. contributed to the screening of BAC libraries and sequencing and analysis of BAC clones. E.C.M. performed structural predictions and variability analyses and created the figures. J.X. performed the transposition and V(D)J recombination assays. All authors contributed to the data interpretation and analysis and to the writing of the paper.

## SUPPLEMENTAL DATA

**Figure S1.**
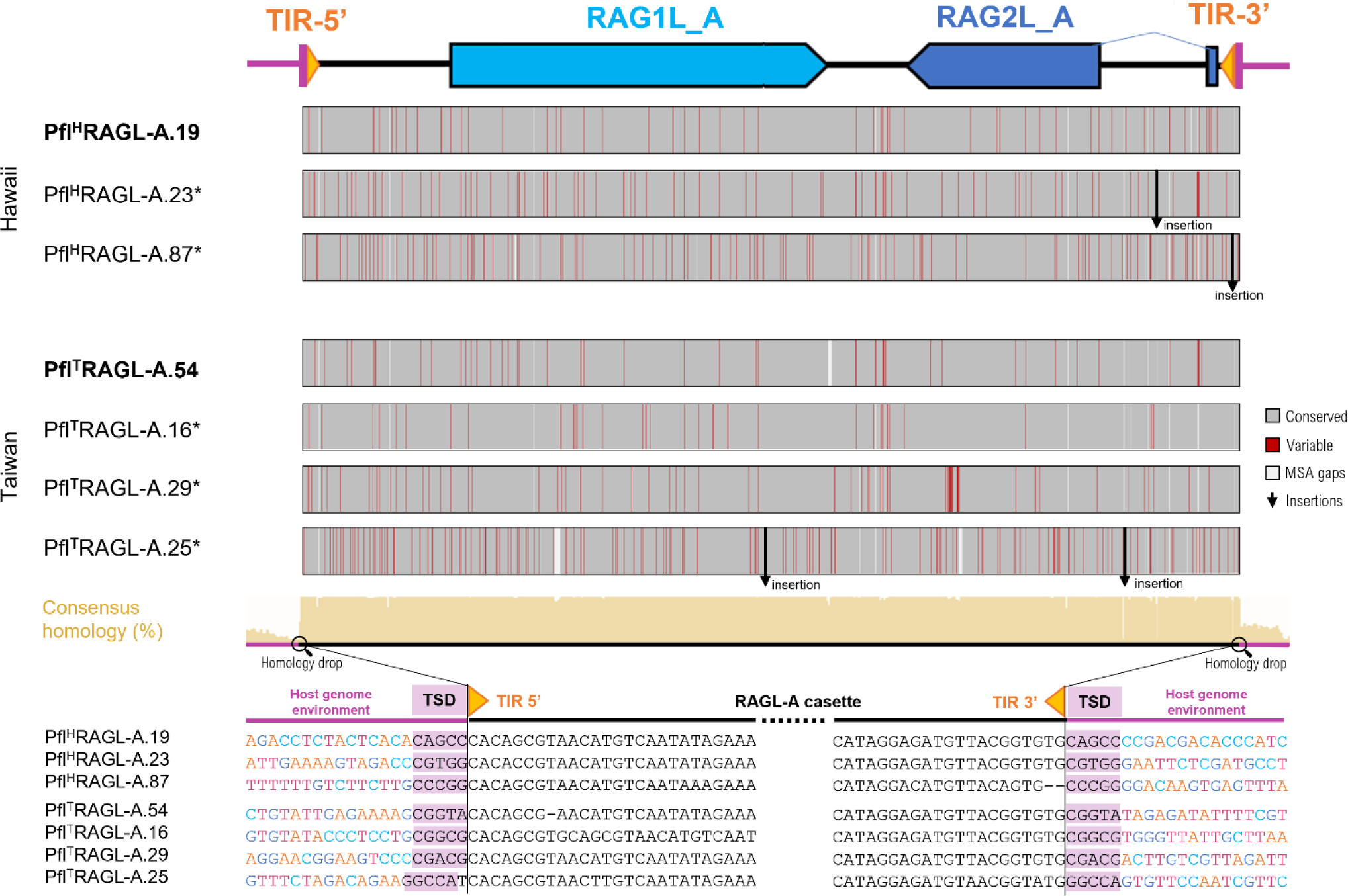
*P. flava* RAGL-A genome cassettes. Comparison of *P. flava* RAGL-A DNA cassettes identified in the Hawaiian and Taiwan populations. Polymorphic positions (red) different from the consensus (gray) are illustrated on the corresponding genomic lane. Short insertions/gaps in the alignment are shown in white, whereas the presence of larger insertions is indicated with an arrow. The consensus line displays the nucleotide conservation percentage as height (taller – more conserved). The margins of the mobile element cassettes are zoomed in at nucleotide level to illustrate the presence of target site duplications (TSD) adjacent to the variable host genome environment.

**Supplemental Figure S2.**
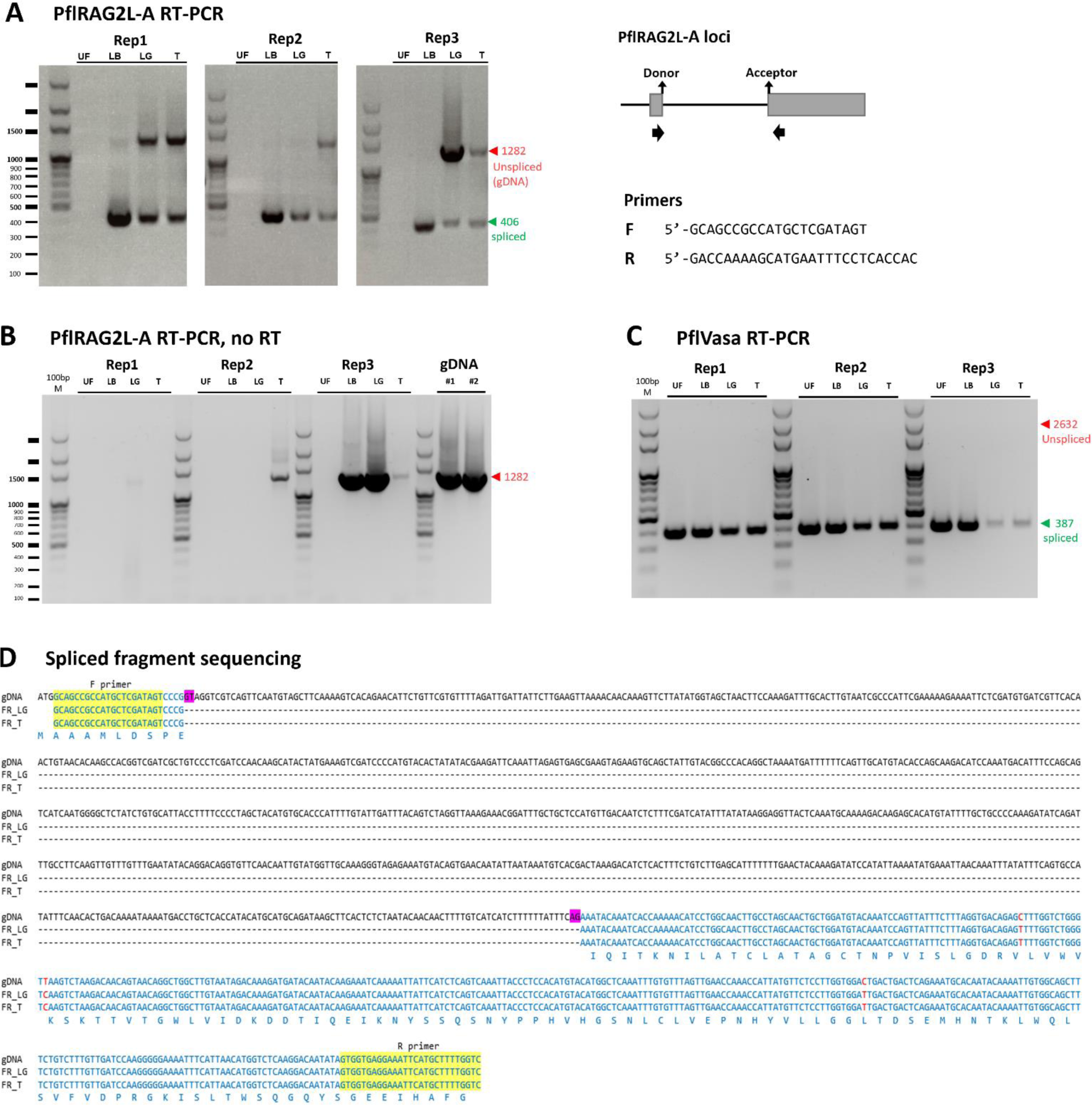
Supporting data for RT-PCR analysis of *PflRAG2L-A* splicing. **A) *PflRAG2L-A* RT-PCR** illustrating PCR products of the size expected from spliced (green) and unspliced (red) mRNA, with mRNA samples from Taiwan *P. flava* of four developmental stages: unfertilized egg (UF), late blastula (LB), late gastrula (LG), and tornaria (T). Replicas (Rep) 1-3 represent different biological replicas performed with independent RNA samples. Rep 1 data are identical to those shown in Figure 1D. **B, C) Control RT-PCR reactions.** B) Control for genomic DNA contamination (in the absence of reverse transcriptase, equal amount of template) demonstrating substantial signal arising from genomic DNA in LB and LG samples in replica 3 and weaker signals in some other lanes. No signal is seen at the size of the product arising from spliced mRNA, indicating that, as expected, signal at this position when reverse transcriptase is added derives from mRNA. PCR using two different extractions of gDNA (genomic DNA) as templates serves as positive controls. C) Control for mRNA integrity by amplifying a fragment of the *Pfl vasa* maternal transcript demonstrating the presence of substantial intact mRNA in the UF samples and reduced levels in LG and T samples, as expected given the decline in *vasa* mRNA levels during development (Lin et al. 2021). **D) Sequence comparison** of the gDNA and the sequenced amplicons corresponding to the spliced variant PCR product from LG and T. Donor/acceptor splice sites are highlighted in magenta, while forward and reverse primer binding sites are depicted in yellow.

**Supplemental Figure S3.**
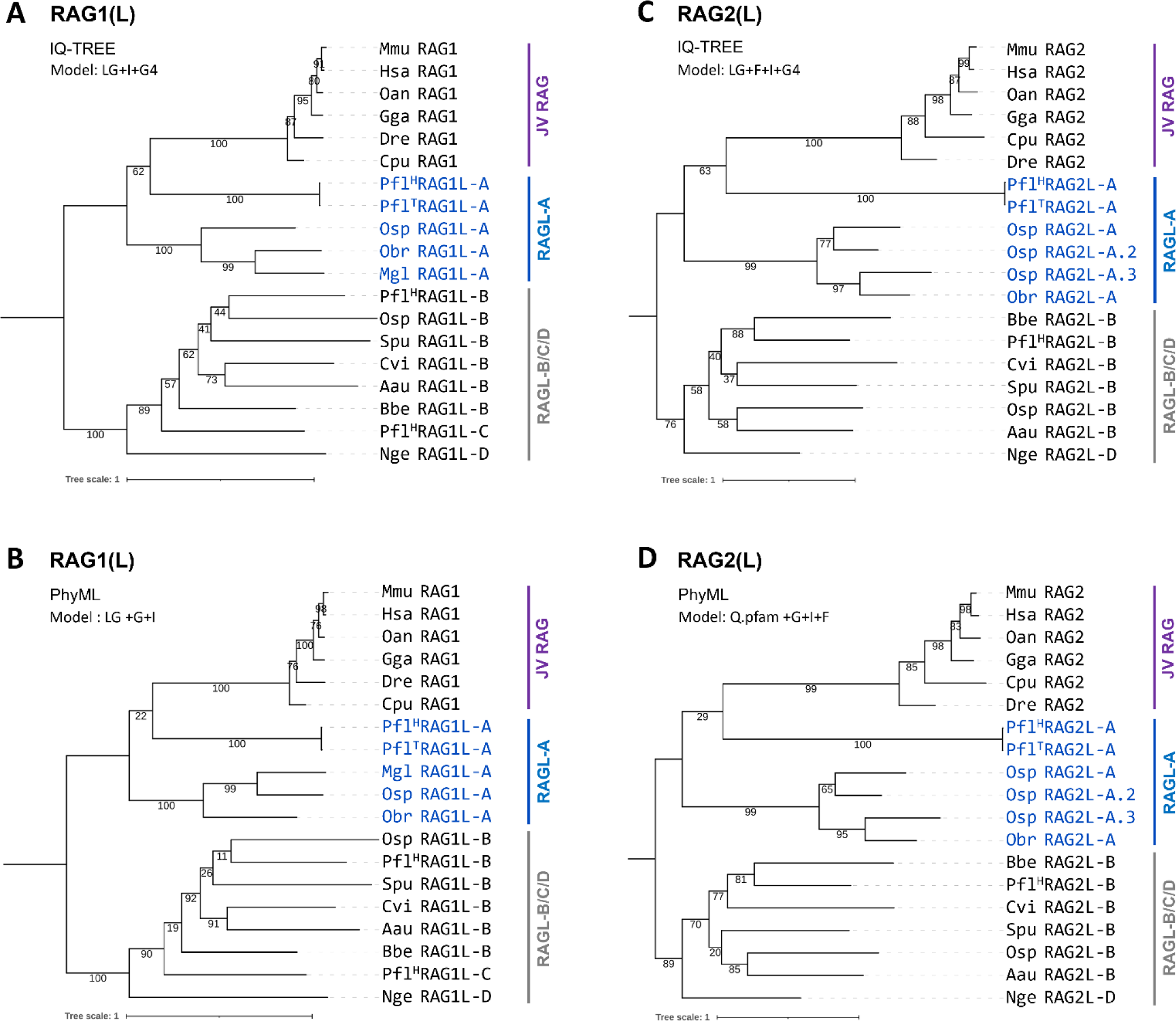
Additional phylogeny analyses of RAG(L) proteins. Additional phylogeny analyses for RAG1L (**A and B**) and RAG2L (**C and D**) sequences using IQ-TREE **(A and C)** and PhyML **(B and D)** with model selection via Bayesian information criterion.

**Supplemental Figure S4.**
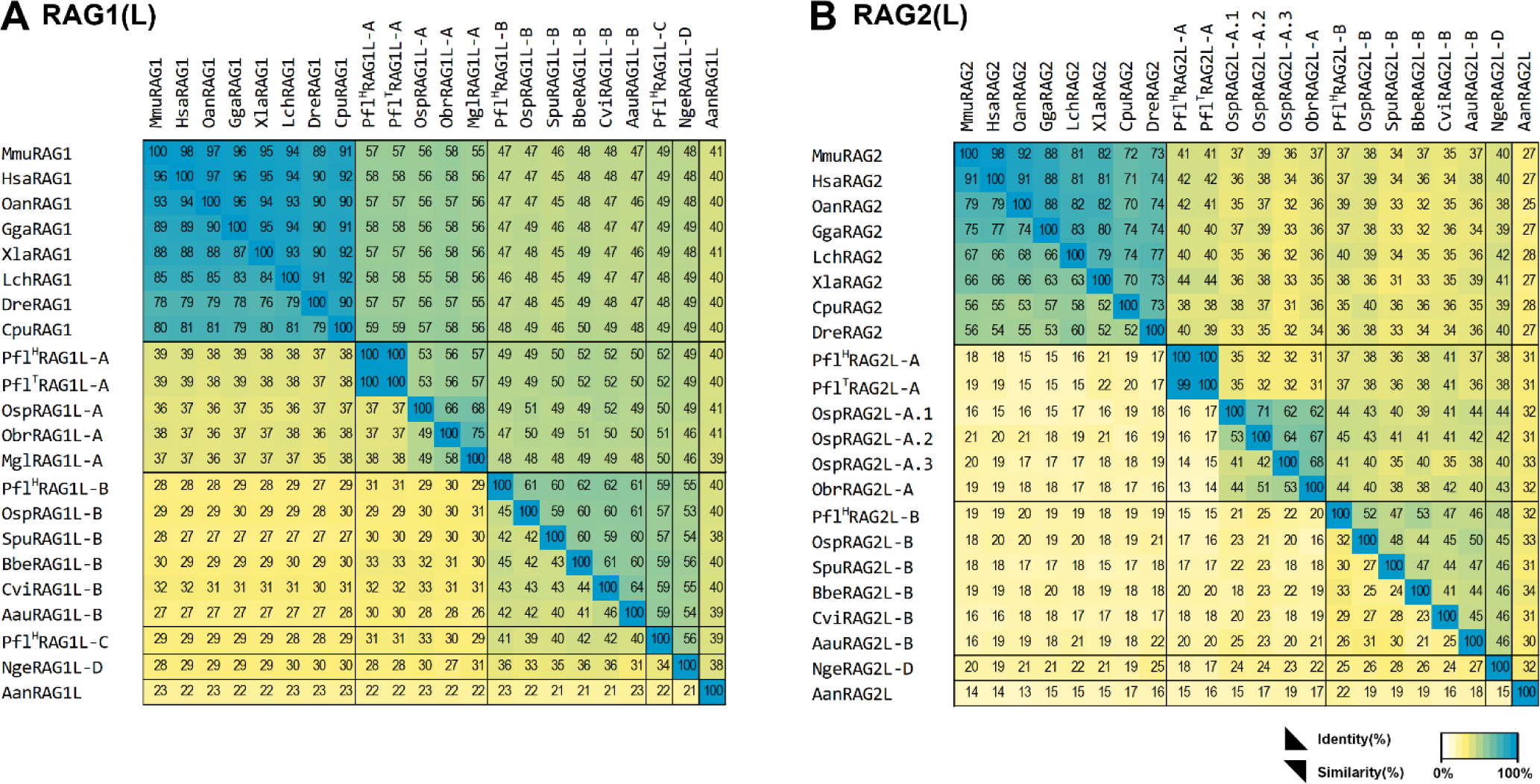
Identity/similarity matrices of RAG(L) proteins. Identity/similarity matrices of RAG1/RAG1L (left) and RAG2/RAG2L (right) sequences from jawed vertebrate and invertebrate A-D lineages. Identity (lower diagonal) and similarity (upper diagonal) are computed on the core of RAG1(L)s (NBD and catalytic core domains) and RAG2(L)s (middle blades 2-5) as detailed in Methods.

**Supplemental Figure S5.**
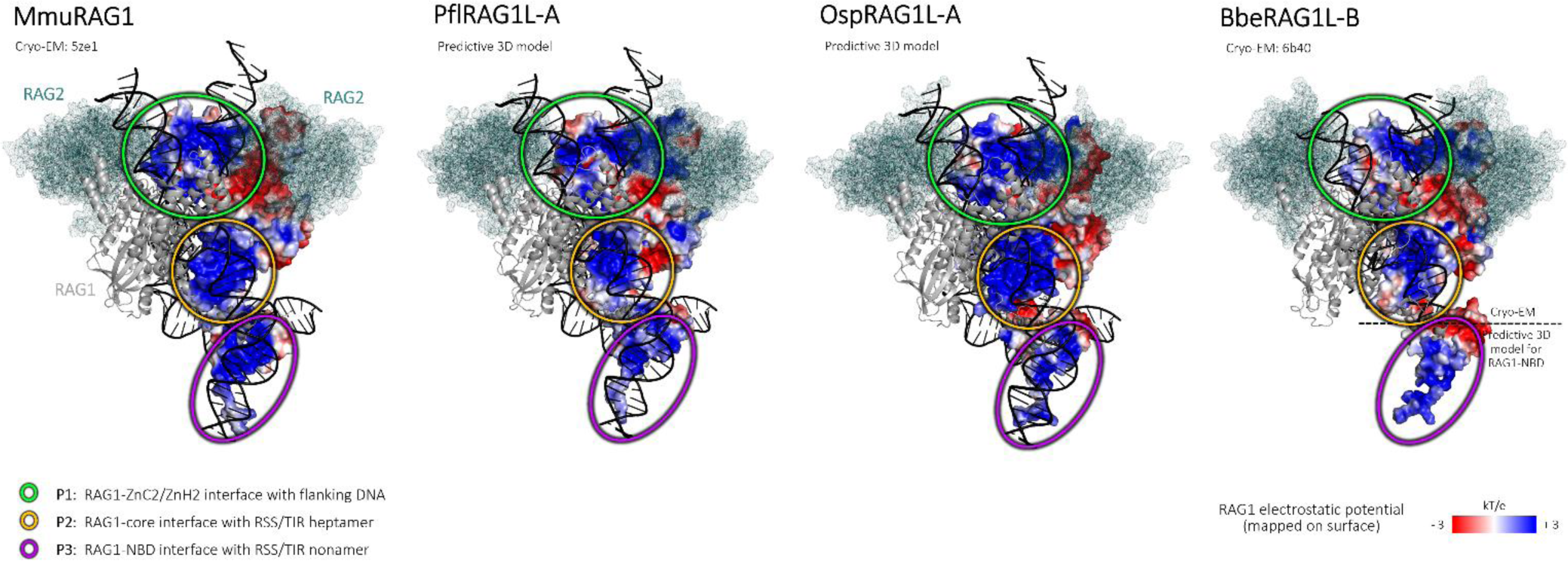
Surface electrostatic charge distribution on RAG1(L) proteins. Comparison of the electrostatic charge distribution on the surface of mouse RAG1 (cryo-EM structure, PDB: 5ZE1), models of *P. flava* and *O. spiculata* RAG1L-A, and BbeRAG1L-B (cryo-EM structure, PDB: 6B40). Because the NBD is not covered by the cryo-EM structure of BbeRAGL, a 3D predictive model of the NBD domain was merged to the tetramer cryo-EM structure. Three positively charged DNA interaction patches are evident in all of the RAG1(L) structures/models and are indicated with colored circles: RAG1 ZnC2/ZnH2 – RSS/TIR flanking DNA (green), RAG1 core – heptamer (orange) and RAG1 NBD – nonamer (purple). Electrostatic surface color code: from blue (positive) to red (negative). The second RAG1(L) subunit of the tetramer is shown in grey, RAG2(L) proteins in teal, and DNA in black.

**Supplemental File S1**

**Genomic data**

Detailed presentation of available genomic and transcriptomic sequence information collected on the *RAGL* loci identified in this study.

**Supplemental File S2**

Multiple sequence alignment of the identified (A) RAG1L-A and (B) RAG2L-A predicted protein sequences. Mapped above the alignment are the functional domains and motifs and the secondary structure assignment. Intermolecular contact interactions are depicted as described in the legend and are derived from the cryo-EM structures of mouse RAG (PDB: 5ZE1) and BbeRAGL-B (PDB: 6B40). In the absence of an experimental structure, consensus secondary predictions of RAGL-A representatives are provided.

