## Supplemental File S1 for "Insights into RAG evolution from the identification of “missing link” family A *RAGL* transposons"

### Supplementary file 1.  Genomic & transcriptomic data

#### Ptychodera flava (Hawaii population)

#### RAGL-A

##### Pfl[H]RAGL-A.19

| Genomic loci | RAG1L status | RAG2L status |
|---|---|---|
| Pfl[H]RAGL-A.19 | potentially functional (1 CDS) | potentially functional (2 CDSs) |

##### FASTA DNA

```
>PflHRAGL-A.19
GGCAGCTGTGCGACTCCAGCACTGCATACATCGCTAACTTTGCACCGTACCTAGGTGTGAAATTTCGGGATCGGACTGATGGGAGACGGCAGGACCGAGGTGTGGTGAAAAGAATTACGATGGAGCTGGTCCAGCCATTTTGTGGATATAATCACAGCCTATTTTGTGTTAGCCTGTTTAGCACGGTCGTTACTGCTAGAGAGCTGATGGAGACCGGGA
TCTACATGGTCGGGAGTTTTAACCGCCGTAATCGTCGCACCATGCCCCCTCAATTAATTCCGCCGGGTAAGAGGAAGCGCCTTCCTCTGTCTGTTGGGGATATGAAAGCCACGACCAGAACGGACTCTCGTCTTAACATCACTGCTTATCAGGATAGTAACGCTCAGGTGCTGATCTTGAATACGGTCTATCCACCCTTGGAGTGCGTGCCAGTGGGG
AGGGAGACGAGCAGCGACAAGTGCCTCTGTCGCTGTATAATTATCGCCGGTACATGGGAGGTGTCGACCGAGCCAACCAGAAGCGCAAGTACTTTCACACTGGGCGCAAGAATCACAGATGGTGGACATATCTGGCATGTTACCTGATTGATGTTGCCCTGGTAAATGCATATATATGCTTCAAGCATGTACACCCTGATAGTAAGCTGACCCATAAAA
TGTTTCATCTGCGTGTCGGGAAACAACTCATTGGCGGTTATTGTGGGCGATCTTAGCCCAGTGTGAGAGAGGTCGAGGCCACCGTTTCTCTTGCCTCGTACATAAATCCACAAAATCAGCCGATGCACGTTGTCAGCCGTTTGGAGGGGGCGGCAGAAATGCTTTAAAGTATGCAGCAGAGAGGGTAAGACCACTGCGAGCGGACGACTGTCTGAGACGT
CCAAAGGATGTAGTCTGTGCGGGGTTCATCTTCACAAGGTCGAGTGTTTTGCGGCTTTTTATCATCGCATGATGCTATACGAGGTAAGAGGAGCGTTGGCGTGCAGACCTCTACTCACACAGCCCACAGCGTAACATGTCAATATAGAAATCGATTAACACTACGATGCGTTGCAGAGAGTATCGACCAATCAGAGAGCAGCATTCATCAAGCCACGAA
CTTTCACGTATGGCTTCGAGATTTCGCGTGATCTAGGGTGTTTACTTCAGCAACTCTACAATCTACTAAATACTAAGGGGTCTCAACTGTTTTACACATGCGCAAACGCCATCTTGAAGAGTGGCTGATGATCATGTCGAAAACCGGAGGTCGAAAGACGTAAGTATCGCCGAGATATCGTAATCTGAATCTCGCGTTGCAATTTTTCACTCGATTATG
TGTATCAAAATTATGCGGATACAATTTTTGGATTTTCTGTCAAACTTGGTTTTTCCTGTTGAAAGTTTGGGTTTTTCTCAGTCCGTAACGACATTGTTTACCGAAGGAACATGTGCTGTACGCTTGGACTTTCTTCAAATGGCTTGGACTTTTAGATCTATACCCATCCAACATGTGTGATTGAGTGTAATGGCCGTTACATTTAAAAATGTTCCTGTG
ATGGTAGCTAAGTCGGGGGAATGATCGTTTTCTGTTTAATATATATTTTTAAGGTTTGAATCAGCGTGTTACGATATTTAGAATTCTGAGCAGCATCCTAGAGGGTCAATTTGGGTGTACAACGCATGGTTCATGCGGTAGCTAGCTAACCCATGCCATATGGCACCAGGTACAGGTTGACTAACGAAGTGTTTTGAGGTTAGCATAGTATGAGTACGTC
AGATTCATATGTCATCGGCATAGAAGGTCGCTTGTGTAAGATAACACCATATTTTTAGGCTGTCATTCATTTCATTTATTTATTTTGCAGTGGGTTGCAGTCAGCAATGAATCATTTACAAGTTTGTTGCACTCTCATGCAATGCAATGCCTTGACCTGATTCAGTATTTCTTGCTTGGGTTCCTGCGGTATGAAGTAGAAAATTAGGCTGGCCACGGT
GTCCTGGGAATTAATGTTTTACCACCAAGGTAAGAATAAAATTCTATTGGATTTATTGTTTACACATATATAAGGATGAAAGGTACAGACCATTGTACCGATTGATACTGTACTTGCCAAATTGATATGCTTAAATAATATTTTAAATATGTTAATTGTTTTCAGAGCGGAATGACGTCAGCTTTTCAACACAGAGAGTACCTTAAGAGGGTGTGTCGT
GTATGTGGTAGAACCCTGAAGACCACATCTTACAGCAACGCAGGACATTACAGACAGGGAGAAGTGTGCATCCCGTACAGTGAACTTTTAGAAGAGTGCTTTAAAGTAAATGTTCAAAATGATGACCCTTTTGTACATCCGCCTAGATTTGGTTCATGTTGTCGTAGAGTTCTTTAAAGTATTCTGTTGCACAGAACCTGGGTAAACATTACTATACC
CATATCAGTGTTTGTAACTGGAACATCATTGTAAGTATGAAAACCAAACTTGTCCTCTTTGCAAACGTTTTTCTCAAAATGGTTCTGGTAGACCTAAAAAACTTAAAAGGGGTCGCTTTAAATTAAATACAAATGAAAATAACGAAACTGACAAGAGTCCTGTTGCAGACAATTTTGCTCATATACTAAATGACCACTGTTATACCACCCAGCTTCC
TGTAAGTTTGCCCAGTCTGTGCAATCAGACCAAAAGACAGATACATATGAATTTTGTACAGAGGAAAATATTCAGACTGTACCCATGCAACAGTCGATTTCCAACAGATTTATGTGTAACTGAACCACATCCAGAATTCATTTGTAGAATGTGTTTAGGAGTCCTAGACACACCACTTCTAACACCGTGTACTCACACTTTCTGTGCAGGCTGCATTAAG
AACTGGTTAAGTCTGTGTCATTTTTGTCCTCTCTGTAAACAACCTATGGCGTCATCAGATTTAAAAGAACCATATAGAGGATTTGTAGAAATTCTTAATGATACTAGAATGAAATGTAATTGTTCAGAAGCAGTTTCAAGCAAAAAAGGTGTCAAGCTCAGTAATTGGCTTCAGCACACTTTAAATTGTGGAAAGGATTTTGAAATTGATGAATGGTTA
GAAAAGTTTTGTCATTGTGGAACGAAATTTCTCTCACTCAAGTTGATTCACATTTAAATAGTACTGAACATTTGCCTGAGACAGATAAGTCAAGTCTGACAACAAGCCATGGCCATCCAGACATTGCTGCAGGTAGCACTGTTGGAAGACCTAAAACTTCACTGTTGTGTCTCCAACAACAAGGACAAAATTAAGATTGAAAAATATCAAAAAACAA
ATAGACGTGTTTGCCAGAGAGAAAGGGGAAGATCTCAGGGTAGTGTATTTTGGCCTTTTGCAAAACTATCTGAGAGCCAACAAGCAAGGTAGTTTAGCTGAACAAGTGAAGTTGCTATATAATGGGGAATCTTGTCAAATGTCACCTCAGGAATGTCTAGCCTTGCGTGTAAATACTTTTTACCAGTTGTAAATCAGTACAAGAAAATTACTCGGCTACA
AATGAGCTTAGTGGAGTTAAAATCTTCCAACCTCTCATCAATGTTGTGAAAGCCGAAGAAAGTTTTCTACCAGGGTCTGTAAAGTTCTACTTTAACCCATCTTTGAAAACTGTTTCTCCCATTTGGCAAGGTACAGAGCATGTTGATGCTCTGTCAGGACTCAGTCGCGATATACCACTTCTAGAGTCACCTGATGTTTATGGGGCAAGGTATAGATAT
GACAAAGCAGTTGCTATGGCTTTGAAAGATATTGATAATGAAATCTCAGAAGGCCTCAGTAAATTAGAAGGGGTGGATACAAATCAACTTGTTCTAACAGCCACTGTCAAAGATGGCAGTGATGGTCTTGGAGATGTGAAAGAAGTCAGATCTCAGGGCCTTGTTCGTTCCAAACAAGGCCTTGCGGTTTTCATTTGTAATTTTAGAAGTCACTGTCAAT
GTAGATGGCAAGATTATTACTGTATTTGAAGAAGATGCTCCAACTCAGAATTATGCACACGCCCTCTCTTGGTAGCTCTGGCAGATGAAAATGACAGAAAAGGGTTATGCATGACACATATACCAATAATGATAGAGAGAGAGGAGATGGAGAAGAGTACCTTGTTTGTAGAAATTGAAGAGGGAAAACAAAGAATCTACAAAGTTGTATTTAAAGGG
ACTATGTATGATGAAAAATTGCAGAGAAGTGGTGAGGGAATGTTAGCGGCAGGGTCATCTTGGTTTTGTACACTTTGTGAGAAAGGAAGACTTGATACCATATCATCAGTCTTGCCAATAACAAGAACACATGAAAACAATATGGCTTTGTATTCAAAGTATGTTGAAAATGCAAATCATATGAAGCAGGCAGATCTAGTGAAAGAAGTCAAAGGCATT
GTTAGAAAACCATTTATAATGTCAGAGCCTTGTGTTGATGCCACGCATGCTGAAATTCACTGGGGGGAGAAAAACTATGATGATATGTATGTACGTGAAATATCAGGGATAATGTGTTGGGGGAAGCCATCCAATGATGATGATGATGCAAAAAAAGTGAAATCAACGAAAAGAATGTTGGATACCGAATTACAGCTGCAATGTGGCTTAAAGAGAGAGCATGGC
TTTATGGTTGATGGAAATTATGCACGTGATTTGGTCAAATCAGAAACAGTTGATGTGGTTTGCTCTTTAATTCAGAGTGAAGCAAGAAAATCTGTTGTTCGTGAATATGTTTCCTACTATCGAGAATTGCGATCAATATACAGGAAACACAATGCTGACTTAGATGATGCCGTGAAATTTAAAACCATAGCTGCAAATTTTTACAAATTGCTACATGAG
AAATTTCCTTATTTGCATCCACTCTCTAACTATCAACACAAGGTATTAGATCACATTCCACAGTTGATCGAGCAATATGGCTCTGTTGGTAAATTTGCCTCTGAAGGCAATGAAAAAGGCAATAAACTTTTCAGAAGATTTCGTAAATTTAATGCACGTAAGTCTCAAATTCATGAAATGCCAGACATTCTGAAATTTCATTGGCTATATACAATGAAA
AGTCTTCAGACTATGGTAAACTGGGAAAAAACTAGATCACTAGTCTGTAGCAAATGTGGTTGTGCTGGTCACAATCAAAGGACATGCATTACATAATTGTGCCCAAGGTCATGATATGAAATCAAAGATGAGACAATGACCATGCATGTTTTATGTATCTGAATCTGTAAGTTTGAAATTAAGTAGCATCTGTATGTATCAGACTACAGAGTCCAAACT
TCAGACACAAAGAGTAGGCTGGCCATACACTCTTATTTACTTAAAAAGTGTTTCTGTTCCATATACTGACTGTACTGTCAAGAGTCATTCTAGCTTCTATGCTAGTCTTAATATACCTAGATGGTTAAAGTGTATTATATTTTGATGTACATAGATGAATTCCACAAGAAATCATCAAATACAAGAAAGCTGAGTTTGTGTTAAATATTTGTATATGT
TTTAATCTTACCTGCCCACACTTTTGAAGGGACTCAAACAAAATCATTCAATTACTTTGGCCAAAATGTCAAAAGTTTGAGTAACTGGCCATGAATATGAGATGAGAAGTGTAAAGTTTGCAATAGAATTTTGTACATATATTCATGAAAGATTGTAACAAGTTGTAAAATGAACTCTCAGTAAAGGTTTCATTCAATTCATGCACATGCTACATGTAC
ATGCTATAATTTCTTGTGCAGTTGTCATGTTCTCAGAAATTACAATCAGACATGTAAATTGATGTACTCTATTTTCCAGTTGTCGTGAAGAAAATCGTTTGCCAGAGCATTTAATGTGAAAAGCTCTAACACATTGTGTACATCGCACTGTTGAAAAGTAACTCTTATCAGTCGCTACAATAATGTTAGTACATGTTGTGAAACATGAAGGG
CATTCAAAGTTTATACAGTCCAGCTTTGTTCTATTTAAGTGATCTAATTCAACACATTTGTCATGGTACCACCTTCCACAGAATACAGTTTGATTGATGTTCATGATGTAAGTTCGATCACATTTCAGCCAGTTGAACTTGGCTTGTTCAGTAGCTGCTAAATTGGCAAATTTGCAGCCATCCAACATACATTCCCATGTTTCCTCCTCACTATCATCTGAT
GTATTGTCTATCTTTCCTGCTCTGCTGCCCTCTTCTGACTCATTCCTGTCTGTGCTAACTTTGACCTCAGCTGCCTCACTTATTCTGTTCTGATCCTCAGTCTCTGAGCCTATACTTTCCTGATCTGTCATGTCTATGCTTATCCTTTCCTCTGCTCTTCCTCCGCTATGCTTACTCTGTCCTCATCTGTTGACCTTTCCTCCTCATCCCCCAACTGACTG
TTATTACTCTGTGTCCTGGTTTCAGAGCTTGTCCCTTCATTTTGAAAACTGAATTTTATCATTTGCTTTCTTACTGTATCACAAACAGCAATTGATCTGAAGAATGGAAGACTGTAGTATTTTACTACTTAAGGTATCATAGTGTGCTGTTTCTATGAAATCTATGGCATCAACAAGGTGACTTGTAAACTGTGTTGTTACTCAATATGTAAACAGTGTTA
TTACATTTGAATGAGACTGGGTTAGGAATACCAATAAATGGTACAGATGTACAGACCATTGAATTCAGGTCAATAACATCAACATCGGGTGGATGTCTTACATTGTTACCTATCACATAAATTCCACCAACAATATACATTTTACTGCTATTTAGAAAAACTGAATGGTAGGCTCTTGGAGTGATCGCTTGTAAACTAATACTTGATAATTTCATAGAT
TGTAAATTAAGAACAGAAAATATGTTTGAAGCCATGGACACATTGTAAAAGCATGACTGAGTTTTTGTGTATATTGTAAGCCTCCATACAGATAGAGAGAATCTCCAAACTTAATTACACTGTGACCAAAAGCATGAATTTCCTCACCACTATATTGTCCTTGAGACCATGTTAATGAAATTTTCCCCCTTGGATCAACAAAGACAGAAAGCTGCCAC
AATTTGTATTGTGCATTCTGAGTCAGTCAATCCACCAAGGAGAACATAATGGTTTGGTTCAACTAAACACAAATTTGAGCCATGTACATGTGGAGGGTAATTTGACTGAGATGAATAATTTTGATTTCTTGTATTGTACCATCTTTGTCTATTACAAGCCAGCCTGTTACTGTTGTCTTAGACTTAACCCAGACCAAAGCTCTGTCACCTAAAGAA
ATAACTGGATTTGTACATCCAGCAGTTGCTAGGCAAGTTGCCAGGATGTTTTTGGTGATTTGTATTTCTGAAATAAAAAAGAGGATGACAAAAGTTGTTGTATTAGAGAGTGAAGCTTATCTGCATGCATGTATGGTGAGCAGGTCATTTTATTTTGTCAGTGTTGAAATATGGCACTGAAATATAAATTTGCTAATTTCATATTTTAATATGGATATC
TTTGTAGTTCAAAAAAAATGCTCAAGACAGAAAGTGAGATGTCTTTAGTCGTGACATTTATTAATATTGTTCACTGTACATTTCTCTACCCTTTGCAACCATACAATTGTTGAACACCTATCCTGTATATTCAAACAAACAACTTGAAGGCAAATCTGATATCTTTGGGGCAGCAAAATACATGTACTCTTGTCTTTTGCATTTGAGTAACCTCCTTAT
ATAAATATGATCGAAAGAGATGTCAACATGGAGCAGCAAATCCGTTTCTTTAACCTAGACTGTAAATCAATACAAAATGGGTGCACATGTAGCTAGGGGAAAAGGTAATGCACAGATAGAGCCCCATTGATGACTGCTGGAAATGTCATTTGGATGTCTTGCTGGTGTACATGCAACTGAAAAAAATCATTTTAGCCTGTGGGCCATACAAT
AGCTGCACTTCTACTTCGCTCACTCTAATTTGAATCTTCGTGTACATGGGGATCGACTTTCATAGTATGCTTGTTGGATCGAGTGACCGTGGCTTGTGTTACAGTTGAGAACGATCACATCGAGAATTTTCTTTTTCGAATGGACGATTACAAGTGCAAATTTTGGAAGTTAGCTACCACATAAGAACTTTGTTGTTTTACTTCAAGAATAATCAAT
CTAAAACAAGAACAGAATGTTCTGTGACTTTTGAAGCTACATTGAACTGACGACCTACCGGGACTATCGAGCATGGCGGCTGCCATCTTGGATATTGTTTACGGCTCACTGAGCATGCTCAGATCAAAAATATTTCCCACAACCCATTGCGGACTTTCTACATTGTAAACATAGGAGATGTTACGGTGTGCAGCCCCGACGACACCCATCAGCAAGAGA
ACCCGACGTAAATAACGGACAGGGGATAGCTTCGCCTCACAGTGGCTGACTGCATTCTTAACATGGACCATGATCACATTTTGAGCACGGTTGCCGTTTGTTTGTGCTTTACGATCCTGCTGATTGTAAATTGAAACACGGATTATTTTGTACAATCCCCCCGATTGTAATCTTTGTAACACGGACCATGCACTCGGACGTCGAGACAGACTCTGAG
ATTTATCATTCGGCATAATGTAAATTATTCCGAATAAAATACTATCTATGGATCGCATATTTTCGTTTGTTTTGTTTAGACGGATTATTTGTACAATTCCCCCTATTATAATTTTTGTAACACGGATCTGCCACCCCGATTGTAATTTTTGAAACACGGACCATGCACTCGGACGTCGAGACAGACTCCGAGATTTTATTGTTCGGCATAATGTAAAT
TATTCCCGAATAAAATACTATCTATGATCGCATATTTTCGTTTGTTTTGTTTTTACCCCCACTTTCGTTTTTGGCAGATCCGTAAGGACGCTATTGTAATGGATGAACGTGCGTTGTATCACGTGGGAGGGGCACAGCCCAACGGAGGCTGTGCTCGTGCGACGTAGACGTTGTACCAACCATCTGTCGTTTGGGTTAGTGCATAGAGCAGTTGCACTG
TCCAGAGCTAAGGTGGTACAAAACTGCACCTTAGTAACACTGCACAACTAACCTGTTAGGAAACGGTAACACTAACGAGCTCAGTGTGCACTGAACTGGCCAAACTACGTACACGCTTTCCTTCAATGGTGAAGCTATGTCGTTGACACTACGTCAAAGCAGACTGCACTGTGCGACTCTTCCAAGTCGTTCAAAGTACGTGGCCAAGTGACACAACC
CAGGGGGAACAGGACAGGTTTGAGGGTGCTGCCGCACCCATAGCACTAACCCCTGGGTCTTCTACTAACTCACGAGCGAGGTGATGTTTCCC
```

### FASTA RNA

**RAG1L-A (Fwd) / partial RAG2L-A (Rev)**

>GDGM01085983.1 TSA: Ptychodera flava comp288179_c0_seq2 transcribed RNA sequence
GTCGGATATCTTTGGAGAAGCAGATCAAAACTCAGGCTGAGTTATGAATTTGAGACTTTCACACAGCTGCTCTCAGAATGTTTCGTTTGGGCATATTTGTCTTAAAATCAAAGCCTTAAGGATAAGTACGCCGAATTCCAACGCGCACGCATTTCCGCGCATTTGACCCCTTAGCTCTACGTAAAATTGTGGTGTTTACCAGTCGGCCATAATAAAAGT
TTCACCGAAGTGCAGCTAGCTACAGCCAACATGAACAGATCTCGTTGGGGTGGAAAATTTTGCTCGAAAACAGAGTTGAAACGACGCAAAAATGTTTCCAAGGCGACCATTGAGCGGTTTGAGAAGAACTCAGTTGATTCACATTCACCAGCGAGAAATGTGGGAACATCGACATCAGATGCTCGTGCACCCATTTGGTCTTCGCCACGCGACGCCGAC
GATACTCAATGCGAACCCTTGATTCCTCCAACTAATCAGCAAATACCGTGGTATGAAGGCGTTAGACTGGTGAATTTGGGAGTGCTTGCAAGAGGATTAGATGCTTGCAATGTTTGCCATGAACACCTGCGTCTGACACAGTGTCAATCTGAAACCAAGTACGGCGGTGGCAGCGTACTCCATATACCCTGTGACCAATGCGGTCTATAAATAATGTA
TATACGAGTAAAACCCATCGGAAAGTTGAGCTGAGCCGTGGTATGCAGGCTTTTGACGTCAACGCCAAAATTGCCACAGGTACGTGCCTATTTTGGTCATATTGAGATTACGTACGGTGAAACGTTGTAACATACGTTTCTACATGTTTCATTTGTGTTCAGTGTGATAACAGCCCAGTACCAGGCGCTAGGCGCGAGACCACTACACAAAAAGTAAAC
ATAGTCGTCACATTTGGTTTTCTCTGTCGGTTGATCATTTCACCAGTGTCCTATTAGATCTGTCACTACTTACAGTTTCTGAAAATTTGTTATTCTTAGTGAACAAGCTGAGGTTCATACAATGTAGCTGCAATAATTATTATCCCGTTGGGGTTGTTTTGACCGTGGCATGGCGCTCATTACTTCCGTATACGCGTATTAGATAGGAAAGCCACGCAC
AGCGTAACATGTCAATATAGAAATCGATTAACACTACGATGAGTCGCAGAGAGTATCGACCAATCAGAGAGCAGCATTCATCAAGCCACGAACTTTCACGTATGGCTTCGAGATTTCGCGTGATCTAGGGTGTTTACTTCAGTAACTCTACAATCTACTAAATACTAAGGGGTCTCAACTGTTCTACACATGCGCAAACGCCATCTTGAAGAGTGGCTG
ATGATCATGTCGAAAACCGGAGGTCGAAAGACTGGGTTGCAGTCAGCAATGAATCATTTACAAGTTTGTTGCACTGTTATGCAATGCAATGCCTTGACCTGATTCAGTATTTCTTGCTTGGGTTCCTGCGGTATGAAGTAGAAAATTAGGCTGGCCACGGTGTCCTTGGAATTAATGTTTTACCACCAAGAGCGGAATGACGTCAGCTTTTCAACACAG
AGAGTACCTTAAGAGGGTGTGTCGTGTATGTGGTAGAACCCTGAAGACCACATCTTACAGCAACTCAGGACATTACAGACAGGGAGAAGTGTGCATCCCGTACAGTGAACTTTTAGAAGAGTGCTTTAAAGTAAATGTTCAAAATGATGATCCTTTTGTACATCCGCCTAGATTTGGTTCATGTTGTCGTAGAGTTCTTTTAAAGTATTCTGTTGCACA
GAACCTGGGTAAACATTACTATACCCATATCAGTGTTTGTAACTGGAAACATCATTGTAAGTATGAAAACCAAACTTGTCCTCTTTGCAAACGTTTTTCTCAAAATGGTTCTGGTAGACCTAAAAAACTTAAAAGGGGTCGCTTTAAATTAAATACAAATGAAAATAATGAAACTGACAAGAGTCCTGTTGCAGACAATTTTGCTCATATACTAAATGA
CCACTGTTATACCACCCCAGCTTCCTGTAAGTTTGCCCAGTCTGTGCAATCAGACCAAAAGACAGATACATATGAATTTTGTACAGAGGAAAATATTCAGACTGTACCCATGACAAGTCGATTTCAACAGATTTATGTGTAACTGAACCACATCCAGAATTCATTTGTAGAATGTGTTTAGGAGTCCTAGACACACCACTTCTAACACCGTGTACTCA
CACTTTCTGTGCAGGCTGCATTAAGAACTGGTTAAGTCTGTGTCATTTTTGTCCTCTCTGTAAACAACCTATGGCGTCATCAGATTTAAAAGAACCATATAGAGGATTTGTAGAAATTCTTAATGATACTAGAATGAAATGTAATTGTCCAGAAGCATTTTCAAGCAAAAAAGGTGTCAAGCTTAGTGATTGGCTTCAGCACACTTTAAATTGTGGAAA
GGATTTTGAAATTGATGAATGGTTAGAAAAAGTTTTGTCATTGTGGAACGAAATTTCTCTCACTCAAGTTGATTCACATTTAAATAGTACTGAACATTTGCCTGAGACAGATAAGTCAAGTCTGACAACAAGCCATGGCCATCCAGACATTGCTGCAGGTAGCACTGTTGGAAGACCTAAAACTTCACTGTTGTGTCTCCAACACAAGGACAAAATTT
AAGATTGAAAAATATCAAAAAACAAATAGACGTGTTTGCCAGAGAGAAAGGGGAAGATCTCAGGGTAGTGTATTTTGGCCTTTTGCAAACTATCTGAGAGCCAACAAGCAAGGTAGTTTAGCTGAACAAGTGAAGTTGCTATATAATGGGGAATCTTGTCAAATGTCACCTCAGGAATGTCTAGCCTTGCGTGTAAATACTTTTACCAGTTGTAATCA
GTACAGAAAATTTACTCGGCTACAAATGAGCTTAGTGGAGTTAAAATCTTCCAACCTCTCATCAATGTTGTGAAAGCTGAAGAAGTTTTCTACCAGGGTCTGTAAAGTTCTACTTTAACCCATCTTTGAAAACTGTTTCTCCCATTTTGGCAAGGTACAGAGCATGTTGATGCTCTGTCAGGACTCAGTCGCGATATACCACTTCTAGAGTCACCTGA
TGTTTATGGGGCAAGGTATAGATATGACAAAGCAGTTGCTATGGCTTTGAAAGATATTGATAATGAAATCTCAGAAGGCCTCAGTAAATTAGAAGGGGTGGATACAAATCAACTTGTTCTAACAGCCACTGTCAAAGATGGCAGTGATGGTCTTGGCGATGTGAAAGAAGTCAGATCTCAGGGGCTTGTCGTTCCAAACAAGGCCTTACGGTTTTCATT
TGTAATTTTAGAAGTCACTGTCAATGTAGATGGCAAGATTATTACTGTATTTGAAGAAGATGCTCCCAACTCAGAATTATGCACACGCCCTCTCTTGGTAGCTCTGGCAGATGAAAATGACAGAAAAGGGTTATGCATGACATATATACCAATAATGATAGAGAGAGAGGAGATTGAGAAAAGTACCTTGTTTGTAGAAATTGAAGAGGGAAAACAAAG
AATCTACAAAGTTGTATTTAAAGGGACTATGTATGATGAAAAATTGCAGAGAGTGGTGAGGGAATGTTTACGCGGCAGGGTCATCTTGGTTTTGTACACTTTGTGAGAAAGGAAGACTTGATCCCATATCATCAGTCTTGCCAATAACAAGAACACATGAAAACAATATGGCTTTGTATTCAAAGTATGTTGAAAATGCAAATCATATGAAGCAGGCAGA
TCTAGTGAAAGAAGTCAAAGGCATTGTTAGAAAACCATTTATAATGTCAGAGCCTTGTGTTGATGCCACGCATGCTGAAATTCACTGGGGAGAGAAAAACTATGATATGTATGTACGTGAAATATCAGGTATAATGTGTTGGGGGAAGCCATCCAATGATGATGATGCAAAAAAAGTGAAATCAACGAAAGAATGTTGGATACCGAATTACAGCTGCA
ATGTGGCTTAAAGAGAGAGCATGGCTTTATGGTTGATGGAAATTATGCACGTGATTTGGTCAAATCAGAAACAGTTGATGTGGTTTGCTCTCTTTAATTCAGAGTGAAGCAAGAAAATCTGTTGTTCGTGAATATGTTTCCTACTATCGAGAATTGCGATCAATATACAGGAAACACAATGCTGACTTAGATGATGCCGTGAAATTTAAAACCATAGCTGC
AAATTTTACAAATGCTACATGAGAAATTCCTTATTTGCATCCACTCTCTAACTATCAACACAAGGTATTAGATCACATTCCACAGTTGATCGAGCAATATGGCTCTGTTGGTAAATTTGCCTCTGAAGGCAATGAAAAAGGCAATAAACTTTTCAGAAGATTTCGTAAATTAATGCACGTAAGTCTCAAATTCATGAAATGCCAGACATTCTGAA
ATTTCATTGGCTATATACAATGAAAGTCTTCAGACTATGGTAAACTGGGAAAAAACTAGATCACTAGTCTGTAGCAAATGTGGTTGTGCTGGTCACAATCAAAGGACATGCATTACATAATTGTGCCCAAGGTCATGATATGAAATCAAAGATGAGACAATGACCATGCATGTTTTATGTATCTGAATCTGTAAGTTTGAAATTAAGTAGCATCTGTA
TGTATCAGACTACAGAGTCCAAACTTCAGACACAAAGAGTAGGCTGGCCATACACTCTTATTTACTTAAAAAGTGTTTCTGTTCCATATACTGACTGTACTGTCAAGAGTCAATTCTAGCTTCTATGCTAGTCTTAATATACCTAGATGGTTAAAGTGTATTATATTTTGATGTACATAGATGAATTCCACAAGAAATCATCAAATACAAGAAAGCTGA
GTTTGTGTTAAATATTTGTATATGTTTTAATCTTACCTGCCCACACTTTTGAAGGGACTCAAACAAAATCATTCAATTACTGTGGCCAAAATGTCAAAAGTTTGAGTAACTGGCCATGAATATGAGATGAGAAGCGTAAGGTTTGCAATAGAATTTTGTACATATATCATGAAAGATTGTAACAAGTTGTAAAATGAACTCTCAGTAAAGGTTTCATT
CAATTCATGCACATGCTACATGTACATGCTATAATTTCTTGTGCAGTTGTCATGTTCTCAGAAATTACAATCAGAACATGTAAATTTGATGTTACTCTATTTTCCAGTTGTCGTGAAGAAAATCGTTTGCCAGAGCATTTAATGTGAAAAGCTCTAACACATTGTGTACATCGCACTGTTGAAAAGTAACTCTTATCAGTCACGCTACAATTAATGTT
AGCACATGTTGTGAAACATGAAGGGCATTCAAAGTTTATACAGTCCAGCTTTGTTCTATTTAAGTGATCTAATTCAACACATTTGTCATGGTACCACCTTCCACAGAATACAGTTTGATTGTTCATGATGTAAGTTCGATCACATTTCAGCCAGTTGAACTTGGCTTGTTCAGTAGCTGCTAAATTGGCAAATTTGCAGCCATCCAACATACATTCCCA
TGTTTCCTCCTCACTATCATCTGATGTATTGTCTATCTTTCCTGCTCTGCTGCCCTCTTCTGACTCATTCCTGTCTGTGCTAACTTTGACCTCAGCTGCCTCACTTATTCCAGTCTCTGAGCCTATACTTTCCTGATCTGTCATGTCTATGCTTATCTTCTTCCTGCTCTTCCTCCGCTATGCTTACTCTGTCCTCATCTGTTGACCTTCCTCCTCATC
CCCCAACTGACTGTTATTACTCTGTGTCCTGGTTTCAGAGCTTGTCCCTTCATTTTGAAAACTGAATTTTATCATTTGCTTTTTTACTGTATCACAAACAGCAATTGAGCTGAAGAATGGAAGACTGTAGTATTTACTACTTAAGGTATCATAGTGTGCTGTTTCTATGAAATCTATGGCATCACAAGGTGACTTGTAAACTGTGTTGTTACTCAATAT
GTAAACAGTGTTATTACATTTGAATGAGACTGGGTTAGGAATACCAATAAATGGTACAGATGTACAGACCATTGAATTCAGGTCAATTACATCAACATCGGGTGGATGTCTTACATTGTTACCTATCACATAAATTCCACCAACAATATACATTTTACTGCTATTACGAAAAACTGAATGGTAGGCTCTTGGAGTGATCGCTTGTAAACTAATACTTGA
TAATTTCATAGATTGTAAATTAAGAACAGAAAATATGTTTGAAGCCATGGACACATTGTAAAAAGCATGACTGAGTTTTTGTGTATATTGTAAGCCTCCATACAGATAAAGAGAATCTCCAAACTTAATTACACTGTGACCAAAAGCATGAATTTCCTCACCACTATATTGTCCTTGAGACCATGTTAATGAAATTTTCCCCCTTGGATCAACAAAGAC
AGAAAGCTGCCACAATTTTGTATTGTGCATTTCTGAGTCAGTCAATCCACCAAGGAGAACATAATGGTTTGGTTCAACTAAACACAAATTTGAGCCATGTACATGTGGAGGGTAATTTGACTGAGATGAATAATTTTTGATTTCTTGTATTGTATCATCTTTGTCTATTACAAGCCAGCCTGTTACTGTTGTCTTAGACTTGACCCAGACCAAAACTCT
GTCACCTAAAGAAATAACTGGATTTGTACATCCAGCAGTTGCTAGGCAAGTTGCCAGGATGTTTTTGGTGATTTGTATTTCTGAAATAAAAAAGATGATGAC

>GDGM01063948.1 TSA: Ptychodera flava comp288179_c0_seq4 transcribed RNA sequence
GTCGGATATCTTTGGAGAAGCAGATCAAAACTCAGGCTGAGTTATGAATTTGAGACTTTCACACAGCTGCTCTCAGAATGTTTCGTTTGGGCATATTTGTCTTAAAATCAAAGCCTTAAGGATAAGTACGCCGAATTCCAACGCGCACGCATTTCCGCGCATTTGACCCCTTAGCTCTACGTAAAATTGTGGTGTTTACCAGTCGGCCATAATAAAAGT
TTCACCGAAGTGCAGCTAGCTACAGCCAACATGAACAGATCTCGTTGGGGTGGAAAATTTTGCTCGAAAACAGAGTTGAAACGACGCAAAAATGTTTCCAAGGCGACCATTGAGCGGTTTGAGAAGAACTCAGTTGATTCACATTCACCAGCGAGAAATGTGGGAACATCGACATCAGATGCTCGTGCACCCATTTGGTCTTCGCCACGCGACGCCGAC
GATACTCAATGCGAACCCTTGATTCCTCCAACTAATCAGCAAATACCGTGGTATGAAGGCGTTAGACTGGTGAATTTGGGAGTGCTTGCAAGAGGATTAGATGCTTGCAATGTTTGCCATGAACACCTGCGTCTGACACAGTGTCAATCTGAAACCAAGTACGGCGGTGGCAGCGTACTCCATATACCCTGTGACCAATGCGGTCTATAAATAATGTA
TATACGAGTAAAACCCATCGGAAAGTTGAGCTGAGCCGTGGTATGCAGGCTTTTGACGTCAACGCCAAAATTGCCACAGGTACGTGCCTATTTTGGTCATATTGAGATTACGTACGGTGAAACGTTGTAACATACGTTTCTACATGTTTCATTTGTGTTCAGTGTGATAACAGCCCAGTACCAGGCGCTAGGCGCGAGACCACTACACAAAAAGTAAAC
ATAGTCGTCACATTTGGTTTTCTCTGTCGGTTGATCATTTCACCAGTGTCCTATTAGATCTGTCACTACTTACAGTTTCTGAAAATTTGTTATTCTTAGTGAACAAGCTGAGGTTCATACAATGTAGCTGCAATAATTATTATCCCGTTGGGGTTGTTTTGACCGTGGCATGGCGCTCATTACTTCCGTATACGCGTATTAGATAGGAAAGCCACGCAC
AGCGTAACATGTCAATATAGAAATCGATTAACACTACGATGAGTCGCAGAGAGTATCGACCAATCAGAGAGCAGCATTCATCAAGCCACGAACTTTCACGTATGGCTTCGAGATTTCGCGTGATCTAGGGTGTTTACTTCAGAAACTCTACAATCTACTACATACTAAGGGGTCTCAACTGTTTTACACATGCGCAAACGCCATCTTGAAGAGTGGCTG
ATGATCATGTCGAAAACCGGAGGTCGAAAGACTGGGTTGCAGTCAGCAATGAATCATTTACAAGTTTGTTGCACTGTTATGCAATGCAATGCCTTGACCTGATTCAGTATTTCTTGCTTGGGTTCCTGCGGTATGAAGTAGAAAATTAGGCTGGCCACGGTGTCCTTGGAATTAATGTTTTACCACCAAGAGCGGAATGACGTCAGCTTTTCAACACAG
AGAGTACCTTAAGAGGGTGTGTCGTGTATGTGGTAGAACCCTGAAGACCACATCTTACAGCAACTCAGGACATTACAGACAGGGAGAAGTGTGCATCCCGTACAGTGAACTTTTAGAAGAGTGCTTTAAAGTAAATGTTCAAAATGATGATCCTTTTGTACATCCGCCTAGATTTGGTTCATGTTGTCGTAGAGTTCTTTTAAAGTATTCTGTTGCACA
GAACCTGGGTAAACATTACTATACCCATATCAGTGTTTGTAACTGGAAACATCATTGTAAGTATGAAAACCAAACTTGTCCTCTTTGCAAACGTTTTTCTCAAAATGGTTCTGGTAGACCTAAAAAACTTAAAAGGGGTCGCTTTAAATTAAATACAAATGAAAATAATGAAACTGACAAGAGTCCTGTTGCAGACAATTTTGCTCATATACTAAATGA
CCACTGTTATACCACCCCAGCTTCCTGTAAGTTTGCCCAGTCTGTGCAATCAGACCAAAAGACAGATACATATGAATTTTGTACAGAGGAAAATATTCAGACTGTACCCATGACAAGTCGATTTCAACAGATTTATGTGTAACTGAACCACATCCAGAATTCATTTGTAGAATGTGTTTAGGAGTCCTAGACACACCACTTCTAACACCGTGTACTCA
CACTTTCTGTGCAGGCTGCATTAAGAACTGGTTAAGTCTGTGTCATTTTTGTCCTCTCTGTAAACAACCTATGGCGTCATCAGATTTAAAAGAACCATATAGAGGATTTGTAGAAATTCTTAATGATACTAGAATGAAATGTAATTGTCCAGAAGCATTTTCAAGCAAAAAAGGTGTCAAGCTTAGTGATTGGCTTCAGCACACTTTAAATTGTGGAAA
GGATTTTGAAATTGATGAATGGTTAGAAAAAGTTTTGTCATTGTGGAACGAAATTTCTCTCACTCAAGTTGATTCACATTTAAATAGTACTGAACATTTGCCTGAGACAGATAAGTCAAGTCTGACAACAAGCCATGGCCATCCAGACATTGCTGCAGGTAGCACTGTTGGAAGACCTAAAACTTCACTGTTGTGTCTCCAACACAAGGACAAAATTT
AAGATTGAAAAATATCAAAAAACAAATAGACGTGTTTGCCAGAGAGAAAGGGGAAGATCTCAGGGTAGTGTATTTTGGCCTTTTGCAAACTATCTGAGAGCCAACAAGCAAGGTAGTTTAGCTGAACAAGTGAAGTTGCTATATAATGGGGAATCTTGTCAAATGTCACCTCAGGAATGTCTAGCCTTGCGTGTAAATACTTTTACCAGTTGTAATCA
GTACAGAAAATTTACTCGGCTACAAATGAGCTTAGTGGAGTTAAAATCTTCCAACCTCTCATCAATGTTGTGAAAGCTGAAGAAGTTTTCTACCAGGGTCTGTAAAGTTCTACTTTAACCCATCTTTGAAAACTGTTTCTCCCATTTTGGCAAGGTACAGAGCATGTTGATGCTCTGTCAGGACTCAGTCGCGATATACCACTTCTAGAGTCACCTGA
TGTTTATGGGGCAAGGTATAGATATGACAAAGCAGTTGCTATGGCTTTGAAAGATATTGATAATGAAATCTCAGAAGGCCTCAGTAAATTAGAAGGGGTGGATACAAATCAACTTGTTCTAACAGCCACTGTCAAAGATGGCAGTGATGGTCTTGGCGATGTGAAAGAAGTCAGATCTCAGGGGCTTGTCGTTCCAAACAAGGCCTTACGGTTTTCATT
TGTAATTTTAGAAGTCACTGTCAATGTAGATGGCAAGATTATTACTGTATTTGAAGAAGATGCTCCCAACTCAGAATTATGCACACGCCCTCTCTTGGTAGCTCTGGCAGATGAAAATGACAGAAAAGGGTTATGCATGACATATATACCAATAATGATAGAGAGAGAGGAGATTGAGAAAAGTACCTTGTTTGTAGAAATTGAAGAGGGAAAACAAAG
AATCTACAAAGTTGTATTTAAAGGGACTATGTATGATGAAAAATTGCAGAGAGTGGTGAGGGAATGTTTACGCGGCAGGGTCATCTTGGTTTTGTACACTTTGTGAGAAAGGAAGACTTGATCCCATATCATCAGTCTTGCCAATAACAAGAACACATGAAAACAATATGGCTTTGTATTCAAAGTATGTTGAAAATGCAAATCATATGAAGCAGGCAGA
TCTAGTGAAAGAAGTCAAAGGCATTGTTAGAAAACCATTTATAATGTCAGAGCCTTGTGTTGATGCCACGCATGCTGAAATTCACTGGGGAGAGAAAAACTATGATATGTATGTACGTGAAATATCAGGTATAATGTGTTGGGGGAAGCCATCCAATGATGATGATGCAAAAAAAGTGAAATCAACGAAAGAATGTTGGATACCGAATTACAGCTGCA
ATGTGGCTTAAAGAGAGAGCATGGCTTTATGGTTGATGGAAATTATGCACGTGATTTGGTCAAATCAGAAACAGTTGATGTGGTTTGCTCTCTTTAATTCAGAGTGAAGCAAGAAAATCTGTTGTTCGTGAATATGTTTCCTACTATCGAGAATTGCGATCAATATACAGGAAACACAATGCTGACTTAGATGATGCCGTGAAATTTAAAACCATAGCTGC
AAATTTTACAAATGCTACATGAGAAATTCCTTATTTGCATCCACTCTCTAACTATCAACACAAGGTATTAGATCACATTCCACAGTTGATCGAGCAATATGGCTCTGTTGGTAAATTTGCCTCTGAAGGCAATGAAAAAGGCAATAAACTTTTCAGAAGATTTCGTAAATTAATGCACGTAAGTCTCAAATTCATGAAATGCCAGACATTCTGAA
ATTTCATTGGCTATATACAATGAAAGTCTTCAGACTATGGTAAACTGGGAAAAAACTAGATCACTAGTCTGTAGCAAATGTGGTTGTGCTGGTCACAATCAAAGGACATGCATTACATAATTGTGCCCAAGGTCATGATATGAAATCAAAGATTAGATAATGACCATGCATGTTTTATGTATCTGAATCTGTAAGTTTGAAATTAAGTAGCATCTGTA
TGTATCAGACTACAGAGTCCAAACTTCAGACACAAAGAGTAGGCTGGCCATACACTCTTATTTACTTAAAAAGTGTTTCTGTTCCATATACTGACTGTACTGTCAAGAGTCAATTCTAGCTTCTATGCTAGTCTTAATATACCTAGATGGTTAAAGTGTATTATATTTTGATGTACATAGATGAATTCCACAAGAAATCATCAAATACAAGAAAGCTGA
GTTTGTGTTAAATATTTGTATATGTTTTAATCTTACCTGCCCACACTTTTGAAGGGACTCAAACAAAATCATTCAATTACTGTGGCCAAAATGTCAAAAGTTTGAGTAACTGGCCATGAATATGAGATGAGAAGCGTAAGGTTTGCAATAGAATTTTGTACATATATCATGAAAGATTGTAACAAGTTGTAAAATGAACTCTCAGTAAAGGTTTCATT
CAATTCATGCACATGCTACATGTACATGCTATAATTTCTTGTGCAGTTGTCATGTTCTCAGAAATTACAATCAGAACATGTAAATTTGATGTTACTCTATTTTCCAGTTGTCGTGAAGAAAATCGTTTGCCAGAGCATTTAATGTGAAAAGCTCTAACACATTGTGTACATCGCACTGTTGAAAAGTAACTCTTATCAGTCACGCTACAATTAATGTT
AGCACATGTTGTGAAACATGAAGGGCATTCAAAGTTTATACAGTCCAGCTTTGTTCTATTTAAGTGATCTAATTCAACACATTTGTCATGGTACCACCTTCCACAGAATACAGTTTGATTGTTCATGATGTAAGTTCGATCACATTTCAGCCAGTTGAACTTGGCTTGTTCAGTAGCTGCTAAATTGGCAAATTTGCAGCCATCCAACATACATTCCCA
TGTTTCCTCCTCACTATCATCTGATGTATTGTCTATCTTTCCTGCTCTGCTGCCCTCTTCTGACTCATTCCTGTCTGTGCTAACTTTGACCTCAGCTGCCTCACTTATTCCGTTCTGATCCTCAGTCTCTGAGCCTATACTTTCCTGATCTGTCATGTCTATGCTTATCTTCTTCCTGCTCTTCCTCCGCTATGCTTACTCTGTCCTCATCTGTTGACCT
TTCCTCCTCATCCCCCAACTGACTGTTATTACTCTGTGTCCTGGTTTCAGAGCTTGTCCCTTCATTTTGAAAACTGAATTTTATCATTTGCTTTTTTACTGTATCACAAACAGCAATTGAGCTGAAGAATGGAAGACTGTAGTATTTACTACTTAAGGTATCATAGTGTGCTGTTTCTATGAAATCTATGGCATCACAAGGTGACTTGTAAACTGTGTT
GTTACTCAATATGTAAACAGTGTTATTACATTTGAATGAGACTGGGTTAGGAATACCAATAAATGGTACAGATGTACAGACCATTGAATTCAGGTCAATTACATCAACATCGGGTGGATGTCTTACATTGTTACCTATCACATAAATTCCACCAACAATATACATTTTACTGCTATTACGAAAAACTGAATGGTAGGCTCTTGGAGTGATCGCTTGTAA
ACTAATACTTGATAATTTCATAGATTGTAAATTAAGAACAGAAAATATGTTTGAAGCCATGGACACATTGTAAAAAGCATGACTGAGTTTTTGTGTATATTGTAAGCCTCCATACAGATAAAGAGAATCTCCAAACTTAATTACACTGTGACCAAAAGCATGAATTTCCTCACCACTATATTGTCCTTGAGACCATGTTAATGAAATTTTCCCCCTTGG
ATCAACAAAGACAGAAAGCTGCCACAATTTTGTATTGTGCATTTCTGAGTCAGTCAATCCACCAAGGAGAACATAATGGTTTGGTTCAACTAAACACAAATTTGAGCCATGTACATGTGGAGGGTAATTTGACTGAGATGAATAATTTTTGATTTCTTGTATTGTATCATCTTTGTCTATTACAAGCCAGCCTGTTACTGTTGTCTTAGACTTAACCCA
GACCAAAGCTCTGTCACCTAAAGAAATAACTGGATTTGTACATCCAGCAGTTGCTAGGCAAGTTGCCAGGATGTTTTTGGTGATTTGTATTTCTGAAATAAAAAAGATGATGAC

### FASTA protein

**RAG1L-A**

>PflHRAG1L-A.19_DNA
MTSAFQHREYLKRVCRVCGRTLKTTSYSNAGHYRQGEVCIPYSELLEECFKVNVQNDDPFVHPPRFGSCCRRVLLKYSVAQNLGKHYYTHISVCNWKHHCKYENQTCPLCKRFSQNGSGRPKKLKRGRFKLNTNENNETDKSPVADNFAHILNDHCYTTPASCKFAQSVQSDQKTDTYEFCTEENIQTVPMTSRFPTDLCVTEPHPEFICRMCLGVLDT
PLLTPCTHTFCAGCIKNWLSLCHFCPLCKQPMASSDLKEPYRGFVEILNDTRMKCNCSEAVSSKKGVKLSNWLQHTLNCGKDFEIDEWLEKVLSLWNEISLTQVDSHLNSTEHLPETDKSSLTTSHGHPDIAAGSTVGRPKTSLLCLQQQGQNLRLKNIKKQIDVFAREKGEDLRVVYFGLLQNYLRANKQGSLAEQVKLLYNGESCQMSPQECLALRV
NTFTSCNQYKKIYSATNELSGVKIFQPLINVVKAEESFLPGSVKFYFNPSLKTVSPIWQGTEHVDALSGLSRDIPLLESPDVYGARYRYDKAVAMALKDIDNEISEGLSKLEGVDTNQLVLTATVKDGSDGLGDVKEVRSQGLVVPNKALRFSFVILEVTVNVDGKIITVFEEDAPNSELCTRPLLVALADENDRKGLCMTHIPIMIEREEMEKSTLFV
EIEEGKQRIYKVVFKGTMYDEKLQRSGEGMLAAGSSWFCTLCEKGRLDTISSVLPITRTHENNMALYSKYVENANHMKQADLVKEVKGIVRKPFIMSEPCVDATHAEIHWGEKNYDMYVREISGIMCWGKPSNDDDAKKVKSTKRMLDTELQLQCGLKREHGFMVDGNYARDLVKSETVDVVCSLIQSEARKSVVREYVSYYRELRSIYRKHNADLDDA
VKFKTIAANFYKLLHEKFPYLHPLSNYQHKVLDHIPQLIEQYGSVGKFASEGNEKGNKLFRRFRKFNARKSQIHEMPDILKFHWLYTMKSLQTMVNWEKTRSLVCSKCGCAGHNQRTCIT

>PflRAG1L-A_RNA_GDGM01085983.1
MTSAFQHREYLKRVCRVCGRTLKTTSYSNSGHYRQGEVCIPYSELLEECFKVNVQNDDPFVHPPRFGSCCRRVLLKYSVAQNLGKHYYTHISVCNWKHHCKYENQTCPLCKRFSQNGSGRPKKLKRGRFKLNTNENNETDKSPVADNFAHILNDHCYTTPASCKFAQSVQSDQKTDTYEFCTEENIQTVPMTSRFPTDLCVTEPHPEFICRMCLGVLDT
PLLTPCTHTFCAGCIKNWLSLCHFCPLCKQPMASSDLKEPYRGFVEILNDTRMKCNCPEAFSSKKGVKLSDWLQHTLNCGKDFEIDEWLEKVLSLWNEISLTQVDSHLNSTEHLPETDKSSLTTSHGHPDIAAGSTVGRPKTSLLCLQQQGQNLRLKNIKKQIDVFAREKGEDLRVVYFGLLQNYLRANKQGSLAEQVKLLYNGESCQMSPQECLALRV
NTFTSCNQYKKIYSATNELSGVKIFQPLINVVKAEESFLPGSVKFYFNPSLKTVSPIWQGTEHVDALSGLSRDIPLLESPDVYGARYRYDKAVAMALKDIDNEISEGLSKLEGVDTNQLVLTATVKDGSDGLGDVKEVRSQGLVVPNKALRFSFVILEVTVNVDGKIITVFEEDAPNSELCTRPLLVALADENDRKGLCMTHIPIMIEREEIEKSTLFV
EIEEGKQRIYKVVFKGTMYDEKLQRSGEGMLAAGSSWFCTLCEKGRLDPISSVLPITRTHENNMALYSKYVENANHMKQADLVKEVKGIVRKPFIMSEPCVDATHAEIHWGEKNYDMYVREISGIMCWGKPSNDDDAKKVKSTKRMLDTELQLQCGLKREHGFMVDGNYARDLVKSETVDVVCSLIQSEARKSVVREYVSYYRELRSIYRKHNADLDDA
VKFKTIAANFYKLLHEKFPYLHPLSNYQHKVLDHIPQLIEQYGSVGKFASEGNEKGNKLFRRFRKFNARKSQIHEMPDILKFHWLYTMKSLQTMVNWEKTRSLVCSKCGCAGHNQRTCIT

>PflRAG1L-A_RNA_GDGM01063948.1
MTSAFQHREYLKRVCRVCGRTLKTTSYSNSGHYRQGEVCIPYSELLEECFKVNVQNDDPFVHPPRFGSCCRRVLLKYSVAQNLGKHYYTHISVCNWKHHCKYENQTCPLCKRFSQNGSGRPKKLKRGRFKLNTNENNETDKSPVADNFAHILNDHCYTTPASCKFAQSVQSDQKTDTYEFCTEENIQTVPMTSRFPTDLCVTEPHPEFICRMCLGVLDT
PLLTPCTHTFCAGCIKNWLSLCHFCPLCKQPMASSDLKEPYRGFVEILNDTRMKCNCPEAFSSKKGVKLSDWLQHTLNCGKDFEIDEWLEKVLSLWNEISLTQVDSHLNSTEHLPETDKSSLTTSHGHPDIAAGSTVGRPKTSLLCLQQQGQNLRLKNIKKQIDVFAREKGEDLRVVYFGLLQNYLRANKQGSLAEQVKLLYNGESCQMSPQECLALRV
NTFTSCNQYKKIYSATNELSGVKIFQPLINVVKAEESFLPGSVKFYFNPSLKTVSPIWQGTEHVDALSGLSRDIPLLESPDVYGARYRYDKAVAMALKDIDNEISEGLSKLEGVDTNQLVLTATVKDGSDGLGDVKEVRSQGLVVPNKALRFSFVILEVTVNVDGKIITVFEEDAPNSELCTRPLLVALADENDRKGLCMTHIPIMIEREEIEKSTLFV
EIEEGKQRIYKVVFKGTMYDEKLQRSGEGMLAAGSSWFCTLCEKGRLDPISSVLPITRTHENNMALYSKYVENANHMKQADLVKEVKGIVRKPFIMSEPCVDATHAEIHWGEKNYDMYVREISGIMCWGKPSNDDDAKKVKSTKRMLDTELQLQCGLKREHGFMVDGNYARDLVKSETVDVVCSLIQSEARKSVVREYVSYYRELRSIYRKHNADLDDA
VKFKTIAANFYKLLHEKFPYLHPLSNYQHKVLDHIPQLIEQYGSVGKFASEGNEKGNKLFRRFRKFNARKSQIHEMPDILKFHWLYTMKSLQTMVNWEKTRSLVCSKCGCAGHNQRTCIT

**RAG2L-A**

>PflHRAG2L-A.19_DNA
MAAAMLDSPEIQITKNILATCLATAGCTNPVISLGDRALVWVKSKTTVTGWLVIDKDGTIQEIKNYSSQSNYPPHVHGSNLCLVEPNHYVLLGGLTDSEMHNTKLWQLSVFVDPRGKISLTWSQGQYSGEEIHAFGHSVIKFGDSLYLYGGLQYTQKLSHAFYNVSMASNIFSVLNLQSMKLSSISLQAITPRAYHSVFLNSSKMYIVGGIYVIGNNVR
HPPDVDVIDLNSMVCTSVPFIGIPNPVSFKCNNTVYILSNNTVYKSPCDAIDFIETAHYDTLSSKYYSLPFFRSIAVCDTVRKQMIKFSFQNEGTSSETRTQSNNSQLGDEEERSTDEDRVSIAEEEQERISIDMTDQESIGSETEDQNRISEAAEVKVSTDRNESEEGSRAGKIDNTSDDSEEETWECMLDGCKFANLAATEQAKFNWLKCDRTYIMN
NQTVFCGRWYHDKCVELDHLNRTKLDCINFECPSCFTTCANINCSVTDKSYFSTVRCTQCVRAFHIKCSGKRFSSRQLENKSNIKFTCSDCNF

>PflRAG2L-A_RNA_GDGM01085983.1_Reverse
VIIFFISEIQITKNILATCLATAGCTNPVISLGDRVLVWVKSKTTVTGWLVIDKDDTIQEIKNYSSQSNYPPHVHGSNLCLVEPNHYVLLGGLTDSEMHNTKLWQLSVFVDPRGKISLTWSQGQYSGEEIHAFGHSVIKFGDSLYLYGGLQYTQKLSHAFYNVSMASNIFSVLNLQSMKLSSISLQAITPRAYHSVFRNSSKMYIVGGIYVIGNNVRHP
PDVDVIDLNSMVCTSVPFIGIPNPVSFKCNNTVYILSNNTVYKSPCDAIDFIETAHYDTLSSKYYSLPFFSSIAVCDTVKKQMIKFSFQNEGTSSETRTQSNNSQLGDEEERSTDEDRVSIAEEEQERISIDMTDQESIGSETGISEAAEVKVSTDRNESEEGSRAGKIDNTSDDSEEETWECMLDGCKFANLAATEQAKFNWLKCDRTYIMNNQTVFC
GRWYHDKCVELDHLNRTKLDCINFECPSCFTTCANINCSVTDKSYFSTVRCTQCVRAFHIKCSGKRFSSRQLENKSNIKFTCSDCNF

>PflRAG2L-A_RNA_GDGM01063948.1_Reverse
VIIFFISEIQITKNILATCLATAGCTNPVISLGDRALVWVKSKTTVTGWLVIDKDDTIQEIKNYSSQSNYPPHVHGSNLCLVEPNHYVLLGGLTDSEMHNTKLWQLSVFVDPRGKISLTWSQGQYSGEEIHAFGHSVIKFGDSLYLYGGLQYTQKLSHAFYNVSMASNIFSVLNLQSMKLSSISLQAITPRAYHSVFRNSSKMYIVGGIYVIGNNVRHP
PDVDVIDLNSMVCTSVPFIGIPNPVSFKCNNTVYILSNNTVYKSPCDAIDFIETAHYDTLSSKYYSLPFFSSIAVCDTVKKQMIKFSFQNEGTSSETRTQSNNSQLGDEEERSTDEDRVSIAEEEQERISIDMTDQESIGSETEDQNGISEAAEVKVSTDRNESEEGSRAGKIDNTSDDSEEETWECMLDGCKFANLAATEQAKFNWLKCDRTYIMNNQ
TVFCGRWYHDKCVELDHLNRTKLDCINFECPSCFTTCANINCSVTDKSYFSTVRCTQCVRAFHIKCSGKRFSSRQLENKSNIKFTCSDCNF

### Pfl$^{H}$RAGL-A.23

| Genomic loci | RAG1L status | RAG2L status |
| --- | --- | --- |
| Pfl$^{H}$RAGL-A.23 | pseudogenised(frameshifts) | potentially functional (2 CDSs) |

#### FASTA DNA

>PflHRAGL-A.23
ACATGCAATAAAGATGGCGTTCATGTTATGTTAGCTGCTTGTTCGTTGCCTATTATTAACGTGTCCCATAACATAACAGCCATACAACATGGTGTCAGGAGTGAACTAAAGTCTGAATGGGCTTATAAAGAACACAGATTTGAGGCATCGGTGTACTAGACTGAAGCAGATCCACGAGAAATTGAAAAGTAGACCCGTGGCACACCGTAACATGTCAATATAGAAATCGATTAACACTAC
GATGCGTCCAAGAGAGTATCGACCAATCAGAGAGCAGCATTCATCAAGGACGACCTTTCACGTATGGCTTCGAGATTTCGCGTGATCTAGGGTGTTTACTTCAGAAACTCTACAATCTACTACATACTAAGGGGTCTCAACTGTTTTACACATGCGCAAACGCCATCTTGAAGAGTGGCTGATGATCATGTCGAAAACCCGGAGGTCGAAAGACGTAAGTATCGCCGAGATAACGTAATC
TGAATCTCGCGTTGCAATTTTTCACTCGATTATGTGTATCAAAATTATGCGGATACAATTTTGGGATTTTCTGTCAAACTTGGTTTTTCCTGTTTAAAGTTTGGGTTTTTCTCAGTCCGTAACGACATAGTTTACCGAAGGAACATGTGCTGTACGCTTGGACTTTCTTCAAATGGCTTGGACTTTTAGATCTATACCCATCCAACATGTGTGATTGAGTGTAATGGCCATTACATTTAA
AAATGTTCCTGTGATGGTAGCTAGGTCAGGGAAATGATCGTTTCCTGTTTAATATATTTTTAAGGTTTGAATGAGCGTGTTACGATATTTAGAATTATGAGCAGCATCCTAGAGGGTCAATTTGGGTGTACAACGCATGGTTCATGCGGTAGCTAGCTAACCCAGGCCATATGGCACCAGGTACAGGTTGACTAACGAAATGTTTTGAGGTTAGCATAGTATGAGTACGTCAGATTCATA
TGTCATTGGCATAGAAGGTCGCTTGTGTAAGATAAGACCATATTTTTAGGCTGTCATTCATTTCATTTATTTATTTTGCAGTGGGTTGCAGTCAGCAATGAATCATTTACAAGTTTGTTGCACTGTTATGCAATGCAATGCCTTGACCTGAATCAGTATTTCTTGCTTGGGTTCCTGCGGTATGAAGTAGAAAATTAGGCTGGCCACGGTGTCCTGGGAATTAATGTTTTACCACTAAGG
TAAGAATAAAATTCTATTGGATTTATTGTTTACACATATATAAGGATGAAAGGTACACACCATTGTACTGATTGGTACTGTACTTGCCAAATTGATATGCTTAAATAATATTTTAAATATGTTAATTGTTTTCAGAGCGGAATGACATCAGCTTTTCAACACAGAGAGTACCTTAAGAGGGTGTGTCGTGTATGTGGTAGAACCCTGAAGACCACATCTTACAGCAGCTCAGGACATTAC
AGACAGGGAGAAGTGTGCATCCCATACAGTGAACTTTTAGAAGAATGCTTTAAAGTAAATGTTTGAAATGATGACCCTTTTGTACATCCGCCTAGATTTGGTTCATGTTGCCGTAGAGTTCTTTTAAAGTATTCTGTTGCACAGAACCTGGGTAAACATTACTATACCCATATCAGTGTTTGTAACTGGAAACATCATTGTAAGTATGAAAACCAAAGTTGTCCTCTTTGCAAACGTTTT
TCTCAAAATGGTTCTGGTAGACCTAAAAAACTTAAAAGGGGTCGCTTTAAATTAAATACAAATGAAAATAATGAAACTGACAAGAGTCCTGTTGCAGACAATTTTGCTCATATACCAAATGACCACTGTTATACCACCCCAGCTTCCTGTAAGTTTGCCCAGTCTGTGCAATCAGACCAAAAGACAGAGACATATGAATTCTGTACGGAGGAAAATATTCAGACTGTACCCATGACAAGT
CGATTCCCAACAGATTTATGTGTAACTGAACCACATCCAGAATTCATTTGTAGAATGTGTTTAGGAGTCCTAGACACACCACTTCTAACACCGTGTACTCACACTTTCTGTGCAGGCTGCATTAAGAACTGGTTAAGTCTGTGTCATTAATGTCCTCTCTGTAAACAACCTATGGCGTTATCAGATTTAAAAGAACCATATAGAGGATTTGTAGAAATTCTTAATGATACTAGAATGAAA
TGTAATTGCCCAGAAGCAGTTTCAAGCAAAAAAGGTGTCAAGCTTAGTAATTGGCTTCAGCACACTTTAAATTGTGGAAAGGATTTTGAAATTGATGAATGGTTAGAAAAAGTTTTGTCATTGTGGAACGAAATTTCTCTCACTCAAGTTAATTCACATTTAAATAGTACTGAACATTTGTCTGAGACAGATAAGTCAAGTCTGACAACAAGCCATGGCCATCCAGACATTGCTGCAGGT
AGCACTGTTGGAAGACCTAAAACTTCACTGTTGTGTCTCCAACAACAAGGACAAAATTTAAGATTGAAAAATATCAAAATACAAATAGACGTGTTTGCCAGAGAGAAAGGGGAAGATCTCAGGGTAGTATATTTTGGCCTTTTGCAAAACTATCTGAGAGCCAACAAGCAAGGTAGTTTAGCTGAACAAGTGAAGTTGCTATATAATGGGGAATCTTGTCAAATGTCACCTCAGGAATGT
CTAGCCTTGCGTGTAAATACTTTTACCAGTTGTAATCAGTACAAGAAAATTTCCTCAGCTACAAATGAGCTTAGTGGAGTTAAAATCTTCCAACCTCTCATCAATGATGTGAAAGCCGAAGAAAGTTTTCTACCAGGGTCTATAAAGTTCTACTTTAACCCATCTTTGAAAACTGTTTCTCCCATTTGGCAAGGTACAGAGCATGTTGATGCTCTGTCAGGACTCAGTCGCAATATACCA
CTTCTAGAATCACCTGATGTTTATGGGGCAAGGTATAGATATGACAAAGCAGTTGCTATGGCTTTGAAAGATATTGATAATGAAATCTCAGAAGGCCTCAGTAAATTAGAAGGGGTGGATACAAATCAACTTGTTCTAACAGCCACTGTCAAAGATGGCAGTGATGGTCTTGGAGATGTGAAAGAAGTCAGATCTCAGGGGCTTGTCATTCCAAACAAGGCCTTGCGGTTTTCATTTGTA
ATTTTAGAAGTCACTGTCAATGTAGATGGCAAGATTATTACTGTATTTGAAGAAGGTTCTCCCAACTCAGAATTATGCACACGCCCTCTCTTGGTAGCTCTGGCAGATGAAAATGACAGAAAAGGGTTATGCATGACACATATACCAATAATGATAGAGAGAGAGGAGATGGAGAAGAGTACCTTGTTTGTAGAAATTGAAGAGGGAAAACAAAGAATCTACAAAGTTGTATTTAAAGGG
ACTATGTATGATGAAAAATTGCAGAGAAGTGGTGAGGGAATGTTAGCGGCAGGGTCATCTTGGTTTTGTACACTTTGTGAGAAAGGAAGACTCGATCCCATATCATCAGTCTTGCCAATAACAAGAACACATGAAAACAATATGGCTTTGTATTCAAAGTATGTTGAAAATGCAAATCATATGAAGCAGGCAGATCTAGTGAAAGAAGTCAAAGGCATTGTTAGAAAACCATTTATAATG
TCAGAGCCTTGTGTTGATGCCACGCATGCTGAAATTCACTGGGGAGAGAAAAACTATGATGTGTATGTACGTGAAATATCAGGGATAATGTGTTGGGGGAAGCCATCCAATGATGATGATGCAAAAAAAGTGAAATCAACAAAAAGAATGTTGGATACCAAATTACAGCTGCAATGTGGCTTAAAGAGAGAGCATGGCTTTATGGTTGATGGAAATTATGCACGTGATTTGGTCAAATCA
GAAACAGTTGATGTGGTTTGCTCTTTAATTCAGAGTGAAGCAAGAAAATCTGTTGTTCGTGAATATGTTTCCTACTATCGAGAATTGCGATCAATATACAGGAAACACAATGCTGACTTAGATGATGCCGTGAAATTTAAAACCATAGCTGCAAATTTTTACAAATTGCTACATGAGAAATTTCCTTATTTGCATCCAATCTCTAACTATCAACACAAGGTATTAGATCACATTCCACAG
TTGATCGAGCAATATGGCTCTGTTGGTAAATTTGCCTCTGAAGGCAATGAAAAAGGCAATAAACTTTTCAGAAGATTTCGTAAATTTAATGCACGTAAGTCTCAAATTCATGAAATGCCAGACATTCTGAAATTTCATTGGCTATATACAATGAAAAGTCTTCAGACTATGGTAAACTGGAAAAAAACTAGATCACTCGTCTGTAGCAAATGTGGTTGTGCTGGTCACAATCAAAGGACA
TGCATTACATAATTGTGCCCAAGGTCATGATATGAAATCAAAGATGAGACAATGACCATGCATGTTTTATGTATCTGAATCTGTAAGTTTGAAATTAAGTAGCATCTGTATGTATCAGACTACAGAGTCCAAACTTCAGACACAAAGAGTAGGCTGGCCATACACTCTTATTTACTTAAAAAGTGTTTCTGTTCCATATACTGACTGTACTTTCAAGAGTCAATTCTAGCTTCTATGCTA
GTCTTAATATACCTAGATGGTTAGAGTGTATTATATTTTGATGTACATAGATGAATTCCACAAGAAATCATCAAATACAAGAAAGCTGAGTTTGTGTTAAATATTTGTATGTGTTTTAATCTTACCAGTAACCTGCCCACACTTTTGAAGGGACTCAAACAAAATCATTTGATTACTTTGGCAGAAAAGTCAAAAGTTTTAGTTATACTGGCCATAAAAAATGAGATGAGAAGCGTAAAG
TTTGCAATAGAAGTTTGTACATATATTCATGAAAGATTGTAACAAGTTGTAAAATGAACTCTCAGTAAAGGTTTCATTCAATTCATGCACATGCTACATGTACATGCTATGATTTCTTGTGCAGTTGTCATGGTCTTCAGAAATTACAATCAGAACATGTAAATTTGATGTTACTCTTATTTTCCAGTTGTCGTGAAGAAAATCGTTTGCCAGAGCATTTAATGTGAAAAGCTCTAACACA
TTGTGTACATCGCACTGTTGAAAAGTAACTCTTATCAGTCACGCTACAATTAATGTTAGCACATGTTGTGAAACATGAAGGGCATTCAAAGTTTATACAGTCCAGCTTTGTTCTATTTAAGTGATCTAATTCAACACATTTGTCATGGTACCACCTTCCACAGAGTACAGTTTGATTGTTCATGATGTAAGTTCGATCACATTTCAGCCAGTTGAACTTGGCTTGTTCAGTAGCTGCTAA
ATTGGCAAATTTGCAGCCATCCAACATACATTCCCATGTTTCCTCCTCACTTTCATCTGATGTATTGTCTATCTTTCCTGCTCTGCTGCCCTCTTCTGACTCATTCCTGTCTGTGCTAACTTTGACCTCAGCTGCCTCACTTATTCCGTTCTGATCCTCAGTCTCTGAGCCTGTACTTTCCTGATCTGTCATGTCTATGCTTATCCTTTCCTGCTCTTCCTCCGCTATGCTTACTCTGTC
CTCATCTGTTGACCTTTCCTCCTCATCACCCAACTGACTGTTATTACTCTGTGTCCTGGTTTCAGAGCTTGTCCCTTCATTTTGAAAACTGAATTTTATCATTTGCTTTTTTACTGTATCACAAACAGCAATTGAGCTGAAGAATGGAAGACTGTAGTATTTACTACTTAAGGTATCATAGTGTGCTGTTTCTATGAAATCTATGGCATCACAAGGTGACTTGTAAACTGTGTTGTTACT
CAATATGTAAACAGTGTCATTACATTTGAATGAGACTGGGTTAGGAGTACCAATAAATGGTACAGATGTACAGACCATTGAATTCAGGTCAATTACGTCAACATCGGGTGGATGTCTTACATTGTTACCTATCACATAAAGTCCACCAACAATATACATTTTACTGCTATTTAGAAAAACTGAATGGTAGGCTCTTGGAGTGATAGCTTGTAAACTAATACTTGATAATTTCATAGATTG
TAAATTAAGAACAGAAAATATGTTTGAAGCCATGGACACATTGTAAAAAGCATGACTGAGTTTTTGTGTATATTGTAAGCCTCCATAAAGATAAAGAGAATCTCCAAACTTAATTACACTGTGACCAAAAGCATGAGTTTCCTCACCACTATATTGTCCTTGAGACCATGTTAATGAAATTTTCCCCCTTGGATCAACAAAGACAGAAAGCTGCCACAATTTTGTATTGTGCATTTCTGA
GTCAGTCAGTCCACCAAGGAGAACATAATGGTTTGGTTCAACTAAACACAAATTTGAGCCATGTACATGTGGAGGGTAATTTGACTGAGATGAATAATTTTTGATTTCTTGTATTGTATCATCTTTGTCTCTTACAAGCCAGCCTGTTACTGTTGTCTTAGACTTAACCCAGACCAAAGCTCTGTCACCTAAAGAAATAACTGGATTTGTACATCCAGCAGTTGCTAGGCAAGTTGCCAG
GATGTTTTTGGTGATTTGTATTTCTGAAATAAAAAAGATGATGACAAAAGTTGTTGTATTAGAGAGTGAAGCTTATCTGCATGCATGTATGGTGAGCAGGTCATTTTATTTTGTCAGTGTTGAAATATGGCACTGAAATATAAATTTGTTAATTTCATATTTTAATATGGATATCTTTGTAGTTAAAAAAAAATGCTCAAGACAGAAAGTGAGATGTCTTTAGTCGTGACATTTATTAAT
ATTGTTTACTGTACATTTCTCTACCCTTTGCAACCATACAATTGTTGAACACCTGTCCTGTATATTCAAACAAACAACTTGAACGAAGGCAAATCTGATATCTTTGGGGCAGCAAAATACATGTGCTCTTGTCTTTTGCATTTGAGTAACCTCCTTATATAAATATGGCTTACACTGATTAGCGGAACGGAATGAGCGGAACGGAACGGAACGGAACGGAATAGCCACACCAAAAAGAAA
AATAGTTTCTCTTAATTAGAGTCCATTTGTTTTCGTTCACTGAATTTTGGAGCGGAACAGCATGGAATTTCGGATCTAAAACAGAAATTTGGATTTAAATAAAATATCACTTTTCAACCCTTAAAGTAATGTATCTGTTTTTCGACGTTTCTCTTGCTGTGATGAAAATCGTGAATAAGACTACCAATAAGAATTTTATTCCATGGAAACACACTTCAGGAACTGTGGTAGAACAAGTAC
AATAAATCAAACATTACACATATAATCAGGCAAGACTGCTCCCGAGACTGCTGCTGGGAATGAATAAGACTACTGGACTGAAGCTATAATTCAAATCTGAAACTACACCTGAATCCAAACATCACCCTGGATATCGCATCTGCTGGAGGATGCTATGGCTACCATGAGGCATGTGAGATCGCGCACAGGCGTTCTGTTGAGTGGGTATAGCTTTTAAAAATGTAGTTCTGCTACGTTCCG
ACGTTAAGGCTACGGAAGAGAACGTCCGAACGTAGCATACTTTTACAAACTACCCACACAACAGAACGCCTATGAGGTGGCGGTCTGTGTAATCCTCGATGGCATGGCGCCTAATTACATTTTTGGCGCGGAGGTGAAAAAAAGAGGGAGGCGGTGACGGATCGTAACTTCTAAAATGGATACATTGAACTGCATTGCTGATTGTTTGTAATTGGCGCGGATGCCACTTCTTTTTCGTTT
TTTGTTGCCAAGGGTCTGCAGACTTACGGGAGCACAAGTTGATTTAGGGGATTGTGAAAATGGAGGTGTAGACAAATTTTTTTTCACCCATTATGACGGCATTTTTTTTCAAAAGCATGCTCGCGAAACAAATTATTTCTGCACAACTGAAGGCAATGCAATATATTTCATTTGTTGTCAAAATGAAGTGCACGTGCATATGTTTTGCATGTCAAAGTTCATTCATGCAACCTGCAATC
TTTATTGCAGAATTTTGCATATCTAACCACTTCCCACGTTAGAAAATTTTTAAATATCCAGATCTCAAAGATCAAAATACTAAGGTCGGTTCTAAACATTCATTTTTGGTAATTCCAGCACCTCGCATCACATTTTTGCCTTCCTTTCAGGGAAGTGGTAACATTTCTTTTCTCACTTTGGGGTTTTGCAGAAAATTATTTTATCATTCTCCGTGGGGGGGGGGGGGTTGACAATGTTCC
CCCTCCCGAATACTCCAGCCCCTCCTTTAGAAATCAAATGATTCTCCCCTCAATCTATCGAATGACTAAACAACTAGGTCGAAGGACATCAGTAATACATACAAAAGGCATCGCTCATACTTACATACACAGTCGATCAACGCGTTAACATGAAACTACGGCCGTTTTGCAATTTAAGTCATTCTTTTGTGATCAGTTTTTGAAGGCAGGTAGTGTTTACACCCCTAATTGTCGTTGTAT
GCCGTACACATGCAATTATTGTTTGAAAAGTATGTTGAGCTATTGCGATCGTACAAAACAGAGTGCACAACTACATGTTGGCCACAGACACGCCCTCTTTTGGGTTGACCACCCCTCCACACTATGGTACTGCCCTCTATAAGGCGTGATTCACGGGCGCGGGCTACAGGTTAGATGTAAATTTAGATGTAGCTTCAAATTGAAATTACAGCCTAGTTTCCCGTGGTATTTGACATAATT
TTCATTTTGGAAACTTCAAAAATTAAGGTGCCTTACTTGTAGAGTTGAATAAATGATATATTTTATTTAAATCAAAATGTCTCTCTTAATTCCATTCTTTTCCGCTCAAAAATTTTGTGGACGAAAAATGGCCTCTAATTAAGAAAAAACTAATTTTTTATTTGGTGTGGCTATTCCGTTCCGTTCCGTTCCGTTCCGCTCATTCCGTTCCGCTAATCAGTGTAAGCCTATAAATATGAT
CGAAAGAGATTGTCAACATGGAGCAGCAAATCCGTTTCTTTAACCTAGACTGTAAATCAATACAAAATGGGTGCACATGTAGCTAGGGGAAAAGGTAATACACAGATAGAGCCCCATTGATGACTGCTGGAAATGTCATTTGGATGTCTTGCTGGTGTACATGCAACTGAAAAAATCATTTTAGCCTGTGGGCCGTACAATAGCTGCACTTCTACTTCGCTCACTCTAATTTGAATCTTC
GTATATAGTGTACATGGGGATCGACTTTCATAGTATGCTTGTTGGATCAAGGGACAGCGATCGACCGTGGCTTGTGTTACAGTTGAGAACGATCACATCGAGAATTTTCTTTTTCGAATGGGCGATTACAAGCGCAAATCTTTGGAAGTTAGCTACCATATAAGAACTTTGTTGTTTTAACTTCAAGAATAATCAATCTAAAACACGAACAGAATGTTCTGTGACTTTTGAAGCTACATT
GAACTGACGACCTACCGGGACTATCGAGCATGGCAGCTGCCATCTTGGATATTGTTTACGGCTCACTGAGCATGCTCAGATCAAAAATATTTCCCACAACCCATTGCGGACTTTCTACATTGTAAACATAGGAGATGTTACGGTGTGCGTGGGAATTCTCGATGCCTTCGATTCTCACACAGTCACTACAGCACATGCACAGTAGCCGCCATTGTGTCTAGCTGACGCAGTTACGCTCGA
TTTGCAGTACGGTGTCTTACGTTACCTCGCAGTAAGTACACATCTTAAAGTGACATTATGGCCGCTTACCAAATACCTGCACCTCAGCCTATGAATGTTAAAGGCGATGTGTCCACAAATTGGAAGAATTTTCAGGCTGCATGGGACTATTTCAACATAGCAACTGAACTCAAAACCAAATCTAAAGAGGTACAGGCAGGCGCACTGTGTAGTGTTATGGGGATGGATTGTGCAAAAGTT
ATGAACAGTCTAGACACTTTGTCCGCAGATGATAAAAAAGATCCAGCAAGAATTCTAAAAGAATTAGGCGATCATTTTATGCCTCAAAAGCATTTGTTGTTCGAGAGAGTTAAGTTTGGATCAGCCAACCAAACTGATGCCGAATCTGTCGATCAGTATGTAGTACGACTACGTCAGCTTGCAGAGTCATGCGAGTTCGAGACATTACTCGACAGTCTAATCAGAGACAGATTGGTTATG
GGTACACATGATTCAGCAACCAGAGATAGATTACTGAGAGAACGTCCTGTACCTAATTTGCAAAGATGTACAGAAGCTTTGCGAGCGAATGAACTGTCCAAGGTACATAAGGAACAACTACGCAGCACAGCAGAAGTACAAGATACTGCACATGTAGCACGTAAGCAGAATACACATGGTAAAGCAAACATCAAGGGGACTCGCCCAAAGCATGCCCGATATACAGATGGAAACACACCA
AAGTCTAGTAATCAGTGCAGGTACTGTGGCACATCTCATGCCAAGGACAAAGCCAAATGTCCAGCGTATGGGAAAAGATGTAATAAATGTGGCCAGTCTAACCACTTTGCAGCTGTATGCCAAGGGCAGAAGAGAATATTGAGGAAAGATGCAAAAAAAGTTCACCAGATCTCAGGATTTGAGAA

#### FASTA protein

**RAG2L-A**

>PflHRAG2L-A.23
MAAAMLDSPEIQITKNILATCLATAGCTNPVISLGDRALVWVKSKTTVTGWLVRDKDDTIQEIKNYSSQSNYPPHVHGSNLCLVEPNHYVLLGGLTDSEMHNTKLWQLSVFVDPRGKISLTWSQGQYSGEETHAFGHSVIKFGDSLYLYGGLQYTQKLSHAFYNVSMASNIFSVLNLQSMKLSSISLQAITPRAYHSVFLNSSKMYIVGGLYVIGNNVR
HPPDVDVIDLNSMVCTSVPFIGTPNPVSFKCNDTVYILSNNTVYKSPCDAIDFIETAHYDTLSSKYYSLPFFSSIAVCDTVKKQMIKFSFQNEGTSSETRTQSNNSQLGDEEERSTDEDRVSIAEEEQERISIDMTDQESTGSETEDQNGISEAAEVKVSTDRNESEEGSRAGKIDNTSDESEEETWECMLDGCKFANLAATEQAKFNWLKCDRTYIMN
NQTVLCGRWYHDKCVELDHLNRTKLDCINFECPSCFTTCANINCSVTDKSYFSTVRCTQCVRAFHIKCSGKRFSSRQLENKSNIKFTCSDCNF

### RAGL-B

#### Pfl[H]RAGL-B.1

| Genomic loci | RAG1L status | RAG2L status |
|---|---|---|
| Pfl[H]RAGL-B.1 | potentially functional (2 CDSs) | potentially functional (2 CDSs) |

##### FASTA DNA

>PflHRAGL-B_01
TCAGGGTTGAATAGGTGTCTGCTTTTAGAAAGCCTATCATATATAACAAAAATACACAACCCCTCAATTTAGAACAGGTCATATGAACTGCTTCATTATAACTTTCAATTTTAGCATTTATACCAGAAATGCGTTCATCCAATCTCGGTAGAATACTTGACACAGCAACAAATGCTTTCGGGAATTTATTGAACATTTTGTTCAATAAAGTATCAAAAC
TGGTGTGACATGATTTGACATCTGAGTCATTTATAAGGTCATTTATACCAACATGAATAATAACTGTCCTTACATCATCACAATTCTCTACATTATCTACAAATATTTGTGCATCAGCCAAGGTTGGAGTAAACACCTTCTTAACACGTCTATAAGGATACATCTTAGTTGGAGACAGTTTTTTGTAGAATAGAGTCACCAATCACAATGGTATCTATA
CCACTACAGTCTATAGTGTTACCACTGTTTAACAAGCTTGACTTTGTCTTGTGTACAATATTATCAACACCATTTTGAGACTCACCCAGAATATTATGTTCATCATTCTTTGTCCCGTCTATTTGTCTAACACTTTGTAATTCATCTTTGTGAATATTTGTTACATTGTTTTCATTGACCTTTTGTTCTTCATCTACAATGTCAAACACCTGCACCTGG
TCCTTTGAATTTATCTCAGCACAAGACTGTGAAACGCCACTCTCATGCGCACGAACATTTGTGCTGATGAAGCTCTCAAATTTTCTCTCCCACTCAGACACAATTAAACTTAATTTCTCTTGTAAATTCACATCAAAGACTCTTTGCATTTCTACTAATTTGTCCGACCAGTTCTTTGTTTGATTCTTGAAGTTCAGTTTCAGCGCGCGATTTTCCGAT
CTAAGTCTACACAGTTCGTCTTCAAGACGACCCACTTTTTGTACAGTACATTTCACACTGTCCAAGTCATTTTTACACGGGTGTACTTCCTGTTGTACGTAATTCAGAACAACCGTAGATTCTACACTTATAGAGCAGCTATTGAAATGTAAATGAGCGCCATCTTGCTAATTATAAATTATAATGACTGTATGATGTCATTCTGGCACAGCGCCCCCA
TCGACAACGGCAGTTTGCCTTGGAAGGCCAGCGGAGAAAAACTGTCCAATAGCAAGGAAGGAAATGCGTGCGCATGCGCATTCGCATTTCCAATACCCAGAGCAGCTTCGGAGCTTGTTTTTTGCCGAAAACCGCATGAAAATCGACGAATTTGCCGGTAAGCTGTGGTTTTTTCTGTCCCGATCTGTTTTGCCAGAAATTTTTGAAGTTTGAAGC
CAAAAATGTAGGGTTTTGTATTCATCGCTGCCTTTTGTGCCGATTTTTTACCATAAACAATTGTTTTTCCGAGGTCATCTACAGTCTAAGCTATCGTTTGAGCATCGTTTCAAGATCGAACTCGCCGGATTTCTTGGCGAACTCGGTTTTTATTCATGACCATGTTTTTATGTTTTCTACAATTTAGATTCTCCACGTTACCACAGAAGGCATGGAA
TATCATTCATCCTGTTTGCAATCTCTTTGTCGCGTGTGTAGTAAACACATTGGGAATAATGCCACTCAAGCAGATGTATATGCTGCAGAACTAGACAAGTTATATAATATCAAAGTCGGCGACGATGATATCCAGGTACTGCTGATCAAATTCATAAGCAAAAGTAATATTGTGAAAATTTCTTAAGATAAAGTCCAGTACGTGCAGTGCAGTGTATCT
TTTGTATTTCTGTGCTGATTATTTTGGCTAAAACAGGCAGATGCACATCAGGTATAATGCATTATAATGATAAAATACAGTTTTCCTGATAATATGTGTGCAAGCTTGTTATATCAATGGTACAGAAATAAGAAATAGGTGTATATGTTTGCGCAATTATTTCAGGTCACTGTTGAAAATCATTTCATGCAGACATTTTTTGTAACATGGTGATTTATTG
TTATTGTATTTACAGATCATCCAACTAAACTCTGTAAAGCATGCACATCCCGCTTGCAAAGATGTAGAAGGTCAGCTGCAGACTTCCAGCCAATGCAAATAGAGGTTGCGACGTTTGTGCCACACCAAGACCAAGGCTGTGTAGTTTGTGATCAGCCAGCACAACTTGCTACAGGCACGGCTCCAAAGGCTAAAGGAAGACCACCGAAAAGAAAATGT
GGTGGAGGTCAGTATGGAGGGCCAGGTAGAGGTCACAAAAAGTCAAGCAGTGATATACCATTGTCCGGTAGTGATATACCGTCGACCAGTAGTAGTATACCATCAACCAGTAGAGGTATACCGTCGACCAGCAGTAGTTCCGAGCTAACAGTTGAATCAGTCAAGAGAAAACTTTTGACCATTGAAAGCCCTGAAAAGGAAGAGGCCATCACAGAAGCG
ACAGAAATTACTAGCATACCGCTGGACAGATTTATAGAGAAAGACATTGCCGAACACTATGTTTGTAGTATTTGTCAGGGCGTCCCAACCACACCATGCATATCACCCTGTAGCCATATTTTTTGTGTTGGTTGTATACAGCAGTGGTTGGCAAATTCATGTGCGTGCCCAAGTTGTAGAGAGATTCTTGAATGTGATGACTGTCAAAACTTAACAGGA
AATCATCTCAACATATATGATAGTCTTAGACTTAGATGCACTTATAGTCATCTTGGTTGTGAAACCATGACTACTTTACCTAATTATATTGACCATGAGTTAACATGTAAATATAAAGCAAAAGGTAAGCGCTCAACATATGGTAAAACAAGAGTAAAACAATCTCTAAGGACAGCAGATAGGCAGTACTGCAAACAAAAAAGGTTGAAAGCTTGTTAT
GATTTTCTCCGAGATTTTTGTACTGCTAACTCAGAGTCCACAGAAGATGTTTTGTTTTTTCTACTCAGATCATATCTATATGACTCTGGTGACAGAGAACGTTCCAATCTTGTGGATGATCTTTGGTCTAAAACACAGAGCAAAATGTCAGCTGATGAATGTTTAGCACTAAGAATAGACAACCTACAAACTAAGAACCGATACAAAGCCCAGTATGAT
ATGTTCAAGAGCAAATCTGTGAGTGTGCTGGTCGCACCAAATCAGTTAGATATGCTAGAACGAACATACATGCCTGGTACTGCAAGGTATGCTATTGTTGGAGAGGAAAATTTTGAACATATATACCAAACTCCAGTAAAATTGCATAGAAAAGACCTTTCCATATCTACTACTGCAGATTCTGTTGAACCTATAGAGTTAAACAGTGAATATCCCAAT
GATAATAAAGAATTTCCAGGTCCAAATTTGGCTGGGGTAAGGTTTAGATACACTGATGCAGTAGCAAAAACACTTGAGGAAATTGAACCTGAAATAAGTGAGAATCTCAAAAGTATTGGTGTGAGCAGTACAAAAACAGAATTGGTCTTGCGAACATTTATCAAAGATGGTTCTGATGGGATGGGGGACGTTGAAATTCATCGGCGCAAGGGGGAGG
ACACTTCCAGCCAATGCCTTCAGAGCTGCATTTGCAGTTGTAAAATGTGAAGTAGAAGTAGGCAATGAAATAAAGACAGTCTGGGACCAAAAGAATCCAAATTCAGTCCGATGCAATAGACCACTAATTGAGGCAATCGCTGAGGAAAATAATGATTCCACAGTTCACTACTGTCTACTGACAATGGAAGGTGAAAGAGAACTTATGAAAAACAAAGTG
ATGAAAATTGACTGTGGTGATTATTGGAGATGTCATTATTTAGTATTTGTTACTTCCATGGTTGATGAAAAGTTAGATAGAAGTGCAGGGGGGTTGCAGGGTGCTGGCTCAGGTTATCCATGTACTCTGTGTGATTGCACCCGTGAAGAAGCAATTTCTAAATTGGGGTCTTTTTCCATCACTAGAAAGAGGACTGAAATAATTGAGAAAGCTGAGATT
CGCAGAGTAAACCCAAAGATTCTTTCACAAAATGAATTGAATTTGAGTTGTAAAGGTGTAAAAAAACATCCTCTTTTGCTTTCTGAACCAGTCGAAAGGGGCATAGATTCAACCCATGCTAACATAAACCTTGCTTCTTTTTTTAAGAAAGTGTTAGTAAGGGAAGTAGCAGAGATCACTCAATGGGAGAAGACTGCAGAATTAAAGTCCAGTCTTGAT
ATGGCTGAGAAAAGGTTAGATGACCATTTGAAGGCTGAAATAGGGATTAATCCTAGCCTAATGATGCCTGGCAACTATGCTAGGGTGTTCTTTGATGAGAAAAATGAACAGGCAATACTCTCCCTAATACCACAAGCTCAAAGACGTGAGGATTTTGCTGCTGTGCTAGCTAAGTTTCGATTTTTGAAGAAGGTGTATTGTGCTAAATTGCCCAAAGTT
GACTACAAAGATGATATCGAAAGTGTGAAAACTGTGGGAATTGAAATGGGCATGCTCCTAATTGATAAATTTGGATATGCAAGATGGCCAAATTACCTTCATAAAGTCATAGAACACACCCAAGAGCTCATTGAAAAGGAAGATAGCCCAGGTACCATTGGTGGAATCTCAGGGGAAGGTAATGAAGCAGGAAATAAATTATTCCGTCAGTTCCGGAAA
CTGCATTCCAGGAAAGGGTCTGTAATGGGTGGGTTGCGCGACACAATTTGGTTGCACTGGCTGTATAGCAGTCCAAAATTATGTCGTCATGCTGAGGTAGCACACAGGAAAAACAGGTGTTCTGCTTGTGGATGTCTAGGGCACAATCGTTTGACATGTTCAAATGAGTCTTGA**GAATGCAGCAGCATTGCAGAGTATGGCAGTCATTGGTAGAAGTTT**
**CAAATGTATTATTTTGAAATAAGTTGAATAATGTTATCATTTGTAACAGCCACTGACCCTTGAAATACAGGGGATGCTTGTGAAATACTTTTAAAAAATAGCACTTTGAGAAAAATATTTTATGACAAACCTCAAATATATATTTTCTAAAGATACTGCTGTATCACATTTATTTACAAACTCTGAAAGCAATATTTCATCATAAAATGTTGATGAAAAA**
**ATAAATAAATGAAC**AATAAATAAAAATCTTTCTGCTTATTTTTGTCTGTTATTAATTTAATGTTTCTTCAATTTTCCTAGACCGCTTTCTCCCCCCAGTGGACTTGCATGAATTGCACACATATTTGCCCTTTGGAATGACTTCAAGTTTAATGCAAAATTGGTGCAGCCACCTTTTACATTTGCCCTCACACTGAATCCATGCGATGGGGCTGATCT
CTGGAGTATCTTTTATGGTACACTCATCCTCTAAATCGCATGGAGCTGGACACATGGCCTTACTTGTAAATACAAAAATATTTTTGTGGGTTCCCCCCAATCCCATGATTGCTGTGTTGTCTAGGGACACCATTGACAAACCAGCAGAGGCAAAGCTATCTGGTGCCTCAAGTCTTGTCAGTACCTCTGAAGTTGTGTTTAACAAGAACATGGATGAAC
TTGGTTTGGCTGGCCTTGCATTTGATTGATATAATGCGCATGGTTGTAACATATCCCCCATAGACAATCACCTGGTCATTGAACTGAAGGGCCCCATGCATTGATACATGGTAAGGCAGAGACTCAGATAAAAATACAGATTTAACAGCGTATTCATTGGTTGCACTGATACTAAGAAGTGTCACCTCGTTGATCTGATGTCGCTTATAAGGCTTTGGT
TCTCTGTATGTCACCCCCCCTACATAGCAAATGGTTGAAGAGCTTGAGAGGTAAGTGGCAGTGTGGAAAGCTCTGGGTTCTAACTGTGGCACATTCAGCTTTTTCCAGGACAAAGAGTCAGTGTTCAAGAGATAAAAGTGACCATCCTTGGTATCTTGCTCAAAACTTGGTACTCTACGAAACCTGTTTGGCATTTCTGCCCCGCCCAAACAGAACAGCAG
CATGTTGTCCTGGGATAGGTGTTAGAGTGTCAACCACAACGAGCTGACGGAATATCTCCTGACTGATGTTTCTCATCTGGGAGGTGGAAGCTCATATCATTTGGATCTTTGTAAATTACGCAGGTGGCAGTTTTCCTTTGTATTTCCATCGTAACTGCCTCGTTGGTAACAAAAGAGTGAACCCAAATGGTATCACCCCCACACCAACAACTTGACTTTGT
CAGGGGTACTGATGTTCAGCATCGAAGCAGCATGCAGTGGTGACAGCATTGCTCCCCTCGTGGTAAATTTCTGTACGCTTTCCACATCAACATCAGTGTCATCTAAAGTAAATGACAATGAGTACAGCTCATTGGAAAGAGACCATGTACTCTCCCTCTTTCCACCTTCCACCACCAAGTGTGTAGCTGTTACCACACCATCACCTACATGAAAAATAC
ATTTATACAAACACATATAACCACCTTATTTTTTAAATGTATTAAATGTTTTAAAAAAATACTTTTTTGTAAAATATATGTAGTTTTAATTTTCTCATTATTTGAACTTTGTGATTGATATTTGACATGACATGATATATGTAGAAAATGATTAAATTACCAAGAAACTATTTCCCTTTTCACAAATGTTTTTGAAAAAATGCACAAAGTACTGTGTAC
TACAGTGTAGTGTGTTACAATACTGGACTTACAGAAAACTTTTTAACGCAAACATCTTTTCAACCTAACTTCAATTGAAAAAGGGGCAGACAGAAAAGAACAGCAAAACAATGTGTGTAAATTACACACTCACATACGCTGTCACAGACATATAAAAACAAAAAAAGACGGCTTTGTCTTTGTAAAATCCATTCACGAAAAGTTACCCACGCGTAAGTGCTTGCGTAACAATGCA
GAGATCTAGAGAACAATAGGATTTAATCAGTGTCCAGCAACAAGTGCAGTTTAACACATATTTATGCTTTACTCACCATTATTTATGGATACATTCATATGTCCTTCGGGGGGGAAGTATTCGCCCAGTTCAAAAAATCTTTTTCTAGTCATTTTTCTCTTGTCACTGGAGCCTTCCGTGAGCGGAATAAAGCGAAAGACATCGAGAGCGAGTTCAGGA
GCAACAGCCATTTTATTAGCATATGAAAGAAAATGCCGTGACCTTTAAAAATGACAACATTGGTGACGTCAAATTTCTCGCGCGATCGTACGGACTTATACGAGAGTACAAACTGCGCATGCGTCAGGCGGCCATGTTGATTTTTTACGATTCGCTATCGGTTACCGGTAATTGATGTAAGTGTTCTATGATCTTTTCTTCCAACACAGCTATCGTACA
ATCTAACCTGGTCCGCACTTTCAGTTGTACCCGAATCTTTTTAACGCAAACATCTTTGGCATCGGTACTGCAAATTTTCCCTGCATATCGGTACTGTCGCATCAGTACAGGGTCGAGTAAGCCCCTCGTCAGCTGTTGACAACTTTTCTTTATCATTACTCTCTCCGTCGACTAAGCTGTACACATCAAAACACTTGGTGTTTATATTTCATCATTT
ATGAACAACCTTTTCACAACAGTGTCAGATTCGGATTTGATAAACTTTCAAAGTCATCAGATGTGACACGCGTCGTATCTGTATCCGTACAGTCATCATGGCCGCCATCGAGAGAATCTACGTCGCAGTCGGAAGCTCAATCCGTTACCTGATTCTGATCGTAGAATTGTTGTGCTTCAGCCTCTGTGTCAAAACCTTTGAAGCAGTTTCCAGGGTATC
CGTTGGTAACTTCATGACATGACTTCCAGTCACAAAAAATACCAGTCGACCTCCCAACAGTTACGGCATAGAACTTTTTTCGTTCCTCGCGTACGCGTTGGTCGCTTCATTGTCAGCTTCACCACTGTAGCGTCATCCCGTACATCTTTATCTTCCGAAAGTAAATGTCGAATTCTTCACCATATATAAAGCCCTCACTGTTATTTAACTTTCTCTCG
CGCTTCTCTGGCGTATGCATTTCGTTCCAACAAAAATTGACTTGTGCAATGAACTAAATCTGCTAAAATCACTGTCAGAATGCAGAGCTGATGAAACACGTGTACACTCTATACTGCCGCCATTGTATAGACTATTGTATTTAACTACCCAAATCTTATGAATGAACCAGTAATAGCAAACTTTATGTCATCTCTCGTTAAGTCAGGTACAGTAGAAC
CTATGAAACATTCATCGGCAATGCCAGTATCAGCCTTCCATGAGCTGTTTATGTCATGGCCAGACAATGAAAAACTGAGTATCAAACA

##### FASTA RNA

**RAG1L-B**

>GDGM01438088.1 TSA: Ptychodera flava comp266024_c0_seq4 transcribed RNA sequence
CCCAGAGCAGCTTCGGAGCTTGTTTTTTGCCGAAAACCGCATGAAAATCGACGAATTTGCCGATTCTCCACGTTACCACAGAAGGCATGGAATATCATTCATCCTGTTTGCAATCTCTTTGTCGCGTGTGTAGTAAACACATTGGGAATAATGCCACTCAAGCAGATGTATATGCTGCAGAACTAGACAAGTTATATAATATCAAAGTCGGCGACGATG
ATATCCAGATTCATCCAACTAAACTCTGTAAAGCATGCACATCCCGCTTGCAAAGATGTAGAAGGTCAGCTGCAGACTTCCAGCCAATGCAAATAGAGGTTGCGACATTTGTGCCACACCAAGACCAAGGCTGTGTAGTTTGTGATCAGCCAGCACAACTTGCTACAGGCACGGCTCCAAAGGCTAAAGGAAGACCACCGAAAAGAAAATGTGGTGGAG
GTCAGTATGGAGGGCCAGGTAGAGGTCACAAAAAGTCAAGCAGTGATATACCATTGTCCGGTAGTGATATACCGTCGACCAGTAGTAGTATACCATCAACCAGTAGAGGTATACCGTCGACCAGCAGTAGTTCCGAGCTAACAGTTGAATCAGTCAAGAGAAAACTTTTGACCATTGAAAGCCCTGAAAAGGAAGAGGCCATCACAGAAGCGACAGAAA
TTACTAGCATACCGCTGGACAGATTTATAGAGAAAGACATTGCCGAACACTATGTTTGTAGTATTTGTCAGGGCGTCCCAACCACACCATGCATATCACCCTGTAGCCATATTTTTTGTGTTGGTTGTATACAGCAGTGGTTGGCAAATTCATGTGCGTGCCCAAGTTGTAGAGAGATTCTTGAATGTGATGACTGTCAAAACTTAACAGGAAATCATC
TCAACATATATGATAGTCTTAGACTTAGATGCACTTATAGTCATCTTGGTTGTGAAACCATGACTACTTTACCTAATTATATTGACCATGAGTTAACATGTAAATATAAAGCAAAAGGTAAGCGCTCAACATATGGTAAAACAAGAGTAAAACAATCTCTAAGGACAGCAGATAGGCAGTACTGCAAACAAAAAAGGTTGAAAGCTTGTTATGATTTTC
TCCGAGATTTTTGTACTGCTAACTCAGAGTCCACAGAAGATGTTTTGTTTTTTCTACTCAGATCATATCTATATGACTCTGGTGACAGAGAACGTTCCAATCTTGTGGATGATCTTTGGTCTAAAACACAGAGCAAAATGTCAGCTGATGAATGTTTAGCACTAAGAATAGACAACCTACAAACTAAGAACCGATACAAAGCCCAGTATGATATGTTCA
AGAGCAAATCTGTGAGTGTGCTGGTCGCACCAAATCAGTTAGATATGCTAGAACGAACATACATGCCTGGTACTGCAAGGTATGCTATTGTTGGAGAGGAAAATTTTGAACATATATACCAAACTCCAGTAAAATTGCATAGAAAAGACCTTTCCATATCTACTACTGCAGATTCTGTTGAACCTATAGAGTTAAACAGTGAATATCCCAATGATAATA
AAGAATTTCCAGGTCCAAATTTGGCTGGGGTAAGGTTTAGATACACTGATGCAGTAGCAAAAACACTTGAGGAAATTGAACCTGAAATAAGTGAGAATCTCAAAAGTATTGGTGTGAGCAGTACAAAAACAGAATTGGTCTTGCGAACATTTATCAAAGATGGTTCTGATGGGATGGGGGACGTTGAAATTCATCGGCGCAAGGGGGAGAGGACACTTC
CAGCCAATGCCTTCAGAGCTGCATTTGCAGTTGTAAAATGTGAAGTAGAAGTAGGCAATGAAATAAAGACAGTCTGGGACCAAAAGAATCCAAATTCAGTGCGATGCAATAGACCACTAATTGAGGCAATCGCTGAGGAAAATAATGATTCCACAGTTCACTACTGTCTACTGACAATGGAAGGTGAAAGAGAACTTATGAAAAACAAAGTGATGAAAA
TTGACTGTGGTGATTATTGGAGATGTCATTATTTAGTATTTGTTACTTCCATGGTTGATGAAAAGTTAGATAGAAGTGCAGGGGGGTTGCAGGGTGCTGGCTCAGGTTATCCATGTACTCTGTGTGATTGCACCCGTGAAGAAGCAATTTCTAAATTGGGGTCTTTTTCCATCAGTAGAAAGAGGACTGAAATAATTGAGAAAGCTGAGATTCGCAGAG
TAAACCCAAAGATTCTTTCACAAAATGAATTGAATTTGAGTTGTAAAGGTGTAAAAAAACATCCTCTTTTGCTTTCTGAACCAGTCGAAAGGGGCATAGATTCAACCCATGCTAACATAAACCTTGCTTCTTTTTTTAAGAAAGTGTTAGTAAGGGAAGTAGCAGAGATCACTCAATGGGAGAAGACTGCAGAATTAAAGTCTAGTCTTGATATGGCTG
AGAAAAGGTTAGATGACCATTTGAAGGCTGAAATAGGGATTAATCCTAGCCTAATGATGCCTGGCAACTATGCTAGGGTGTTCTTTGATGAGAAAAATGAACAGGCAATACTCTCCCTAATACCACAAGCTCAAAGACGTGAGGATTTTGCTGCTGTGCTAGCTAAGTTTCGATTTTTGAAGAAGGTGTATTGTGCTAAATTGCCCAAAGTTGACTACA
AAGATGATATCGAAAGTGTGAAAACTGTGGGAATTGAAATGGGCATGCTCCTAATTGATAAATTTGGATATGCAAGATGGCCAAATTACCTTCATAAAGTCATAGAACACACCCAAGAGCTCATTGAAAAGGAAGATAGCCCAGGTACCATTGGTGGAATCTCAGGGGAAGGTAATGAAGCAGGAAATAAATTATTCCGTCAGTTCCGGAAACTGCATT
CCAGGAAAGGGTCTGTAATGGGTGGGTTGCGCGACACAATTTGGTTGCACTGGCTGTATAGCAGTCCAAAATTATGTCGTCATGCTGAGGTAGCACACAGGAAAAACAGGTGTTCTGCTTGTGGATGTCTAGGGCACAATCGTTTGACATGTTCAAATGAGTCTTGAGAATGCAGCAGCATTGCAGAGTATGGCAGTCATTGGTAGAAGTTTCAAATGT

ATTATTTTGAAATAAGTTGAATAATGTTATCATTTGTAACAGCCACTGACCCTTGAAATACAGGGGATGCTTGTGAAATACTTTTAAAAAATAGCACTTTGAGAAAAATATTTTATGACAAACCTCAAATATATATTTTCTAAAGATACTGCTGTATCACATTTATTACAAACTCTGAAAGCAATATTTCATCATAAAATGTTGATGAAAAAATAAATA
AATGAAC

>GDGM01014902.1 TSA: Ptychodera flava comp266024_c0_seq2 transcribed RNA sequence
GGTATATGCTGCATCAAGCAGATGTATATGCTGCAGAACTAGACAAGTTATATAATATCAAAGTCGGCGACGATGATATCCAGATTCATCCAACTAAACTCTGTAAAGCATGCACATCCCGCTTGCAAAGATGTAGAAGGTCAGCTGCAGACTTCCAGCCAATGCAAATAGAGGTTGCGACATTTGTGCCACACCAAGACCAAGGCTGTGTAGTTTGTG
ATCAGCCAGCACAACTTGCTACAGGCACGGCTCCAAAGGCTAAAGGAAGACCACCGAAAAGAAAATGTGGTGGAGGTCAGTATGGAGGGCCAGGTAGAGGTCACAAAAAGTCAAGCAGTGATATACCATTGTCCGGTAGTGATATACCGTCGACCAGTAGTAGTATACCATCAACCAGTAGAGGTATACCGTCGACCAGCAGTAGTTCCGAGCTAACAG
TTGAATCAGTCAAGAGAAAACTTTTGACCATTGAAAGCCCTGAAAAGGAAGAGGCCATCACAGAAGCGACAGAAATTACTAGCATACCGCTGGACAGATTTATAGAGAAAGACATTGCCGAACACTATGTTTGTAGTATTTGTCAGGGCGTCCCAACCACACCATGCATATCACCCTGTAGCCATATTTTTTGTGTTGGTTGTATACAGCAGTGGTTGG
CAAATTCATGTGCGTGCCCAAGTTGTAGAGAGATTCTTGAATGTGATGACTGTCAAAACTTAACAGGAAATCATCTCAACATATATGATAGTCTTAGACTTAGATGCACTTATAGTCATCTTGGTTGTGAAACCATGACTACTTTACCTAATTATATTGACCATGAGTTAACATGTAAATATAAAGCAAAAGGTAAGCGCTCAACATATGGTAAAACAA
GAGTAAAACAATCTCTAAGGACAGCAGATAGGCAGTACTGCAAACAAAAAAGGTTGAAAGCTTGTTATGATTTTCTCCGAGATTTTTGTACTGCTAACTCAGAGTCCACAGAAGATGTTTTGTTTTTTCTACTCAGATCATATCTATATGACTCTGGTGACAGAGAACGTTCCAATCTTGTGGATGATCTTTGGTCTAAAACACAGAGCAAAATGTCAG
CTGATGAATGTTTAGCATTAAGAATAGACAACCTACAAACTAAGAACCGATACAAAGCCCAGTATGATATGTTCAAGAGCAAATCTGTGAGTGTGCTGGTCGCACCAAATCAGTTAGATATGCTAGAACGAACATACATGCCTGGTACTGCAAGGTATGCTATTGTTGGAGAGGAAAATTTTGAACATATATACCAAACTCCAGTAAAATTGCATAGAA
AAGACCTTTCCATATCTACTACTGCAGATTCTGTTGAACCTATAGAGTTAAACAGTGAATATCCCAATGATAATAAAGAATTTCCAGGTCCAAATTTGGCTGGGGTAAGGTTTAGATACACTGATGCAGTAGCAAAAACACTTGAGGAAATTGAACCTGAAATAAGTGAGAATCTCAAAAGTATTGGTGTGAGCAGTACAAAAACAGAATTGGTCTTGC
GAACATTTATCAAAGATGGTTCTGATGGGATGGGGGACGTTGAAATTCATCGGCGCAAGGGGGAGAGGACACTTCCAGCCAATGCCTTCAGAGCTGCATTTGCAGTTGTAAAATGTGAAGTAGAAGTAGGCAATGAAATAAAGACAGTCTGGGACCAAAAGAATCCAAATTCAGTGCGATGCAATAGACCACTAATTGAGGCAATCGCTGAGGAAAATA
ATGATTCTACAGTTCACTACTGTCTACTGACAATGGAAGGTGAAAGAGAACTTATGAAAAACAAAGTGATGAAAATTGACTGTGGTGATTATTGGAGATGTCATTATTTAGTATTTGTTACTTCCATGGTTGATGAAAAGTTAGATAGAAGTGCAGGGGGGTTGCAGGGTGCTGGCTCAGGTTATCCATGTACTCTGTGTGATTGCACCCGTGAAGAAG
CAATTTCTAAATTGGGGTCTTTTTCCATCAGTAGAAAGAGGACTGAAATAATTGAGAAAGCTGAGATTCGCAGAGTAAACCCAAAGATTCTTTCACAAAATGAATTGAATTTGAGTTGTAAAGGTGTAAAAAAACATCCTCTTTTGCTTTCTGAACCAGTCGAAAGGGGCATAGATTCAACCCATGCTAACATAAACCTTGCTTCTTTTTTTAAGAAAG
TGTTAGTAAGGGAAGTAGCAGAGATCACTCAATGGGAGAAGACTGCAGAATTAAAGTCTAGTCTTGATATGGCTGAGAAAAGGTTAGATGACCATTTGAAGGCTGAAATAGGGATTAATCCTAGCCTAATGATGCCTGGCAACTATGCTAGGGTGTTCTTTGATGAGAAAAATGAACAGGCAATACTCTCCCTAATACCACAAGCTCAAAGACGTGAGG
ATTTTGCTGCTGTGCTAGCTAAGTTTCGATTTTTGAAGAAGGTGTATTGTGCTAAATTGCCCAAAGTTGACTACAAAGATGATATCGAAAGTGTGAAAACTGTGGGAATTGAAATGGGCATGCTCCTAATTGATAAATTTGGATATGCAAGATGGCCAAATTACCTTCATAAAGTCATAGAACACACCCAAGAGCTCATTGAAAAGGAAGATAGCCCAG
GTACCATTGGTGGAATCTCAGGGGAAGGTAATGAAGCAGGAAATAAATTATTCCGTCAGTTCCGGAAACTGCATTCCAGGAAAGGGTCTGTAATGGGTGGGTTGCGCGACACAATTTGGTTGCACTGGCTGTATAGCAGTCCAAAATTATGTCGTCATGCTGAGGTAGCACACAGGAAAAACAGGTGTTCTGCTTGTGGATGTCTAGGGCACAATCGTT
TGACATGTTCAAATGAGTCTTGAGAATGCAGCAGCATTGCAGAGTATGGCAGTCATTGGTAGAAGTTTCAAATGTATTATTTTGAAATAAGTTGAATAATGTTATCATTTGTAACAGCCACTGACCCTTGAAATACAGGGGATGCTTGTGAAATACTTTTAAAAAATAGCACTTTGAGAAAAATATTTTATGACAAACCTCAAATATATATTTTCTAAA
GATACTGCTGTATCACATTTATTACAAACTCTGAAAGCAATATTTCATCATAAAATGTTGATGAAAAAATAAATAAATGAAC

>GDGM01481817.1 TSA: Ptychodera flava comp266024_c0_seq1 transcribed RNA sequence
CTACAGGCACGGCTCCAAAGGCTAAAGGAAGACCACAAAAAAGAAAATGTGATGGAGGGTCAGTATGGAGGGCCAGGTAGAGGTCACAAAAAGTCAAGCAGTGATATACCATTGTCCAGTAGCGATAAACCGTCGACCAGTAGTGGTATACCGTCGACCAGCAATAGTTCAGAACTAACAGTTGAATCAGTCAAGAGAAAACTTTTGACCATTGAAATC
TCTGAAAAGGAAGAGGCCATCACAGAAGCGACAGAAATTACTAGCATACCGCTGGACAGATTTATAGAGAAAGACATTGCCGAACACTATGTTTGTAGTATTTGTCAGGGCGTCCCAACCACACCATGCATATCACCCTGTAGCCATATTTTTTGTGTTGGTTGTATACAGCAGTGGTTGGCAAATTCATGTGCGTGCCCAAGTTGTAGAGAGATTCTT
GAATGTGATGACTGTCAAAACTTAACAGGAAATCATCTCAACATATATGATAGTCTTAGACTTAGATGCACTTATAGTCATCTTGGTTGTGAAACCATGACTACTTTACCTAATTATATTGACCATGAGTTAACATGTAAATATAAAGCAAAAGGTAAGCGCTCAACATATGGTAAAACAAGAGTAAAACAATCTCTAAGGACAGCAGATAGGCAGTAC
TGCAAACAAAAAAGGTTGAAAGCTTGTTATGATTTTCTCCGAGATTTTTGTACTGCTAACTCAGAGTCCACAGAAGATGTTTTGTTTTTTCTACTCAGATCATATCTATATGACTCTGGTGACAGAGAACGTTCCAATCTTGTGGATGATCTTTGGTCTAAAACACAGAGCAAAATGTCAGCTGATGAATGTTTAGCATTAAGAATAGACAACCTACAA
ACTAAGAACCGATACAAAGCCCAGTATGATATGTTCAAGAGCAAATCTGTGAGTGTGCTGGTCGCACCAAATCAGTTAGATATGCTAGAACGAACATACATGCCTGGTACTGCAAGGTATGCTATTGTTGGAGAGGAAAATTTTGAACATATATACCAAACTCCAGTAAAATTGCATAGAAAAGACCTTTCCATATCTACTACTGCAGATTCTGTTGAA
CCTATAGAGTTAAACAGTGAATATCCCAATGATAATAAAGAATTTCCAGGTCCAAATTTGGCTGGGGTAAGGTTTAGATACACTGATGCAGTAGCAAAAACACTTGAGGAAATTGAACCTGAAATAAGTGAGAATCTCAAAAGTATTGGTGTGAGCAGTACAAAAACAGAATTGGTCTTGCGAACATTTATCAAAGATGGTTCTGATGGGATGGGGGAC
GTTGAAATTCATCGGCGCAAGGGGGAGAGGACACTTCCAGCCAATGCCTTCAGAGCTGCATTTGCAGTTGTAAAATGTGAAGTAGAAGTAGGCAATGAAATAAAGACAGTCTGGGACCAAAAGAATCCAAATTCAGTGCGATGCAATAGACCACTAATTGAGGCAATCGCTGAGGAAAATAATGATTCTACAGTTCACTACTGTCTACTGACAATGGAA
GGTGAAAGAGAACTTATGAAAAACAAAGTGATGAAAATTGACTGTGGTGATTATTGGAGATGTCATTATTTAGTATTTGTTACTTCCATGGTTGATGAAAAGTTAGATAGAAGTGCAGGGGGGTTGCAGGGTGCTGGCTCAGGTTATCCATGTACTCTGTGTGATTGCACCCGTGAAGAAGCAATTTCTAAATTGGGGTCTTTTTCCATCAGTAGAAAG
AGGACTGAAATAATTGAGAAAGCTGAGATTCGCAGAGTAAACCCAAAGATTCTTTCACAAAATGAATTGAATTTGAGTTGTAAAGGTGTAAAAAAACATCCTCTTTTGCTTTCTGAACCAGTCGAAAGGGGCATAGATTCAACCCATGCTAACATAAACCTTGCTTCTTTTTTTAAGAAAGTGTTAGTAAGGGAAGTAGCAGAGATCACTCAATGGGAG
AAGACTGCAGAATTAAAGTCTAGTCTTGATATGGCTGAGAAAAGGTTAGATGACCATTTGAAGGCTGAAATAGGGATTAATCCTAGCCTAATGATGCCTGGCAACTATGCTAGGGTGTTCTTTGATGAGAAAAATGAACAGGCAATACTCTCCCTAATACCACAAGCTCAAAGACGTGAGGATTTTGCTGCTGTGCTAGCTAAGTTTCGATTTTTGAAG
AAGGTGTATTGTGCTAAATTGCCCAAAGTTGACTACAAAGATGATATCGAAAGTGTGAAAACTGTGGGAATTGAAATGGGCATGCTCCTAATTGATAAATTTGGATATGCAAGATGGCCAAATTACCTTCATAAAGTCATAGAACACACCCAAGAGCTCATTGAAAAGGAAGATAGCCCAGGTACCATTGGTGGAATCTCAGGGGAAGGTAATGAAGCA
GGAAATAAATTATTCCGTCAGTTCCGGAAACTGCATTCCAGGAAAGGGTCTGTAATGGGTGGGTTGCGCGACACAATTTGGTTGCACTGGCTGTATAGCAGTCCAAAATTATGTCGTCATGCTGAGGTAGCACACAGGAAAAACAGGTGTTCTGCTTGTGGATGTCTAGGGCACAATCGTTTGACATGTTCAAATGAGTCTTGAGAATGCAGCAGCATT
GCAGAGTATGGCAGTCATTGGTAGAAGTTTCAAATGTATTATTTTGAAATAAGTTGAATAATGTTATCATTTGTAACAGCCACTGACCCTTGAAATACAGGGGATGCTTGTGAAATACTTTTAAAAAATAGCACTTTGAGAAAAATATTTTATGACAAACCTCAAATATATATTTTCTAAAGATACTGCTGTATCACATTTATTACAAACTCTGAAAGC
AATATTTCATCATAAAATGTTGATGAAAAAATAAATAAATGAAC

#### FASTA protein

**RAG1L-B**

>PflRAG1L-B_01_DNA
MEYHSSCLQSLCRVCSKHIGNNATQADVYAAELDKLYNIKVGDDDIQIHPTKLCKACTSRLQRCRRSAADFQPMQIEVATFVPHQDQGCVVCDQPAQLATGTAPKAKGRPPKRKCGGGQYGGPGRGHKKSSSDIPLSGSDIPSTSSSIPSTSRGIPSTSSSSELTVESVKRKLLTIESPEKEEAITEATEITSIPLDRFIEKDIAEHYVCSICQGVPTT
PCISPCSHIFCVGCIQQWLANSCACPSCREILECDDCQNLTGNHLNIYDSLRLRCTYSHLGCETMTTLPNYIDHELTCKYKAKGKRSTYGKTRVKQSLRTADRQYCKQKRLKACYDFLRDFCTANSESTEDVLFFLLRSYLYDSGDRERSNLVDDLWSKTQSKMSADECLALRIDNLQTKNRYKAQYDMFKSKSVSVLVAPNQLDMLERTYMPGTARYA
IVGEENFEHIYQTPVKLHRKDLSISTTADSVEPIELNSEYPNDNKEFPGPNLAGVRFRYTDAVAKTLEEIEPEISENLKSIGVSSTKTELVLRTFIKDGSDGMGDVEIHRRKGERTLPANAFRAAFAVVKCEVEVGNEIKTVWDQKNPNSVRCNRPLIEAIAEENNDSTVHYCLLTMEGERELMKNKVMKIDCGDYWRCHYLVFVTSMVDEKLDRSAGG
LQGAGSGYPCTLCDCTREEAISKLGSFSITRKRTEIIEKAEIRRVNPKILSQNELNLSCKGVKKHPLLLSEPVERGIDSTHANINLASFFKKVLVREVAEITQWEKTAELKSSLDMAEKRLDDHLKAEIGINPSLMMPGNYARVFFDEKNEQAILSLIPQAQRREDFAAVLAKFRFLKKVYCAKLPKVDYKDDIESVKTVGIEMGMLLIDKFGYARWPN
YLHKVIEHTQELIEKEDSPGTIGGISGEGNEAGNKLFRQFRKLHSRKGSVMGGLRDTIWLHWLYSSPKLCRHAEVAHRKNRCSACGCLGHNRLTCSNES

>PflRAG1L-B_RNA_GDGM01438088.1
MEYHSSCLQSLCRVCSKHIGNNATQADVYAAELDKLYNIKVGDDDIQIHPTKLCKACTSRLQRCRRSAADFQPMQIEVATFVPHQDQGCVVCDQPAQLATGTAPKAKGRPPKRKCGGGQYGGPGRGHKKSSSDIPLSGSDIPSTSSSIPSTSRGIPSTSSSSELTVESVKRKLLTIESPEKEEAITEATEITSIPLDRFIEKDIAEHYVCSICQGVPTT
PCISPCSHIFCVGCIQQWLANSCACPSCREILECDDCQNLTGNHLNIYDSLRLRCTYSHLGCETMTTLPNYIDHELTCKYKAKGKRSTYGKTRVKQSLRTADRQYCKQKRLKACYDFLRDFCTANSESTEDVLFFLLRSYLYDSGDRERSNLVDDLWSKTQSKMSADECLALRIDNLQTKNRYKAQYDMFKSKSVSVLVAPNQLDMLERTYMPGTARYA
IVGEENFEHIYQTPVKLHRKDLSISTTADSVEPIELNSEYPNDNKEFPGPNLAGVRFRYTDAVAKTLEEIEPEISENLKSIGVSSTKTELVLRTFIKDGSDGMGDVEIHRRKGERTLPANAFRAAFAVVKCEVEVGNEIKTVWDQKNPNSVRCNRPLIEAIAEENNDSTVHYCLLTMEGERELMKNKVMKIDCGDYWRCHYLVFVTSMVDEKLDRSAGG
LQGAGSGYPCTLCDCTREEAISKLGSFSISRKRTEIIEKAEIRRVNPKILSQNELNLSCKGVKKHPLLLSEPVERGIDSTHANINLASFFKKVLVREVAEITQWEKTAELKSSLDMAEKRLDDHLKAEIGINPSLMMPGNYARVFFDEKNEQAILSLIPQAQRREDFAAVLAKFRFLKKVYCAKLPKVDYKDDIESVKTVGIEMGMLLIDKFGYARWPN
YLHKVIEHTQELIEKEDSPGTIGGISGEGNEAGNKLFRQFRKLHSRKGSVMGGLRDTIWLHWLYSSPKLCRHAEVAHRKNRCSACGCLGHNRLTCSNES

>PflRAG1L-B_RNA_GDGM01014902.1
MLHQADVYAAELDKLYNIKVGDDDIQIHPTKLCKACTSRLQRCRRSAADFQPMQIEVATFVPHQDQGCVVCDQPAQLATGTAPKAKGRPPKRKCGGGQYGGPGRGHKKSSSDIPLSGSDIPSTSSSIPSTSRGIPSTSSSSELTVESVKRKLLTIESPEKEEAITEATEITSIPLDRFIEKDIAEHYVCSICQGVPTTPCISPCSHIFCVGCIQQWLAN
SCACPSCREILECDDCQNLTGNHLNIYDSLRLRCTYSHLGCETMTTLPNYIDHELTCKYKAKGKRSTYGKTRVKQSLRTADRQYCKQKRLKACYDFLRDFCTANSESTEDVLFFLLRSYLYDSGDRERSNLVDDLWSKTQSKMSADECLALRIDNLQTKNRYKAQYDMFKSKSVSVLVAPNQLDMLERTYMPGTARYAIVGEENFEHIYQTPVKLHRKD
LSISTTADSVEPIELNSEYPNDNKEFPGPNLAGVRFRYTDAVAKTLEEIEPEISENLKSIGVSSTKTELVLRTFIKDGSDGMGDVEIHRRKGERTLPANAFRAAFAVVKCEVEVGNEIKTVWDQKNPNSVRCNRPLIEAIAEENNDSTVHYCLLTMEGERELMKNKVMKIDCGDYWRCHYLVFVTSMVDEKLDRSAGGLQGAGSGYPCTLCDCTREEAI
SKLGSFSISRKRTEIIEKAEIRRVNPKILSQNELNLSCKGVKKHPLLLSEPVERGIDSTHANINLASFFKKVLVREVAEITQWEKTAELKSSLDMAEKRLDDHLKAEIGINPSLMMPGNYARVFFDEKNEQAILSLIPQAQRREDFAAVLAKFRFLKKVYCAKLPKVDYKDDIESVKTVGIEMGMLLIDKFGYARWPNYLHKVIEHTQELIEKEDSPGT
IGGISGEGNEAGNKLFRQFRKLHSRKGSVMGGLRDTIWLHWLYSSPKLCRHAEVAHRKNRCSACGCLGHNRLTCSNES

>PflRAG1L-B_RNA_GDGM01481817.1
MEGQYGGPGRGHKKSSSDIPLSSSDKPSTSSGIPSTSNSSELTVESVKRKLLTIEISEKEEAITEATEITSIPLDRFIEKDIAEHYVCSICQGVPTTPCISPCSHIFCVGCIQQWLANSCACPSCREILECDDCQNLTGNHLNIYDSLRLRCTYSHLGCETMTTLPNYIDHELTCKYKAKGKRSTYGKTRVKQSLRTADRQYCKQKRLKACYDFLRDFC
TANSESTEDVLFFLLRSYLYDSGDRERSNLVDDLWSKTQSKMSADECLALRIDNLQTKNRYKAQYDMFKSKSVSVLVAPNQLDMLERTYMPGTARYAIVGEENFEHIYQTPVKLHRKDLSISTTADSVEPIELNSEYPNDNKEFPGPNLAGVRFRYTDAVAKTLEEIEPEISENLKSIGVSSTKTELVLRTFIKDGSDGMGDVEIHRRKGERTLPANAF
RAAFAVVKCEVEVGNEIKTVWDQKNPNSVRCNRPLIEAIAEENNDSTVHYCLLTMEGERELMKNKVMKIDCGDYWRCHYLVFVTSMVDEKLDRSAGGLQGAGSGYPCTLCDCTREEAISKLGSFSISRKRTEIIEKAEIRRVNPKILSQNELNLSCKGVKKHPLLLSEPVERGIDSTHANINLASFFKKVLVREVAEITQWEKTAELKSSLDMAEKRLD
DHLKAEIGINPSLMMPGNYARVFFDEKNEQAILSLIPQAQRREDFAAVLAKFRFLKKVYCAKLPKVDYKDDIESVKTVGIEMGMLLIDKFGYARWPNYLHKVIEHTQELIEKEDSPGTIGGISGEGNEAGNKLFRQFRKLHSRKGSVMGGLRDTIWLHWLYSSPKLCRHAEVAHRKNRCSACGCLGHNRLTCSNES

**RAG2L-B**

>PflRAG2L-B_01_DNA
MAVAPELALDVFRFIPLTEGSSDKRKMTRKRFFELGEYFPPEGHMNVSINNGDGVVTVYTLGGGRWKEESTWSLSNELYSLSFTLDDTDVDESVQKFTTRGAMLSPLHAASMLNISTPDKVKLLVWGGYHLGSLFCTNEAVTMEIQRKTATCVIYKDPNDMSFHLPDEKHQSGDIPSARCGHTLTPIPGQHAAVLFGGAEMPNRFRRVPSFEQDTKDG
HFYLLNTDSLSWKKLNVPQLEPRAFHTATYLSSSSTICYVGGVTYRDQKPYKRHQINEVTLLSISATNEYAVKSVFLSESLPYHVSMHGALQFNDQVIVYGGYVTPCALYQSNARPAKPSSSMFLLNTTSEVLTRLEAPDSFASAGLSMVSLDNTAIMGLGGTHKNIFVFTSKAMCPAPCDLEDECTIKDTPEISPIAWIQCEGKCKRWLHQFCIKLEV
IPKGKYVCNSCKSTGGRKRSRKN

### Ptychodera flava (Taiwan population)

#### RAGL-A

##### Pfl$^{T}$RAGL-A.54

| Genomic loci | RAG1L status | RAG2L status |
|---|---|---|
| Pfl$^{T}$RAGL-A_54_v1 | *pseudogenic (frameshifts)* | *pseudogenic (frameshifts)* |
| Pfl$^{T}$RAGL-A_54_v2 | potentially functional (1 complete CDS) | potentially functional (2 complete CDSs) |

```
          3401                                                                                                                                                                                                    3600
PflT_54_v1  GGCCGCTATCCGTGACTATGATGTATGACAGATGTCACATGAATGTATCACAGTCAAATTCATGTTGCTGATAAATCACGGCGATCGGCAGATTTCAGATTATAATAACGGTCAGAGCAGGAAGTCTGTCCGGGTTCTCGGAATTCAGTCTGCTTTACTCAGAAAAAGACTAGTCTCCATAGTTACAAAGCAAATTTCGG
PflT_54_v2  GGCCGCTATCCGTGACTGTGATGTTTGACAGATGTCACATGAATGTATCACAGTCAAATTCATGATGCTGACAAATCGCTGCGATCGGCAGATTTTAGATTATAATAACGGTCAGAGCAGGAAGTCTGTCCGGGTTCTCGGAATTCAGTCTGCTTTACTCAGAAAAAGACTAGTCTCCATAGTTACAATGCAAATTTCGG

          3601                                                                                                                                                                                                    3800
PflT_54_v1  CCTTGACGTGGTGAGTATTCCGCTGTCTTCGAGAGGCAGAGACTATAAATTTAATTTTCTCTCGCTATCCCGGGCAATCACAAATGTACCGTGTGTGTTGATTTTCCGCATTTGACACCATCGCTCGATCGTGTTTTTACTGTCGTGTACACAGCTATGAAAGGAATTTGCTTTGGCGTACGCGGTTGTTGCTACATGTA
PflT_54_v2  CCTTGACGTGGTGAGTATTCCGCTGTCTTCGAGAGGCAGAGACTATAAATTTAATTTTCTCTCGCTATCCCGGGCAATCACAAATGTACCGTGTGTGTTGATTTTCCGCATTTGACACCATCGCTCGATCGTGTTTTTACTGTCGTGTACACAGCTATGAAAGGAATTTGTTTTGGCGTACGCGGTTGTTGCTACATGTA

          3801                                                                                                                                                                                                    4000
PflT_54_v1  GTTGTTTTGCCAATGTGGGTTTCACGTTAATTACACGCCCTGTACAGGACAAACTGCCAAAGTTCACGATCACTTCACCAATGCGACCATCTAACAATTTATTCATTCGAATCACTGTTTTGTCAGTCCGCTGACATTTAGCGACTCAAATTCTGCGGCTCATATAAAACATGGCAGGAACAAAAACATCGAATTACAAA
PflT_54_v2  GCTGGTTTGCCAATGTGGGATTCACGTTAATTACACGCCCTGTACAGGACAAACTGCCAAAGTTCACGATCACTTCACCAATGCGACCATCTAACAATTTATTCATTCGAATCACTGTTTTGTCAGTCCGCTGACATTTAGCGACTCAAATTCTGCGGGTCATATAAAACATGGCAGGAACAAAAACATCGAATTACAAA

          4001                                                                                                                                                                                                    4200
PflT_54_v1  CACGATAACAGTCGAACATCGAGCGCGAGCTGGTGAACGCGAAGCGAGCACACAGCTCCTTGTAAAATAATATAAAATGATATCATTTAAAGTGGTAAGCTGGACACGAGAGAGAGAGAGAGAGAGAGAGAGAGAGAGAGAGAGAGAGAGAGAGAGAGAGAGAGAGAGTGTGGATGAACTGGTAACTCTA
PflT_54_v2  CACGATAACAGTCGAACATCGAGCGCGAGCTGGTGAACGCGAAGCGAGCACACAGCTCCTTGTAAAATAATATAAAATGATATCATTTAAAGTGGTAAGCTGGACACGAGAGAGAGAGAGAGAGAGAGAGAGAGAGAGAGAGAGAGAGAGAGAGAGAGAGAGAGAGA-TGTGGATGAACTGGTAACTCTA

          4201                                                                                                                                                                                    TSD         5'-TIR5
PflT_54_v1  TCGTATCTTTCTCACGTTGATTTACATACCACAAAGATGCCACAAAAAGCCGATCGACTTTTTCCACTGAACACTGACACGTTAACATTCAAACGAAATAACACAAACTCACTTTTATTTTTTTTTTGGAGCCTGGGGAAATCTATGAAGTGGCATAAAGCAAAATCTGTATTGAGAAAAGCGGTA------CACAGCGT
PflT_54_v2  TCGTATCTTTCTCACGTTGATTTACATACCACAAAGATGCCACAAAAAGCCGATCGACTTTTTCCACTGAACACTGACACGTTAACATTCAAACGAAATAACACAAACTCACTTTTATTTTTTTTTTGGAGCCTGGGGAAATCTATGAAGTGGCATAAAGCAAAATCTGTATTGAGAAAAGCGGTACCGACTCACAGCGT

          4401    5'-TIR5                                                                                                                                                                                 4600
PflT_54_v1  AACATGTCAATATAGAAATCGATTAACACTACGATGCGTCCCAGAGAGTATCGACCAATCAGAGAGCAGCATTCATCAAGCCACGACCTTTCACGTATGGCTTCGAGATTTCGCGTGATCTAGGGTGTTTACTTCAGAAACTCTACAATCTACTACATACTAAGGGGTCTCAACTGTTTTACACATGCGCAAACGCCATC
PflT_54_v2  AACATGTCAATATAGAAATCGATTAACACTACGATGCGTCCCAGAGAGTATCGACCAATCAGAGAGCAGCATTCATCAAGCCACGACCTTTCACGTATGGCTTCGAGATTTCGCGTGATCTAGGGTGTTTACTTCAGAAACTCTACAATCTACTACATACTAAGGGGTCTCAACTGTTTTACACATGCGCAAACGCCATC

          4601                                                                                                                                                                                                    4800
PflT_54_v1  TTGAAGAGTGGCTGATGATCATGTCGAAAACCGGAGGTCGAAAGACGTAAGTATCGCCGAGATATCGTAATCTGAATCTCGCGTTGCAATTTTTCACTCGATTATGTGTATCAAAATTATGCGGATACAATTTTGGGATTTTCTGTCAAACTTGTTTTTCCTG--TTAAAGTTTGGGTTTTTCTCAGTCCGTAACGACAT
PflT_54_v2  TTGAAGAGTGGCTGATGATCATGTCGAAAACCGGAGGTCGAAAGACGTAAGTATCGCCGAGATATCGTAATCTGAATCTCGCGTTGCAATTTTTCACTCGATTATGTGTATCAAAATTATGCGGATACAATTTTGGGATTTTCTGTCAAACTTGGTTTTTCCTGTTTAAAGTTTGGGTTTTTCTCAGTCCGTAACGACAT

          4801                                                                                                                                                                                                    5000
PflT_54_v1  TGTTTACCGAAGGAACATGTGCTGTACGCTTGGACTTTCTTCAAATGGCTTGGACTTTTAGATCTATACCCATCCAACATGTGTGATTGAGTGTAATGGCCGTTACATTTAAAAATGTTC-CTGTGATGGTAGCTAGGTCAGGGAAATGATTGTTTTGTGTTTAATATATTTTTAAGGTTTGAATGAGCGTGTTACGATA
PflT_54_v2  TGTTTACCGAAGGAACATGTGCTGTACGCTTGGACTTTCTTCAAATGGCTTGGACTTTTAGATCTATACCCATCCAACATGTGTGATTGAGTGTAATGGCCGTTACATTTAAAAATGTTC□CTGTGATGGTAGCTAGGTCAGGGAAATGATTGTTTTGTGTTTAATATATTTTTAAGGTTTGAATGAGCGTGTTACGATA

          5001                                                                                                                                                                                                    5200
PflT_54_v1  TTTAGAATTCTGAGCAGCATCCTAGAGGGTCAATTTGGGTGTACAACGCATGGTTCATGCGGTAGCTAGCTAACCCAGGCCATATGGCACCAGGTACAGGTTGACTAACGAAATGTTTTGAGGTTAGCATAGTATGAGTACGTCAGATTCATATGTCATCGGCATAGAAGGTCGCTTGTGTAAGATAAGACCATATTTTT
PflT_54_v2  TTTAGAATTCTGAGCAGCATCCTAGAGGGTCAATTTGGGTGTACAACGCATGGTTCATGCGGTAGCTAGCTAACCCAGGCCATATGGCACCAGGTACAGGTTGACTAACGAAATGTTTTGAGGTTAGCATAGTATGAGTACGTCAGATTCATATGTCATCGGCATAGAAGGTCGCTTGTGTAAGATAAGACCATATTTTT

          5201                                                                                                                                                                                                    5400
PflT_54_v1  AGGCTGTCATTCATTTCATTTATTTATTTTGCAGTGGGTTGCTGTCAGCAATGAATCATTTACAAGTTTGTTGCACTGTTATGCAATGCAATGCCTTGACCTGAATCAGTATTTCTTGCTTGGGTTCCTGCGGTATGAAGTAGAAA-TTAGGCTGGCCACGGTGTCCTGGGAATTAATGTTTTACCACTAAGGTAAGAAT
PflT_54_v2  AGGCTGTCATTCATTTCATTTATTTATTTTGCAGTGGGTTGCAGTCAGCAATGAATCATTTACAAGTTTGTTGCACTGTTATGCAATGCAATGCCTTGACCTGAATCAGTATTTCTTGCTTGGGTTCCTGCGGTATGAAGTAGAAAATTAGGCTGGCCACGGTGTCCTGGGAATTAATGTTTTACCACTAAGGTAAGAAT

          5401                                                                                                                                                   RAG1L-A                                          5600
PflT_54_v1  AAAATTCTATTGGATTTATTGTTTACACATATATAAGGATGAAAGGTACAGACCATTGTACTGATTGGTACTGTACTTGCCAAATTGATATGCTTAAATAATATTTTAAATATGTTAATTGTTTTCAGAGCGGAATGACATCAGCTTTTCAACACAGAGAGTACCTTAAGAGGGTGTGTCGTGTATGTGGTAGAACCCTG
PflT_54_v2  AAAATTCTATTGGATTTATTGTTTACACATATATAAGGATGAAAGGTACAGACCATTGTACTGATTGGTACTGTACTTGCCAAATTGATATGCTTAAATAATATTTTAAATATGTTAATTGTTTTCAGAGCGGAATGACATCAGCTTTTCAACACAGAGAGTACCTTAAGAGGGTGTGTCGTGTATGTGGTAGAACCCTG

          5601                                                                                                                                                                                                    5800
PflT_54_v1  AAGACCACATCTTACAGCAGCTCAGGACATTACAGACAGGGAGAAGTGTGCATCCCATACAGTGAACTTTTAGAAGAATGCTTTAAAGTAAATGTTCGAAATGATGACCCTTTTGTACATCCGCCTAGATTTGGTTCATGTTGCCGTAGAGTTCTTTTAAAGTATTCTGTTGCACAGAACCTGGGTAAACATTACTATAC
PflT_54_v2  AAGACCACATCTTACAGCAGCTCAGGACATTACAGACAGGGAGAAGTGTGCATCCCATACAGTGAACTTTTAGAAGAATGCTTTAAAGTAAATGTTCGAAATGATGACCCTTTTGTACATCCGCCTAGATTTGGTTCATGTTGCCGTAGAGTTCTTTTAAAGTATTCTGTTGCACAGAACCTGGGTAAACATTACTATAC

          5801                                                                                                                                                                                    F       6000
PflT_54_v1  CCATATCAGTGTTTGTAACTGGAAACATCATTGTAAGTATGAAAACCAAAGTTGTCCTCTTTGCAAACGTTTTTCTCAAAATGGTTCTGGTAGACCTAAAAAACTTAAAAGGGGTCGCTTTAAATTAAATACAAATGAAAAAAATGAAACTGACAAGAGTCCTGTTGCAGACAATTT-GCTCATATACCAAATGACCACT
PflT_54_v2  CCATATCAGTGTTTGTAACTGGAAACATCATTGTAAGTATGAAAACCAAAGTTGTCCTCTTTGCAAACGTTTTTCTCAAAATGGTTCTGGTAGACCTAAAAAACTTAAAAGGGGTCGCTTTAAATTAAATACAAATGAAAAAAATGAAACTGACAAGAGTCCTGTTGCAGACAATTTTGCTCATATACCAAATGACCACT

          6001                                                                                                                                                                                                    6200
PflT_54_v1  GTTATACCACCCCAGCTTCCTGTAAGTTTGCCCAGTCTGTGCAATCAGACCAAAAGACAGAGACATATGAATTCTGTACAGAGGAAAATATTCAGACTGTACCCATGACAAGTCGATTTCCAACAGATTTATGTGTAACTGAACCACATCCAGAATTCATTTGTAGAATGTGTTTAGGAGTCCTAGACACACCACTTCTA
```

```
PflT_54_v2  GTTATACCACCCCAGCTTCCTGTAAGTTTGCCCAGTCTGTGCAATCAGACCAAAAGACAGAGACATATGAATTCTGTACAGAGGAAAATATTCAGACTGTACCCATGACAAGTCGATTTCCAACAGATTTATGTGTAACTGAACCACATCCAGAATTCATTTGTAGAATGTGTTTAGGAGTCCTAGACACACCACTTCTA

            6201                                                                                                                                                                                                   6400
PflT_54_v1  ACACCGTGTACTCACACTTTCTGTGCAGGCTGCATTAAGAACTGGTTAAGTCTGTGTCATTTTTGTCCTCTCTGTAAACAACCTATGGCGTCATCAGATTTAAAAGAACCATATAGAGGATTTGTAGAAATTCTTAATGATACTAGAATGAAATGTAATTGTCCAGAAGCAGTTTCAAGCAAAAAAGGTGTCAAGCTTAG
PflT_54_v2  ACACCGTGTACTCACACTTTCTGTGCAGGCTGCATTAAGAACTGGTTAAGTCTGTGTCATTTTTGTCCTCTCTGTAAACAACCTATGGCGTCATCAGATTTAAAAGAACCATATAGAGGATTTGTAGAAATTCTTAATGATACTAGAATGAAATGTAATTGTCCAGAAGCAGTTTCAAGCAAAAAAGGTGTCAAGCTTAG

            6401                                                                                                                                                                                                   6600
PflT_54_v1  TAATTGGCTTCAGCACACTTTAAATTGTGGAAAGGATTTTGAAATTGATGAATGGTTAGAAAAAGTTTTGTCATTGTGGAACGAAATTTCTCTCACTCAAGTTGATTCACATTTAAATAGTACTGAACATTTGCCTGAGACAGATAAGTCAAATCTGACAACAAGCCATGGCCATCCAGACATTGCTGCAGGTAGCACTG
PflT_54_v2  TAATTGGCTTCAGCACACTTTAAATTGTGGAAAGGATTTTGAAATTGATGAATGGTTAGAAAAAGTTTTGTCATTGTGGAACGAAATTTCTCTCACTCAAGTTGATTCACATTTAAATAGTACTGAACATTTGCCTGAGACAGATAAGTCAAATCTGACAACAAGCCATGGCCATCCAGACATTGCTGCAGGTAGCACTG

            6601                                                                                                                                                                                                   6800
PflT_54_v1  TTGGAAGACCTAAAACTTCACTGTTGTGTCTCCAACAACAAGGACAAAATTTAAGATTGAAAAATATCAAAAAACAAATAGACGTGTTTGCCAGAGAGAAAGGGGAAGATCTCAGGGTAGTATATTTTGGCCTTTTGCAAAACTATCTGAGAGCCAACAAGCAAGGTAGTTTAGCTGAACAAGTGAAGTTGCTATATAAT
PflT_54_v2  TTGGAAGACCTAAAACTTCACTGTTGTGTCTCCAACAACAAGGACAAAATTTAAGATTGAAAAATATCAAAAAACAAATAGACGTGTTTGCCAGAGAGAAAGGGGAAGATCTCAGGGTAGTATATTTTGGCCTTTTGCAAAACTATCTGAGAGCCAACAAGCAAGGTAGTTTAGCTGAACAAGTGAAGTTGCTATATAAT

            6801                                                                                                                                                                                                   7000
PflT_54_v1  GGGGAATCTTGTCAAATGTCACCTCAGGAATGTCTAGCCTTGCGTGTAAATACTTTTACCAGTTGTAATCAGTACAAGAAAATTTACTCAGCTACAAATGAGCTTAGTGGAGTTAAAATCTTCCAACCTCTCATCAATGTTGTGAAAGCCGAAGAAAGTTTTCTACCAGGGTCTGTAAAGTTCTACTTTAACCCATCTTT
PflT_54_v2  GGGGAATCTTGTCAAATGTCACCTCAGGAATGTCTAGCCTTGCGTGTAAATACTTTTACCAGTTGTAATCAGTACAAGAAAATTTACTCAGCTACAAATGAGCTTAGTGGAGTTAAAATCTTCCAACCTCTCATCAATGTTGTGAAAGCCGAAGAAAGTTTTCTACCAGGGTCTGTAAAGTTCTACTTTAACCCATCTTT

            7001                                                                                                                                                                                                   7200
PflT_54_v1  GAAAACTGTTTCTCCCATTTGGCAAGGTACAGAGCATGTTGATGCTCTGTCAGGACTCAGTCGCGATATACCACTTCTAGAATCACCTGATGTTTATGGGGCAAGGTATAGATATGACAAAGCAGTTGCTATGGCTTTGAAAGATATTGATAATGAAATCTCAGAAGGCCTCAGTAAATTAGAAGGGGTGGATACAAATC
PflT_54_v2  GAAAACTGTTTCTCCCATTTGGCAAGGTACAGAGCATGTTGATGCTCTGTCAGGACTCAGTCGCGATATACCACTTCTAGAATCACCTGATGTTTATGGGGCAAGGTATAGATATGACAAAGCAGTTGCTATGGCTTTGAAAGATATTGATAATGAAATCTCAGAAGGCCTCAGTAAATTAGAAGGGGTGGATACAAATC

            7201                                                                                                                                                                                                   7400
PflT_54_v1  AACTTGTTCTAACAGCCACTGTCAAAGATGGCAGTGATGGTCTTGGAGATGTGAAAGAAGTCAGATCTCAGGGGCTTGTCGTTCCAAACAAGGCCTTGCGGTTTTCATTTGTAATTTTAGAAGTCACTGTCAATGTAGATGGCAAGATTATTACTGTATTTGAAGAAGGTTCTCCCAACTCAGAATTATGCACACGCCCT
PflT_54_v2  AACTTGTTCTAACAGCCACTGTCAAAGATGGCAGTGATGGTCTTGGAGATGTGAAAGAAGTCAGATCTCAGGGGCTTGTCGTTCCAAACAAGGCCTTGCGGTTTTCATTTGTAATTTTAGAAGTCACTGTCAATGTAGATGGCAAGATTATTACTGTATTTGAAGAAGGTTCTCCCAACTCAGAATTATGCACACGCCCT

            7401                                                                                                                    F                                                                              7600
PflT_54_v1  CTCTTGGTAGCTCTGGCAGATGAAAATGACAGAAAAGGGTTATGCATGACACATATACCAATAATGATAGAGAGAGAGGAGATGGAGAAAAGTACCTTGTTTGTAGAAATTGAAGAGGGAAA-CAAAGAATCTACAAAGTTGTATTTAAAGGGACTATGTATGATGAAAAATTGCAGAGAAGTGGTGAGGGAATGTTAGC
PflT_54_v2  CTCTTGGTAGCTCTGGCAGATGAAAATGACAGAAAAGGGTTATGCATGACACATATACCAATAATGATAGAGAGAGAGGAGATGGAGAAAAGTACCTTGTTTGTAGAAATTGAAGAGGGAAAACAAAGAATCTACAAAGTTGTATTTAAAGGGACTATGTATGATGAAAAATTGCAGAGAAGTGGTGAGGGAATGTTAGC

            7601                                                                                                                                                                                                   7800
PflT_54_v1  GGCAGGGTCATCTTGGTTTTGTACACTTTGTGAGAAAGGAAGACTCGATCCCATATCATCAGTCTTGCCAATAACAAGAACACATGAAAACAATATGGCTTTGTATTCAAAGTATGTTGAAAATGCAAATCATATGAAGCAGGCAGATCTAGTGAAAGAAGTCAAAGGCATTGTTAGAAAACCATTTATAATGTCAGAGC
PflT_54_v2  GGCAGGGTCATCTTGGTTTTGTACACTTTGTGAGAAAGGAAGACTCGATCCCATATCATCAGTCTTGCCAATAACAAGAACACATGAAAACAATATGGCTTTGTATTCAAAGTATGTTGAAAATGCAAATCATATGAAGCAGGCAGATCTAGTGAAAGAAGTCAAAGGCATTGTTAGAAAACCATTTATAATGTCAGAGC

            7801                                                                                                                                                                                                   8000
PflT_54_v1  CTTGTGTTGATGCCACGCATGCTGAAATTCACTGGGGAGAGAAAAACTATGATATGTATGTACGTGAAATATCAGGGATAATGTGTTGGGGGAAGCCATCCAATGATGATGATGCAAAAAAAGTGAAATCAACAAAAAGAATGTTGGATACCGAATTACAGCTGCAATGTGGCTTAAAGAGAGAGCATGGCTTTATGGTT
PflT_54_v2  CTTGTGTTGATGCCACGCATGCTGAAATTCACTGGGGAGAGAAAAACTATGATATGTATGTACGTGAAATATCAGGGATAATGTGTTGGGGGAAGCCATCCAATGATGATGATGCAAAAAAAGTGAAATCAACAAAAAGAATGTTGGATACCGAATTACAGCTGCAATGTGGCTTAAAGAGAGAGCATGGCTTTATGGTT

            8001                                                                                                                                                                                                   8200
PflT_54_v1  GATGGAAATTATGCACGTGATTTGGTCAAATCAGAAACAGTTGATGTGGTTTGCTCTTTAATTCAGAGTGAAGCAAGAAAATCTGTTGTTCGTGAATATGTTTCCTACTATCGAGAATTGCGATCAATATACAGGAAACACAATGCTGACTTAGATGATGCCGTGAAATTTAAAACCATAGCTGCAATTTTTTACAAATT
PflT_54_v2  GATGGAAATTATGCACGTGATTTGGTCAAATCAGAAACAGTTGATGTGGTTTGCTCTTTAATTCAGAGTGAAGCAAGAAAATCTGTTGTTCGTGAATATGTTTCCTACTATCGAGAATTGCGATCAATATACAGGAAACACAATGCTGACTTAGATGATGCCGTGAAATTTAAAACCATAGCTGCAAATTTTTACAAATT

            8201                                                                                                                                                                                                   8400
PflT_54_v1  TCTACATGAGAAATTTTCTTATTTGCATCCACTCTCTAACTATCAACACAAGGTATTAGATCACATTCCACAGTTGATCGAGCAATATGGCTCTGTTGGTAAATTTGCCTCTGAAGGCAATGAAAAAGGCAATAAACTTTTCAGAAGATTTCGTAAATTTAATGCACGTAAGTCTCAAATTCATGAAATGCCAGACATTC
PflT_54_v2  GCTACATGAGAAATTTCCTTATTTGCATCCACTCTCTAACTATCAACACAAGGTATTAGATCACATTCCACAGTTGATCGAGCAATATGGCTCTGTTGGTAAATTTGCCTCTGAAGGCAATGAAAAAGGCAATAAACTTTTCAGAAGATTTCGTAAATTTAATGCACGTAAGTCTCAAATTCATGAAATGCCAGACATTC

            8401                                                                                                                                                                                                   8600
PflT_54_v1  TGAAATTTCATTGGCTATATACAATGAAAAGTCTTCAGACTATGGTAAACTGGGAAAAAACTAGATCACTAGTCTGTAGCAAATGTGGTTGTGCTGGTCACAATCAAAGGACATGATATGAAATCAAAGATGAGACAATGACCATGCATGTTTTATGTATCTGAATCTGTAAGTTTGAAATTAAGTAGCATCTGTATGTA
PflT_54_v2  TGAAATTTCATTGGCTATATACAATGAAAAGTCTTCAGACTATGGTAAACTGGGAAAAAACTAGATCACTAGTCTGTAGCAAATGTGGTTGTGCTGGTCACAATCAAAGGACATGATATGAAATCAAAGATGAGACAATGACCATGCATGTTTTATGTATCTGAATCTGTAAGTTTGAAATTAAGTAGCATCTGTATGTA

            8601                                                                                                                                                                                                   8800
PflT_54_v1  TCAGACTACAGAGTCCAAACTTCAGACACAAAGAGTAGGCTGGCCATACACTCTTATTTACTTAAAAAGTGTTTCTGTTCCATATACTGACTGTACTTTCAAGAGTCAATTCTAGCTTCTATGCTAGTCTTAATATACCTAGATGGTTAAAGTGTATTATATATTTGATGTACATAGATGAATTCCACAAGAATCATCAA
PflT_54_v2  TCAGACTACAGAGTCCAAACTTCAGACACAAAGAGTAGGCTGGCCATACACTCTTATTTACTTAAAAAGTGTTTCTGTTCCATATACTGACTGTACTTTCAAGAGTCAATTCTAGCTTCTATGCTAGTCTTAATATACCTAGATGGTTAAAGTGTATTATATATTTGATGTACATAGATGAATTCCACAAGAATCATCAA

            8801                                                                                                                                                                                                   9000
PflT_54_v1  ATACAAGAAAGCTGAGTTTGTGTTAAATATTTGTATGTGTTTTAATCTTACCAGTAACCTGCCCACACTTTTGAAGGGACTCAAACAAAATCATTTGATTACTTTGGCAGAAAAGCCAAAAGTTTTAGTTATACTGGCCATAAAAAATGAGATGAGAAGCGTAAAGTTTGCAATAGAATTTTGTACATATATTCATGAAA
PflT_54_v2  ATACAAGAAAGCTGAGTTTGTGTTAAATATTTGTATGTGTTTTAATCTTACCAGTAACCTGCCCACACTTTTGAAGGGACTCAAACAAAATCATTTGATTACTTTGGCAGAAAAGCCAAAAGTTTTAGTTATACTGGCCATAAAAAATGAGATGAGAAGCGTAAAGTTTGCAATAGAATTTTGTACATATATTCATGAAA

                                                                                                                                          RAG2L-A
            9001                                                                                                                                                                                                   9200
PflT_54_v1  GATTGTAACAAGTTGTAAAATGAACTCTCAGTAAAGGTTTCATTCAATTCATGCACATGCTACATGTACATGCTATGATTTCTTGTGCAGTTGTCATGTTCTCAGAAATTACAATCAGAACATGTAAATTTGATGTTACTCTTATTTTCCAGTTGTCGTGAAGAAAATCGTTGCCAGAGCATTTAATGTGAAAAGCTCT
PflT_54_v2  GATTGTAACAAGTTGTAAAATGAACTCTCAGTAAAGGTTTCATTCAATTCATGCACATGCTACATGTACATGCTATGATTTCTTGTGCAGTTGTCATGTTCTCAGAAATTACAATCAGAACATGTAAATTTGATGTTACTCTTATTTTCCAGTTGTCGTGAAGAAAATCGTTGCCAGAGCATTTAATGTGAAAAGCTCT

            9201                                                                                                                                                                                                   9400
PflT_54_v1  AACACATTGTGTACATCGCACTGTTGAAAAGTAACTCTTATCAGTCACGCTACAATTAATGTTAGCACATGTTGTGAAACACGAAGGGCATTCAAAGTTTATACAGTCCAGCTTTGTTCTATTTAAGTGATCTAATTCAACACATTTGTCATGGTACCACCTTCCACAGAGTACAGTTTGATTGTTCATGATGTAAGTTC
PflT_54_v2  AACACATTGTGTACATCGCACTGTTGAAAAGTAACTCTTATCAGTCACGCTACAATTAATGTTAGCACATGTTGTGAAACACGAAGGGCATTCAAAGTTTATACAGTCCAGCTTTGTTCTATTTAAGTGATCTAATTCAACACATTTGTCATGGTACCACCTTCCACAGAGTACAGTTTGATTGTTCATGATGTAAGTTC

            9401                                                      F                                                                                                                                            9600
PflT_54_v1  GATCACATTTCAGCCAGTTGAACTTGGCTTGTTCAGTAGCTGCTAAATTGGCAA-TTTGCAGCCATCCAACATACATTCCCATGTTTCCTCCTCACTATCATCTGATGTATTGTCTATCTTTCCTGCTCTGCTGCCCTCTTCTGACTCATTCCTGTCTGTGCTAACTTTGACCTCAGCTGCCTCACTTATTCCGTTCTGA
PflT_54_v2  GATCACATTTCAGCCAGTTGAACTTGGCTTGTTCAGTAGCTGCTAAATTGGCAAATTTGCAGCCATCCAACATACATTCCCATGTTTCCTCCTCACTATCATCTGATGTATTGTCTATCTTTCCTGCTCTGCTGCCCTCTTCTGACTCATTCCTGTCTGTGCTAACTTTGACCTCAGCTGCCTCACTTATTCCGTTCTGA

            9601                                                                                                                                                                                                   9800
PflT_54_v1  TCCTCAGTCTCTGAGCCTATACTTTCCTGATCTGTCATGTCTATGCTTATCCTTTCCTGCTCTTCCTCCGCTATGCTTACTCTGTCCTCATCTGTTGACCTTTCCTCCTCATCCCCCAACTGACTGTTATTACTCTGTGTCCTGGTTTCAGAGCTTGTCCCTTCATTTGAAAACTGAATTTATCATTTGCTTTTTTAC
PflT_54_v2  TCCTCAGTCTCTGAGCCTATACTTTCCTGATCTGTCATGTCTATGCTTATCCTTTCCTGCTCTTCCTCCGCTATGCTTACTCTGTCCTCATCTGTTGACCTTTCCTCCTCATCCCCCAACTGACTGTTATTACTCTGTGTCCTGGTTTCAGAGCTTGTCCCTTCATTTGAAAACTGAATTTATCATTTGCTTTTTTAC

            9801                                                                                                                                                                                                   10000
PflT_54_v1  TGTATCACAAACAGCAATTGAGCTGAAGAATGGAAGACTGTAGTATTTACTACTTAAGGTATCATAGTGTGCTGTTTCTATGAATCTATGGCATCACAAGGTGACTTGTAAACTGTGTTGTTACTCAATATGTAAACAGTGTCATTACATTTGAATGAGACTGGGTTAGGAATACCAATAAATGGTACAGATGTACAGA
PflT_54_v2  TGTATCACAAACAGCAATTGAGCTGAAGAATGGAAGACTGTAGTATTTACTACTTAAGGTATCATAGTGTGCTGTTTCTATGAATCTATGGCATCACAAGGTGACTTGTAAACTGTGTTGTTACTCAATATGTAAACAGTGTCATTACATTTGAATGAGACTGGGTTAGGAATACCAATAAATGGTACAGATGTACAGA
```

```
10001                                                                                                                                                                                                   10200
PflT_54_v1  CCATTGAATTCAGGTCAATTACGTCAACATCGGGTGGATGTCTTACATTGTTACCTATCACATAAATTCCACCAACAATATACATTTTACTGCTATTTAGAAAAACTGAATGGTAGGCTCTTGGAGTGATAGCTTGTAAACTAATACTTGATAATTTCATAGATTGTAAATTAAGAACAGAAAATATGTTTGAAGCCATG
PflT_54_v2  CCATTGAATTCAGGTCAATTACGTCAACATCGGGTGGATGTCTTACATTGTTACCTATCACATAAATTCCACCAACAATATACATTTTACTGCTATTTAGAAAAACTGAATGGTAGGCTCTTGGAGTGATAGCTTGTAAACTAATACTTGATAATTTCATAGATTGTAAATTAAGAACAGAAAATATGTTTGAAGCCATG

10201                                                                                                                                                                                                   10400
PflT_54_v1  GACACATTGTAAAAAGCATGACTGAGTTTTTGTGTATATTGTAAGCCTCCATAAAGATAAAGAGAATCTCCAAACTTAATTACACTGTGACCAAAAGCATGAATTTCCTCACCACTATATTGTCCTTGAGACCATGTTAATGAAATTTTCCCCCTTGGATCAACAAAGACAGAAAGCTGCCACAATTTTGTATTGTGCAT
PflT_54_v2  GACACATTGTAAAAAGCATGACTGAGTTTTTGTGTATATTGTAAGCCTCCATAAAGATAAAGAGAATCTCCAAACTTAATTACACTGTGACCAAAAGCATGAATTTCCTCACCACTATATTGTCCTTGAGACCATGTTAATGAAATTTTCCCCCTTGGATCAACAAAGACAGAAAGCTGCCACAATTTTGTATTGTGCAT

10401                                                                                         F                                                                                                         10600
PflT_54_v1  TTCTGAGTCAGTCAGTCCACCAAGGAGAACATAATGGTTTGGTTCAACTAAACACAAATTTGAGCCATGTACATGTGGAGGGTAATTTGACTAAGATGAATAATTTTGTGATTTCTTGTATTGTATCATCTTTGTCTATTACAAGCCAGCCTGTTACTGTTGTCTTAGACTTAACCCAGACCAAAGCTCTGTCACCTAAA
PflT_54_v2  TTCTGAGTCAGTCAGTCCACCAAGGAGAACATAATGGTTTGGTTCAACTAAACACAAATTTGAGCCATGTACATGTGGAGGGTAATTTGACTGAGATGAATAATTTT-TGATTTCTTGTATTGTATCATCTTTGTCTATTACAAGCCAGCCTGTTACTGTTGTCTTAGACTTAACCCAGACCAAAGCTCTGTCACCTAAA

10601                                                                                                                                                                                                   10800
PflT_54_v1  GAAATAACTGGATTTGTACATCCAGCAGTTGCTAGGCAAGTTGCCAGGATGTTTTTGGTGATTTGTATTTCTGAAATAAAAAAGATGATGACAAAAGTTGTTGGATTAGAGAGTGAAGCTTATCTGCATGCATGTATGGTGAGCAGGTCATTTTATTTTGTCATTGTTGAAATATGGCTCTGAAATATAAATTTGTTAAT
PflT_54_v2  GAAATAACTGGATTTGTACATCCAGCAGTTGCTAGGCAAGTTGCCAGGATGTTTTTGGTGATTTGTATTTCTGAAATAAAAAAGATGATGACAAAAGTTGTTGTATTAGAGAGTGAAGCTTATCTGCATGCATGTATGGTGAGCAGGTCATTTTATTTTGTCAGTGTTGAAATATGGCACTGAAATATAAATTTGTTAAT

10801                                                                                                                                                                                                   11000
PflT_54_v1  TTCATATTTTAATATGGGTATCTTTGTAGTTTAAAAAAATGCTCAAGACAGAAAGTGAGATGTCTTTAGTCGTGACATTTGTTAATATTGTTCACTGTACATTTCTGTACCCTTTGCAACCATACAATTGTTGAACACCTGTCCTGTATATTCAAACAAACAACTTGAAGGCAAATCTGATATCTTTGGGGCAGCAAAAT
PflT_54_v2  TTCATATTTTAATATGGATATCTTTGTAGTTCAAAAAAATGCTCAAGACAGAAAGTGAGATGTCTTTAGTCGTGACATTTATTAATATTGTTCACTGTACATTTCTCTACCCTTTGCAACCATACAATTGTTGAACACCTGTCCTGTATATTCAAACAAACAACTTGAAGGCAAATCTGATATCTTTGGGGCAGCAAAAT

11001                                                                                                                                                                                                   11200
PflT_54_v1  ACATGTGCTCTTGTCTTTTGCATTTGAGTAACCTCCTTATATAAATATGATCGAAAGAGATTGTCAACATGGAGCAGCAAATCCGTTTCTTTAACCTAGACTGTAAATCAATACAAAATGGGTGCACATGTAGCTAGGGGAAAAGGTAATGCACAGATAGAGCCCCATTGATGACTGCTGGAAATGTCATTTGGATGTCT
PflT_54_v2  ACATGTGCTCTTGTCTTTTGCATTTGAGTAACCTCCTTATATAAATATGATCGAAAGAGATTGTCAACATGGAGCAGCAAATCCGTTTCTTTAACCTAGACTGTAAATCAATACAAAATGGGTGCACATGTAGCTAGGGGAAAAGGTAATGCACAGATAGAGCCCCATTGATGACTGCTGGAAATGTCATTTGGATGTCT

11201                                                                                                                                                                                                   11400
PflT_54_v1  TGCTGGTGTACATGCAACTGAAAAAATCATTTTAGCCTGTGGGCCGTACAATAGCTGCACTTCTACTTCGCTCACTCTAATTTGAATCTTCGTATATAGTGTACATGGGGATCGACTTTCATAGTATGCTTGTTGGATCGAGGGACAGCGATCGACCGTGGCTTGTGTTACAGTTGTGAACGATCACATCGAGAATTTTC
PflT_54_v2  TGCTGGTGTACATGCAACTGAAAAAATCATTTTAGCCTGTGGGCCGTACAATAGCTGCACTTCTACTTCGCTCACTCTAATTTGAATCTTCGTATATAGTGTACATGGGGATCGACTTTCATAGTATGCTTGTTGGATCGAGGGACAGCGATCGACCGTGGCTTGTGTTACAGTTGTGAACGATCACATCGAGAATTTTC

11401                                                                                                                                                           RAG2L-A                                 11600
PflT_54_v1  TTTTTCGAATGGGCGATTACAAGTGCAAATCTTTGGAAGTTAGCTACCATATAAGAACTTTGTTGTTTTAACTTCAAGAATAATCAATCTAAAACACGAACAGAATGTTCTGTGACTTTTGAAGCTACATTGAACTGACGACCTACCGGGACTATCGAGCATGGCGGCTGCCATCTTGGATATTGTTTACGGCTCACTGA
PflT_54_v2  TTTTTCGAATGGGCGATTACAAGTGCAAATCTTTGGAAGTTAGCTACCATATAAGAACTTTGTTGTTTTAACTTCAAGAATAATCAATCTAAAACACGAACAGAATGTTCTGTGACTTTTGAAGCTACATTGAACTGACGACCTACCGGGACTATCGAGCATGGCGGCTGCCATCTTGGATATTGTTTACGGCTCACTGA

11601                                                                   3'-TIR  TSD                                                                                                                     11800
PflT_54_v1  GCATGTTCAGATCAAAAATATTTCCCACA-CCCATTGCGGACTTTCTACATTGTAAACATAGGAGATGTTACGGTGTGCGGTATAGAGATATTTTCGTTCTACGGAAAACATCAACTTGGAAACGGCAACGGTATAAATATCGCATGTGTCAGAAGTTATCGGAGCTATCTATGCTTACGTTATTTGTAGTCCTGTGATC
PflT_54_v2  GCATGCTCAGATCAAAAATATTTCCCACAACCCATTGCGGACTTTCTACATTGTAAACATAGGAGATGTTACGGTGTGCGGTATAGAGATATTTTCGTTCTACGGAAAACATCAACTTGGAAACGGCAACGGTATAAATATCGCATGTGTCAGAAGTTATCGGAGCTGTCTATGCTTACGTTATTTGTAGTCCTGTGATC

11801                                                                                                                                                                                                   12000
PflT_54_v1  AGACGTGACACCTCACACTCTGAAAAAGCCATAAAATAACTTACCTGCTGTATGTATGTTCACAACATATCACTGTACTGCCCTCAAAGTCATATCGATCACGGCTAGCACGTAACCATGCCATGTCAGCGTTGGATCAACTCTCTCCACCTGTTCTGTTCACCAGAACGCTGACTGTATGCGCTTGCGCACTTGATGTT
PflT_54_v2  AGACGTGACACCTCACACTCTGAAAAAGCCATAAAATAACTTACCTGCTGTATGTATGTTCACAACATATCACTGTACTGCCCTCAAAGTCATATGGATAACGGCTAGCACGTAACCATGCCATGTCAGCGTTGGATCAACTCTCTCCACCTGTTCTGTTCACCATAACGCTGACTGTATGCGCTTGCGCACTTGATGTT

12001                                                                                                                                                                                                   12200
PflT_54_v1  GCACGTGATCTAGGGCTGGGTCGTTTTGTACTATAAGGTGGGAGTTTTGAATACTGTATAACTTTTTGTCTCTATAGGACATTGTTACCGAGGTAGTGGA-AACCCAGGGATTATTCAAAATTAGCCATGGCATAGGATTACATGTAGAGATGTCAGAAAACATACTTGGAATCTAGTCAGTTTCCTGAAACTTCCACAA
PflT_54_v2  GCACGTGATCTAGGGCTGGGTCGTTTTGTACTATAAGGTGGGAGTTTTGAATACTGTATGACTTTTTGTATCTATAGGACATTGCCACCGAGGTAGTAGACAACCCAGGGATTATTCAAAAGTAGCCATGGCATAGGATTACATGTAGAAATGTCAGAAAACATACTTGGAATCTAGTCAGTTTCCTTAAACTTCCACAA

12201                                                                                                                                                                                                   12400
PflT_54_v1  ACGATTCAATGAAAGATGCTTAATTGAAAAAAATACCATCCCATCATGCTTTATTTTTAGAGAACGTGTTGGTAGTAACTTTTCCTTTGAGTTTGAATAGCCGCTGATGCAAAAATGCTGACTTAGCACTTTTTTCCACAAGTTTTAATAATCAAATAGGTGGCATCTTTATTATTGGATCAGTAACTTCATATCTGTTG
PflT_54_v2  ATGATTCACTGAAAGATGCTTTATTGAAAAAAATACCATCCCATCATGCTTTATTTTTAGAGAACGTGTTGGTAGTAACTTTTCCTTTGAGTTTGAATAGCCGCTGATGCAAAAATGCTGACTTAGCACTTTTTTCCACAAGTTTTAATAATCAAATAGGTGGCATCTTTATTATTGGATCAGTAACTTCATATCTGTTG

12401                                                                                                                                                                                                   12600
PflT_54_v1  TCTATATCGGAAATAGTACATGTAGTGATTAAATAACAATATTTTTATCCACCATGGTTATTTTTTTCTTTATAAGGGTCAAATTTGCAATAAAGTCGTGTTTGGTCTTTGTTACTCAAAATCATGGGTAATTTGCGATTTAATGTAAATTATCTCAAAAAGCAGAATGACAACACCAATTTT-CTAGTCATAAATTAAT
PflT_54_v2  TCTATATCGGAAATAGTACATGTAGTGATTAAATAACAATATTTTTATCCACCATGGTTATTTTTTTCTTTATAAGGGTCAAATTTGCAATAAAGTCGTGTTTGGTGTTTGTTACTCAAAATCATGGGTAATTTGTGATTTAATGTATATTATCTCAAAAAGCAGAATGACAACACCAATTTTTCTAGTCATAATATAAT

12601
PflT_54_v1  TCTCACCCTTCCAGGTAAAACATATGTGATTCTAGAAAAAATGACAATATTCTTGGAATTGGACAGTCCTCAACCTAA
PflT_54_v2  TCTCACCCTTCCAGCTGAAACATATGTGATTCTAGAAAAAATGACAATATTCTTGGAATTGGGCAGTCCTCAACCTAA
```

#### FASTA DNA

```
>PflTRAGL-A_54_v1
CGAGTAGTTTTTCTACCACGGGATCTATTATCGGAGTTTTCCTCGGCTCAGTCAGTCATAGTGTACATCACTCTTGCAGAGTCGTATTTGCCTTGATAATGAAGCATCGTTTTGTCCAAACGCCGTAGTAGAAATGCCGCATTTCAGGAAAGAACGGGATAAAATCAAAACAGTTCTGAAAGAACTGCAAATTCGCCTACTAGATGTCCAAATTTATC
GAGTGTGGTTATTTCAATGGACGACGTTGGACATTTACCACAATGCTCAACGGCCGAGTCGTCTACCAAATTTTGTAGAGCTTTTGTCTTACAACATATCCTTTCCTCATTCTACCTTAGTTTAGCGAGGCAGGGCTGTGTCTCCATTATAAAACGTAGTTTTCCACAGTTCTTAGATGGTAGTTGTAGTTTCCTTTTCAAAATTTCTTTGTCTCATGA
ACTCGGCGCTGTCGCGGATGGGAGGGTACCGACGACTATCCCCCGGCTTCGATTACAAGTTTTATGGTGACCGAAATGAAATGTGGCGTCGATTTGTTGTCTTATGTTTATTTCAAAAAGCTTCTGAAAAACACTTACTTTAGAAATCAAACCTGTTAATATTTAATCACCCAATTTTAAAATCTGCATAAAGTGATCGCATGATCTCTCGGAGCTTTC
ATTTCAATGTTCATCCCTCCTTACAAATGGTGAGTTCTAGTATTTCGCTACTTTGAACGATCCCGCCAAAATCGAGTCCACTCGCACTTCTACACTCTCCACGCAGTCGCGATACTAGTTCATTACAAACGCGTGGTAGTCAGAGTCAGACTGCTAATCATACTATGTCAACTGGCTTTATGAAGATAGCGGTCACTGATGAGCACTAGAGGTAGCCTC
GGGAATTTTCTCTGCCAGAATGAATTGGAGCTAGCTGTAATGGGGATCGAGCGCGGACATTTGAGTGCAACTTATGGGACTGATTGGTACGGTGACCTCAGCTGCCGCCGGCGACGTTTGTCAACTCGACGTCTGATCAGCATAAGTGACCCAGCAGTGACGACCGAATATGCCGAAGGCGATAGCAGGCGTCAAAAAAATTCCGGTTTTGTCAGCACA
TTGGATTCTGGAGGCTACTTGTTTTTTGTTACATAGGCATGCACGGCTTTAGATCGGGAAAGAAAATGTATATCACTTTTGGAAAAAGCTAAGCGTAAGCGATAGTTACGCATATCGCATTGCGATGATTGCGCTTTCAACAGTACGTTTACTCGAGGGTTAAACTTGAAATTTGAAAAGAAGACAGAAGATATACAAACAGTGCAGACAAAGTGAGAG
ATCATGCCAGAATTTGTGCTTGCACTTGTCCGTGGTCTCAACCCGAACATACAGACGTACTTTATGTCGGGCATAAATGACAGTATAGGCCCTCCTGTCTGTAAAATGTACGGCTCAGGTAGGCCAACATAAATAATCAAAGTAAAACTTAAGTTGCCCACGCGGTAATAGTTGAAAAATGCAAGATTAATTGAACAATATTATTTGCCACAATACAAA
TTTACCAAACGAAAAGAATTTGTGTACCTATAATATACAAACAGGCATGGGATCTCTGAAATTTCATGAGCTCAATGAAAGATCTAGACCAAAATATTTCTATTAACATGACATGACACATAAATATTAATCAGTCTTTTATTAGAGTAAACAAAGCAAGACATTATTCTTAGAATGGACAAATAATATGCATTATTCAAGTAACACCATACAATGCTA
TGAAAAATGCAATGAAAGAACAAACTTAATAAGTTACACAACAACTACAATAAAGTTAGAGAAATCTAAACTTGTGAAGTGTAACTTTGAAATTTTGTATAATTGTGATATGAATGATTCCAGATTAGTTGTTTTGTTTGACTGTTATGCATCGTGTAGGGGTAATGACATGATCAGATCAAAGTAAGTGAATTTGGTATCATTGACAGCAAAAGTTTA
TCAATTCTATGCCTGTATGTCAAATTTTCCAGTTGTTTTGTCTGAAAAAGCATCTTTCACTGTTGTTTCTATGCTCCCTGGAGCTGAATGCCTCAAGCTATAATCGGTTATTAATTATGATCAGCCGTTCAGGAATCTGTGGAGAGAAAAAGGGATGTATAGATAAAATAATAACTAATAATGTAAAGCTCATCAAACTTCACAAAATGTTTTCAATAA
ATATTACCAATGTAAGTATGGTGTAAACATAGCATGTTCATTATGCAAAAGTATGTTAAATGCAACTTGGATCAAATCCACTCCACTAAGTTTTGTTTATGGCCTAATTACTAAAGGCATTAAATGTGTAAATGAGCTAAGTATAATATGAGACACTCATGAAATTGAAGCATGAATAAAAGTATAGCTGCTGAAGTTAAGAAGTCCAATCGAATTAA
GTTTCAAATATCAGATTCACTTTCGTCATATAGCCGAAGTACAGAATTTCATAATTCTAAGTATAACCAATGTACCTGAGAAGTCTAATGAAATGTCTTTTGATACACTTCCTTATTAAAAATTTGCATACATATGCAAATAATCTATGGATTAAAATGTCATAGATTACTGCAACCAACACTTCATATGATCAACAAATGGCAAGGGACCCATTTAAA
TTGTTGCACTGGATATTGATACTAGCCTAATTACAGAAATTCATTAAACAAGTAAATGAGCTTGAATCAAAAATCTCACTGTCCGTGAATCGTTTATGTGCTGAATGCTACAAATGAAAATGAAAGCCCTGGAAACGGAGAATGAAGGGCAAGCATTCACATTTGTGTATATGTAAAATCCGAATCGGGTCAAGTATAGAACTGATGAAGTCCTTGCCA
TGGTGTGTTGTCTTTATTGTCTGGCCCGACACCGTTACGCTAAAATTACCAGGAGAATACTCACTAGGTGAGTATTATTGTAATATTGTATCAAACTGTAATCGGACCCATAGAGGTGGTAGAATCTGCATGCCGAAGCTTAGTCTCGCATGCCAGACCTCGAACCTGCATGCGGCGCGAGCGGGAACAGCAACACGAGTGTGACAGAACTATTGGTTC
```

AAAGTGAGAATAATCATGAAGCCTATTGGCTACAAGTCTTCCAAAGCCATGCGTTATTAACAAAGTCTGTTTTCGATGATGTTCACACGGTGACGTCATACAACTTTAGAATCGCATTTGAACGTCAGGCTATTACATGCAAAATACGAAACAAATGAAAATTGTTACGAACAACTATCGACAATTTCACAATCTTGCACGAACAAGACATACATCTGA
ACACAGTGGCGATATACGACACCGTTCAGGCCTCCGGCCCATGGCCATGCATGGTGTCCAAAAATATCGATCCGCTACAGAGTTATGGTTTATTCGTGAATGTTCATGATCCCGGGGCCGCTATCCGTGACTATGATGTATGACAGATGTCACATGAATGTATCACAGTCAAATTCATGTTGCTGATAAATCACGGCGATCGGCAGATTTCAGATTATA
ATAACGGTCAGAGCAGGAAGTCTGTCCGGGTTCTCGGAATTCAGTCTGCTTTACTCAGAAAAAGACTAGTCTCCATAGTTACAAAGCAAATTTCGGCCTTGACGTGGTGAGTATTCCGCTGTCTTCGAGAGGCAGAGACTATAAATTTAATTTTCTCTCGCTATCCCGGGCAATCACAAATGTACCGTGTGTGTTGATTTTCCGCATTTGACACCATCG
CTCGATCGTGTTTTTACTGTCGTGTACACAGCTATGAAAGGAATTTGCTTTGGCGTACGCGGTTGTTGCTACATGTAGTTGTTTTGCCAATGTGGGTTTCACGTTAATTACACGCCCTGTACAGGACAAACTGCCAAAGTTCACGATCACTTCACCAATGCGACCATCTAACAATTTATTCATTCGAATCACTGTTTTGTCAGTCCGCTGACATTTAGC
GACTCAAATTCTGCGGCTCATATAAAACATGGCAGGAACAAAAACATCGAATTACAAACACGATAACAGTCGAACATCGAGCGCGAGCTGGTGAACGCGAAGCGAGCACACAGCTCCTTGTAAAATAATATAAAATGATATCATTTAAAGTGGTAAGCTGGACACGAGAGAGAGAGAGAGAGAGAGAGAGAGAGAGAGAGAGAGAGAGAGAGAGAGAGA
GAGAGAGAGAGAGAGAGTGTGGATGAACTGGTAACTCTATCGTATCTTTCTCACGTTGATTTACATACCACAAAGATGCCACAAAAAGCCGATCGACTTTTTCCACTGAACACTGACACGTTAACATTCAAACGAAATAACACAAACTCACTTTTATTTTTTTTTTGGAGCCTGGGGAAATCTATGAAGTGGCATAAAGCAAAATCTGTATTGAGAAAA
GCGGTACACAGCGTAACATGTCAATATAGAAATCGATTAACACTACGATGCGTCCCAGAGAGTATCGACCAATCAGAGAGCAGCATTCATCAAGCCACGACCTTTCACGTATGGCTTCGAGATTTCGCGTGATCTAGGGTGTTTACTTCAGAAACTCTACAATCTACTACATACTAAGGGGTCTCAACTGTTTTACACATGCGCAAACGCCATCTTGAA
GAGTGGCTGATGATCATGTCGAAAACCGGAGGTCGAAAGACGTAAGTATCGCCGAGATATCGTAATCTGAATCTCGCGTTGCAATTTTTCACTCGATTATGTGTATCAAAATTATGCGGATACAATTTTGGGATTTTCTGTCAAACTTGTTTTTCCTGTTAAAGTTTGGGTTTTTCTCAGTCCGTAACGACATTGTTTACCGAAGGAACATGTGCTGTA
CGCTTGGACTTTCTTCAAATGGCTTGGACTTTTAGATCTATACCCATCCAACATGTGTGATTGAGTGTAATGGCCGTTACATTTAAAAATGTTCCTGTGATGGTAGCTAGGTCAGGGAAATGATTGTTTTGTGTTTAATATATTTTTAAGGTTTGAATGAGCGTGTTACGATATTTAGAATTCTGAGCAGCATCCTAGAGGGTCAATTTGGGTGTACAA
CGCATGGTTCATGCGGTAGCTAGCTAACCCAGGCCATATGGCACCAGGTACAGGTTGACTAACGAAATGTTTTGAGGTTAGCATAGTATGAGTACGTCAGATTCATATGTCATCGGCATAGAAGGTCGCTTGTGTAAGATAAGACCATATTTTTAGGCTGTCATTCATTTCATTTATTTATTTTGCAGTGGGTTGCTGTCAGCAATGAATCATTTACAA
GTTTGTTGCACTGTTATGCAATGCAATGCCTTGACCTGAATCAGTATTTCTTGCTTGGGTTCCTGCGGTATGAAGTAGAAATTAGGCTGGCCACGGTGTCCTGGGAATTAATGTTTTACCACTAAGGTAAGAATAAAATTCTATTGGATTTATTGTTTACACATATATAAGGATGAAAGGTACAGACCATTGTACTGATTGGTACTGTACTTGCCAAAT
TGATATGCTTAAATAATATTTTAAATATGTTAATTGTTTTCAGAGCGGA*ATGACATCAGCTTTTCAACACAGAGAGTACCTTAAGAGGGTGTGTCGTGTATGTGGTAGAACCCTGAAGACCACATCTTACAGCAGCTCAGGACATTACAGACAGGGAGAAGTGTGCATCCCATACAGTGAACTTTTAGAAGAATGCTTTAAAGTAAATGTTCGAAATGA*
*TGACCCTTTTGTACATCCGCCTAGATTTGGTTCATGTTGCCGTAGAGTTCTTTTAAAGTATTCTGTTGCACAGAACCTGGGTAAACATTACTATACCCATATCAGTGTTTGTAACTGGAAACATCATTGTAAGTATGAAAACCAAAGTTGTCCTCTTTGCAAACGTTTTTCTCAAAATGGTTCTGGTAGACCTAAAAAACTTAAAAGGGGTCGCTTTAA*
*ATTAAATACAAATGAAAAAAATGAAACTGACAAGAGTCCTGTTGCAGACAATTTGCTCATATACCAAATGACCACTGTTATACCACCCCAGCTTCCTGTAAGTTTGCCCAGTCTGTGCAATCAGACCAAAAGACAGAGACATATGAATTCTGTACAGAGGAAAATATTCAGACTGTACCCATGACAAGTCGATTTCCAACAGATTTATGTGTAACTGAA*
*CCACATCCAGAATTCATTTGTAGAATGTGTTTAGGAGTCCTAGACACACCACTTCTAACACCGTGTACTCACACTTTCTGTGCAGGCTGCATTAAGAACTGGTTAAGTCTGTGTCATTTTTGTCCTCTCTGTAAACAACCTATGGCGTCATCAGATTTAAAAGAACCATATAGAGGATTTGTAGAAATTCTTAATGATACTAGAATGAAATGTAATTGT*
*CCAGAAGCAGTTTCAAGCAAAAAAGGTGTCAAGCTTAGTAATTGGCTTCAGCACACTTTAAATTGTGGAAAGGATTTTGAAATTGATGAATGGTTAGAAAAAGTTTTGTCATTGTGGAACGAAATTTCTCTCACTCAAGTTGATTCACATTTAAATAGTACTGAACATTTGCCTGAGACAGATAAGTCAAATCTGACAACAAGCCATGGCCATCCAGAC*
*ATTGCTGCAGGTAGCACTGTTGGAAGACCTAAAACTTCACTGTTGTGTCTCCAACAACAAGGACAAAATTTAAGATTGAAAAATATCAAAAAACAAATAGACGTGTTTGCCAGAGAGAAAGGGGAAGATCTCAGGGTAGTATATTTTGGCCTTTTGCAAAACTATCTGAGAGCCAACAAGCAAGGTAGTTTAGCTGAACAAGTGAAGTTGCTATATAAT*
*GGGGAATCTTGTCAAATGTCACCTCAGGAATGTCTAGCCTTGCGTGTAAATACTTTTACCAGTTGTAATCAGTACAAGAAAATTTACTCAGCTACAAATGAGCTTAGTGGAGTTAAAATCTTCCAACCTCTCATCAATGTTGTGAAAGCCGAAGAAAGTTTTCTACCAGGGTCTGTAAAGTTCTACTTTAACCCATCTTTGAAAACTGTTTCTCCCATT*
*TGGCAAGGTACAGAGCATGTTGATGCTCTGTCAGGACTCAGTCGCGATATACCACTTCTAGAATCACCTGATGTTTATGGGGCAAGGTATAGATATGACAAAGCAGTTGCTATGGCTTTGAAAGATATTGATAATGAAATCTCAGAAGGCCTCAGTAAATTAGAAGGGGTGGATACAAATCAACTTGTTCTAACAGCCACTGTCAAAGATGGCAGTGAT*
*GGTCTTGGAGATGTGAAAGAAGTCAGATCTCAGGGGCTTGTCGTTCCAAACAAGGCCTTGCGGTTTTCATTTGTAATTTTAGAAGTCACTGTCAATGTAGATGGCAAGATTATTACTGTATTTGAAGAAGGTTCTCCCAACTCAGAATTATGCACACGCCCTCTCTTGGTAGCTCTGGCAGATGAAAATGACAGAAAAGGGTTATGCATGACACATATA*
*CCAATAATGATAGAGAGAGAGGAGATGGAGAAAAGTACCTTGTTTGTAGAAATTGAAGAGGGAAACAAAGAATCTACAAAGTTGTATTTAAAGGGACTATGTATGATGAAAAATTGCAGAGAAGTGGTGAGGGAATGTTAGCGGCAGGGTCATCTTGGTTTTGTACACTTTGTGAGAAAGGAAGACTCGATCCCATATCATCAGTCTTGCCAATAACAA*
*GAACACATGAAAACAATATGGCTTTGTATTCAAAGTATGTTGAAAATGCAAATCATATGAAGCAGGCAGATCTAGTGAAAGAAGTCAAAGGCATTGTTAGAAAACCATTTATAATGTCAGAGCCTTGTGTTGATGCCACGCATGCTGAAATTCACTGGGGAGAGAAAAACTATGATATGTATGTACGTGAAATATCAGGGATAATGTGTTGGGGAAGC*
*CATCCAATGATGATGATGCAAAAAAAGTGAAATCAACAAAAAGAATGTTGGATACCGAATTACAGCTGCAATGTGGCTTAAAGAGAGAGCATGGCTTTATGGTTGATGGAAATTATGCACGTGATTTGGTCAAATCAGAAACAGTTGATGTGGTTTGCTCTTTAATTCAGAGTGAAGCAAGAAAATCTGTTGTTCGTGAATATGTTTCCTACTATCGAG*
*AATTGCGATCAATATACAGGAAACACAATGCTGACTTAGATGATGCCGTGAAATTTAAAACCATAGCTGCAATTTTTTACAAATTTCTACATGAGAAATTTTCTTATTTGCATCCACTCTCTAACTATCAACACAAGGTATTAGATCACATTCCACAGTTGATCGAGCAATATGGCTCTGTTGGTAAATTTGCCTCTGAAGGCAATGAAAAAGGCAATA*
*AACTTTTCAGAAGATTTCGTAAATTTAATGCACGTAAGTCTCAAATTCATGAAATGCCAGACATTCTGAAATTTCATTGGCTATATACAATGAAAAGTCTTCAGACTATGGTAAACTGGGAAAAAACTAGATCACTAGTCTGTAGCAAATGTGGTTGTGCTGGTCACAATCAAAGGACATGA*TATGAAATCAAAGATGAGACAATGACCATGCATGTTT
TATGTATCTGAATCTGTAAGTTTGAAATTAAGTAGCATCTGTATGTATCAGACTACAGAGTCCAAACTTCAGACACAAAGAGTAGGCTGGCCATACACTCTTATTTACTTAAAAAGTGTTTCTGTTCCATATACTGACTGTACTTTCAAGAGTCAATTCTAGCTTCTATGCTAGTCTTAATATACCTAGATGGTTAAAGTGTATTATATTTTGATGTAC
ATAGATGAATTCCACAAGAAATCATCAAATACAAGAAAGCTGAGTTTGTGTTAAATATTTGTATGTGTTTTAATCTTACCAGTAACCTGCCCACACTTTTGAAGGGACTCAAACAAAATCATTTGATTACTTTGGCAGAAAAGCCAAAAGTTTTAGTTATACTGGCCATAAAAAATGAGATGAGAAGCGTAAAGTTTGCAATAGAATTTTGTACATATA
TTCATGAAAGATTGTAACAAGTTGTAAAATGAACTCTCAGTAAAGGTTTCATTCAATTCATGCACATGCTACATGTACATGCTATGATTTCTTGTGCAGTTGTCATGTTC*TCAGAAATTACAATCAGAACATGTAAATTTGATGTTACTCTTATTTCCAGTTGTCGTGAAGAAAATCGTTTGCCAGAGCATTTAATGTGAAAAGCTCTAACACATTGT*
*GTACATCGCACTGTTGAAAAGTAACTCTTATCAGTCACGCTACAATTAATGTTAGCACATGTTGTGAAACACGAAGGGCATTCAAAGTTTATACAGTCCAGCTTTGTTCTATTTAAGTGATCTAATTCAACACATTTGTCATGGTACCACCTTCCACAGAGTACAGTTTGATTGTTCATGATGTAAGTTCGATCACATTTCAGCCAGTTGAACTTGGCT*
*TGTTCAGTAGCTGCTAAATTGGCAATTTGCAGCCATCCAACATACATTCCCATGTTTCCTCCTCACTATCATCTGATGTATTGTCTATCTTTCCTGCTCTGCTGCCCTCTTCTGACTCATTCCTGTCTGTGCTAACTTTGACCTCAGCTGCCTCACTTATTCCGTTCTGATCCTCAGTCTCTGAGCCTATACTTTCCTGATCTGTCATGTCTATGCTTA*
*TCCTTTCCTGCTCTTCCTCCGCTATGCTTACTCTGTCCTCATCTGTTGACCTTTCCTCCTCATCCCCCAACTGACTGTTATTACTCTGTGTCCTGGTTTCAGAGCTTGTCCCTTCATTTTGAAAACTGAATTTTATCATTTGCTTTTTTACTGTATCACAAACAGCAATTGAGCTGAAGAATGGAAGACTGTAGTATTTACTACTTAAGGTATCATAGT*
*GTGCTGTTTCTATGAAATCTATGGCATCACAAGGTGACTTGTAAACTGTGTTGTTACTCAATATGTAAACAGTGTCATTACATTTGAATGAGACTGGGTTAGGAATACCAATAAATGGTACAGATGTACAGACCATTGAATTCAGGTCAATTACGTCAACATCGGGTGGATGTCTTACATTGTTACCTATCACATAAATTCCACCAACAATATACATTT*
*TACTGCTATTTAGAAAAACTGAATGGTAGGCTCTTGGAGTGATAGCTTGTAAACTAATACTTGATAATTTCATAGATTGTAAATTAAGAACAGAAAATATGTTTGAAGCCATGGACACATTGTAAAAAGCATGACTGAGTTTTTGTGTATATTGTAAGCCTCCATAAAGATAAAGAGAATCTCCAAACTTAATTACACTGTGACCAAAAGCATGAATTT*
*CCTCACCACTATATTGTCCTTGAGACCATGTTAATGAAATTTTCCCCCTTGGATCAACAAAGACAGAAAGCTGCCACAATTTTGTATTGTGCATTTCTGAGTCAGTCAGTCCACCAAGGAGAACATAATGGTTTGGTTCAACTAAACACAAATTTGAGCCATGTACATGTGGAGGGTAATTTGACTAAGATGAATAATTTTGTGATTTCTTGTATTGTA*
*TCATCTTTGTCTATTACAAGCCAGCCTGTTACTGTTGTCTTAGACTTAACCCAGACCAAAGCTCTGTCACCTAAAGAAATAACTGGATTTGTACATCCAGCAGTTGCTAGGCAAGTTGCCAGGATGTTTTTGGTGATTTGTATTTCT*GAAATAAAAAAGATGATGACAAAAGTTGTTGGATTAGAGAGTGAAGCTTATCTGCATGCATGTATGGTGAGC
AGGTCATTTTATTTTGTCATTGTTGAAATATGGCTCTGAAATATAAATTTGTTAATTTCATATTTTAATATGGGTATCTTTGTAGTTTAAAAAAATGCTCAAGACAGAAAGTGAGATGTCTTTAGTCGTGACATTTGTTAATATTGTTCACTGTACATTTCTGTACCCTTTGCAACCATACAATTGTTGAACACCTGTCCTGTATATTCAAACAAACAA
CTTGAAGGCAAATCTGATATCTTTGGGGCAGCAAAATACATGTGCTCTTGTCTTTTGCATTTGAGTAACCTCCTTATATAAATATGATCGAAAGAGATTGTCAACATGGAGCAGCAAATCCGTTTCTTTAACCTAGACTGTAAATCAATACAAAATGGGTGCACATGTAGCTAGGGGAAAAGGTAATGCACAGATAGAGCCCCATTGATGACTGCTGGA
AATGTCATTTGGATGTCTTGCTGGTGTACATGCAACTGAAAAAATCATTTTAGCCTGTGGGCCGTACAATAGCTGCACTTCTACTTCGCTCACTCTAATTTGAATCTTCGTATATAGTGTACATGGGGATCGACTTTCATAGTATGCTTGTTGGATCGAGGGACAGCGATCGACCGTGGCTTGTGTTACAGTTGTGAACGATCACATCGAGAATTTTCT
TTTTCGAATGGGCGATTACAAGTGCAAATCTTTGGAAGTTAGCTACCATATAAGAACTTTGTTGTTTTAACTTCAAGAATAATCAATCTAAAACACGAACAGAATGTTCTGTGACTTTTGAAGCTACATTGAACTGACGACCTACCGGGACTATCGAGCATGGCGGCTGCCATCTTGGATATTGTTTACGGCTCACTGAGCATGTTCAGATCAAAAATA
TTTCCCACACCCATTGCGGACTTTCTACATTGTAAACATAGGAGATGTTACGGTGTGCGGTATAGAGATATTTTCGTTCTACGGAAAACATCAACTTGGAAACGGCAACGGTATAAATATCGCATGTGTCAGAAGTTATCGGAGCTATCTATGCTTACGTTATTTGTAGTCCTGTGATCAGACGTGACACCTCACACTCTGAAAAAGCCATAAAATAAC
TTACCTGCTGTATGTATGTTCACAACATATCACTGTACTGCCCTCAAAGTCATATCGATCACGGCTAGCACGTAACCATGCCATGTCAGCGTTGGATCAACTCTCTCCACCTGTTCTGTTCACCAGAACGCTGACTGTATGCGCTTGCGCACTTGATGTTGCACGTGATCTAGGGCTGGGTCGTTTTGTACTATAAGGTGGGAGTTTTGAATACTGTAT
AACTTTTTGTCTCTATAGGACATTGTTACCGAGGTAGTGGAAACCCAGGGATTATTCAAAATTAGCCATGGCATAGGATTACATGTAGAGATGTCAGAAAACATACTTGGAATCTAGTCAGTTTCCTGAAACTTCCACAAACGATTCAATGAAAGATGCTTAATTGAAAAAAATACCATCCCATCATGCTTTATTTTTAGAGAACGTGTTGGTAGTAAC
TTTTCCTTTGAGTTTGAATAGCCGCTGATGCAAAAATGCTGACTTAGCACTTTTTTCCACAAGTTTTAATAATCAAATAGGTGGCATCTTTATTATTGGATCAGTAACTTCATATCTGTTGTCTATATCGGAAATAGTACATGTAGTGATTAAATAACAATATTTTTATCCACCATGGTTATTTTTTTCTTTATAAGGGTCAAATTTGCAATAAAGTCG
TGTTTGGTCTTTGTTACTCAAAATCATGGGTAATTTGCGATTTAATGTAAATTATCTCAAAAAGCAGAATGACAACACCAATTTTCTAGTCATAAATTAATTCTCACCCTTCCAGGTAAAACATATGTGATTCTAGAAAAAATGACAATATTCTTGGAATTGGACAGTCCTCAACCTAAAATCAATAATAAATAAGTGTTTTATGCCATTATTGGCTGT
TTATTTGGGGCTATGGATAATTAAAAGAAGCGAACGGCAACGCAACTTCCACTCTTAGGGGAGAGGGTTTGGCTGTATTTCCATTATCGTGTGCAAAAATATCACTACATGGTTTTTTGTATCGCAACATCTCGCTACACGTTTTATGTATTACAAAATATCGCTCTTTTGTTTTGCCTGTACAAGGCCAGACGGGCATGCACTGGCGCCACACCATTC
TTTTGACATATATGTGGTTCAGTGCATGTTTAAAATAACGTACCAATCATTTTGGAAATACTGTTTCAATTCATTCATTCTATTGGTTGTGTCATTGTCAAGTTTTCGCTGAACAGAGATTGATTTAAACGGGGTATACCAGGCTGATGGGAGTGTAGGTTTGATTTTATAGCGCCGTAATTACTGATTGTAGGTATGCTGATGTAGAATGAGGATGA
GACTTGGTTAGATTTTCACCCTTCCAAACTTCTTTTGGAGGGGATGAGGGCAACACAATTAGTTTTGTTTACCATGCACAAGCTTTAATTATCCACGGCCTCTTAGCTTTTAGGCAATTACTCTTTAATTAGGTTTGCTGTCACAAAATAGACATGAACACGCTATTATGAGCAATATAAGACACTTACATTAGAAAAATTCCTAAATTTTGTGAAGGT
TTTGACCGTTGGTAATTTTGTCTGACATATTCACAGTATTGTATTTGTGGTATTTACACTAGATATGAAATGAAAGTGCTCCTGTGTTATTGTATATGTTGCTGTTGTCTGTAATTTGACACTTTTCTCCAAGGTTATGCCATTAGATTTTAACTTGATTGAAATGCTTATGTTTAACATCACTAGCAATATCCATTGTGTATAGCTTCAATTTAATAT
CTAAATGTGAAAGATCGGAAAATATGTGAAATACAAATTAGATGTGAAATTGCCTAATCTATTCCATACACAAGACAGTGAAACAAAAATACACACTACTATACATTAAACGCAAATCACCAATATATGAACGATGTGTGGAGTTGCTTTCCATTTATTGACTAATTTCTTTCACTTCTACCCATAACAGTCCGTCCAGTTAAGGTAGTTATATGTGCA
AATCGGCATATATAGGTGATATCACAATGCTAAATGTCTTTCACCACAATACGTAATCATATCATCAAAACAGCAAAATTCATAAACTTTTGTTATATTCTTCCTTCATTCCTTCTTTTCCCTAAAAGCATTTAATATGCAAACTACTAAAATGCAGATTTAAAACTGTTACATGAACATATTAGTGTCGTATCATAGAATCGTCAATATACATGGCAT
TTTGCCGACTTTGGTAATCTATACAGATATATATTCCAAATTAGAAAAGTCTCTTAAATACCTTAGTTACTTATTAGGTTTGTAAAAATGCTAAATGACTTTCACTTATGTTATATCAGAGTATTTTCAATTTATATAGCAAGTTTTGTCAACTTTGGTCAGTCAATCCAGATGTATATCTGTAATAAAGAAAGTTCATTACATATACAAACAAACAGT
AACATTGTCAAAATTCTTAAACAGACAAACAAACAGACAAACACTAATACAGACATACACAGAGAATTAAAAACATCAGTACCACAATAGCATTCGAATGAACACACGAGCTAAACATCATAGTCACAGTAAATACTATTTATCCAAGTCCTGTGTGCCCCACTACACATAATTTAGTGCATCCTCCCTTGCTATGTAAATGAGCAAAAATAAGTCAAA
ATATTACTTCAAAAATTAATTTCTGCTGAATTATGCAACCAACAGCCCCAAAAACTTTATTAGATATAGTTTCCATCCCAAAGATTCTGTGTACCTAATTTCATGACATTTGATGAAGGAGTTTGAGAGATATGGCCCATAACACACACACAGACACACACACATACATATTACAGACACACATACAGACTAACAAACGCCACCCTATGACAATGGCCC
ACTTAGGTGGGATATGCCCACCGCGTTGGCCAAAAAACGGTCCTTTTAAGGACCAACAAATTTTCCTAAAAGAGAACTACCTAATCA

>PflTRAGL-A_54_v2
TGACCCCGGGGCCGCTATCCGTGACTGTGATGTTTGACAGATGTCACATGAATGTATCACAGTCAAATTCATGATGCTGACAAATCGCTGCGATCGGCAGATTTTAGATTATAATAACGGTCAGAGCAGGAAGTCTGTCCGGGTTCTCGGAATTCAGTCTGCTTTACTCAGAAAAAGACTAGTCTCCATAGTTACAATGCAAATTTCGGCCTTGACGTG
GTGAGTATTCCGCTGTCTTCGAGAGGCAGAGACTATAAATTTAATTTTCTCTCGCTATCCCGGGCAATCACAAATGTACCGTGTGTGTTGATTTTCCGCATTTGACACCATCGCTCGATCGTGTTTTTACTGTCGTGTACACAGCTATGAAAGGAATTTGTTTTGGCGTACGCGGTTGTTGCTACATGTAGCTGGTTTGCCAATGTGGGATTCACGTTA
ATTACACGCCCTGTACAGGACAAACTGCCAAAGTTCACGATCACTTCACCAATGCGACCATCTAACAATTTATTCATTCGAATCACTGTTTTGTCAGTCCGCTGACATTTAGCGACTCAAATTCTGCGGGTCATATAAAACATGGCAGGAACAAAAACATCGAATTACAAACACGATAACAGTCGAACATCGAGCGCGAGCTGGTGAACGCGAAGCGAG
CACACAGCTCCTTGTAAAATAATATAAAATGATATCATTTAAAGTGGTAAGCTGGACACGAGAGAGAGAGAGAGAGAGAGAGAGAGAGAGAGAGAGAGAGAGAGAGAGAGAGAGAGAGAGAGAGAGAGATGTGGATGAACTGGTAACTCTATCGTATCTTTCTCACGTTGATTTACATACCACAAAGATGCCACAAAAAGCCGATCGACTTTTTCCACT
GAACACTGACACGTTAACATTCAAACGAAATAACACAAACTCACTTTTATTTTTTTTTTGGAGCCTGGGGAAATCTATGAAGTGGCATAAAGCAAAATCTGTATTGAGAAAAGCGGTACCGACTCACAGCGTAACATGTCAATATAGAAATCGATTAACACTACGATGCGTCCCAGAGAGTATCGACCAATCAGAGAGCAGCATTCATCAAGCCACGAC
CTTTCACGTATGGCTTCGAGATTTCGCGTGATCTAGGGTGTTTACTTCAGAAACTCTACAATCTACTACATACTAAGGGGTCTCAACTGTTTTACACATGCGCAAACGCCATCTTGAAGAGTGGCTGATGATCATGTCGAAAACCGGAGGTCGAAAGACGTAAGTATCGCCGAGATATCGTAATCTGAATCTCGCGTTGCAATTTTTCACTCGATTATG
TGTATCAAAATTATGCGGATACAATTTTGGGATTTTCTGTCAAACTTGGTTTTTCCTGTTTAAAGTTTGGGTTTTTCTCAGTCCGTAACGACATTGTTTACCGAAGGAACATGTGCTGTACGCTTGGACTTTCTTCAAATGGCTTGGACTTTTAGATCTATACCCATCCAACATGTGTGATTGAGTGTAATGGCCGTTACATTTAAAAATGTTC[2]CTGT
GATGGTAGCTAGGTCAGGGAAATGATTGTTTTGTGTTTAATATATTTTTAAGGTTTGAATGAGCGTGTTACGATATTTAGAATTCTGAGCAGCATCCTAGAGGGTCAATTTGGGTGTACAACGCATGGTTCATGCGGTAGCTAGCTAACCCAGGCCATATGGCACCAGGTACAGGTTGACTAACGAAATGTTTTGAGGTTAGCATAGTATGAGTACGTC
AGATTCATATGTCATCGGCATAGAAGGTCGCTTGTGTAAGATAAGACCATATTTTTAGGCTGTCATTCATTTCATTTATTTATTTTGCAGTGGGTTGCAGTCAGCAATGAATCATTTACAAGTTTGTTGCACTGTTATGCAATGCAATGCCTTGACCTGAATCAGTATTTCTTGCTTGGGTTCCTGCGGTATGAAGTAGAAAATTAGGCTGGCCACGGT
GTCCTGGGAATTAATGTTTTACCACTAAGGTAAGAATAAAATTCTATTGGATTTATTGTTTACACATATATAAGGATGAAAGGTACAGACCATTGTACTGATTGGTACTGTACTTGCCAAATTGATATGCTTAAATAATATTTTAAATATGTTAATTGTTTTCAGAGCGGAATGACATCAGCTTTTCAACACAGAGAGTACCTTAAGAGGGTGTGTCGT
GTATGTGGTAGAACCCTGAAGACCACATCTTACAGCAGCTCAGGACATTACAGACAGGGAGAAGTGTGCATCCCATACAGTGAACTTTTAGAAGAATGCTTTAAAGTAAATGTTCGAAATGATGACCCTTTTGTACATCCGCCTAGATTTGGTTCATGTTGCCGTAGAGTTCTTTTAAAGTATTCTGTTGCACAGAACCTGGGTAAACATTACTATACC
CATATCAGTGTTTGTAACTGGAAACATCATTGTAAGTATGAAAACCAAAGTTGTCCTCTTTGCAAACGTTTTTCTCAAAATGGTTCTGGTAGACCTAAAAAACTTAAAAGGGGTCGCTTTAAATTAAATACAAATGAAAAAAATGAAACTGACAAGAGTCCTGTTGCAGACAATTTTGCTCATATACCAAATGACCACTGTTATACCACCCCAGCTTCC
TGTAAGTTTGCCCAGTCTGTGCAATCAGACCAAAAGACAGAGACATATGAATTCTGTACAGAGGAAAATATTCAGACTGTACCCATGACAAGTCGATTTCCAACAGATTTATGTGTAACTGAACCACATCCAGAATTCATTTGTAGAATGTGTTTAGGAGTCCTAGACACACCACTTCTAACACCGTGTACTCACACTTTCTGTGCAGGCTGCATTAAG
AACTGGTTAAGTCTGTGTCATTTTTGTCCTCTCTGTAAACAACCTATGGCGTCATCAGATTTAAAAGAACCATATAGAGGATTTGTAGAAATTCTTAATGATACTAGAATGAAATGTAATTGTCCAGAAGCAGTTTCAAGCAAAAAAGGTGTCAAGCTTAGTAATTGGCTTCAGCACACTTTAAATTGTGGAAAGGATTTTGAAATTGATGAATGGTTA
GAAAAAGTTTTGTCATTGTGGAACGAAATTTCTCTCACTCAAGTTGATTCACATTTAAATAGTACTGAACATTTGCCTGAGACAGATAAGTCAAATCTGACAACAAGCCATGGCCATCCAGACATTGCTGCAGGTAGCACTGTTGGAAGACCTAAAACTTCACTGTTGTGTCTCCAACAACAAGGACAAAATTTAAGATTGAAAAATATCAAAAAACAA
ATAGACGTGTTTGCCAGAGAGAAAGGGGAAGATCTCAGGGTAGTATATTTTGGCCTTTTGCAAAACTATCTGAGAGCCAACAAGCAAGGTAGTTTAGCTGAACAAGTGAAGTTGCTATATAATGGGGAATCTTGTCAAATGTCACCTCAGGAATGTCTAGCCTTGCGTGTAAATACTTTTACCAGTTGTAATCAGTACAAGAAAATTTACTCAGCTACA
AATGAGCTTAGTGGAGTTAAAATCTTCCAACCTCTCATCAATGTTGTGAAAGCCGAAGAAAGTTTTCTACCAGGGTCTGTAAAGTTCTACTTTAACCCATCTTTGAAAACTGTTTCTCCCATTTGGCAAGGTACAGAGCATGTTGATGCTCTGTCAGGACTCAGTCGCGATATACCACTTCTAGAATCACCTGATGTTTATGGGGCAAGGTATAGATAT
GACAAAGCAGTTGCTATGGCTTTGAAAGATATTGATAATGAAATCTCAGAAGGCCTCAGTAAATTAGAAGGGGTGGATACAAATCAACTTGTTCTAACAGCCACTGTCAAAGATGGCAGTGATGGTCTTGGAGATGTGAAAGAAGTCAGATCTCAGGGGCTTGTCGTTCCAAACAAGGCCTTGCGGTTTTCATTTGTAATTTTAGAAGTCACTGTCAAT
GTAGATGGCAAGATTATTACTGTATTTGAAGAAGGTTCTCCCAACTCAGAATTATGCACACGCCCTCTCTTGGTAGCTCTGGCAGATGAAAATGACAGAAAAGGGTTATGCATGACACATATACCAATAATGATAGAGAGAGAGGAGATGGAGAAAAGTACCTTGTTTGTAGAAATTGAAGAGGGAAACAAAGAATCTACAAAGTTGTATTTAAAGGG
ACTATGTATGATGAAAAATTGCAGAGAAGTGGTGAGGGAATGTTAGCGGCAGGGTCATCTTGGTTTTGTACACTTTGTGAGAAAGGAAGACTCGATCCCATATCATCAGTCTTGCCAATAACAAGAACACATGAAAACAATATGGCTTTGTATTCAAAGTATGTTGAAAATGCAAATCATATGAAGCAGGCAGATCTAGTGAAAGAAGTCAAAGGCATT
GTTAGAAAACCATTTATAATGTCAGAGCCTTGTGTTGATGCCACGCATGCTGAAATTCACTGGGGAGAGAAAAACTATGATATGTATGTACGTGAAATATCAGGGATAATGTGTTGGGGGAAGCCATCCAATGATGATGATGCAAAAAAAGTGAAATCAACAAAAAGAATGTTGGATACCGAATTACAGCTGCAATGTGGCTTAAAGAGAGAGCATGGC

TTTATGGTTGATGGAAATTATGCACGTGATTTGGTCAAATCAGAAACAGTTGATGTGGTTTGCTCTTTAATTCAGAGTGAAGCAAGAAAATCTGTTGTTCGTGAATATGTTTCCTACTATCGAGAATTGCGATCAATATACAGGAAACACAATGCTGACTTAGATGATGCCGTGAAATTTAAAACCATAGCTGCAAATTTTTACAAATTGCTACATGAG
AAATTTCCTTATTTGCATCCACTCTCTAACTATCAACACAAGGTATTAGATCACATTCCACAGTTGATCGAGCAATATGGCTCTGTTGGTAAATTTGCCTCTGAAGGCAATGAAAAAGGCAATAAACTTTTCAGAAGATTTCGTAAATTTAATGCACGTAAGTCTCAAATTCATGAAATGCCAGACATTCTGAAATTTCATTGGCTATATACAATGAAA
AGTCTTCAGACTATGGTAAACTGGGAAAAAACTAGATCACTAGTCTGTAGCAAATGTGGTTGTGCTGGTCACAATCAAAGGACATGATATGAAATCAAAGATGAGACAATGACCATGCATGTTTTATGTATCTGAATCTGTAAGTTTGAAATTAAGTAGCATCTGTATGTATCAGACTACAGAGTCCAAACTTCAGACACAAAGAGTAGGCTGGCCATA
CACTCTTATTTACTTAAAAAGTGTTTCTGTTCCATATACTGACTGTACTTTCAAGAGTCAATTCTAGCTTCTATGCTAGTCTTAATATACCTAGATGGTTAAAGTGTATTATATTTTGATGTACATAGATGAATTCCACAAGAAATCATCAAATACAAGAAAGCTGAGTTTGTGTTAAATATTTGTATGTGTTTTAATCTTACCAGTAACCTGCCCACA
CTTTTGAAGGGACTCAAACAAAATCATTTGATTACTTTGGCAGAAAAGCCAAAAGTTTTAGTTATACTGGCCATAAAAAATGAGATGAGAAGCGTAAAGTTTGCAATAGAATTTTGTACATATATTCATGAAAGATTGTAACAAGTTGTAAAATGAACTCTCAGTAAAGGTTTCATTCAATTCATGCACATGCTACATGTACATGCTATGATTTCTTGT
GCAGTTGTCATGTTCTCAGAAATTACAATCAGAACATGTAAATTTGATGTTACTCTTATTTTCCAGTTGTCGTGAAGAAAATCGTTTGCCAGAGCATTTAATGTGAAAAGCTCTAACACATTGTGTACATCGCACTGTTGAAAAGTAACTCTTATCAGTCACGCTACAATTAATGTTAGCACATGTTGTGAAACACGAAGGGCATTCAAAGTTTATACA
GTCCAGCTTTGTTCTATTTAAGTGATCTAATTCAACACATTTGTCATGGTACCACCTTCCACAGAGTACAGTTTGATTGTTCATGATGTAAGTTCGATCACATTTCAGCCAGTTGAACTTGGCTTGTTCAGTAGCTGCTAAATTGGCAAATTTGCAGCCATCCAACATACATTCCCATGTTTCCTCCTCACTATCATCTGATGTATTGTCTATCTTTCC
TGCTCTGCTGCCCTCTTCTGACTCATTCCTGTCTGTGCTAACTTTGACCTCAGCTGCCTCACTTATTCCGTTCTGATCCTCAGTCTCTGAGCCTATACTTTCCTGATCTGTCATGTCTATGCTTATCCTTTCCTGCTCTTCCTCCGCTATGCTTACTCTGTCCTCATCTGTTGACCTTTCCTCCTCATCCCCCAACTGACTGTTATTACTCTGTGTCCT
GGTTTCAGAGCTTGTCCCTTCATTTTGAAAACTGAATTTTATCATTTGCTTTTTTACTGTATCACAAACAGCAATTGAGCTGAAGAATGGAAGACTGTAGTATTTACTACTTAAGGTATCATAGTGTGCTGTTTCTATGAAATCTATGGCATCACAAGGTGACTTGTAAACTGTGTTGTTACTCAATATGTAAACAGTGTCATTACATTTGAATGAGAC
TGGGTTAGGAATACCAATAAATGGTACAGATGTACAGACCATTGAATTCAGGTCAATTACGTCAACATCGGGTGGATGTCTTACATTGTTACCTATCACATAAATTCCACCAACAATATACATTTTACTGCTATTTAGAAAAACTGAATGGTAGGCTCTTGGAGTGATAGCTTGTAAACTAATACTTGATAATTTCATAGATTGTAAATTAAGAACAGA
AAATATGTTTGAAGCCATGGACACATTGTAAAAAGCATGACTGAGTTTTTGTGTATATTGTAAGCCTCCATAAAGATAAAGAGAATCTCCAAACTTAATTACACTGTGACCAAAAGCATGAATTTCCTCACCACTATATTGTCCTTGAGACCATGTTAATGAAATTTTCCCCCTTGGATCAACAAAGACAGAAAGCTGCCACAATTTTGTATTGTGCAT
TTCTGAGTCAGTCAGTCCACCAAGGAGAACATAATGGTTTGGTTCAACTAAACACAAATTTGAGCCATGTACATGTGGAGGGTAATTTGACTGAGATGAATAATTTTTGATTTCTTGTATTGTATCATCTTTGTCTATTACAAGCCAGCCTGTTACTGTTGTCTTAGACTTAACCCAGACCAAAGCTCTGTCACCTAAAGAAATAACTGGATTTGTACA
TCCAGCAGTTGCTAGGCAAGTTGCCAGGATGTTTTTGGTGATTTGTATTTCTGAAATAAAAAAGATGATGACAAAAGTTGTTGTATTAGAGAGTGAAGCTTATCTGCATGCATGTATGGTGAGCAGGTCATTTTATTTTGTCAGTGTTGAAATATGGCACTGAAATATAAATTTGTTAATTTCATATTTTAATATGGATATCTTTGTAGTTCAAAAAAA
TGCTCAAGACAGAAAGTGAGATGTCTTTAGTCGTGACATTTATTAATATTGTTCACTGTACATTTCTCTACCCTTTGCAACCATACAATTGTTGAACACCTGTCCTGTATATTCAAACAAACAACTTGAAGGCAAATCTGATATCTTTGGGGCAGCAAAATACATGTGCTCTTGTCTTTTGCATTTGAGTAACCTCCTTATATAAATATGATCGAAAGA
GATTGTCAACATGGAGCAGCAAATCCGTTTCTTTAACCTAGACTGTAAATCAATACAAAATGGGTGCACATGTAGCTAGGGGAAAAGGTAATGCACAGATAGAGCCCCATTGATGACTGCTGGAAATGTCATTTGGATGTCTTGCTGGTGTACATGCAACTGAAAAAATCATTTTAGCCTGTGGGCCGTACAATAGCTGCACTTCTACTTCGCTCACTC
TAATTTGAATCTTCGTATATAGTGTACATGGGGATCGACTTTCATAGTATGCTTGTTGGATCGAGGGACAGCGATCGACCGTGGCTTGTGTTACAGTTGTGAACGATCACATCGAGAATTTTCTTTTTCGAATGGGCGATTACAAGTGCAAATCTTTGGAAGTTAGCTACCATATAAGAACTTTGTTGTTTTAACTTCAAGAATAATCAATCTAAAACA
CGAACAGAATGTTCTGTGACTTTTGAAGCTACATTGAACTGACGACCTACCGGGACTATCGAGCATGGCGGCTGCCATCTTGGATATTGTTTACGGCTCACTGAGCATGCTCAGATCAAAAATATTTCCCACAACCCATTGCGGACTTTCTACATTGTAAACATAGGAGATGTTACGGTGTGCGGTATAGAGATATTTTCGTTCTACGGAAAACATCAA
CTTGGAAACGGCAACGGTATAAATATCGCATGTGTCAGAAGTTATCGGAGCTGTCTATGCTTACGTTATTTGTAGTCCTGTGATCAGACGTGACACCTCACACTCTGAAAAAGCCATAAAATAACTTACCTGCTGTATGTATGTTCACAACATATCACTGTACTGCCCTCAAAGTCATATGGATAACGGCTAGCACGTAACCATGCCATGTCAGCGTTG
GATCAACTCTCTCCACCTGTTCTGTTCACCATAACGCTGACTGTATGCGCTTGCGCACTTGATGTTGCACGTGATCTAGGGCTGGGTCGTTTTGTACTATAAGGTGGGAGTTTTGAATACTGTATGACTTTTTGTATCTATAGGACATTGCCACCGAGGTAGTAGACAACCCAGGGATTATTCAAAAGTAGCCATGGCATAGGATTACATGTAGAAATG
TCAGAAAACATACTTGGAATCTAGTCAGTTTCCTTAAACTTCCACAAATGATTCACTGAAAGATGCTTTATTGAAAAAAATACCATCCCATCATGCTTTATTTTTAGAGAACGTGTTGGTAGTAACTTTTCCTTTGAGTTTGAATAGCCGCTGATGCAAAAATGCTGACTTAGCACTTTTTCCACAAGTTTTAATAATCAAATAGGTGGCATCTTTAT
TATTGGATCAGTAACTTCATATCTGTTGTCTATATCGGAAATAGTACATGTAGTGATTAAATAACAATATTTTTATCCACCATGGTTATTTTTTTCTTTATAAGGGTCAAATTTGCAATAAAGTCGTGTTTGGTGTTTGTTACTCAAAATCATGGGTAATTTGTGATTTAATGTATATTATCTCAAAAAGCAGAATGACAACACCAATTTTTCTAGTCA
TAATATAATTCTCACCCTTCCAGCTGAAACATATGTGATTCTAGAAAAAATGACAATATTCTTGGAATTGGGCAGTCCTCAACCTAA

#### FASTA RNA

**RAG1L-A (Fwd) & RAG2L-A (Rev)**

>DN90864
ATCATTTACAAGTTGTTGCACTCTCATGCAATGCAATGCCTTGACCTGATTCAGTATTTCTTGCTTTGGTTCCTGCGGTATGAAGTAGAAAATTAGGCTGGCCACGGTGTCCTGGGAATTAATGTTTTACCACTAAGAGCGGAATGACATCAGCTTTTCAACACAGAGAGTACCTTAAGAGGGTGTGTCGTGTATGTGGTAGAACCCTGAAGACCACAT
CTTACAGCAACTCAGGACATTACAGACAGGGAGAAGTGTGCATCCCATACAGTGAACTTTTAGAAGAATGCTTTAAAGTAAATGTTCGAAATGATGACCCTTTTATACATCCGCCTAGATTTGGTTCATGTTGTCGTAGAGTTCTTTTAAAGTATTCTGTTGCACAGAACCTGGGTAAACATTACTATACCCATATCAGTGTTTGTAACTGGAAACATC
ATTGTAAGTATGAAAACCAAACTTGTCCTCTTTGCAAACGTTTTTCTCAAAATGGTTCTGGTAGACCTAAAAACTTAAAAGGGGTCGCTTTAAATTAAATACAAATGAAAATAATGAAACTGACAAGAGTCCTGTTGCAGACAATTTTGCTCATATACTAAATGACCACTGTTATACCACCCCAGCTTCCTGTAAGTTTGCCCAGTCTGTGCAATCAG
ACCAAAAGACAGATACATATGAATTTTGTACAGAGGAAAATATTCAGACTGTACCCATGACAAGTCGATTTCCAACAGATTTATGTGTAACTGAACCACATCCAGAATTCATTTGTAGAATGTGTTTAGGAGTCCTAGACACACCACTTCTAACACCGTGTACTCACACTTTCTGTGCAGGCTGCATTAAGAACTGGTTAAGTCTGTGTCATTTTTGTC
CTCTCTGTAAACAACCTATGGTGTCATCAGATTTAAAAGAACCATATAGAGGATTTGTAGAAATTCTTAATGATACTAGAATGAAATGTAATTGTCCAGAAGCATTTTCAAGCAAAAAAGGTGTCAAGCTTAGTGATTGGCTTCAGCACACTTTAAATTGTGGAAAGGATTTTGAAATTGATGAATGGTTAGAAAAAGTTTTGTCATTGTGGAACGAAA
TTTCTCTCACTCAAGTTGATTCACATTTAAATAGTACTGAACATTTGCCTGAGACAGATAAGTCTAGTCTGACAACAAGCCATGGCCATCCAGACATTGCTGCAGGTAGCACTGTTGGAAGACCTAAAACTTCACTGTTGTGTCTCCAACAACAAGGACAAAATTTAAGATTGAAAAATATCAAAAAACAAATAGACGTGTTTGCCAGAGAGAAAGGGG
AAGATCTCAGGGTAGTGTATTTTGGCCTTTTGCAAAACTATCTGAGAGCCAACAAGCAAGGTAGTTTAGCTGAACAAGTGAAGTTGCTATATAATGGGGAATCTTGTCAAATGTCACCTCAGGAATGTCTAGCCTTGCGTGTAAATACTTTTACCAGTTGTAATCAGTACAAGAAAATTTACTCGGCTACAAATGAGCTTAGTGGAGTTAAAATCTTCC
AACCTCTCATCAATGTTGTGAAAGCTGAAGAAAGTTTTCTACCAGGGTCTGTAAAGTTCTACTTTAACCCATCTTTGAAAACTGTTTCTCCCATTTGGCAAGGTACAGAGCATGTTGATGCTCTGTCAGGACTCAGTCGCGATATACCACTTCTAGAGTCACCTGATGTTTATGGGGCAAGGTATAGATATGACAAAGCAGTTGCTATGGCTTTGAAAG
ATATTGATAATGAAATCTCAGAAGGCCTCAGTAAATTAGAAGGGGTGGATACAAATCAACTTGTTCTAACAGCCACTGTCAAAGATGGCAGTGATGGTCTTGGCGATGTGAAAGAAGTCAGATCTCAGGGGCTTGTCGTTCCAAACAAGGCCTTACGGTTTTCATTTGTAATTTTAGAAGTCACTGTCAATGTAGATGGCAAGATTATTACTGTATTTG
AAGAAGATGCTCCCAACTCAGAATTATGCACACGCCCTCTCTTGGTAGCTCTGGCAGATGAAAATGACAGAAAAGGGTTATGCATGACACATATACCAATAATGATAGAGAGAGAGGAGATTGAGAAAAGTACCTTGTTTGTAGAAATTGAAGAGGGAAAACAAAGAATCTACAAAGTTGTATTTAAAGGGACTATGTATGATGAAAAATTGCAGAGAA
GTGGTGAGGGAATGTTAGCGGCAGGGTCATCTTGGTTTTGTACACTTTGTGAGAAAGGAAGACTTGATCCCATATCATCAGTCTTGCCAATAACAAGAACACATGAAAACAATATGGCTTTGTATTCAAAGTATGTTGAAAATGCAAATCATATGAAGCAGGCAGATCTAGTGAAAGAAGTCAAAGGCATTGTTAGAAAACCATTTATAATGTCAGAGC
CTTGTGTTGATGCCACGCATGCTGAAATTCACTGGGGAGAGAAAAACTATGATATGTATGTACGTGAAATATCAGGTATAATGTGTTGGGGGAAGCCATCCAATGATGATGATGCAAAAAAAGTGAAATCAACGAAAAGAATGTTGGATACCGAATTACAGCTGCAATGTGGCTTAAAGAGAGAGCATGGCTTTATGGTTGATGGAAATTATGCACGTG
ATTTGGTCAAATCAGAAACAGTTGATGTGGTTTGCTCTTTAATTCAGAGTGAAGCAAGAAAATCTGTTGTTCGTGAATATGTTTCCTACTATCGAGAATTGCGATCAATATACAGGAAACACAATGCTGACTTAGATGATGCCGTGAAATTTAAAACCATAGCTGCAAATTTTTACAAATTGCTACATGAGAAATTTCCTTATTTGCATCCACTCTCTA
ACTATCAACACAAGGTATTAGATCACATTCCACAGTTGATCGAGCAATATGGCTCTGTTGGTAAATTTGCCTCTGAAGGCAATGAAAAAGGCAATAAACTTTTCAGAAGATTTCGTAAATTTAATGCACGTAAGTCTCAAATTCATGAAATGCCAGACATTCTGAAATTTCATTGGCTATATACAATGAAAAGTCTTCAGACTATGGTAAACTGGGAAA
AAACTAGATCACTAGTCTGTAGCAAATGTGGTTGTGCTGGTCACAATCAAAGGACATGCATTACATAATTGTGCCCAAGGTCATGATATGAAATCAAAGATTAGATAATGACCATGCATGTTTTATGTATCTGAATCTGTAAGTTTGAAATTAAGTAGCATCTGTATGTATCAGACTACAGAGTCCAAACTTCAGACACAAAGAGTAGGCTGGCCATAC
ACTCTTATTTACTTAAAAAGTGTTTCTGTTCCATATACTGACTGTACTGTCAAGAGTCAATTCTAGCTTCTATGCTAGTCTTAATATACCTAGATGGTTAAAGTGTATTATATTTTGATGTACATAGATGAATTCCACAAGAAATCATCAAATACAAGAAAGCTGAGTTTGTGTTAAATATTTGTATATGTTTTAATCTTACCTGCCCACACTTTTGAA
GGGACTCAAACAAAATCATTCAATTACTGTGGCCAAAATGTCAAAAGTTTGAGTAACTGGCCATGAATATGAGATGAGAAGCGTAAGGTTTGCAATAGAATTTTGTACATATATTCATGAAAGATTGTAACAAGTTGTAAAATGAACTCTCAGTAAAGGTTTCATTCAATTCATGCACATGCTACATGTACATGCTATAATTTCTTGTGCAGTTGTCAT
GTTCTCAGAAATTACAATCAGAACATGTAAATTTGATGTTACTCTTATTTTCCAGTTGTCGTGAAGAAAATCGTTTGCCAGAGCATTTAATGTGAAAAGCTCTAACACATTGTGTACATCGCACTGTTGAAAAGTAACTCTTATCAGTCACGCTACAATTAATGTTAGCACATGTTGTGAAACATGAAGGGCATTCAAAGTTTATACAGTCCAGCTTTG
TTCTATTTAAGTGATCTAATTCAACACATTTGTCATGGTACCACCTTCCACAGAATACAGTTTGATTGTTCATGATGTAAGTTCGATCACATTTCAGCCAGTTGAACTTGGCTTGTTCAGTAGCTGCTAAATTGGCAAATTTGCAGCCATCCAACATACATTCCCATGTTTCCTCCTCACTATCATCTGATGTATTGTCTATCTTTCCTGCTCTGCTGC
CCTCTTCTGACTCATTCCTGTCTGTGCTAACTTTGACCTCAGCTGCCTCACTTATTCCGTTCTGATCCTCAGTCTCTGAGCCTATACTTTCCTGATCTGTCATGTCTATGCTTATCCTTTCCTGCTCTTCCTCCGCTATGCTTACTCTGTCCTCATCTGTTGACCTTTCCTCCTCATCCCCCAACTGACTGTTATTACTCTGTGTCCTGGTTTCAGAGC
TTGTCCCTTCATTTTGAAAACTGAATTTTATCATTTGCTTTTTTACTGTATCACAAACAGCAATTGAGCTGAAGAATGGAAGACTGTAGTATTTACTACTTAAGGTATCATAGTGTGCTGTTTCTATGAAATCTATGGCATCACAAGGTGACTTGTAAACTGTGTTGTTACTCAATATGTAAACAGTGTTATTACATTTGAATGAGACTGGGTTAGGAA
TACCAATAAATGGTACAGATGTACAGACCATTGAATTCAGGTCAATTACATCAACATCGGGTGGATGTCTTACATTGTTACCTATCACATAAATTCCACCAACAATATACATTTTACTGCTATTACGAAAAACTGAATGGTAGGCTCTTGGAGTGATCGCTTGTAAACTAATACTTGATAATTTCATAGATTGTAAATTAAGAACAGAAAATATGTTTG
AAGCCATGGACACATTGTAAAAGCATGACTGAGTTTTTGTGTATATTGTAAGCCTCCATACAGATAAAGAGAATCTCCAAACTTAATTACACTGTGACCAAAAGCATGAATTTCCTCACCACTATATTGTCCTTGAGACCATGTTAATGAAATTTTCCCCCTTGGATCAACAAAGACAGAAAGCTGCCACAATTTTGTATTGTGCATTTCTGAGTCAG
TCAATCCACCAAGGAGAACATAATGGTTTGGTTCAACTAAACACAAATTTGAGCCATGTACATGTGGAGGGTAATTTGACTGAGAAGAATAATTTTTGATTTCTTGTATTGTATCATCTTTGTCTATTACAAGCCAGCCTGTTACTGTTGTCTTAGACTTGACCCAGACCAAAACTCTGTCACCTAAAGAAATAACTGGATTTGTACATCCAGCAGTTG
CTAGGCAAGTTGCCAGGATGTTTTTGGTGATTTGTATTTCTGAAATAAAAAAGATGATGACAAAAGTTGTTGTATTAGAGAGTG

**RAG2L-A (Fwd)**

>DN47023
TAAAATGACCTGCTCACCATACATGCATGCAGATAAGCTACACTCTCTAATACAACAACTTTTGTCATCATCTTTTTTATTTCAGAAATACAAATCACCAAAAACATCCTGGCAACTTGCCTAGCAACTGCTGGATGTACAAATCCAGTTATTTCTTTAGGTGACAGAGTTTTGGTCTGGGTCAAGTCTAAGACAACAGTAACAGGCTGGCTTGTAATA
GACAAAGATGATACAATACAAGAAATCAAAAATTATTCATCTCAGTCAAATTACCCTCCACATGTACATGGCTCAAATTTGTGTTTAGTTGAACCAAACCATTATGTTCTCCTTGGTGGATTGACTGACTCAGAAATGCACAATACAAAATTGTGGCAGCTTTCTGTCTTTGTTGATCCAAGGGGGAAAATTTCATTAACATGGTCTCAAGGACAATAT
AGTGGTGAGGAAATTCATGCTTTTGGTCACAGTGTAATTAAGTTTGGAGATTCTCTTTATCTGTATGGAGGCTTACAATATACACAAAAACTCAGTCATGCTTTTTACAATGTGTCCATGGCTTCAAACATATTTTCTGTTCTTAATTTACAATCTATGAAATTATCAAGTATTAGTTTACAAGCTATCACTCCAAGAGCCTACCATTCAGTTTTTCGT
AATAGCAGTAAAATGTATATTGTTGGTGGAATTTATGTGATAGGTAACAATGTAAGACATCCACCCGATGTTGATGTAATTGACCTGAATTCAATGGTCTGTACATCTGTACCATTTATTGGTATTCCTAACCCAGTCTCATTCAAATGTAATAACACTGTTTACATATTGAGTAACAACACAGTTTACAAGTCACCTTGTGATGCCATAGATTTCATA
GAAACAGCACACTATGATACCTTAAGTAGTAAATACTACAGTCTTCCATTCTTCAGCTCAATTGCTGTTTGTGATACAGTAAAAAAGCAAATGATAAAATTCAGTTTTCAAAATGAAGGGACAAGCTCTGAAACCAGGACACAGAGTAATAACAGTCAGTTGGGGGATGAGGAGGAAAGGTCAACAGATGAGGACAGAGTAAGCATAGCGGAGGAAGAG
CAGGAAAGGATAAGCATAGACATGACAGATCAGGAAAGTATAGGCTCAGAGACTGAGGATCAGAACGGAATAAGTGAGGCAGCTGAGGTCAAAGTTAGCACAGACAGGAATGAGTCAGAAGAGGGCAGCAGAGCAGGAAAGATAGACAATACATCAGATGATAGTGAGGAGGAAACATGGGAATGTATGTTGGATGGCTGCAAATTTGCCAATTTAGCA
GCTACTGAACAAGCCAAGTTCAACTGGCTGAAATGTGATCGAACTTACATCATGAACAATCAAACTGTACTCTGTGGAAGGTGGTACCATGACAAGTGTGTTGAATTAGATCACTTAAATAGAACAAAGCTGGACTGTATAAACTTTGAATGCCCTTCATGTTTCACAACATGTGCTAACATTAATTGTAGCGTGACTGATAAGAGTTACTTTTCAACA
GTGCGATGTACCAATGTGTTAGAGCTTTTCACATTAAATGCTCTGGCAAACGATTTTCTTCACGACAACTGGAAAATAAGAGTAACATCAAATTTACATGTTCTGATAGTAATTTCTGAGAACATGACAACTGCACAAGAAATTATAGCATGTACATGTAGCATGTGCATGAATTGAATGAAACCTTTACTGAGAGTTCATTTTACAACTTGTTACAAT
CTTTCATGAATATATGTACAAAATTCTATTGCAAACTTTACG

### FASTA protein

**RAG1L-A**

>PflTRAG1L-A_54_DNA (from v2)
MTSAFQHREYLKRVCRVCGRTLKTTSYSSSGHYRQGEVCIPYSELLEECFKVNVRNDDPFVHPPRFGSCCRRVLLKYSVAQNLGKHYYTHISVCNWKHHCKYENQSCPLCKRFSQNGSGRPKKLKRGRFKLNTNEKNETDKSPVADNFAHIPNDHCYTTPASCKFAQSVQSDQKTETYEFCTEENIQTVPMTSRFPTDLCVTEPHPEFICRMCLGVLDT
PLLTPCTHTFCAGCIKNWLSLCHFCPLCKQPMASSDLKEPYRGFVEILNDTRMKCNCPEAVSSKKGVKLSNWLQHTLNCGKDFEIDEWLEKVLSLWNEISLTQVDSHLNSTEHLPETDKSNLTTSHGHPDIAAGSTVGRPKTSLLCLQQQGQNLRLKNIKKQIDVFAREKGEDLRVVYFGLLQNYLRANKQGSLAEQVKLLYNGESCQMSPQECLALRV
NTFTSCNQYKKIYSATNELSGVKIFQPLINVVKAEESFLPGSVKFYFNPSLKTVSPIWQGTEHVDALSGLSRDIPLLESPDVYGARYRYDKAVAMALKDIDNEISEGLSKLEGVDTNQLVLTATVKDGSDGLGDVKEVRSQGLVVPNKALRFSFVILEVTVNVDGKIITVFEEGSPNSELCTRPLLVALADENDRKGLCMTHIPIMIEREEMEKSTLFV
EIEEGKQRIYKVVFKGTMYDEKLQRSGEGMLAAGSSWFCTLCEKGRLDPISSVLPITRTHENNMALYSKYVENANHMKQADLVKEVKGIVRKPFIMSEPCVDATHAEIHWGEKNYDMYVREISGIMCWGKPSNDDDAKKVKSTKRMLDTELQLQCGLKREHGFMVDGNYARDLVKSETVDVVCSLIQSEARKSVVREYVSYYRELRSIYRKHNADLDDA
VKFKTIAANFYKLLHEKFPYLHPLSNYQHKVLDHIPQLIEQYGSVGKFASEGNEKGNKLFRRFRKFNARKSQIHEMPDILKFHWLYTMKSLQTMVNWEKTRSLVCSKCGCAGHNQRT

>PflRAG1L-A_RNA_DN90864
MTSAFQHREYLKRVCRVCGRTLKTTSYSNSGHYRQGEVCIPYSELLEECFKVNVRNDDPFIHPPRFGSCCRRVLLKYSVAQNLGKHYYTHISVCNWKHHCKYENQTCPLCKRFSQNGSGRPKKLKRGRFKLNTNENNETDKSPVADNFAHILNDHCYTTPASCKFAQSVQSDQKTDTYEFCTEENIQTVPMTSRFPTDLCVTEPHPEFICRMCLGVLDT
PLLTPCTHTFCAGCIKNWLSLCHFCPLCKQPMVSSDLKEPYRGFVEILNDTRMKCNCPEAFSSKKGVKLSDWLQHTLNCGKDFEIDEWLEKVLSLWNEISLTQVDSHLNSTEHLPETDKSSLTTSHGHPDIAAGSTVGRPKTSLLCLQQQGQNLRLKNIKKQIDVFAREKGEDLRVVYFGLLQNYLRANKQGSLAEQVKLLYNGESCQMSPQECLALRV
NTFTSCNQYKKIYSATNELSGVKIFQPLINVVKAEESFLPGSVKFYFNPSLKTVSPIWQGTEHVDALSGLSRDIPLLESPDVYGARYRYDKAVAMALKDIDNEISEGLSKLEGVDTNQLVLTATVKDGSDGLGDVKEVRSQGLVVPNKALRFSFVILEVTVNVDGKIITVFEEDAPNSELCTRPLLVALADENDRKGLCMTHIPIMIEREEIEKSTLFV
EIEEGKQRIYKVVFKGTMYDEKLQRSGEGMLAAGSSWFCTLCEKGRLDPISSVLPITRTHENNMALYSKYVENANHMKQADLVKEVKGIVRKPFIMSEPCVDATHAEIHWGEKNYDMYVREISGIMCWGKPSNDDDAKKVKSTKRMLDTELQLQCGLKREHGFMVDGNYARDLVKSETVDVVCSLIQSEARKSVVREYVSYYRELRSIYRKHNADLDDA
VKFKTIAANFYKLLHEKFPYLHPLSNYQHKVLDHIPQLIEQYGSVGKFASEGNEKGNKLFRRFRKFNARKSQIHEMPDILKFHWLYTMKSLQTMVNWEKTRSLVCSKCGCAGHNQRTCIT

**RAG2L-A**

>PflTRAG2L-A_54_DNA(from v2)
MAAAMLDSPEIQITKNILATCLATAGCTNPVISLGDRALVWVKSKTTVTGWLVIDKDDTIQEIKNYSSQSNYPPHVHGSNLCLVEPNHYVLLGGLTDSEMHNTKLWQLSVFVDPRGKISLTWSQGQYSGEEIHAFGHSVIKFGDSLYLYGGLQYTQKLSHAFYNVSMASNIFSVLNLQSMKLSSISLQAITPRAYHSVFLNSSKMYIVGGIYVIGNNVR
HPPDVDVIDLNSMVCTSVPFIGIPNPVSFKCNDTVYILSNNTVYKSPCDAIDFIETAHYDTLSSKYYSLPFFSSIAVCDTVKKQMIKFSFQNEGTSSETRTQSNNSQLGDEEERSTDEDRVSIAEEEQERISIDMTDQESIGSETEDQNGISEAAEVKVSTDRNESEEGSRAGKIDNTSDDSEEETWECMLDGCKFANLAATEQAKFNWLKCDRTYIMN
NQTVLCGRWYHDKCVELDHLNRTKLDCINFECPSCFTTCANINCSVTDKSYFSTVRCTQCVRAFHIKCSGKRFSSRQLENKSNIKFTCSDCNF

>PflRAG2L-A_RNA_DN47023
ATLSNTTTFVIIFFISEIQITKNILATCLATAGCTNPVISLGDRVLVWVKSKTTVTGWLVIDKDDTIQEIKNYSSQSNYPPHVHGSNLCLVEPNHYVLLGGLTDSEMHNTKLWQLSVFVDPRGKISLTWSQGQYSGEEIHAFGHSVIKFGDSLYLYGGLQYTQKLSHAFYNVSMASNIFSVLNLQSMKLSSISLQAITPRAYHSVFRNSSKMYIVGGIY
VIGNNVRHPPDVDVIDLNSMVCTSVPFIGIPNPVSFKCNNTVYILSNNTVYKSPCDAIDFIETAHYDTLSSKYYSLPFFSSIAVCDTVKKQMIKFSFQNEGTSSETRTQSNNSQLGDEEERSTDEDRVSIAEEEQERISIDMTDQESIGSETEDQNGISEAAEVKVSTDRNESEEGSRAGKIDNTSDDSEEETWECMLDGCKFANLAATEQAKFNWLKC
DRTYIMNNQTVLCGRWYHDKCVELDHLNRTKLDCINFECPSCFTTCANINCSVTDKSYFSTVRCTNVLELFTLNALANDFLHDNWKIRVTSNLHVLIVISENMTTAQEIIACTCSMCMN

>PflRAG2L-A_RNA_DN90864_Reverse
LSNTTTFVIIFFISEIQITKNILATCLATAGCTNPVISLGDRVLVWVKSKTTVTGWLVIDKDDTIQEIKNYSSQSNYPPHVHGSNLCLVEPNHYVLLGGLTDSEMHNTKLWQLSVFVDPRGKISLTWSQGQYSGEEIHAFGHSVIKFGDSLYLYGGLQYTQKLSHAFYNVSMASNIFSVLNLQSMKLSSISLQAITPRAYHSVFRNSSKMYIVGGIYVI
GNNVRHPPDVDVIDLNSMVCTSVPFIGIPNPVSFKCNNTVYILSNNTVYKSPCDAIDFIETAHYDTLSSKYYSLPFFSSIAVCDTVKKQMIKFSFQNEGTSSETRTQSNNSQLGDEEERSTDEDRVSIAEEEQERISIDMTDQESIGSETEDQNGISEAAEVKVSTDRNESEEGSRAGKIDNTSDDSEEETWECMLDGCKFANLAATEQAKFNWLKCDR
TYIMNNQTVFCGRWYHDKCVELDHLNRTKLDCINFECPSCFTTCANINCSVTDKSYFSTVRCTQCVRAFHIKCSGKRFSSRQLENKSNIKFTCSDCNF

### Pfl$^{T}$RAGL-A_16

| Genomic loci | RAG1L status | RAG2L status |
|---|---|---|
| Pfl$^{T}$RAGL-A_16_v1 | *pseudogenic (frameshifts)* | potentially functional (2 complete CDSs) |
| Pfl$^{T}$RAGL-A_16_v2 | *pseudogenic (frameshifts)* | *pseudogenic (frameshifts)* |

```
           1                                                                                                                                                                                                      200
PflT_16_v1 GAATGAGACAATCTATTTTTAGTATGCCAAAAAAACAGGACTATTGTTCATATGTAAATTTACGAAAAAATTGTTACTTTTTCTTTTGATTTGAACTCTTGACCTGTTTTCTGTGTCTGCAGCAGGTATTTTAATGATAGTCTTCGCGTGCGTTGCATGCGAATAAAATGCAATGTTGATGGACGTATGCGTATTTATTG
PflT_16_v2 GAATGAGACAATCTATTTTTAGTATGCCAAAAAAACAGGACTATTGTTCATATGTAAATTTACGAAAAAATTGTTACTTTTTCTTTTGATTTGAACTCTTGACCTGTTTTCTGTGTCTGCAGCAGGTATTTTAATGATAGTCTTCGCGTGCGTTGCATGCGAATAAAATGCAATGTTGATGGACGTATGCGTATTTATTG

           201                                                                                                                                                                                                    400
PflT_16_v1 TACGATACTGTGTGCGTGTACGTAGGGCCTAATCCCAACATAAAAATGCTCGGTCTCGGGCGCAGAATTTCCGACGATCAATCTCTCACACGCCCATTTTCTGCATTTGGGGCTTAAAGTGACGAAAATACCCAAATAAGACAACAAAGACACACAAGTTGATATAACATAATAGCAATAACCCCCTTCGCAGTCGGGTA
PflT_16_v2 TACGATACTGTGTGCGTGTACGTAGGGCCTAATCCCAACATAAAAATGCTCGGTCTCGGGCGCAGAATTTCCGACGATCAATCTCTCACACGTCCATTTTCTGCATTTGGGGCTTAAAGTGACGAAAATACCCAAATAAGACAACAAAGACACACAAGTTGATATAACATAATAGCAATAACCCCCTTCGCAGTCGGGTA

           401                                                                                                                                                                                                    600
PflT_16_v1 TACACTCGATTTTGGTACAATTCGCTCCATATACCGGTAGCACTCGCCTATCGGCTCGTGCTATATGTCGCGCATTGTACCAAAATCTCGTGTATACCCTCCTGCGGCGCACAGCGTGCAGCGTAACATGTCAATATAGAAATCGATTAACACTACGATGCGTCGCAGAGAGTATCGACCAATCAGAGAGCAGCATTCAT
PflT_16_v2 TACACTCGATTTTGGTACAATTCGCTCCATATA------GCACTCGCCTATCGGCTCGTGCTATATGTCGCGCATTGTACCAAAATCTCGTGTATACCCTCCTGCGGCGCACAGCGTGCAGCGTAACATGTCAATATAGAAATCGATTAACACTACGATGCGTCGCAGAGAGTATCGACCAATCAGAGAGCAGCATTCAT

           601                                                                                                                                                                                                    800
PflT_16_v1 CAAGCCACGAACTTTCACGTATGGCTTCGAGATTTCGCGTGATCTAGGGTGTTTACTTCAGTAACTCTACAATCTACTAAATACTAAGGGGTCTCAACTGTTTTACACATGCGCAAACGCCATCTTGAAGAGTGGCTGATGATCATGTCGAAAACCGGAGGTCGAAAGACGTAAGTATCGCCGAGATATCGTAATCTGAA
PflT_16_v2 CAAGCCACGAACTTTCACGTATGGCTTCGAGATTTCGCGTGATCTAGGGTGTTTACTTCAGTAACTCTACAATCTACTAAATACTAAGGGGTCCCAACTGTTCTACACATGCGCAAACGCCATCTTGAAGAGTGGCTGATGATCATGTCGAAAACCGGAGGTCGAAAGACGTAAGTATCGCCGAGATATCGTAATCTGAA

           801                                                                                                                                                                                                    1000
PflT_16_v1 TCTCGCGTTGCAATTTTTCACTCGATTATGTGTATCAAAATTATGCGGATACAATTTTTGGATTTTCTGTCAAACTTGGTTTTTCCTGTTGAAAGTTTGGGTTTTTCTCAGTCCGTAACGACATTGTTTACCGAAGGAACATGTGCTGTACGCTTGGACTTTCTTCAAATGGCTTGGACTTTTAGATCTATACCCATCCA
PflT_16_v2 TCTCGCGTTGCAATTTTTCACTCGATTATGTGTATCAAAATTATGCGGATACAATTTTTGGATTTTCTGTCAAACTTGGTTTTTCCTGTTGAAAGTTTGGGTTTTTCTCAGTCCGTAACGACATTGTTTACCGAAGGAACATGTGCTGTACGCTTGGACTTTCTTCAAATGGCTTGGACTTTTAGATCTATACCCATCCA

           1001                                                                                                                                                                                                   1200
PflT_16_v1 ACATGTGTGATTGAGTGTAATGGCCGTTACATTTAAAAATGTTCCTGTGATGGTAGCTAAGTCGGGGAAATGATCGTTTTCTGTTTAATATATTTTTTAAGGTTTGAATCAGCGTGTTACGATATTTAGAATTCTGAGCAGCATCCTAGAGGGTCAATTTGGGTGTACAACGCATGGTTCATGCGGTAGCTAGCTAACCC
PflT_16_v2 ACATGTGTGATTGAGTGTAATGGCCGTTACATTTAAAA-TGTTCCTGTGATGGTAGCTAAGTCGGG-AAATGATCGTTTTCTGTTTAATATATTTTTTAAGGTTTGAATCAGCGTGTTACGATATTTAGAATTCTGAGCAGCATCCTAGAGGGTCAATTTGGGTGTACAACGCATG-TTCATGCGGTAGCTAGCTAACCC

           1201                                                                                                                                                                                                   1400
PflT_16_v1 AGGCCATATGGCACCAGGTACAGGTTGACTAACGAAGTGTTTTGAGGTTAGCATAGTATGAGTACGTCAGATTCATATGTCATCGGCATAGAAGGTCGCTTGTGTAAGATAACACCATATTTTTAGGCTGTCATTCATTTCATTTATTTATTTTGCAGTGGGTTGCAGTCAGCAATGAATCATTTACAAGTTTGTTGCAC
PflT_16_v2 AGGCCATATGGCACCAGGTACAGGTTGACTAACGAAGTGTTTTGAGGTTAGCATAGTATGAGTACGTCAGATTCATATGTCATCGGCATAGAAGGTCGCTTGTGTAAGATAACACCATATTTTTAGGCTGTCATTCATTTCATTTATTTATTTTGCAGTGGGTTGCAGTCAGCAATGAATCATTTACAAGTTTGTTGCAC

           1401                                                                                                                                                                                                   1600
PflT_16_v1 TGTTATGCAATGCAATGCCTTGACCTGATTCAGTATTTCTTGCTTGGGTTCCTGCGGTATGAAGTAGAAAATTAGGCTGGCCACGGTGTCCTGGGAATTAATGTTTTACCACCAAGGTAAGAATAAAATTCTATTGGATTTATTGTTTACACATATATAAGGATGAAAGGTACAGACCATTGTACTGATTGGTACTGTAC
PflT_16_v2 TGTTATGCAATGCAATGCCTTGACCTGATTCAGTATTTCTTGCTTGGGTTCCTGCGGTATGAAGTAGAAAATTAGGCTGGCCACGGTGTCCTGGGAATTAATGTTTTACCACCAAGGTAAGAATAAAATTCTATTGGATTTATTGTTTACACATATATAAGGATGAAAGGTACAGACCATTGTACTGATTGGTACTGTAC

           1601                                                                                                                                                                                                   1800
PflT_16_v1 TTGCCAAATTGATATGCTTAAATAATATTTTAAATATGTTAATTGTTTTCAGAGCGGAATGACGTCAGCTTTTCAACACAGAGAGTACCTTAAGAGGGTGTGTCGTGTATGTGGTAGAACCCTGAAGACCACATCTTACAGCAACTCAGGACATTACAGACAGGGAGAAGTGTGCATCCCGTACAGTGAACTTTTAGAAG
PflT_16_v2 TTGCCAAATTGATATGCTTAAATAATATTTTAAATATGTTAATTGTTTTCAGAGCGGAATGACGTCAGCTTTTCAACACAGAGAGTACCTTAAGAGGGTGTGTCATGTATGTGGTAGAACCCTGAAGACCACATCTTACAGCAACTCAGGACATTACAGACAGGGAGAAGTGTGCATCCCGTACAGTGAACTTTTAGAAG

           1801                                                                                                                                                                                                   2000
PflT_16_v1 AGTGCTTTAAAGTAAATGTTCAAAATGATGACCCTTTTGTACATCCGCCTAGATTTGGTTCATGTTGTCGTAGAGTTCTTTTAAAGTATTCTGTTGCACAGAACCTGGGTAAACATTACTATACCCATATCAGTGTTTGTAACTGGAAACATCATTGTAAGTATGAAAACCAAACTTGTCCTCTTTGCAAACGTTTTTCT
PflT_16_v2 AGTGCTTTAAAGTAAATGTTCAAAATGATGACCCTTTTGTACATCCGCCTAGATTTGGTTCATGTTGCCGTAGAGTTCTTTTAAAGTATTCTGTTGCACAGAACCTGGGTAAACATTACTATACCCATATCAGTGTTTCTAACTGGAAACATCATTGTAAGTATGAAAACCAAACTTGTCCTCTTTGCAAACGTTTTTCT

           2001                                                                                                                                                                                                   2200
PflT_16_v1 CAAAATGGTTCTGGTAGACCTAAAAAACTTAAAAGGGGTCGCTTTAAATTAAATACAAATGAAAATAATGAAACTGACAAGAGTCCTGTTGCAGACAATTTTGCTCATATACTAAATGACCACTGTTATACCACCCCAGCTTCCTGTAAGTTTGCCCAGTCTGTGCAATCAGACCAAAAGACAGATACATATGAATTTTG
PflT_16_v2 CAAAATGGTTCTGGTAGACCTAAAAAACTTAAAAGGGGTCGCTTTAAATTAAATACAAATGAAAATAATGAAACTGACAAGAGTCCTGTTGCAGACAATTTTGCTCATATACTAAATGACCACTGTTATACCACCCCAGCTTCCTGTAAGTTTGCCCAGTCTGTGCAATCAGACCAAAAGACAGATACATATGAATTTTG

           2201                                                                                                                                                                                                   2400
PflT_16_v1 TACAGAGGAAAATATTCAGACTGTACCCATGACAAGTCGATTTCCAACAGATTTATGTGTAACTGAACCACATCCAGAATTCATTTGTAGAATGTGTTTAGGAGTCCTAGACACACCACTTCTAACACCGTGTACTCACACTTTCTGTGCAGGCTGCATTAAGAACTGGTTAAGTCTGTGTCATTTTTGTCCTCTCTGTA
PflT_16_v2 TACAGAGGAAAATATTCAGACTGTACCCATGACAAGTCGATTTCCAACAGATTTATGTGTAACTGAACCACATCCAGAATTCATTTGTAGAATGTGTTTAGGAGTCCTAGACACACCACTTCTAACACCGTGTACTCACACTTTCTGTGCAGGCTGCATTAAGAACTGGTTAAGTCTGTGTCATTTTTGTCCTCTCTGTA

           2401                                                                                                                   F                                                                               2600
PflT_16_v1 AACAACCTATGGTGTCATCAGATTTAAAAGAACCATATAGAGGATTTGTAGAAATTCTTAATGATACTAGAATGAAATGTAATTGTCCAGAAGCAGTTTCAAGCAAAAAAGGTGTCAAGCTCAGTAATTGGCTTCAGCACACTTTAAATTGTGGAAAGGATTTTGAAATTGATGAATGGTTAGAAAAAGTTTTGTCATTG
PflT_16_v2 AACAACCTATGGTGTCATCAGATTTAAAAGAACCATATAGAGGATTTGTAGAAATTCTTAATGATACTAGAATGAAATGTAATTGTCCAGAAGCAGTTTCAAGCAAAAAAGGTGTCAAGCTCAGTAATTGGCT-CAGCACACTTTAAATTGTGGAAAGGATTTTGAAATTGATGAATGGTTAGAAAAAGTTTTGTCATTG

           2601                                                                                                                                                                                                   2800
PflT_16_v1 TGGAACGAAATTTCTGTCATTCAAGTTGATTCATATTTAAATAGTACGGAACATTTGCCTGAGACAGATAAGTCAAGTCTGACAACAAGCCGTGGCCATCAAGACATTGCTGCAGGTAGCACTGTTGGAAGACCTAAACTTCACTGTTGTGTCTCCAACAACAAGGACAAAATTTAAGATTGAAAAATATCAAAAAACAA
PflT_16_v2 TGGAACGAAATTTCTGTCATTCAAGTTGATTCATATTTAAATAGTACTGAACATTTGCCTGAGACAGATAAGTCAAGTCTGACAACAAGCCGTGGCCATCAAAACATTGCTGCAGGTAGCACTGTTGGAAGACCTAAACTTCACTGTTGTGTCTCCAACAACAAGGACAAAATTTAAGATTGAAAAATATCAAAAAACAA

           2801                                                                                                                                                                                                   3000
PflT_16_v1 ATAGACGTGTTTGCCAGAGAGAAAGGGGAAGATCTCAGGGTAGTGTATTTTGGCCTTTTGCAAAACTATCTGAGAGCCAACAAGCAAGGTAGTTTAGCTGAACAAGTGAAGTTGCTATATAATGGGGAATCTTGTCAAATGTCACCTCAGGAATGTCTAGCCTTGCGTGTAAATACTTTTACCAGTTGTAATCAGTACAA
PflT_16_v2 ATAGACGTGTTTGCCAGAGAGAAAGGGGAAGATCTCAGGGTAGTGTATTTTGGCCTTTTGCAAAACTATCTGAGAGCCAACAAGCAAGGTAGTTTAGCTGAACAAGTGAAGTTGCTATATAATGGGGAATCTTGTCAAATGTCACCTCAGGAATGTCTAGCCTTGCGTGTAAATACTTTTACCAGTTGTAATCAGTACAA

           3001                                                                                                                                                                                                   3200
PflT_16_v1 GAAAATTTACTCGGCTACAAATGAGCTTAGTGGAGTTAAAATCTTCCAACCTCTCATCAATGTTGTGAAAGCCGAAGAAAGTTTTCTACCAGGGTCTGTAAAGTTCTACTTTAACCCATCTTTGAAAACTGTTTCTCCATTTGGCAAGGTACAGAGTATGTTGATGCTCTGTCAGGACTCAGTCACGATATACCACTTCT
PflT_16_v2 GAAAATTTACTCGGCTACAAATGAGCTTAGTGGAGTTAAAATCTTCCAACCTCTCATCAATGTTGTGAAAGCCGAAGAAAGTTTTCTACCAGGGTCTGTAAAGTTCTACTTTAACCCATCTTTGAAAACTGTTTCTCCATTTGGCAAGGTACAGAGTATGTTGATGCTCTGTCAGGACTCAGTCACGATATACCACTTCT

           3201                                                                                                                                                                                                   3400
PflT_16_v1 AGAATCACCTGATGTGTATGGGGCAAGGTATAGATATGACAAAGCAGTTGCTATGGCTTTGAAAGATATTGATAATGAAATCTCAGAAGGCCTCAGTAAATTAGAAGGGGTGGATACAAATCAACTTGTTCTAACAGCCACTGTCAAAGATGGCAGTGATGGTCTTGGAGATGTGAAAGAAGTCAGATCTCAGGGGCTTG
PflT_16_v2 AGAATCACCTGATGTGTATGGGGCAAGGTATAGATATGACAAAGCAGTTGCTATGGCTTTGAAAGATATTGATAATGAAATCTCAGAAGGCCTCAGTAAATTAGAAGGGGTGGATACAAATCAACTTGTTCTAACAGCCACTGTCAAAGATGGCAGTGATGGTCTTGGAGATGTGAAAGAAGTCAGATCTCAGGGGCTTG
```

```
           3401                                                                                                                                                                                                   3600
PflT_16_v1 TCGTTCCAAACAAGGCCTTGCGGTTTTCATTTGTAATTTTAGAAGTCACTGTCAATGTAGATGGCAAGATTATTACTGTATTTGAAGAAGATGCTCCCAACTCAGAATTATGCACACGCCCTCTCTTGGTAGCTCTGGCAGATGAAAATGACAGAAAAGGGTTATGCATGACACATATACCAATAATGATAGAGAGAGAG
PflT_16_v2 TCGTTCCAAACAAGGCCTTGCGGTTTTCATTTGTAATTTTAGAAGTCACTGTCAATGTAGATGGCAAGATTATTACTGTATTTGAAGAAGATGCTCCCAACTCAGAATTATGCACACGCCCTCTCTTGGTAGCTCTGGCAGATGAAAATGACAGAAAAGGGTTATGCATGACACATATACCAATAATGATAGAGAGAGAG

           3601                                                                                                                                                                                                   3800
PflT_16_v1 GAGATTGAGAAAAGTACCTTGTTTGTAGAAATTGAAGAGGGAAAACAAAGAATCTACAAAGTTGTATTTAAAGGGACTATGTATGATGAAAAATTGCAGAGAAGTGGTGAGGGAATATTAGCAGCAGGGTCATCTTGGTTTTGTACACTTTGTGAGAAAGGAAGACTTGATCCCATATCATCAGTCTTGCCAATAACAAG
PflT_16_v2 GAGATTGAGAAAAGTACCTTGTTTGTAGAAATTGAAGAGGGAAAACAAAGAATCTACAAAGTTGTATTTAAAGGGACTATGTATGATGAAAATTTGCAGAGAAGTGGTGAGGGAATGTTAGCTGCAGGGTCATCTTGGTTTTGTACACTTTGTGAGAAAGGAAGACTTGATCCCATATCATCAGTCTTGCCAATAACAAG

           3801                                                                                                                                                                                                   4000
PflT_16_v1 AACACATGAAAACAATATGGCTTTGTATTCAAAATATGTTGAAAATGCAAATCATATGAAGCAGGCAGATCTAGTGAAAGAAGTCAAAGGCATTGTTAGAAAACCATTTATAATGTCAGAGCCTTGTGTTGATGCCACGCATGCTGAAATTCACTGGGGAGAGAAAAACTATGATATGTATGTACGTGAAATATCAGGGA
PflT_16_v2 AACACATGGAAACAATATGGCTTTGTATTCAAAATATGTTGAAAATGCAAATCATATGAAGCAGGCAGATCTAGTGAAAGAAGTCAAAGGCATTGTTAGAAAACCATTTATAATGTCAGAGCCTTGTGTTGATGCCACGCATGCTGAAATTCACTGGGGAGAGAAAAACTATGATATGTATGTACGTGAAATATCAGGGA

           4001                                                                                                                                                                                   F               4200
PflT_16_v1 TAATGTGTTGGGGGAAGCCATCCAATGATGATGATGCAAAAAAAGTGAAATCAACGAAAAGAATGTTGGATACCGAATTACAGCTGCAATGTGGCTTAAAGAGAGAGCATGGCTTTATGGTTGATGGAAATTATGCACGTGATTTGGTCAAATCAGAAACAGTTGATGTGGTTTGCTCTTTAATTCAGAGTGAAGCAAGA
PflT_16_v2 TAATGTGTTGGGGGAAGCCATCCAATGATGATGATGCAAAAAAAGTGAAATCAACGAAAAGAATGTTGGATACCGAATTACAGCTGCAATGTGGCTTAAAGAGAGAGCATGGCTTTATGGTTCATGGAAATTATGCACGTGATTTGGTCAAATCAGAAACAGTTGATGTGGTT-GCTCTTTAATTCAGAGTGAAGCAAGA

           4201                                                                                                                                                                                                   4400
PflT_16_v1 AAATCTGTTGTTCGTGAATATGTTTCCTACTATCGAGAATTGCGATCAATATACAGGAAACACAATGCTGACTTAGATGATGCCGTGAAATTTAAAACCATAGCTGCAAATTTTTACAAATTGCTACATGAGAAATTTCCTTATTTGCATCCACTCTCTAACTATCAACACAAGGTATTAGATCACATTCCACAGTTGAT
PflT_16_v2 AAATCTGTTGTTCGTGAATATGTTTCCTACTATCGAGAATTGCGATCAATATACAGGAAACACAATGCTGACTTAGATGATGCCGTGAAATTTAAAACCATAGCTGCAAATTTTTACAAATTGCTACATGAGAACTTTCCTTATTTGCATCCACTCTCTAACTATCAACACAAGGTATTAGATCACATTCCACAGTTGAT

           4401  F                    F                                                                                                                                                                   4600
PflT_16_v1 CGAGCAATATGGCTCTGTTGGTAAATTTGCCTCTGAAGGCAATGAAAAAGGCAATAAACTTTTCAGAAGATTTCGTAAATTTAATGCACGTAAGTCTCAAATTCATGAAATGCCAGACATTCTGAAATTTCATTGGCTATATACAATGAAAAGTCTTCAGACTATGGTAAACTGGGAAAAAACTAGATCACTAGTCTGTA
PflT_16_v2 CGAGCA-TATGGCTCTGTTGGTAGATT-GCCTCTGAAGGCAATGAAAAAGGCAATAAACTTTTCAGAAGATTTCGTAAATTTAATGCACGTAAGTCTCAAATTCATGAAATGCCAGACATTCTGAAATTTCATTGGCTATATACAATGAAAAGTCTTCAGACTATGGTAAACTGGGAAAAAACTAGATCACTAGTCTGTA

           4601                                                                                                                                                                                                   4800
PflT_16_v1 GCAAATGTGGTTGTGCTGGTCACAATCAAAGGACATGCATTACATAATTGTGCCCAAGGTCATGATATGAAATCAAAGATGAGACAATGACCATGCATGTTTTATGTATCTGAATCTGTAAGTTTGAAATTAAGTAGCATCTGTATGTATCAGACTACAGAGTCCAAACTTCAGACACAAAGAGTAGGCTGGCCATACAC
PflT_16_v2 GCAAATGTGGTTGTGCTGGTCACAATCAAAGGACATGCATTACATAATTGTGCCCAAGGTCATGATATGAAATCAAAGATGAGACAATGATCATGCATGTTTTATGTATCTGAATCTGTAAGTTTGAAATTAAGTAGCATCTGTATGTATCAGACTACAGAGTCCAAACTTCAGACACAAAGAGTAGGCTGGCCATACAC

           4801                                                                                                                                                                                                   5000
PflT_16_v1 TCTTATTTACTTAAAAA-GTGTTTCTGTTCCATATACTGACTGTACTGTCAAGAGTCAATTCTAGCTTCTATGCTAGTCTTAATATACCTAGATGGTTAAAGTGTATTATATTTTGATGTACATAGATGAATTCCACAAGAAATCATCAAATACAAGAAAGCTGAGTTTGTGTTAAATATTTGTATATGTTTTAATCTTA
PflT_16_v2 TCTTATTTACTTAAAAAAGTGTTTCTGTTCCATATACTGACTGTACTGTCAAGAGTCAATTCTAGCTTCTATGCTAGTCTTAATATACCTAGATGGTTAAAGTGTATTATATTTTGATGTACATAGATGAATTCCACAAGAAATCATCAAATACAAGAAAGCTGAGTTTGTGTTAAATATTTGTATATGTTTTAATCTTA

           5001                                                                                                                                                                                                   5200
PflT_16_v1 CCTGCCCACACTTTTGAAGGGACTCAAACAAAATCATTCAATTACTTTGGCCAAAATGTCAAAAGTTTGAGTAACTGGCCATGAATATGAGATGAGAAGTGTAAAGTTTGCAATAGAATTTTGTACATATATTCATGAAAGATTGTAACAAGTTGTAAAATGAACTCTCAGTAAAGGTTTCATTCAATTCATGCACATGC
PflT_16_v2 CCTGCCCACACTTTTGAAGGGACTCAAACAAAATCATTCAATTACTTTGGCCAAAATGTCAAAAGTTTGAGTAACTGGCCATGAATATGAGATGAGAAGTGTAAAGTTTGCAATAGAATTTTGTACATATATTCATGAAAGATTGTAACAAGTTGTAAAATGAACTCTCAGTAAAGGTTTCATTCAATTAATGCACATGC

           5201                                                                                                                                                                                                   5400
PflT_16_v1 TACATGTACATGCTATAATTTCTTGTGCAGTTGTCATGTTCTCAGAAATTACAATCAGAACATGTAAATTTGATGTTACTCTTATTTTCCAGTTGTCGTGAAGAAAATCGTTTGCCAGAGCATTTAATGTGAAAAGCTCTAACACATTGTGTACATCGCACTGTTGAAAAGTAACTCTTATCAGTCACGCTACAATTAAT
PflT_16_v2 TACATGTACATGCTATAATTTCTTGTGCAGTTCTCATGTTCTCAGAAATTACAATCAGAACATGTAAATTTGATGTTACTCTTATTTTCCAGTTGTCGTGAAGAAAATCGTTTGCCAGAGCATTTAATGTGAAAAGCTCTAACACATTGTGTACATCGCACTGTTGAAAAGTAACTCTTATCAGTCATGCTACAATTAAT

           5401                                                                                                                                                                                                   5600
PflT_16_v1 GTTAGCACATGTTGTGAAACATGAAGGGCATTCAAAGTTTATACAGTCCAGCTTTGTTCTATTTAAGTGATCTAATTCAACACATTTGTCATGGTACCACCTTCCACAGAATACAGTTTGATTGTTCATGATGTAAGTTCGATCACATTTCAGCCAGTTGAACTTGGCTTGTTCAGTAGCTGCTAAATTGGCAAATTTGC
PflT_16_v2 GTTAGCACATGTTGTGAAACATGAAGGGCATTCAAAGTTTATACAGTCCAGCTTTGTTCTATTTAAGTGATCTAATTCAACACATTTGTCATGGTACCACCTTCCACAGAATACAGTTTGATTGTTCATGATGTAAGTTCGATCACATTTCAGCCAGTTGAACTTGGCTTGTTCAGTAGCTGCTAAATTGGCAAATTTGC

           5601                                                                                                                                                                                                   5800
PflT_16_v1 AGCCATCCAACATACATTCCCATGTTTCCTCCTCACTATCATCTGATGTATTGTCTATCTTTCCTGCTCTGCTGCCCTCTTCTGACTCATTCCTGTCTGTGCTAACTTTGACCTCAGCTGCCTCACTTATTCCGTTCTGATCCTCAGTCTCTGAGCCTATACTTTCCTGATCTGTCATGTCTATGCTTATCCTTTCCTGC
PflT_16_v2 AGCCATCCAACATACATTCCCATGTTTCCTCCTCACTATCATCTGATGTATTGTCTATCTTTCCTGCTCTGCTGCCCTCTTCTGACTCATTCCTGTCTGTGCTAACTTTGACCTCAGCTGCCTCACTTATTCCGTTCTGATCCTCAGTCTCTGAGCCTATACTTTCCTGATCTGTCATGTCTATGCTTATCCTTTCCTGC

           5801                                                                                                                                                                                                   6000
PflT_16_v1 TCTTCCTCCGCTATGCTTACTCTGTCCTCATCTGTTGACCTTTCCTCCTCATCCCCCAACTGACTGTTATTACTCTGTGTCCTGGTTTCAGAGCTTGTCCCTTCATTTTGAAAACTGAATTTTATCATTTGCTTTTTTACTGTATCACAAACAGCAATTGAGCTGAAGAATGGAAGACTGTAGTATTTACTACTTAAGGT
PflT_16_v2 TCTTCCTCCGCTATGCTTACTCTGTCCTCATCTGTTGACCTTTCCTCCTCATCCCCCAACTGACTGTTATTACTCTGTGTCCTGGTTTCAGAGCTTGTCCCTTCATTTTGAAAACTGAATTTTATCATTTGCTTTTTTACTGTATCACAAACAGCAATTGAGCTGAAGAATGGAAGACTGTAGTATTTACTACTTAAGGT

           6001                                                                                                                                                                                                   6200
PflT_16_v1 ATCATAGTGTGCTGTTTCTATGAAATCTATGGCATCACAAGGTGACTTGTAAACTGTGTTGTTACTCAATATGTAAACAGTGTTATTACATTTGAATGAGACTGGGTTAGGAATACCAATAAATGGTACAGATGTACAGACCATTGAATTCAGGTCAATAACATCAACATCGGGTGGATGTCTTACATTGTTACCTATCA
PflT_16_v2 ATCATAGTGTGCTGTTTCTATGAAATCTATGGCATCACAAGGTGACTTGTAAACTGTGTTGTTACTCAATATGTAAACAGTGTTATTACATTTGAATGAGACTGGGTTAGGAATACCAATAAATGGTACAGATGTACAGACCATTGAATTCAGGTCAATAACATCAACATCGGGTGGATGTCTTACATTGTTACCTATCA

           6201                                                                                                                                                                                 F                 6400
PflT_16_v1 CATAAATTCCACCAACAATATACATTTTACTGCTATTTAGAAAAACTGAATGGTAGGCTCTTGGAGTGATAGCTTGTAAACTAATACTTGATAATTTCATAGATTGTAAATTAAGAACAGAAAATATGTTTGAAGCCATGGACACATTGTAAAAAGCATGACTGAGTTTTT-GTGTATATTGTAAGCCTCCATACAGATA
PflT_16_v2 CATAAATTCCACCAACAATATACATTTTACTGCTATTTAGAAAAACTGAATGGTAGGCTCTTGGAGTGATAGCTTGTAAACTAATACTTGATAATTTCATAGATTGTAAATTAAGAACAGAAAATATGTTTGAAGCCATGGACACATTGTAAAAAGCATGACTGAGTTTTTTGTGTATATTGTAAGCCTCCATACAGATA

           6401                                 F                                                                                                                                                         6600
PflT_16_v1 AAGAGAATCTCCAAACTTAATTACACTGTGACCAAAAGCATGAATTTCCTCACCACTATATTGTCCTTGAGACCATGTTAATGAAATTTTCCCCCTTGGATCAACAAAGACAGAAAGCTGCCACAATTTTGTATTGTGCATTTCTGAGTCAGTCAATCCACCAAGGAGAACATAATGGTTTGGTTCAACTAAACACAAAT
PflT_16_v2 AAGAGAATCTCCAAACTTAATTACACTGTGACCAAA-GCATGAATTTCCTCACCACTATATTGTCCTTGAGACCATGTTAATGAAATTTTCCCCCTTGGATCAACAAAGACAGAAAGCTGCCACAATTTTGTATTGTGCATTTCTGAGTCAGTCAATCCACCAAGGAGAACATAATGGTTTGGTTCAACTAAACACAAAT

           6601                                                                                                                                                                                                   6800
PflT_16_v1 TTGAGCCATGTACATGTGGAGGGTAATTTGACTGAGATGAATAATTTTTGATTTCTTGTATTGTATCATCTTTGTCTATTACAAGCCAGCCTGTTACTGTTGTCTTAGACTTAACCCAGACCAAAACTCTGTCACCTAAAGAAATAACTGGATTTGTACATCCAGCAGTTGCTAGGCAAGTTGCCAGGATGTTTTTGGTG
PflT_16_v2 TTGAGCCATGTACATGTGGAGGGTAATTTGACTGAGATGAATAATTTTTGATTTCTTGTATTGTATCATCTTTGTCTATTACAAGCCAGCCTGTTACTGTTGTCTTAGACTTAACCCAGACCAAAACTCTGTCACCTAAAGAAATAACTGGATTTGTACATCCAGCAGTTGCTAGGCAAGTTGCCAGGATGTTTTTGGTG

           6801                                                                                                                                                                                                   7000
PflT_16_v1 ATTTGTATTTCTGAAATAAAAAAGATGATGACAAAAGTTGTTGTATTAGAGAGTGAAGCTTATCTGCATGCATGTATGGTGAGCAGGTCATTTTATTTTGTCAGTGTTGAAATATGGCACTGAAATATAAATTTGTTAATTTCATATTTTAATATGGATATCTTTGTAGTTCAAAAAATGCTCAAGACAGAAAGTGAGAT
PflT_16_v2 ATTTGTATTTCTGAAATGAAAAAGATGATGACAAAAGTTGTTGTATTAGAGAGTGAAGCTTATCTGCATGCATGTATGGTGAGCAGGTCATTTTATTTTGTCAGTGTTGAAATATGGCACTGAAATATAAATTTGTTAATTTCATATTTTAGTATGGATATCTTTGTAGTTCAAAAAATGCTCAAGACAGAAAGTGAGAT

           7001                                                                                                                                                                                                   7200
PflT_16_v1 GTCTTTAGTCGTGACATTTATTAATATTGTTCACTGTACATTTCTCTACCCTTTGCAACCATACAATTGTTGAACACCTGTCCTGTATATTCAAACAAACAACTTGAAGGCAAATCTGATATCTTTGGGCAGCATGTTCTCTTGTCTTTTGCATTTGAGTAACCTCCTTATATAAATATGATCGAAAGAGATTGTCAACA
PflT_16_v2 GTCTTTAGTCGTGACATTTATTAATATTGTTCACTGTACATTTCTCTACCCTTTGCAACCATACAATTGTTGAACACCTATCCTGTATATTCAAACAAACAACTTGAAGGCAAATCTGATATCTTTGGGCAGCATGTTCTCTTGTCTTTTGCATTTGAGTAACCTCCTTATATAAATATGATCGAAAGAGATTGTCAACA
```

```
         7201                                                                                                                                                                                              7400
PflT_16_v1  TGGAGCAGCAAATCCGTTTCTTTAACCTAGACTGTAAATCAATACAAAATGGGTGCACATGTAGCTAGGGGAAAAGGTAATGCACAGATAGAGCCCCATTGATGACTGCTGGAAATGTCATTTGGATGTCTTGCTGGTGTACATGCAACTGAAAAAATCATTTTAGCCTGTGGGCCGTACAATAGCTGCACTTCTACTTC
PflT_16_v2  TGGAGCAGCAAATCCGTTTCTTTAACCTAGACTGTAAATCAATACAAAATGGGTGCACATGTAGCTAGGGGAAAAGGTAATGCACAGATAGAGCCCCATTGATGACTGCTGGAAATGTCATTTGGATGTCTTGCTGGTGTACATGCAACTGAAAAAATCATTTTAGCCTGTGGGCCGTACAATAGCTGCACTTCTACTTC

         7401                                                                                                                                                                                              7600
PflT_16_v1  GCTCACTCTAATTTGAATCTTCGTATATAGTGTACATGGGGATCGACTTTCATAGTATGCTTGTTGGATCGAGGGACCGTGGCTTGTGTTACAGTTGAGAACGATCACATCGAGAATTTTCTTTTTCGAATGGGCGATTACAAGTGCAAATCTTTGGAAGTTAGCTACCATATAAGAACTTTGTTGTTTTAACTTCAAGA
PflT_16_v2  GCTCACTCTAATTTGAATCTTCGTATATAGTGTACATGGGGATCGACTTTCATAGTATGCTTGTTGGATCGAGGGACCGTGGCTTGTGTTACAGTTGAGAACGATCACATCGAGAATTTTCTTTTTCGAATGGGCGATTACAAGTGCAAATCTTTGGAAGTTAGCTACCATATAAGAACTTTGTTGTTTTAACTTCAAGA

         7601                                                                                                                                                                                              7800
PflT_16_v1  ATAATCAATCTAAAACACGAACAGAATGTTCTGTGACTTTTGAAGCTACATTGAACTGACGACCTACCGGGACTATCGAGCATGGCGGCTGCCATCTTGGATATTGTTTACGGCTCACTGAGCATGCTCAGATCAAAAATATTTCCCACAACCCATTGCGGACTTTCTACATTGTAAACATAGGAGATGTTACGGTGTGC
PflT_16_v2  ATAATCAATCTAAAACACGAACAGAATGTTCTGTGACTTTTGAAGCTACATTGAACTGACAACCTACCGGGACTATCGAGCATGGCGGCTGCCATCTTGGATATTGTTTACGGCTCACTGAGCATGCTCAGATCAAAAATATTTCCCACAACCCATTGCGGACTTTCTACATTGTAAACATAGGAGATGTTACGGTGTGC

         7801                                                                                                                                                      7941
PflT_16_v1  GGCGTGGGTTATTGCTTAATTTTTTGCGCTGCATTTTTATCGGCTAACTTCATACGTAATCGATGATGGAAATAGAATTATGCCGTCAATTCCCCTCTAAAATACAGTGAAAATATGATAATAAGGTAAGTCTTTGGTCAA
PflT_16_v2  GGCGTGGGTTATTGCTTAATTTTTTGCGCTGCATTTTTATCGGCTAACTTCATACGTAATCGATGATGGAAATAGAATTATGCCGTCAATTCCCCTCTAAAATACAGTGAAAATATGATAATAAGGTAAGTCTTTGGTCAA
```

#### FASTA DNA

```
>PflT_16_v1
AATTATTGCATTGAAGCGACAGCGCACCGTGCACTTTTCTTGATATTGCGGGAATCGAGACGTGATATATCCTACCGCTCTTTAAAAATGAAGCTAGTATTCGTTCACACACTAATAAATTGACACTTCCTCACAGTAACAGTATGTCAGATATTCTTTACCAAAGTTTGGGCTGCAACAGACCAGGGTGTCCTAACGATGTAGACCAAGAAGTTCAAA
ATAAAAATGGGACGACTCCTGGCGACTGCGACAAAATTTGACATCAAATGCTACGAACAATATCAGCGTCCAGCTCTCGCTGCATCGGGCAACACACACTGTACATCATTACAATTGCAGTCATCGTACGTCTGGATTATTTCAAACTGCATGTTTCTAGTATGTCCGAATTATCCATCAAAAATACATCCTGTAACTTCACTGTACCACCTCTAAAT
GTCGCGACGGTCGACGTACACCGTATACGCGCGTAAGCGGTACAGAATGAGACAATCTATTTTTAGTATGCCAAAAAAACAGGACTATTGTTCATATGTAAATTTACGAAAAAATTGTTACTTTTTCTTTTGATTTGAACTCTTGACCTGTTTTCTGTGTCTGCAGCAGGTATTTTAATGATAGTCTTCGCGTGCGTTGCATGCGAATAAAATGCAATG
TTGATGGACGTATGCGTATTTATTGTACGATACTGTGTGCGTGTACGTAGGGCCTAATCCCAACATAAAAATGCTCGGTCTCGGGCGCAGAATTTCCGACGATCAATCTCTCACACGCCCATTTTCTGCATTTGGGGCTTAAAGTGACGAAAATACCCAAATAAGACAACAAAGACACACAAGTTGATATAACATAATAGCAATAACCCCCTTCGCAGT
CGGGTATACACTCGATTTTGGTACAATTCGCTCCATATACCGGTAGCACTCGCCTATCGGCTCGTGCTATATGTCGCGCATTGTACCAAAATCTCGTGTATACCCTCCTGCGGCGCACAGCGTGCAGCGTAACATGTCAATATAGAAATCGATTAACACTACGATGCGTCGCAGAGAGTATCGACCAATCAGAGAGCAGCATTCATCAAGCCACGAACT
TTCACGTATGGCTTCGAGATTTCGCGTGATCTAGGGTGTTTACTTCAGTAACTCTACAATCTACTAAATACTAAGGGGTCTCAACTGTTTTACACATGCGCAAACGCCATCTTGAAGAGTGGCTGATGATCATGTCGAAAACCGGAGGTCGAAAGACGTAAGTATCGCCGAGATATCGTAATCTGAATCTCGCGTTGCAATTTTTCACTCGATTATGTG
TATCAAAATTATGCGGATACAATTTTTGGATTTTCTGTCAAACTTGGTTTTTCCTGTTGAAAGTTTGGGTTTTTCTCAGTCCGTAACGACATTGTTTACCGAAGGAACATGTGCTGTACGCTTGGACTTTCTTCAAATGGCTTGGACTTTTAGATCTATACCCATCCAACATGTGTGATTGAGTGTAATGGCCGTTACATTTAAAATGTTCCTGTGAT
GGTAGCTAAGTCGGGAAATGATCGTTTTCTGTTTAATATATTTTTTAAGGTTTGAATCAGCGTGTTACGATATTTAGAATTCTGAGCAGCATCCTAGAGGGTCAATTTGGGTGTACAACGCATGGTTCATGCGGTAGCTAGCTAACCCAGGCCATATGGCACCAGGTACAGGTTGACTAACGAAGTGTTTTGAGGTTAGCATAGTATGAGTACGTCAG
ATTCATATGTCATCGGCATAGAAGGTCGCTTGTGTAAGATAACACCATATTTTTAGGCTGTCATTCATTTCATTTATTTATTTTGCAGTGGGTTGCAGTCAGCAATGAATCATTTACAAGTTTGTTGCACTGTTATGCAATGCAATGCCTTGACCTGATTCAGTATTTCTTGCTTGGGTTCCTGCGGTATGAAGTAGAAAATTAGGCTGGCCACGGTGT
CCTGGGAATTAATGTTTTACCACCAAGGTAAGAATAAAATTCTATTGGATTTATTGTTTACACATATATAAGGATGAAAGGTACAGACCATTGTACTGATTGGTACTGTACTTGCCAAATTGATATGCTTAAATAATATTTTAAATATGTTAATTGTTTTCAGAGCGGAATGACGTCAGCTTTTCAACACAGAGAGTACCTTAAGAGGGTGTGTCGTGT
ATGTGGTAGAACCCTGAAGACCACATCTTACAGCAACTCAGGACATTACAGACAGGGAGAAGTGTGCATCCCGTACAGTGAACTTTTAGAAGAGTGCTTTAAAGTAAATGTTCAAAATGATGACCCTTTTGTACATCCGCCTAGATTTGGTTCATGTTGTCGTAGAGTTCTTTTAAAGTATTCTGTTGCACAGAACCTGGGTAAACATTACTATACCCA
TATCAGTGTTTGTAACTGGAAACATCATTGTAAGTATGAAAACCAAACTTGTCCTCTTTGCAAACGTTTTTCTCAAAATGGTTCTGGTAGACCTAAAAAACTTAAAAGGGGTCGCTTTAAATTAAATACAAATGAAAATAATGAAACTGACAAGAGTCCTGTTGCAGACAATTTTGCTCATATACTAAATGACCACTGTTATACCACCCCAGCTTCCTG
TAAGTTTGCCCAGTCTGTGCAATCAGACCAAAAGACAGATACATATGAATTTTGTACAGAGGAAAATATTCAGACTGTACCCATGACAAGTCGATTTCCAACAGATTTATGTGTAACTGAACCACATCCAGAATTCATTTGTAGAATGTGTTTAGGAGTCCTAGACACACCACTTCTAACACCGTGTACTCACACTTTCTGTGCAGGCTGCATTAAGAA
CTGGTTAAGTCTGTGTCATTTTTGTCCTCTCTGTAAACAACCTATGGTGTCATCAGATTTAAAAGAACCATATAGAGGATTTGTAGAAATTCTTAATGATACTAGAATGAAATGTAATTGTCCAGAAGCAGTTTCAAGCAAAAAAGGTGTCAAGCTCAGTAATTGGCTTCAGCACACTTTAAATTGTGGAAAGGATTTTGAAATTGATGAATGGTTAGA
AAAAGTTTTGTCATTGTGGAACGAAATTTCTGTCATTCAAGTTGATTCATATTTAAATAGTACGGAACATTTGCCTGAGACAGATAAGTCAAGTCTGACAACAAGCCGTGGCCATCAAGACATTGCTGCAGGTAGCACTGTTGGAAGACCTAAACTTCACTGTTGTGTCTCCAACAACAAGGACAAAATTAAGATTGAAAAATATCAAAAAACAAATA
GACGTGTTTGCCAGAGAGAAAGGGGAAGATCTCAGGGTAGTGTATTTTGGCCTTTTGCAAAACTATCTGAGAGCCAACAAGCAAGGTAGTTTAGCTGAACAAGTGAAGTTGCTATATAATGGGGAATCTTGTCAAATGTCACCTCAGGAATGTCTAGCCTTGCGTGTAAATACTTTTACCAGTTGTAATCAGTACAAGAAAATTTACTCGGCTACAAAT
GAGCTTAGTGGAGTTAAAATCTTCCAACCTCTCATCAATGTTGTGAAAGCCGAAGAAAGTTTTCTACCAGGGTCTGTAAAGTTCTACTTTAACCCATCTTTGAAAACTGTTTCTCCATTTGGCAAGGTACAGAGTATGTTGATGCTCTGTCAGGACTCAGTCACGATATACCACTTCTAGAATCACCTGATGTGTATGGGGCAAGGTATAGATATGACA
AAGCAGTTGCTATGGCTTTGAAAGATATTGATAATGAAATCTCAGAAGGCCTCAGTAAATTAGAAGGGGTGGATACAAATCAACTTGTTCTAACAGCCACTGTCAAAGATGGCAGTGATGGTCTTGGAGATGTGAAAGAAGTCAGATCTCAGGGGCTTGTCGTTCCAAACAAGGCCTTGCGGTTTTCATTTGTAATTTAGAAGTCACTGTCAATGTAG
ATGGCAAGATTATTACTGTATTTGAAGAAGATGCTCCCAACTCAGAATTATGCACACGCCCTCTCTTGGTAGCTCTGGCAGATGAAAATGACAGAAAAGGGTTATGCATGACACATATACCAATAATGATAGAGAGAGAGGAGATTGAGAAAAGTACCTTGTTTGTAGAAATTGAAGAGGGAAAACAAAGAATCTACAAAGTTGTATTTAAAGGACTA
TGTATGATGAAAAATTGCAGAGAAGTGGTGAGGGAATATTAGCAGCAGGGTCATCTTGGTTTTGTACACTTTGTGAGAAAGGAAGACTTGATCCCATATCATCAGTCTTGCCAATAACAAGAACACATGAAAACAATATGGCTTTGTATTCAAAATATGTTGAAAATGCAAATCATATGAAGCAGGCAGATCTAGTGAAAGAAGTCAAAGGCATTGTTA
GAAAACCATTTATAATGTCAGAGCCTTGTGTTGATGCCACGCATGCTGAAATTCACTGGGGAGAGAAAAACTATGATATGTATGTACGTGAAATATCAGGGATAATGTGTTGGGGGAAGCCATCCAATGATGATGATGCAAAAAAAGTGAAATCAACGAAAAGAATGTTGGATACCGAATTACAGCTGCAATGTGGCTTAAAGAGAGAGCATGGCTTTA
TGGTTGATGGAAATTATGCACGTGATTTGGTCAAATCAGAAACAGTTGATGTGGTTTGCTCTTTAATTCAGAGTGAAGCAAGAAAATCTGTTGTTCGTGAATATGTTTCCTACTATCGAGAATTGCGATCAATATACAGGAAACACAATGCTGACTTAGATGATGCCGTGAAATTTAAAACCATAGCTGCAAATTTTACAAATGCTACATGAGAAAT
TTCCTTATTTGCATCCACTCTCTAACTATCAACACAAGGTATTAGATCACATTCCACAGTTGATCGAGCAATATGGCTCTGTTGGTAAATTTGCCTCTGAAGGCAATGAAAAAGGCAATAAACTTTTCAGAAGATTTCTGTAAATTTAATGCACGTAAGTCTCAAATTCATGAAATGCCAGACATTCTGAAATTTCATTGGCTATATACAATGAAAAGTC
TTCAGACTATGGTAAACTGGGAAAAAACTAGATCACTAGTCTGTAGCAAATGTGGTTGTGCTGGTCACAATCAAAGGACATGCATTACATAATTGTGCCCAAGGTCATGATATGAAATCAAAGATGAGACAATGACCATGCATGTTTTATGTATCTGAATCTGTAAGTTTGAAATTAAGTAGCATCTGTATGTATCAGACTACAGAGTCCAAACTTCAG
ACACAAAGAGTAGGCTGGCCATACACTCTTATTTACTTAAAAAGTGTTTCTGTTCCATATACTGACTGTACTGTCAAGAGTCAATTCTAGCTTCTATGCTAGTCTTAATATACCTAGATGGTTAAAGTGTATTATATTTTGATGTACATAGATGAATTCCACAAGAAATCATCAAATACAAGAAAGCTGAGTTTGTGTTAAATATTTGTATATGTTTTA
ATCTTACCTGCCCACACTTTTGAAGGGACTCAAACAAAATCATTCAATTACTTTGGCCAAAATGTCAAAAGTTTGAGTAACTGGCCATGAATATGAGATGAGAAGTGTAAAGTTTGCAATAGAATTTTGTACATATATTCATGAAAGATTGTAACAAGTTGTAAAATGAACTCTCAGTAAAGGTTTCATTCAATTCATGCACATGCTACATGTACATGC
TATAATTTCTTGTGCAGTTGTCATGTTCTCAGAAATTACAATCAGAACATGTAAATTTGATGTTACTCTTATTTTCCAGTTGTCGTGAAGAAAATCGTTTGCCAGAGCATTTAATGTGAAAAGCTCTAACACATTGTGTACATCGCACTGTTGAAAAGTAACTCTTATCAGTCACGCTACAATTAATGTTAGCACATGTTGAAACATGAAGGGCATT
CAAAGTTTATACAGTCCAGCTTTGTTCTATTTAAGTGATCTAATTCAACACATTTGTCATGGTACCACCTTCCACAGAATACAGTTTGATTGTTCATGATGTAAGTTCGATCACATTTCAGCCAGTTGAACTTGGCTTGTTCAGTAGCTGCTAAATTGGCAAATTTGCAGCCATCCAACATACATTCCCATGTTTCCTCCTCACTATCATCTGATGTAT
TGTCTATCTTTCCTGCTCTGCTGCCCTCTTCTGACTCATTCCTGTCTGTGCTAACTTTGACCTCAGCTGCCTCACTTATTCCGTTCTGATCCTCAGTCTCTGAGCCTATACTTTCCTGATCTGTCATGTCTATGCTTATCCTTTCCTGCTCTTCCTCCGCTATGCTTACTCTGTCCTCATCTGTTGACCTTTCCTCCTCATCCCCCAACTGACTGTTAT
TACTCTGTGTCCTGGTTTCAGAGCTTGTCCCTTCATTTTGAAAACTGAATTTTATCATTTGCTTTTTTACTGTATCACAAACAGCAATTGAGCTGAAGAATGGAAGACTGTAGTATTTACTACTTAAGGTATCATAGTGTGCTGTTTCTATGAAATCTATGGCATCACAAGGTGACTTGTAAACTGTGTTGTTACTCAATATGTAAACAGTGTTATTAC
ATTTGAATGAGACTGGGTTAGGAATACCAATAAATGGTACAGATGTACAGACCATTGAATTCAGGTCAATAACATCAACATCGGGTGGATGTCTTACATTGTTACCTATCACATAAATTCCACCAACAATATACATTTTACTGCTATTTAGAAAAACTGAATGGTAGGCTCTTGGAGTGATAGCTTGTAAACTAATACTTGATAATTTCATAGATTGTA
AATTAAGAACAGAAAATATGTTTGAAGCCATGGACACATTGTAAAAAGCATGACTGAGTTTTTGTGTATATTGTAAGCCTCCATACAGATAAAGAGAATCTCCAAACTTAATTACACTGTGACCAAAAGCATGAATTTCCTCACCACTATATTGTCCTTGAGACCATGTTAATGAAATTTTCCCCCTTGGATCAACAAAGACAGAAAGCTGCCACAATT
TTGTATTGTGCATTTCTGAGTCAGTCAATCCACCAAGGAGAACATAATGGTTTGGTTCAACTAAACACAAATTTGAGCCATGTACATGTGGAGGGTAATTTGACTGAGATGAATAATTTTTGATTTCTTGTATTGTATCATCTTTGTCTATTACAAGCCAGCCTGTTACTGTTGTCTTAGACTTAACCCAGACCAAACTCTGTCACCTAAAGAAATAA
CTGGATTTGTACATCCAGCAGTTGCTAGGCAAGTTGCCAGGATGTTTTTGGTGATTTGTATTTCTGAAATAAAAAAGATGATGACAAAAGTTGTTGTATTAGAGAGTGAAGCTTATCTGCATGCATGTATGGTGAGCAGGTCATTTTATTTTGTCAGTGTTGAAATATGGCACTGAAATATAAATTTGTTAATTTCATATTTTAATATGGATATCTTTG
TAGTTCAAAAAATGCTCAAGACAGAAAGTGAGATGTCTTTAGTCGTGACATTTATTAATATTGTTCACTGTACATTTCTCTACCCTTTGCAACCATACAATTGTTGAACACCTGTCCTGTATATTCAAACAAACAACTTGAAGGCAAATCTGATATCTTTGGGCAGCATGTTCTCTTGTCTTTTGCATTTGAGTAACCTCCTTATATAAATATGATCGA
AAGAGATTGTCAACATGGAGCAGCAAATCCGTTTCTTTAACCTAGACTGTAAATCAATACAAAATGGGTGCACATGTAGCTAGGGGAAAAGGTAATGCACAGATAGAGCCCCATTGATGACTGCTGGAAATGTCATTTGGATGTCTTGCTGGTGTACATGCAACTGAAAAAATCATTTTAGCCTGTGGGCCGTACAATAGCTGCACTTCTACTTCGCTC
ACTCTAATTTGAATCTTCGTATATAGTGTACATGGGGATCGACTTTCATAGTATGCTTGTTGGATCGAGGGACCGTGGCTTGTGTTACAGTTGAGAACGATCACATCGAGAATTTTCTTTTTCGAATGGGCGATTACAAGTGCAAATCTTTGGAAGTTAGCTACCATATAAGAACTTTGTTGTTTTAACTTCAAGAATAATCAATCTAAAACACGAACA
GAATGTTCTGTGACTTTTGAAGCTACATTGAACTGACGACCTACCGGGACTATCGAGCATGGCGGCTGCCATCTTGGATATTGTTTACGGCTCACTGAGCATGCTCAGATCAAAAATATTTCCCACAACCCATTGCGGACTTTCTACATTGTAAACATAGGAGATGTTACGGTGTGCGGCGTGGGTTATTGCTTAATTTTTTGCGCTGCATTTTTATCG
GCTAACTTCATACGTAATCGATGATGGAAATAGAATTATGCCGTCAATTCCCCTCTAAAATACAGTGAAAATATGATAATAAGGTAAGTCTTTGGTCAAGAAGAACAATTTTAAGATGCATTAAGTAAGTTTTTGAAGAGGAACTAATGACAAACCAAAAACATCAAAAAAATTCGAACATGCATATTGCTATGATATCCCTACAAACATGCATACAAA
ATTTCAAATAATATGTGCTTTATAGAAACACTTTTGACTATATAAACAATTATGAATATCCAATATTTCTGAAATAAAGAAGCGTTGATGTGTAAGTGAAAATTTGAATGGCCATAAAATCTAGCTCCCCGTCCACAGACTCTCTTTTTCACTTTCCTTGAGAGATTTTTCTATATTACAAATGATGTTGACAAAATCATTCACACGTTCGACATGTGT
TTGATCTCGAATTTTCGAAAATACTAATCGGTTTACTGATTGGTCGGAATGTCTTGCATGCTAATAATGCACTTGGCTCTGTTATTTTCCAGACTGTATTTGTTAGACAACGAACTGGAGTCGATAATAAGTCTCGACTTGAATTCTGAAAATGCTGAACCCTATACTTTCAAAAGAATGAATGATTTACAATGGTTGTTTTCAATCGATATAGACCCG
GTAGGTATGACACAGAAATTTAAAATAAATTTTAAATTTTATTGCAGAAAAACGTTGGCCAAAGTTGTGTCGACGAATCTGTCGATAGTCTACAGATTTCTGCATCGAGGATTGAGAATGTTATGTATGAGACTGACTTTGAACACTTATTAAAATGTCTGATATCAATCACTCAATATACAACTAGACGTTCACCCACATATCTTTCTTTCATGCTTT
TGTGTGCAATGTATTTTCTTCGGTACATTGGTTCTAAAAGCTTTATGTCTGTGCTAGTAAATCTTTAACGTAGTAATGT

>PflT_16_v2
GAAATAATATTATATTATATCATGCTACCATGCGCCATTCTGATTGGACGAGAGCTCATTCCACCGATGCTAAAATACTGTTTTAGCACTGATAGCAGTGCTAAAACGGGCCCCTAAACCAGCATTTTCGAAAATTTCCCACTCGCTGCATTTGAACGTACCATAGACCCCCGCGAAAAACCGAAACGGTCCGCTTTGTTTCAAAGAAAGAATACCGAA
TTTTGATACACAGATAAAGTGTGGGTCTGTAACTCACCTCTTGGTGAAAATAAGTTGGGATCGATGATAAAATACATCTCTGAAGCAGCACATTTTAGTGTGATGTAGACGAATTATTGCATTGAAGCGACAGCGCACCGTGCACTTTTCTTGATATTGCGGGAATCGAGACGTGATATATCCTACCGCTCTTTAAAAATGAAGCTAGTATTCGTTCAC
ACACTAATAAATTGACACTTCCTCACAGTAACAGTATGTCAGATATTCTTTACCAAAGTTTGGGCTGCAACAGACCAGGGTGTCCTAACGATGTAGACCAAGAAGTTCAAAATAAAAATGGGACGACTCCTGGCGACTGCGACAAAATTTGACATCAAATGCTACGAACAATATCAGCGTCCAGCTCTCGCTGCATCGGGCAACACACACTGTACATCA
TTACAATTGCAGTCATCGTACGTCTGGATTATTTCAAACTGCATGTTTCTAGTATGTCCGAATTATCCATCAAAAATACATCCTGTAACTTCACTGTACCACCTCTAAATGTCGCGACGGTCGACGTACACCGTATACGCGCGTAAGCGGTACAGAATGAGACAATCTATTTTTAGTATGCCAAAAAAACAGGACTATTGTTCATATGTAAATTTACG
AAAAAATTGTTACTTTTTCTTTTGATTTGAACTCTTGACCTGTTTTCTGTGTCTGCAGCAGGTATTTTAATGATAGTCTTCGCGTGCGTTGCATGCGAATAAAATGCAATGTTGATGGACGTATGCGTATTTATTGTACGATACTGTGTGCGTGTACGTAGGGCCTAATCCCAACATAAAAATGCTCGGTCTCGGGCGCAGAATTTCCGACGATCAATC
TCTCACACGTCCATTTTCTGCATTTGGGGCTTAAAGTGACGAAAATACCCAAATAAGACAACAAAGACACACAAGTTGATATAACATAATAGCAATAACCCCCTTCGCAGTCGGGTATACACTCGATTTTGGTACAATTCGCTCCATATAGCACTCGCCTATCGGCTCGTGCTATATGTCGCGCATTGTACCAAAATCTCGTGTATACCCTCCTGCGGC
GCACAGCGTGCAGCGTAACATGTCAATATAGAAATCGATTAACACTACGATGCGTCGCAGAGAGTATCGACCAATCAGAGAGCAGCATTCATCAAGCCACGAACTTTCACGTATGGCTTCGAGATTTCGCGTGATCTAGGGTGTTTACTTCAGTAACTCTACAATCTACTAAATACTAAGGGGTCCCAACTGTTCTACACATGCGCAAACGCCATCTTG
AAGAGTGGCTGATGATCATGTCGAAAACCGGAGGTCGAAAGACGTAAGTATCGCCGAGATATCGTAATCTGAATCTCGCGTTGCAATTTTTCACTCGATTATGTGTATCAAAATTATGCGGATACAATTTTTGGATTTTCTGTCAAACTTGGTTTTTCCTGTTGAAAGTTTGGGTTTTTCTCAGTCCGTAACGACATTGTTTACCGAAGGAACATGTGC
TGTACGCTTGGACTTTCTTCAAATGGCTTGGACTTTTAGATCTATACCCATCCAACATGTGTGATTGAGTGTAATGGCCGTTACATTTAAAATGTTCCTGTGATGGTAGCTAAGTCGGGAAATGATCGTTTTCTGTTTAATATATTTTTTAAGGTTTGAATCAGCGTGTTACGATATTTAGAATTCTGAGCAGCATCCTAGAGGGTCAATTTGGGTGTA
CAACGCATGTTCATGCGGTAGCTAGCTAACCCAGGCCATATGGCACCAGGTACAGGTTGACTAACGAAGTGTTTTGAGGTTAGCATAGTATGAGTACGTCAGATTCATATGTCATCGGCATAGAAGGTCGCTTGTGTAAGATAACACCATATTTTTAGGCTGTCATTCATTTCATTTATTTATTTTGCAGTGGGTTGCAGTCAGCAATGAATCATTTAC
AAGTTTGTTGCACTGTTATGCAATGCAATGCCTTGACCTGATTCAGTATTTCTTGCTTGGGTTCCTGCGGTATGAAGTAGAAAATTAGGCTGGCCACGGTGTCCTGGGAATTAATGTTTTACCACCAAGGTAAGAATAAAATTCTATTGGATTTATTGTTTACACATATATAAGGATGAAAGGTACAGACCATTGTACTGATTGGTACTGTACTTGCCA
AATTGATATGCTTAAATAATATTTTAAATATGTTAATTGTTTTCAGAGCGGAATGACGTCAGCTTTTCAACACAGAGAGTACCTTAAGAGGGTGTGTCATGTATGTGGTAGAACCCTGAAGACCACATCTTACAGCAACTCAGGACATTACAGACAGGGAGAAGTGTGCATCCCGTACAGTGAACTTTTAGAAGAGTGCTTTAAAGTAAATGTTCAAAA
```

TGATGACCCTTTTGTACATCCGCCTAGATTTGGTTCATGTTGCCGTAGAGTTCTTTTAAAGTATTCTGTTGCACAGAACCTGGGTAAACATTACTATACCCATATCAGTGTTTCTAACTGGAAACATCATTGTAAGTATGAAAACCAAACTTGTCCTCTTTGCAAACGTTTTTCTCAAAATGGTTCTGGTAGACCTAAAAAACTTAAAAGGGGTCGCTT
TAAATTAAATACAAATGAAAATAATGAAACTGACAAGAGTCCTGTTGCAGACAATTTTGCTCATATACTAAATGACCACTGTTATACCACCCCAGCTTCCTGTAAGTTTGCCCAGTCTGTGCAATCAGACCAAAAGACAGATACATATGAATTTTGTACAGAGGAAAATATTCAGACTGTACCCATGACAAGTCGATTTCCAACAGATTTATGTGTAAC
TGAACCACATCCAGAATTCATTTGTAGAATGTGTTTAGGAGTCCTAGACACACCACTTCTAACACCGTGTACTCACACTTTCTGTGCAGGCTGCATTAAGAACTGGTTAAGTCTGTGTCATTTTTGTCCTCTCTGTAAACAACCTATGGTGTCATCAGATTTAAAAGAACCATATAGAGGATTTGTAGAAATTCTTAATGATACTAGAATGAAATGTAA
TTGTCCAGAAGCAGTTTCAAGCAAAAAAGGTGTCAAGCTCAGTAATTGGCTCAGCACACTTTAAATTGTGGAAAGGATTTTGAAATTGATGAATGGTTAGAAAAAGTTTTGTCATTGTGGAACGAAATTTCTGTCATTCAAGTTGATTCATATTTAAATAGTACTGAACATTTGCCTGAGACAGATAAGTCAAGTCTGACAACAAGCCGTGGCCATCAA
AACATTGCTGCAGGTAGCACTGTTGGAAGACCTAAACTTCACTGTTGTGTCTCCAACAACAAGGACAAAATTTAAGATTGAAAAATATCAAAAAACAAATAGACGTGTTTGCCAGAGAGAAAGGGGAAGATCTCAGGGTAGTGTATTTTGGCCTTTTGCAAAACTATCTGAGAGCCAACAAGCAAGGTAGTTTAGCTGAACAAGTGAAGTTGCTATATA
ATGGGGAATCTTGTCAAATGTCACCTCAGGAATGTCTAGCCTTGCGTGTAAATACTTTTACCAGTTGTAATCAGTACAAGAAAATTTACTCGGCTACAAATGAGCTTAGTGGAGTTAAAATCTTCCAACCTCTCATCAATGTTGTGAAAGCCGAAGAAAGTTTTCTACCAGGGTCTGTAAAGTTCTACTTTAACCCATCTTTGAAAACTGTTTCTCCAT
TTGGCAAGGTACAGAGTATGTTGATGCTCTGTCAGGACTCAGTCACGATATACCACTTCTAGAATCACCTGATGTGTATGGGGCAAGGTATAGATATGACAAAGCAGTTGCTATGGCTTTGAAAGATATTGATAATGAAATCTCAGAAGGCCTCAGTAAATTAGAAGGGGTGGATACAAATCAACTTGTTCTAACAGCCACTGTCAAAGATGGCAGTGA
TGGTCTTGGAGATGTGAAAGAAGTCAGATCTCAGGGGCTTGTCGTTCCAAACAAGGCCTTGCGGTTTTCATTTGTAATTTTAGAAGTCACTGTCAATGTAGATGGCAAGATTATTACTGTATTTGAAGAAGATGCTCCCAACTCAGAATTATGCACACGCCCTCTCTTGGTAGCTCTGGCAGATGAAAATGACAGAAAAGGTTATGCATGACACATAT
ACCAATAATGATAGAGAGAGAGGAGATTGAGAAAAGTACCTTGTTTGTAGAAATTGAAGAGGGAAAACAAAGAATCTACAAAGTTGTATTTAAAGGGACTATGTATGATGAAAATTTGCAGAGAAGTGGTGAGGGAATGTTAGCTGCAGGGTCATCTTGGTTTTGTACACTTTGTGAGAAAGGAAGACTTGATCCCATATCATCAGTCTTGCCAATAAC
AAGAACACATGGAAACAATATGGCTTTGTATTCAAAATATGTTGAAAATGCAAATCATATGAAGCAGGCAGATCTAGTGAAAGAAGTCAAAGGCATTGTTAGAAAACCATTTATAATGTCAGAGCCTTGTGTTGATGCCACGCATGCTGAAATTCACTGGGGAGAGAAAAACTATGATATGTATGTACGTGAAATATCAGGGATAATGTGTTGGGGAA
GCCATCCAATGATGATGATGCAAAAAAAGTGAAATCAACGAAAAGAATGTTGGATACCGAATTACAGCTGCAATGTGGCTTAAAGAGAGAGCATGGCTTTATGGTTCATGGAAATTATGCACGTGATTTGGTCAAATCAGAAACAGTTGATGTGGTTGCTCTTTAATTCAGAGTGAAGCAAGAAAATCTGTTGTTCGTGAATATGTTTCCTACTATCGA
GAATTGCGATCAATATACAGGAAACACAATGCTGACTTAGATGATGCCGTGAAATTTAAAACCATAGCTGCAAATTTTTACAAATTGCTACATGAGAACTTTCCTTATTTGCATCCACTCTCTAACTATCAACACAAGGTATTAGATCACATTCCACAGTTGATCGAGCATATGGCTCTGTTGGTAGATTGCCTCTGAAGGCAATGAAAAAGGCAATAA
ACTTTTCAGAAGATTTCGTAAATTTAATGCACGTAAGTCTCAAATTCATGAAATGCCAGACATTCTGAAATTTCATTGGCTATATACAATGAAAAGTCTTCAGACTATGGTAAACTGGGAAAAAACTAGATCACTAGTCTGTAGCAAATGTGGTTGTGCTGGTCACAATCAAAGGACATGCATTACATAATTGTGCCCAAGGTCATGATATGAAATCAA
AGATGAGACAATGATCATGCATGTTTTATGTATCTGAATCTGTAAGTTTGAAATTAAGTAGCATCTGTATGTATCAGACTACAGAGTCCAAACTTCAGACACAAAGAGTAGGCTGGCCATACACTCTTATTTACTTAAAAAAGTGTTTCTGTTCCATATACTGACTGTACTGTCAAGAGTCAATTCTAGCTTCTATGCTAGTCTTAATATACCTAGATG
GTTAAAGTGTATTATATTTTGATGTACATAGATGAATTCCACAAGAAATCATCAAATACAAGAAAGCTGAGTTTGTGTTAAATATTTGTATATGTTTTAATCTTACCTGCCCACACTTTTGAAGGGACTCAAACAAAATCATTCAATTACTTTGGCCAAAATGTCAAAAGTTTGAGTAACTGGCCATGAATATGAGATGAGAAGTGTAAAGTTTGCAAT
AGAATTTTGTACATATATTCATGAAAGATTGTAACAAGTTGTAAAATGAACTCTCAGTAAAGGTTTCATTCAATTAATGCACATGCTACATGTACATGCTATAATTTCTTGTGCAGTTCTCATGTTCTCAGAAATTACAATCAGAACATGTAAATTTGATGTTACTCTTATTTTCCAGTTGTCGTGAAGAAAATCGTTTGCCAGAGCATTTAATGTGAA
AAGCTCTAACACATTGTGTACATCGCACTGTTGAAAAGTAACTCTTATCAGTCATGCTACAATTAATGTTAGCACATGTTGTGAAACATGAAGGGCATTCAAAGTTTATACAGTCCAGCTTTGTTCTATTTAAGTGATCTAATTCAACACATTTGTCATGGTACCACCTTCCACAGAATACAGTTTGATTGTTCATGATGTAAGTTCGATCACATTTCA
GCCAGTTGAACTTGGCTTGTTCAGTAGCTGCTAAATTGGCAAATTTGCAGCCATCCAACATACATTCCCATGTTTCCTCCTCACTATCATCTGATGTATTGTCTATCTTTCCTGCTCTGCTGCCCTCTTCTGACTCATTCCTGTCTGTGCTAACTTTGACCTCAGCTGCCTCACTTATTCCGTTCTGATCCTCAGTCTCTGAGCCTATACTTTCCTGAT
CTGTCATGTCTATGCTTATCCTTTCCTGCTCTTCCTCCGCTATGCTTACTCTGTCCTCATCTGTTGACCTTTCCTCCTCATCCCCCAACTGACTGTTATTACTCTGTGTCCTGGTTTCAGAGCTTGTCCCTTCATTTTGAAAACTGAATTTTATCATTTGCTTTTTTACTGTATCACAAACAGCAATTGAGCTGAAGAATGGAAGACTGTAGTATTTAC
TACTTAAGGTATCATAGTGTGCTGTTTCTATGAAATCTATGGCATCACAAGGTGACTTGTAAACTGTGTTGTTACTCAATATGTAAACAGTGTTATTACATTTGAATGAGACTGGGTTAGGAATACCAATAAATGGTACAGATGTACAGACCATTGAATTCAGGTCAATAACATCAACATCGGGTGGATGTCTTACATTGTTACCTATCACATAAATTC
CACCAACAATATACATTTTACTGCTATTTAGAAAAACTGAATGGTAGGCTCTTGGAGTGATAGCTTGTAAACTAATACTTGATAATTTCATAGATTGTAAATTAAGAACAGAAAATATGTTTGAAGCCATGGACACATTGTAAAAAGCATGACTGAGTTTTTTGTGTATATTGTAAGCCTCCATACAGATAAAGAGAATCTCCAAACTTAATTACACTG
TGACCAAAGCATGAATTTCCTCACCACTATATTGTCCTTGAGACCATGTTAATGAAATTTTCCCCCTTGGATCAACAAAGACAGAAAGCTGCCACAATTTTGTATTGTGCATTTCTGAGTCAGTCAATCCACCAAGGAGAACATAATGGTTTGGTTCAACTAAACACAAATTTGAGCCATGTACATGTGGAGGGTAATTTGACTGAGATGAATAATTTT
TGATTTCTTGTATTGTATCATCTTTGTCTATTACAAGCCAGCCTGTTACTGTTGTCTTAGACTTAACCCAGACCAAAACTCTGTCACCTAAAGAAATAACTGGATTTGTACATCCAGCAGTTGCTAGGCAAGTTGCCAGGATGTTTTTGGTGATTTGTATTTCTGAAATGAAAAAGATGATGACAAAAGTTGTTGTATTAGAGAGTGAAGCTTATCTGC
ATGCATGTATGGTGAGCAGGTCATTTTATTTTGTCAGTGTTGAAATATGGCACTGAAATATAAATTTGTTAATTTCATATTTTAGTATGGATATCTTTGTAGTTCAAAAAATGCTCAAGACAGAAAGTGAGATGTCTTTAGTCGTGACATTTATTAATATTGTTCACTGTACATTTCTCTACCCTTTGCAACCATACAATTGTTGAACACCTATCCTGT
ATATTCAAACAAACAACTTGAAGGCAAATCTGATATCTTTGGCAGCATGTTCTCTTGTCTTTTGCATTTGAGTAACCTCCTTATATAAATATGATCGAAAGAGATTGTCAACATGGAGCAGCAAATCCGTTTCTTTAACCTAGACTGTAAATCAATACAAAATGGGTGCACATGTAGCTAGGGGAAAAGGTAATGCACAGATAGAGCCCCATTGATGA
CTGCTGGAAATGTCATTTGGATGTCTTGCTGGTGTACATGCAACTGAAAAAATCATTTTAGCCTGTGGGCCGTACAATAGCTGCACTTCTACTTCGCTCACTCTAATTTGAATCTTCGTATATAGTGTACATGGGGATCGACTTTCATAGTATGCTTGTTGGATCGAGGGACCGTGGCTTGTGTTACAGTTGAGAACGATCACATCGAGAATTTTCTTT
TTCGAATGGGCGATTACAAGTGCAAATCTTTGGAAGTTAGCTACCATATAAGAACTTTGTTGTTTTAACTTCAAGAATAATCAATCTAAAACACGAACAGAATGTTCTGTGACTTTTGAAGCTACATTGAACTGACAACCTACCGGGACTATCGAGCATGGCGGCTGCCATCTTGGATATTGTTTACGGCTCACTGAGCATGCTCAGATCAAAAATATT
TCCCACAACCCATTGCGGACTTTCTACATTGTAAACATAGGAGATGTTACGGTGTGCGGCGTGGGTTATTGCTTAATTTTTTGCGCTGCATTTTTATCGGCTAACTTCATACGTAATCGATGATGGAAATAGAATTATGCCGTCAATTCCCCTCTAAAATACAGTGAAAATATGATAATAAGGTAAGTCTTTGGTCAAGAAGAACAATTTTAAGATGCA
TTAAGTAAGTTTTTGAAGAGGAACTAATGACAAACCAAAAACATCAAAAAAATTCGAACATGCATATTGCTATGATATCCCTACAAACATGCATACAAAATTTCAAATAATATGTGCTTTATAGAAACACTTTTGACTATATAAACAATTATGAATATCCAATATTTCTGAAATAAAGAAGCGTTGATGTGTAAGTGAAAATTTGAATGGCCATAAAAT
CTAGCTCCCCGTCCACAGACTCTCTTTTTCACTTTCCTTGAGAGATTTTTCTATATTACAAATGATGTTGACAAAATCATTCACACGTTCGACATGTGTTTGATCTCGAATTTTCGAAAATACTAATCGGTTTACTGATTGGTCGGAATGTCTTGCATGCTAATAATGCACTTGGCTCTGTTATTTTCCAGACTGTATTTGTTAGACAACGAACTGGAG
TCGATAATAAGTCTCGACTTGAATTCTGAAAATGCTGAACCCTATACTTTCAAAAGAATGAATGATTTACAATGGTTGTTTTCAATCGATATAGACCCGGTAGGTATGACACAGAAATTTAAAATAAATTTTAAATTTTATTGCAGAAAAACGTTGGCCAAAGTTGTGTCGACGAATCTGTCGATAGTCTACAGATTTCTGCATCGAGGATTGAGAATG
TTATGTATGAGACTGACTTTGAACACTTATTAAAATGTCTGATATCAATCACTCAATATACAACTAGACGTTCACCCACATATCTTTCTTTCATGCTTTTGTGTGCAATGTATTTTCTTCGGTACATTGGTTCTAAAAGCTTTATGTCTGTGCTAGTAAATCTTTAACGTAGTAATGTGATTTGTGAATTTACATAGCATTGAATTTCTTAAATTTTAT
TAATTTTTGTACCTTTATATTCACATCGTTACATTCCAAATGTTTGATATTCATACTTTCCTTTGCATTTCTCGAAATAGTAAGCCGTCTGTGCACTTACGTTATGTAAAATCAAATAATGCACAAGGCAATTTTTTATGAAACCAATGGGAGCCTAATCCATTACTTTTACCTCTTGTTTGCAGAACTTTGGTTTTTTTGTCACGGATAGAGAAAATA
AATTGCATTACATTCCATATGATCCGAGTATGATTCAAGATCTGTAGGGCTGGGAGATTATGTACCATACGGAGTAGCATTCTTTTCAAACCAAAGACAGCCTAAAATTTCTAGTAAGTAGCAAATTTCGAAGATTAAACCAACTGTGAAGTATGGAATACCTCTCCTGGGCGAATACACATAGGTGGATTAAAGCATATTATGATTTACTATAAGTTG
TGTTCTTTTTCTTGAGGATTTTCCAACCTTGTATATTGAATTCCTTCATTGTAATAACGTGATTTTAAGCGTCATTTTTCTTAGCGCCCCATACCCACTTGTTTATCATTTAAAACGGATGATGATATAGAACTGACACGGATAATTGGATATCCTGTGTACGTATTTTATTGATCATGTCATTTTGCATATTGACAGAATTTCTATAATTGGAGTTTT
AATATAATATACAAGAACTTTACGATTTTCAGCGATGTCTGAAGATCCAAGAACACTTTCATGATGTCCCTTTATTTTATTCACGCTGCTTCTAGTAATATTCTTTAAGTCAAGAAGACCGTCATCTGTACATCATCTTTATTATACTTGTAGAAAGGTAGCTGATGATGTAAAAGCCTACCAGTTCACTTATCTAGGTTCAATGGTAGCTCCCATGAC
TTTATTCGGAAATACGTAAGGCCTCGTTTCATCTATGACGGAGTTTTGTAAATTATGTTACATTCAAATATTGATAGTGCAAATACAGCGATCTGATTGGTCGAAACCTAAAAAGAACCGTACAGCAAGGTTGGCACATGCGGAAGCTTTTGGCAAGAGTAAAAATCCTATTTGAGGCTAGTCAACACTAGTCAATTTTCCAATTTATTCACTCATAAT
TTATTCATTCAAATACGTGATAGAAATTGTAATAATATATACATTTTAAAGGTTTCGTTTTCATTTGCAAATGAAGTGCTTAACTTTGGCCATTTTCGACATAAAAGGATAATCGACTCCGAATCATAAGAAATTGGTAAAAGTCCTTGTGAGCATAATACTTTACGGCTAGCAGGTATATGTCATTTGCAAACGTCATCGTAAATGCTGATAGGCTT
GGAAACATTCCCTGTTCATTTATTCTGACCACTCTGTGCAGTATGTAGTGGAAATCGGTAACTTTT

#### FASTA protein

>PflTRAG2L-A_16(from v1)
MAAAMLDSPEIQITKNILATCLATAGCTNPVISLGDRVLVWVKSKTTVTGWLVIDKDDTIQEIKNYSSQSNYPPHVHGSNLCLVEPNHYVLLGGLTDSEMHNTKLWQLSVFVDPRGKISLTWSQGQYSGEEIHAFGHSVIKFGDSLYLYGGLQYTQKLSHAFYNVSMASNIFSVLNLQSMKLSSISLQAITPRAYHSVFLNSSKMYIVGGIYVIGNNVR
HPPDVDVIDLNSMVCTSVPFIGIPNPVSFKCNNTVYILSNNTVYKSPCDAIDFIETAHYDTLSSKYYSLPFFSSIAVCDTVKKQMIKFSFQNEGTSSETRTQSNNSQLGDEEERSTDEDRVSIAEEEQERISIDMTDQESIGSETEDQNGISEAAEVKVSTDRNESEEGSRAGKIDNTSDDSEEETWECMLDGCKFANLAATEQAKFNWLKCDRTYIMN
NQTVFCGRWYHDKCVELDHLNRTKLDCINFECPSCFTTCANINCSVTDKSYFSTVRCTQCVRAFHIKCSGKRFSSRQLENKSNIKFTCSDCNF

### PflTRAGL-A_29

| Genomic loci | RAG1L status | RAG2L status |
|---|---|---|
| PflTRAGL-A_29_v1 | *pseudogenic (frameshifts)* | *pseudogenic (frameshifts)* |
| PflTRAGL-A_29_v2 | *pseudogenic (frameshifts)* | *pseudogenic (frameshifts)* |

```
           1                                                                                                                                                                                                      200
PflT_29_v1 TGCTCCTTCTGCATTGATAATGTAGTGTTGTAAGAGCTCATCAAATTTATTTGGTAAACTGTCAGCATTAGTATATATACAATATACGTTATGTCCACGAAAACTATGTACATTCCTATTTACAATATTTTTCCTCTCGTTAGATTGTATCCTTTCCTTTCACTTTGATACTTTTCCTTTCCAAGGTGGTCCTCTCACCA
PflT_29_v2 TGCTCCTTCTGCATTGATAATGTAGTGTTGTAAGAGCTCATCAAATTTATTTGGTAAACTGTCAGCATTAGTATATATACAATATACGTTATGTCCACGAAAACTATGTACATTCCTATTTACAATATTTTTCCTCTCGTTAGATTGTATCCTTTCCTTTCACTTTGATACTTTTCCTTTCCAAGGTGGTCCTCTCACCA

           201            TSD         5'-TIR5                                                                                                                                                                 400
PflT_29_v1 GGAACGGAAGTCCCCGACGCACAGCGTAACATGTCAATATAGAAATCGATTAACACTACGATGAGTCGCAGAGAGTATCGACCAATCAGAGAGCAGCATTCATCAAGCCACGAACTTTCACGTATGGCTTCGAGATTTCGCGTGATCTAGGGTGTTTACTTCAGTAACTCTACAATCTACTAAATACTAAGGGGTCTCAA
PflT_29_v2 GGAACGGAAGTCCCCGATGCACAGCGTAACATGTCAATATAGAAATCGATTAACACTACGATGAGTCGCAGAGAGTATCGACCAATCAGACAGCAGCATTCATCAAGCCACGAACTTTCACGTATGGCTTCGAGATTTCGCGTGATCTAGGGTGTTTACTTCAGTAACTCTACAATCTACTAAATACTAAGGGGTCTCAA

           401                                                                                                                                                                                                    600
PflT_29_v1 CTGTTCTACACATGCGCAAACGCCATCTTGAAGAGTGGCTGATGATCATGTCGAAAACCGGAGGTCGAAAGACGTAAGTATCGCCGAGATATCGTAATCTGAATCTCACGTTGCAATTTTTCACTCGATTATGTGTATCAAAATTATGCGGATACAATTTTTGGATTTTCTGTCAAACTTGGTTTTTCCTGTTGAAAGTT
PflT_29_v2 CTGTTCTACACATGCGCAAACGCCATCTTGAAGAGTGGCTGATGATCATGTCGAAAACCGGAGGTCGAAAGACGTAAGTATCGCCGAGATATCGTAATCTGAATCTCACGTTGCAATTTTTCACTCGATTATGTGTATCAAAATTATGCGGATACAATTTTTGGATTTTCTGTCAAACTTGGTTTTTCCTGTTGAAAGTT

           601                                                                                                                                                                                                    800
PflT_29_v1 TGGGTTTTTCTCAGTCCGTAACGACATTGTTTACCGAAGGAACATGTGCTGTACGCTTGGACTTTCTTCAAATGGCTTGGACTTTTAGATCTATACCCATCCAACATGTGTGATTGAGTGTAATGGCCGTTATATTTAAAA-TGTTCCTGTGATGGTAGCTAAGTCGGGGAAATGATCGTTTTCTGTTTAATATATTTTT
PflT_29_v2 TGGGTTTTTCTCAGTCCGTAACGACATTGTTTACCGAAGGAACATGTGCTGTACGCTTGGACTTTCTTCAAATGGCTTGGACTTTTAGATCTATACCCATCCAACATGTGTGATTGAGTGTAATGGCCGTTATATTTAAAAATGTTCCTGTGATGGTAGCTAAGTCGGGGAAATGATCGTTTTCTGTTTAATGTATTTTT

           801                                                                                                                                                                                                    1000
PflT_29_v1 TAAGGTTTGAATCAGCGTGTTACGATATTTAGAATTCCGAGCAGCATCCTAGAGGGTCAATTTGGGTGTACAACGCATGGTTCATGCGGTAGCAAGCTAACCCAGGCCATATGGCACCAGGTACAGGTTGACTAACGAAGTGTTTTGAGGTTAGCATAGTATGAGTACGTCAGATTCATATGTCATCGGCATAGAAGGTC
PflT_29_v2 TAAGGTTTGAATCAGCGTGTTACGATATTTAGAATTCTGAGCAGCATCCTAGAGGGTCAATTTGGGTGTACAACGCATGGTTCATGCGGTAGCAAGCTAACCCAGGCCATATGGCACCAGGTACAGGTTGACTAACGAAGTGTTTTGAGGTTAGCATAGTATGAGTACGTCAGATTCATATGTCATCGGCATAGAAGGTC

           1001                                                                                                                                                                                                   1200
PflT_29_v1 GCTTGTGTAAGATAACACCATATTTTTAGGCTGTCATTCATTTCATTTATT-ATTTTGCAGTGGGTTGCAGTCAGCAATGAATCATTTACAAGTTTGTTGCACTGTTATGCAATGCAATGCCTTGACCTGATTCAGTATTTCTTGCTTTGGTCCTGCGGTATGAAGTAGAAAATTAGGCTGGCCACGGTGTCCTTGGAA
PflT_29_v2 GCTTGTGTAAGATAACACCATATTTTTAGGCTGTCATTCATTTCATTTATTTATTTTGCAGTGGGTTGCAGTCAGCAATGAATCATTTACAAGTTTGTTGCACTGTTATGCAATGCAATGCCTTGACCTGATTCAGTATTTCTTGCTTTGGTCCTGCGGTATGAAGTAGAAAATTAGGCTGGCCACGGTGTCCTTGGAA

           1201                                                                                                                                                                                                   1400
PflT_29_v1 TTAATGTTTTACCACCAAGGTAAGAATAAAATTCTATTGGATTTATTGTTTACACATATATAAGGATGAAAGGTACAGACCATTGAACTGATTGGTACTGTACTTGCCAAATTGATATGCTTAAATAATATTTTAAATATGTTAATTGTTTTCAGAGCGGAATGACGTCAGCTTTTCAACACAGAGAGTACCTTAAGAGG
PflT_29_v2 TTAATGTTTTACCACCAAGGTAAGAATAAAATTCTATTGGATTTATTGTTTACACATATATAAGGATGAAAGGTACAGACCATTGAACTGATTGGTACTGTACTTGCCAAATTGATATGCTTAAATAATATTTTAAATATGTTAATTGTTTTCAGAGCGGAATGACGTCAGCTTTTCAACACAGAGAGTACCTTAAGAGG

           1401                                                                                                                                                                                                   1600
PflT_29_v1 GTGTGTCGTGTATGTGGTAGAACCCTGAAGACCACATCTTACAGCAACTCAGGACATTACAGACAGGGAGAAGTGTGCATCCCGTACAGTGAACTTTTAGAAGAGTGCTTTAAAGTAATGTTCAAAATGATGATCCTTTTGTACATCCGCCTAGATTTGGTTCATGTTGTCGTAGAGTTCTTTTAA-GTATTCTGTTGC
PflT_29_v2 GTGTGTCGTGTATGTGGTAGAACCCTGAAGACCACATCTTACAGCAACTCAGGACATTACAGACAGGGAGAAGTGTGCATCCCGTACAGTGAACTTTTAGAAGAGTGCTTTAAAGTAATGTTCAAAATGATGATCCTTTTGTACATCCGCCTAGATTTGGTTCATGTTGTCGTAGAGTTCTTTTAAAGTATTCTGTTGC

           1601                                                                                                                                                                                                   1800
PflT_29_v1 ACAGAACCTGGGTAAACATTACTATACCCATATCAGTGTTTGTAACTGGAAACATCATTGTAAGTATGAAAACCAAACTTGTCCTCTTTGCAAACGTTTTTCTCAAAATGGTTCTGGTAGACCTAAAAAACTTAAAAGGGGTCGCTTAAATTAAATACAAATGAAAATAATGAAACTGACAAGAGTCCTGTTGCAGACAA
PflT_29_v2 ACAGAACCTGGGTAAACATTACTATACCCATATCAGTGTTTGTAACTGGAAACATCATTGTAAGTATGAAAACCAAACTTGTCCTCTTTGCAAACGTTTTTCTCAAAATGGTTCTGGTAGACCTAAAAAACTTAAAAGGGGTCGCTTAAATTAAATACAAATGAAAATAATGAAACTGACAAGAGTCCTGTTGCAGACAA

           1801                                                                                                                                                                                                   2000
PflT_29_v1 TTTTGCTCATATACTAAATGACCACTGTTATACCACCCCAGCTTCCTGTAAGTTTGCCCAGTCTGTGCAATCAGACCAAAAGACAGATACATATGAATTTGTACAGAGGAAAATATTCAGACTGTACCCATGACAAGTCGATTTCCAACAGATTTATGTGTAACTGAACCACATCCAGAATTCATTTGTAGAATGTGTTT
PflT_29_v2 TTTTGCTCATATACTAAATGACCACTGTTATACCACCCCAGCTTCCTGTAAGTTTGCCCAGTCTGTGCAATCAGACCAAAAGACAGATACATATGAATTTGTACAGAGGAAAATATTCAGACTGTACCCATGACAAGTCGATTTCCAACAGATTTATGTGTAACTGAACCACATCCAGAATTCATTTGTAGAATGTGTTT

           2001                                                                                                                                                                                                   2200
PflT_29_v1 AGGAGTCCTAGACACACCACTTCTAACACCGTGTACTCACACTTTCTGCGCAGGCTGCATTAAGAACTGGTTAAGTCTGTGTCATTTTTGTCCTCTCTGTAAACAACCTATGGTGTCATCAGATTTAAAAGAACCATATAGAGGATTTGTAGAAATTCTTAATGATACTAGAATGAAATGTAATTGTCCAGAAGCATTTT
PflT_29_v2 AGGAGTCCTAGACACACCACTTCTAACACCGTGTACTCACACTTTCTGCGCAGGCTGCATTAAGAACTGGTTAAGTCTGTGTCATTTTTGTCCTCTCTGTAAACAACCTATGGTGTCATCAGATTTAAAAGAACCATATAGAGGATTTGTAGAAATTCTTAATGATACTAGAATGAAATGTAATTGTCCAGAAGCATTTT

           2201                                                                                                                                                                                                   2400
PflT_29_v1 CAAGCAAAAAAGGTGTCAAGCTTAGTGATTGGCTTCAGCACACTTTAAATTGTGGAAACGATTTTGAAATTGATGAATGGTTAGAAAAAGTTTTGTCATTGTGGAACGAAATTTCTCTCACTCAAGTTGATTCACATTTAAATAGTACTGAACATTTGCCTGAGACAGATAAGTCTAGTCTGACAACAAGCCATGGCCAT
PflT_29_v2 CAAGCAAAAAAGGTGTCAAGCTTAGTGATTGGCTTCAGCACACTTTAAATTGTGGAAACGATTTTGAAATTGATGAATGGTTAGAAAAAGTTTTGTCATTGTGGAACGAAATTTCTCTCACTCAAGTTGATTCACATTTAAATAGTACTGAACATTTGCCTGAGACAGATAAGTCCAGTCTGACAACAAGCCATGGCCAT

           2401                                                                                                                                                                                                   2600
PflT_29_v1 CCAGACATTGCTGCAGGTAGCACTGTGGAAGACCTAAAACTTCACTGTTGTGTCTCCAACAACAAGGACAAAATTTAAGATTGAAAAATATCAAAAAACAAATAGACGTGTTTGCCAGAGAGAAAGGGGAAGATCTCAGGGTAGTGTATTTTGGCCTTTTGCAAAAACTATCTGAGAGCCAACAAGCAAGGTAGTTTAGC
PflT_29_v2 CAAGACATTGCTGCAGGTAGCACTGTGGAAGACCTAAAACTTCACTGTTGTGTCTCCAACAACAAGGACAAAATTTAAGATTGAAAAATATCAAAAAACAAATAGACGTGTTTGCCAGAGAGAAAGGGGAAGATCTCAGGGTAGTGTATTTTGGCCTTTTGCAAAAACTATCTGAGAGCCAACAAGCAAGGTAGTTTAGC

           2601                                                                                                                                                                                                   2800
PflT_29_v1 TGAACAAGTGAAGTTGCTATATAATGGGGAATCTTGTCAAATGTCACCTCAGGAATGTCTAGCCTTGCGTGTAAATACTTT-ACCAGTTGTAATCAGTACAAGAAAATTTACTCGGCTACAAATGAGCTTAGTGGAGTTAAAATCTTCCAACCTCTCATCAATGTTGTGAAAGCTGAAGAAAGTTTTCTACCAGGGTCTG
PflT_29_v2 TGAACAAGTGAAGTTGCTATATAATGGGGAATCTTGTCAAATGTCACCTCAGGAATGTCTAGCCTTGCGTGTAAATACTTTTACCAGTTGTAATCAGTACAAGAAAATTTACTCGGCTACAAATGAGCTTAGTGGAGTTAAAATCTTCCAACCTCTCATCAATGTTGTGAAAGCTGAAGAAAGTTTTCTACCAGGGTCTG

           2801                                                                                                                                                                                                   3000
PflT_29_v1 TAAAGTTCTACTT-AACCCATCTTTGAAAACTGTTTCTCCCATTTGGCAAGGTACAGAGCATGTTGATGCTCTGTCAGGACTCAGTCGCGATATACCACTTCTAGAGTCACCTGATGTTTATGGGGCAAGGTATAGATATGACAAAGCAGTTGCTATGGCTTTGAAAGATATTGATAATGAAATCTCAGAAGGCCTCAGT
PflT_29_v2 TAAAGTTCTACTTTAACCCATCTTTGAAAACTGTTTCTCCCATTTGGCAAGGTACAGAGCATGTTGATGCTCTGTCAGGACTCAGTCGCGATATACCACTTCTAGAGTCACCTGATGTTTATGGGGCAAGGTATAGATATGACAAAGCAGTTGCTATGGCTTTGAAAGATATTGATAATGAAATCTCAGAAGGCCTCAGT

           3001                                                                                                                                                                                                   3200
PflT_29_v1 AAATTAGAAGGGGTGGATACAAATCAACTTGTTCTAACAGCCACTGTCAAAGATGGCAGTGATGGTCTTGGCGATGTGAAAGAAGTCAGATCTCAGGGGCTTGTCGTTCCAAACAAGGCCTTACGGTTTTCATTTGTAATTTTAGAAGTCACTGTCAATGTAGATGGCAAGATTATTACTGTATTTGAAGAAGATGCTCC
PflT_29_v2 AAATTAGAAGGGGTGGATACAAATCAACTTGTTCTAACAGCCACTGTCAAAGATGGCAGTGATGGTCTTGGCGATGTGAAAGAAGTCAGATCTCAGGGGCTTGTCGTTCCAAACAAGGCCTTACGGTTTTCATTTGTAATTTTAGAAGTCACTGTCAATGTAGATGGCAAGATTATTACTGTATTTGAAGAAGATGCTCC

           3201                                                                                                                                                                                                   3400
PflT_29_v1 CAACTCAGAATTATGCACACGCCCTCTCTTGGTAGCTCTGGCAGATGAAAATGACAGAAAAGGGTTATGCATGACACATATACCAATAATGATAGAGAGAGAGGAGATTGAGAAAAGTACCTTGTTTGTAGAAATTGAAGAGGGAAAACAAAGAATCTACAAAGTTGTATT-AAAGGGACTATGTATGATGAAAAATTGC
PflT_29_v2 CAACTCAGAATTATGCACACGCCCTCTCTTGGTAGCTCTGGCAGATGAAAATGACAGAAAAGGGTTATGCATGACACATATACCAATAATGATAGAGAGAGAGGAGATTGAGAAAAGTACCTTGTTTGTAGAAATTGAAGAGGGAAAACAAAGAATCTACAAAGTTGTATTTAAAGGGACTATGTATGATGAAAAATTGC
```

```
           3401                                                                                                                                                                                                  3600
PflT_29_v1  AGAGAAGTGGTGAGGGAATGTTAGCGGCAGGGTCATCTTGGTTTTGTACACTTTGTGAGAAAGGAAGACTTGATCCCATATCATCAGTCTTGCCAATAACAAGAACACATGAAAACAATATGGCTTTGTATTCAAAGTATGTTGAAAATGCAAATCATATGAAGCAGGCAGATCTAGTGAAAGAAGTCAAAGGCATTGTT
PflT_29_v2  AGAGAAGTGGTGAGGGAATGTTAGCGGCAGGGTCATCTTGGTTTTGTACACTTTGTGAGAAAGGAAGACTTGATCCCATATCATCAGTCTTGCCAATAACAAGAACACATGAAAACAATATGGCTTTGTATTCAAAGTATGTTGAAAATGCAAATCATATGAAGCAGGCAGATCTAGTGAAAGAAGTCAAAGGCATTGTT

           3601                                                                                                                                                                                                  3800
PflT_29_v1  AGAAAACCATTTATAATGTCAGAGCCTTGTGTTGATGCCACGCATGCTGAAATTCACTGGGGAGAGAAAAACTATGATATGTATGTACGTGAAATATCAGGTATAATGTGTTGGGGGAAGCCATCCAATGATGATGATGCAAAAAAAGTGAAATCAACGAAAAGAATGTTGGATACCGAATTACAGCTGCAATGTGGCTT
PflT_29_v2  AGAAAACCATTTATAATGTCAGAGCCTTGTGTTGATGCCACGCATGCTGAAATTCACTGGGGAGAGAAAAACTATGATATGTATGTACGTGAAATATCAGGTATAATGTGTTGGGGGAAGCCATCCAATGATGATGATGCAAAAAAAGTGAAATCAACGAAAAGAATGTTGGATACCGAATTACAGCTGCAATGTGGCTT

           3801                                                                                                                                                                                                  4000
PflT_29_v1  AAAGAGAGAGCATGGCTTTATGGTTGATGGAAATTATGCACGTGATTTGGTCAAATCAGAAACAGTTGATGTGGTTTGCTCTTTAATTCAGAGTGAAGCAAGAAAATCTGTTGTTCGTGAATATGTTTCCTACTATCGAGAATTGCGATCAATATACAGGAAACACAATGCTGACTTAGATGATGCCGTGAAATTTAAAA
PflT_29_v2  AAAGAGAGAGCATGGCTTTATGGTTGATGGAAATTATGCACGTGATTTGGTCAAATCAGAAACAGTTGATGTGGTTTGCTCTTTAATTCAGAGTGAAGCAAGAAAATCTGTTGTTCGTGAATATGTTTCCTACTATCGAGAATTGCGATCAATATACAGGAAACACAATGCTGACTTAGATGATGCCGTGAAATTTAAAA

           4001                                                                                                                                                                                                  4200
PflT_29_v1  CCATAGCTGCAAATTTT-ACAAATTGCTACATGAGAAATTTCCT-ATTTGCATCCACTCTCTAACTATCAACACAAGGTATTAGATCACATTCCACAGTTGATCGAGCAATATGGCTCTGTTGGTAAATTTGCCTCTGAAGGCAATGAAAAAGGCAATAAACTTTTCAGAAGATTTCGTAAATTTAATGCACGTAAGTCT
PflT_29_v2  CCATAGCTGCAAATTTTTACAAATTGCTACATGAGAAATTTCCTTATTTGCATCCACTCTCTAACTATCAACACAAGGTATTAGATCACATTCCACAGTTGATCGAGCAATATGGCTCTGTTGGTAAATTTGCCTCTGAAGGCAATGAAAAAGGCAATAAACTTTTCAGAAGATTTCGTAAATTTAATGCACGTAAGTCT

           4201                                                                                                                                                                                                  4400
PflT_29_v1  CAAATTCATGAAATGCCAGACATTCTGAAATTTCATTGGCTATATACAATGAAAAGTCTTCAGATTATGGTAAACTGGGAAAAAACTAGATCACTAGTCTGTAGCAAATGTGGTTGTGCAGGTCACAATCAAAGGACATGCATTACATAATTGTGCCCAAGGCCATGATATGAAATCAAAGATTAGATAATGACCATGCA
PflT_29_v2  CAAATTCATGAAATGCCAGACATTCTGAAATTTCATTGGCTATATACAATGAAAAGTCTTCAGACTATGGTAAACTGGGAAAAAACTAGATCACTAGTCTGTAGCAAATGTGGTTGTGCTGGTCACAATCAAAGGACATGCATTACATAATTGTGCCCAAGGTCATGATATGAAATCAAAGATTAGATAATGACCATGCA

           4401                                                                                                                                                                                                  4600
PflT_29_v1  TGTTTTATGTATCTGAATCTGTAAGTTTGAAATTAAGTAGCATCTGTATGTATCAGACTACAGAGTCCAAACTTCAGACACAAAGAGTAGGCTGGCCATACACTCTTATTTACTTGAAAAGTGTTTCTGTTCCATATACTGACTGTACTGTCAGGAGTCAATTCTAGCTTCTATGCTTCTATACCTAGATAGTTAAAGTG
PflT_29_v2  TGTTTTATGTATCTGAATCTGTAAGTTTGAAATTAAGTAGCATCTGTATGTATCAGACTACAGAGTCCAAACTTCAGACACAAAGAGTAGGCTGGCCATACACTCTTATTTACTTGAAAAGTGTTTCTGTTCCATATACTGACTGTACTGTCAGGAGTCAATTCTAGCTTCTATGCTTCTATACCTAGATAGTTAAAGTG

           4601                                                                                                                                                                                                  4800
PflT_29_v1  TATTATATTTTGATGTACATAGATGAATTCCACAAGAAAACATCAAATATAAGAAAGCTGAGTTTGTGTTAAATATTTGTATGTGTTTTAATCTTACCTGCCCACACTTTTGAAGGGACTCAAACAAAATAATTTGATTACTGTGGCCAAAATGTCAAAAGTTTGAGTAACTGGCCATGAATATGAGATGAGAAGCTTAA
PflT_29_v2  TATTATATTTTGATGTACATAGATGAATTCCACAAGAAAACATCAAATATAAGAAAGCTGAGTTTGTGTTAAATATTTGTATGTGTTTTAATCTTACCTGCCCACACTTTTGAAGGGACTCAAACAAAATAATTTGATTACTTTGGCCAAAATGTCAAAAGTTTGAGTAACTGGCCATGAATATGAGATGAGAAGCGTAA
 Consensus  TATTATATTTTGATGTACATAGATGAATTCCACAAGAAAACATCAAATATAAGAAAGCTGAGTTTGTGTTAAATATTTGTATGTGTTTTAATCTTACCTGCCCACACTTTTGAAGGGACTCAAACAAAATAATTTGATTACTgTGGCCAAAATGTCAAAAGTTTGAGTAACTGGCCATGAATATGAGATGAGAAGCgTAA

           4801                                                                                                                                                                                                  5000
PflT_29_v1  GGTTTGCAATAGAATTTTGTACATATATTCATGAAAGATTGTAACAAGTTGTAAAATGAACTCTCAGTAAAGGTTTCATTCAATTCATGCACATGCTACATGTACATGCTATGATTTCTTGTGCAGTTGTCATGTTCTCAGAAATTACAATCAGAACATGTAAATTTGATGTTACTCTTATTTTCCAGTTGTCGTGAAGA
PflT_29_v2  GGTTTGCAATAGAATTTTGTACATATATTCATGAAAGATTGTAACAAGTTGTAAAATGAACTCTCAGTAAAGGTTTCATTCAATTCATGCACATGCTACATGTACATGCTATGATTTCTTGTGCAGTTGTCATGTTCTCAGAAATTACAATCAGAACATGTAAATTTGATGTTACTCTTATTTTCCAGTTGTCGTGAAGA

           5001                                                                                                                                                                                                  5200
PflT_29_v1  AAATCGTTTGCCAGAGCATTTAATGTGAAAAGCTCTAACACATTGTGTACATCGCACTGTTGAAAAGTAACTCTTATCAGTCACGCTACAATTAATGTTAGCACATGTTGTGAAACATGAAGGGCATTCAAAGTTTATACAGTCCAGCTTTGTTCTATTTAAGTGATCTAATTCAACACATTTGTCATGGTACCACCTTC
PflT_29_v2  AAATCGTTTGCCAGAGCATTTAATGTGAAAAGCTCTAACACATTGTGTACATCGCACTGTTGAAAAGTAACTCTTATCAGTCACGCTACAATTAATGTTAGCACATGTTGTGAAACATGAAGGGCATTCAAAGTTTATACAGTCCAGCTTTGTTCTATTTAAGTGATCTAATTCAACACATTTGTCATGGTACCACCTTC

           5201                                                                                                                                                                                                  5400
PflT_29_v1  CACAGAATACAGTTTGATTGTTCATGATGTAAGTTCGCTCACATTTCAGCCAATTGAACTCTGCTTTATCACTTGCTGGTAAACTAGCAAACTTGCAGCCATCCAACATACATTCCCATGTTTCATTCGTCACTGTCATCTGATGTATTGTCTATCTTTCCTGCTCTGCTGCCCTCTTCTGACTCATTCCTGTCTGTGCT
PflT_29_v2  CACAGAATACAGTTTGATTGTTCATGATGTAAGTTCGCTCACATTTCAGCCAATTGAACTCTGCTTTATCACTTGCTGGTAAACTAGCAAACTTGCAGCCATCCAACATACATTCCCATGTTTCATTTGTTACTGTCATCTTATGTATTGTCTATCTTTCCTGCTCTGCTGCCCTCTTCTGACTCATTCCTGTCTGTGCT

           5401                                                                                                                                                                                                  5600
PflT_29_v1  AACTTTGACCTCAGCTGCCTCACTTATTCCGTTCTGATCCTCAGTCTCTGAGCCTATACTTTCCTGATCTGTCATGTCTATGCTTATCCTTTCCTGCTCTTCCTCCGCTATGCTTACTCTGTCCTCATCTGTTGACCTTTCCTCCTCATCCCCCAACTGACTGTTATTACTCTGTGTCCTGGTTTCAGAGCTTGTCCCTT
PflT_29_v2  AACTTTGACCTCAGCTGCCTCACTTATTCCGTTCTGATCCTCAGTCTCTGAGCCTATACTTTCCTGATCTGTCATGTCTATGCTTATCCTTTCCTGCTCTTCCTCCGCTATGCTTACTCTGTCCTCATCTGTTGACCTTTCCTCCTCATCCCCCAACTGACTGTTATTACTCTGTGTCCTGGTTTCAGAGCTTGTCCCTT

           5601                                                                                                                                                                                                  5800
PflT_29_v1  CATTTTGAAAACTGAATTTTATCATTTGCTTTTTTACTGTATCACAAACAGCAATTGAGCTGAAGAATGGAAGACTGTAGTATT-ACTACTTAAGGTATCATAGTGTGCTGTTTCTATGAAATCTATGGCATCACAAGGTGACTTGTAAACTGTGTTGTTACTCAATATGTAAACAGTGTTACTACATTTGAATGAGACT
PflT_29_v2  CATTTTGAAAACTGAATTTTATCATTTGCTTTTTTACTGTATCACAAACAGCAATTGAGCTGAAGAATGGAAGACTGTAGTATTTACTACTTAAGGTATCATAGTGTGCTGTTTCTATGAAATCTATGGCATCACAAGGTGACTTGTAAACTGTGTTGTTACTCAATATGTAAACAGTGTTACTACATTTGAATGAGACT

           5801                                                                                                                                                                                                  6000
PflT_29_v1  GGGTTAGGAATACCAATAAATGGTACAGATGTACAGACCATTGAATTCAGGTCAATTACATCAACATCGGGTGGATGTCTTACATTGTTACCTATCACATAAATTCCACCAACAATATACATTTTACTGCTATTACGAAAAACTGAATGGTAGGCTCTTGGAGTGATCGCTTGTAAACTAATACTTGATAATTTCATAGA
PflT_29_v2  GGGTTAGGAATACCAATAAATGGTACAGATGTACAGACCATTGAATTCAGGTCAATTACATCAACATCGGGTGGATGTCTTACATTGTTACCTATCACATAAATTCCACCAACAATATACATTTTACTGCTATTACGAAAAACTGAATGGTAGGCTCTTGGAGTGATCGCTTGTAAACTAATACTTGATAATTTCATAGA

           6001                                                                                                                                                                                                  6200
PflT_29_v1  TTGTAAATTAAGAACAGAAAATATGTTTGAAGCCATGGACACATTGTAAAAAGCATGACTGAGTTTTTGTGTATATTGTAAGCCTCCATACAGATAAAGAGAATCTCCAAACTTAATTACACTGTGACCAAAAGCATGAATTTCCTCACCACTATATTGTCCTTTAGACCATGTTAATGAAATTTTCCCCCTTGGATCAA
PflT_29_v2  TTGTAAATTAAGAACAGAAAATATGTTTGAAGCCATGGACACATTGTAAAAAGCATGACTGAGTTTTTGTGTATATTGTAAGCCTCCATACAGATAAAGAGAATCTCCAAACTTAATTACACTGTGACCAAAAGCATGAATTTCCTCACCACTATATTGTCCTTGAGACCATGTTAATGAAATTTTCCCCCTTGGATCAA

           6201                                                                                                                                                                                                  6400
PflT_29_v1  CAAAGACAGAAAGCTGCCACAATTTTGTATTGTGCATTTCTGAGTCAGTCAATCCACCAAGGAGAACATAATGGTTTGGTTCAACTAAACACAAATTTGAGCCATGTACATGTGGAGG-TAATTTGACTGAGAAGAATAATTTTTGATTTCTTGTATTGTATCATCTTTGTCTATTACAAGCCAGCCTGTTACTGTTGTC
PflT_29_v2  CAAAGACAGAAAGCTGCCACAATTTTGTATTGTGCATTTCTGAGTCAGTCAATCCACCAAGGAGAACATAATGGTTTGGTTCAACTAAACACAAATTTGAGCCATGTACATGTGGAGGGTAATTTGACTGAGATGAATAATTTTTGATTTCTTGTATTGTATCATCTTTGTCTATTACAAGCCAGCCTGTTACTGTTGTC

           6401                                                                                                                                                                                                  6600
PflT_29_v1  TTAGACTTGACCCAGACCAAAACTCTGTCACCTAAAGAAATAACTGGATTTGTACATCCAGCAGTTGCTAGGCAAGTTGCCAGGATGTTTTTGGTGATTTGTATTTCTGAAATAAAAAAGATGATGACAAAAGTTGTTGTATTAGAGAGTGAAGCTTATCTGCATGCATGTATGGTGAGCAGGTCATTTTATTTTGTCAG
PflT_29_v2  TTAGACTTGACCCAGACCAAAACTCTGTCACCTAAAGAAATAACTGGATTTGTACATCCAGCAGTTGCTAGGCAAGTTGCCAGGATGTTTTTGGTGATTTGTATTTCTGAAATAAAAAAGATGATGACAAAAGTTGTTGTATTAGAGAGTGAAGCTTATCTGCATGCATGTATGGTGAGCAGGTCATTTTATTTTGTCAG

           6601                                                                                                                                                                                                  6800
PflT_29_v1  TGTTGAAATATGGCACTGAAATATAAATTTGTTAATTTCATATTTTAGTATATGGATATCTTTGTAGTTCAAAAAAATGCTCAAGACAGAAAGTGAGATGTCTTTAGTCGTGACATTTATTAATATTGTTCACTGTACATTTCTCTACCCTTTGCAACCATACAATTGTTGAACACCTATCCTGTATATTCAAACAAACA
PflT_29_v2  TGTTGAAATATGGCACTGAAATATAAATTTGTTAATTTCATATTTTAGTATATGGATATCTTTGTAGTTCAAAAAAATGCTCAAGACAGAAAGTGAGATGTCTTTAGTCGTGACATTTATTAATATTGTTCACTGTACATTTCTCTACCCTTTGCAACCATACAATTGTTGAACACCTATCCTGTATATTCAAACAAACA

           6801                                                                                                                                                                                                  7000
PflT_29_v1  ACTTAAAGGCAAATCTGATATCTTTGGGGCAGCAAAATACATGTTCTCTTGTCTTTTGCATTTGAGTAACCTCCTTATATAAATATGATCGAAAGAGATTGTCAACATGGAGCAGCAAATCCGTTTCTTTAACCTAGACTGTAAATCAATACAAAATGGGTGCACATGTAGCTAGGGGAAAAGGTAATGCACAGATAGAG
PflT_29_v2  ACTTAAAGGCAAATCTGATATCTTTGGGGCAGCAAAATACATGTTCTCTTGTCTTTTGCATTTGAGTAACCTCCTTATATAAATATGATCGAAAGAGATTGTCAACATGGAGCAGCAAATCCGTTTCTTTAACCTAGACTGTAAATCAATACAAAATGGGTGCACATGTAGCTAGGGGAAAAGGTAATGCACAGATAGAG

           7001                                                                                                                                                                                                  7200
PflT_29_v1  CCCCATTGATGACTGCTGGAAATGTCATTTGGATGTCTTGCTGGTGTACATGCAACTGAAAAAAATAATTTTAGCCTGTGGGCCGTACAATAGCTGCACTTCTACTTCGCTCACTGTAATTTGAATCTTCGTATATAGTGTACATGGGGATCGACTTTCATAGTATGCTTGTTGGATCGAGGGACCGTGGCTTGTGTTAC
PflT_29_v2  CCCCATTGATGACTGCTGGAAATGTCATTTGGATGTCTTGCTGGTGTACATGCAACTGAAAAAAATCATTTTAGCCTGTGGGCCGTACAATAGCTGCACTTCTACTTCGCTCACTGTAATTTGAATCTTCGTATATAGTGTACATGGGGATCGACTTTCATAGTATGCTTGTTGGATCGAGGGACCGTGGCTTGTGTTAC
```

```
            7201                                                                                                                                                                                                    7400
PflT_29_v1  AGTTGAGAACGATCACATCGAGAATTTTCTTTTTCGAATGGGCGATTACAAGTGCAAATTTTTGGAAGTTAGCTACCATATAAGAACTTTGTTGTTTTAACTTCAAGAATAATCAATCTAAAACACGAACAGAATGTTCTGTGACTTTTGAAGCTACATTGAACTGACGACCTACCGGGACTATCGAGCATGGCCGCTGC
PflT_29_v2  AGTTGAGAACGATCACATCGAGAATTTTCTTTTTCGAATGGGCGATTACAAGTGCAAATTTTTGGAAGTTAGCTACCATATAAGAACTTTGTTGTTTTAACTTCAAGAATAATCAATCTAAAACACGAACAGAATGTTCTGTGACTTTTGAAGCTACATTGAACTGACGACCTACCGGGACTATCGAGCATGGCGGCTGC

            7401                                                                                                                                                                                                    7600
PflT_29_v1  CATCTTGGATATTGTTTACGACTCACTGAGCATGCTCAGATCAAAAATATTTCCCACAACCCATTGCGGACTTTCTACATTGTAA-CATAGGAGATGTTACGGTGTG------CGACGACTTGTCGTTA-GATTTCTTGCCTCCATCATCAACTTTTTGTTCAGTTCT------TCTCTTGTCATGTCATGGGTAACATG
PflT_29_v2  CATCTTGGATATTGTTTACGGCTCACTGAGCATGCTCAGATCAAAAATATTTCCCACAACCCATTGCGGACTTTCTACATTGTAAACATAGGAGATGTTACGGTGTGAACCTGCGACGACTTGTCGTTAAGATTTCTTGCCTCCATCATCAACTTTTTGTTCAGTTCTCGCTCTTATCTTGTCATGTCATGGGTAACATA

            7601                                                                                                                                                                                                    7800
PflT_29_v1  AATTTTTTGAAGAGACTATGTGAAGGGACCGTCCCAAAGTTTTGGAAGACATTTCATGAAATGCTTTTTTGCAGCAACATCAGTCATGCTGACCTTAATTGGTCTGGGTCTGACAGAGTCTGCTTCAGATGAATTATTGATTTTTCCTATTCGGACAACATTCTGTATATGTGTACAATCTGCTTCAATAACATCAGCTA
PflT_29_v2  AATTTTTTGAAGAGACTATGTGAAGGGACCGTCCCAAAGTTTTGGAAGACATTTCATGAAATGCTTTTTTGCAGCAACATCAGTCATGCTGACCTTAATTGGTCTGGGTCTGACAGAGTCTGCTTCAGATGAATTATTGATTTTTCCTATTCGGACAACATTCTGTATATGTGTACAATCTGCTTCAATAACATCAGCTA

            7801                                                                                                                  7920
PflT_29_v1  ATTCACGAAGTACATTAGTATCCGCCATTTTTCTTCCAGTTGGGTCAGGTTTATCTGCTCAGGAACACCATAAACAATCATATTCTTCTCTCTGTCGTTACTCTCTCTCTCAGTTCCTGT
PflT_29_v2  ATTCACGAAGTACATTAGTATCCGCCATTTTTCTTCCAGTTGGGTCAGGTTTATCTGCTCAGGAACACCATAAACAATCATATTCTTCTCTCTGTCGTTACTCTCTCTCTCAGTTCCTGT
```

#### FASTA DNA

>PflTRAGL-A_29_v1
TCTAGGGGACTAACGGTAGACAAGTTCCTGCAAAACAAAAGATGGCCTGTGGCAACGAGAATGGCCTACACAACAGGACATCTCCAGAGCGCTATGTAATAAAGACCCAGAAGTCAAGCGGGAAGCATCCGTGTACAGCACCATACTAGTAGAGCAAGAATTTGGGGTAGAGAAGATCTTACTACGCTACTCCAGCATGATGAAACTAAAGAAGATAGT
CGGATGGTTCCTGCTAGCGAAGAAAAACCTTCAAGATAAATGCCGCAAAAGAAAACAAGGCGATGAAGACACCAACGCAGATACTAGCCAGAAGAGCTGTGAAACAACAATCCCACCTGTTTCACTAGAGAATCTGCAAAGGGCAGAAGATGCAATCATGAGTATGTGCAGCGAAAGCATTTTCCAGATAAAATGGCACTCCTACAGCGAAAGAAGCAA
GTAAGGAGATACAGCCCATTATATAAGTTGGACCCAATTTTGGATAAAGGAATAATACGAGTTGGCGGACGCCTGGAACGTGCCTCAGTGTCACTAGACAGCAAGCACCCGATAGTGCTGCCTAAAGACTCACCAGTGTCAACAATCATACTCCAAGAAACCCACAAAGAAGTCGGACATCTTGGAAGAAACTCGATGCTGGCGAAGCTAAGAGAGAAA
TATTGGATTCTACGGGCACCTACAGCGATCCGCAACCTCGTAGCCAAATGTGTCATCTGTAGAAAGTGTAGAGCGAAAGTTTGTGAGCAGAAACTGTCAGGCCTACCTAAGGACTGCATCACCCCTGAAGAGCCGCCATTTACTCGCACAGATGTAGATTACTTCGGACCACTGGAGTCAAACGTGGCCGTGTCACTGAGAAGAGGTTTGGAGCGATCT
TCACATGTATGGCAACTCGTGCAGTACACTTAGAAGTAGCCTGCTCATTAGACACAGATTCGTGCATCAGAAGATTTATAGCACGAAGAGGACAAGTTAAGTCCATCAGATCAGACAACGGAACTAACTTGTCACTGCGGGGCCAAAGGCCCTCCATTTTGTTTCATATTAGCGTGTCACTTTTTCGCCGCACATACGTCGTGTACGCGTACAGTGAGT
CAAATCCGCCTTTGACGCTATGTATTAAATTTCGCAAAATTTCCACGCTGCACAGCTTAATATTGATGGAAGAGTTGCAGATTCAATTTCCTCATATGTTGACCTTTCCATTATGATATTTTGGCAATAATTTCATGAAGGAATTATGAAAATACATATGTGATAATTATTCATATGCAAAAATAGGTGAAAACACAGTCAACAGCCAAACGATCGTTA
ACTCGTCATTTCATAAAAATAAGAGGGAAAATCAAAAAAGTAAAACGATAGAGTTTTAGATAACCTTCATGAGATGCGCTATTTTAAATATCGTAAGAAATTACAGCGAAATGTAAAAGTTAAACCAATGGTTTTTCAGATATATATTTCATTTTACGACGCGGCGAATAAGCTTGATTTCCCGTAGAAGACAATGGCGTGTAATATACCGAACAGAT
TGCACACTTCTCGTCGGTTCAAATTCCTCAATTTTGAACAAATAATGATGAAAATTGACATGGTGATCCTTTGATTAATATACAAAATTCCTATCGTTTTAGATTTTTGAATTTCTTCACGAAAATTGTTTTATTTGAATTTTATTTATTCCGATCACTGAAATATACTGTTGCCGTCAATGATTTGGCGAGCTGACGGTGTGTACATATGCAGAACAC
GGGCCTTAAATTGACGCATTTTTATCAGTATGCGTCTCGTAAGATGTACAGACCAAATGCATGTCATGTTGTTGTAATTTATACCAAATTTGGCGAATATCAACGATGAAAATTCCAGTAAATTTGAAAAAATAACACAGAACGGAGGCGAACGGGCGCAGACTCTCCACTGTAAACGTGCAGTGAACCTTATCGATCGATATTCTTTTATTGGAAATT
GATACATTGTTGTCGTGCTCACTCGCAGGTCATATTCAGCTCACAATGGCGCCAAACTTGCAAAGAGCGCCAAAATGTGAACGCATCGCCAAAACTGTAAAAGGTGTCGACGATTCAATCTAAGTACAAGAAACCAACGGATACGTTGATTTTATGCTATCTAAGTCAATATATACGGGTTATGTGGACATATTAATGATTTGACCTCGACACATGAG
CCCGTACGCCAGCAGATTTCGATCGATATTCTATTGAAAATAAATTGATACAATGTTCGCTGTCGTCACTAGCAACCGGCTGAAAGCGCTCTCAGGTAGAATATATGCTTCGCGGAATACGAAACATTCAGGCATACGTTCGTCCCGCAGTGTGTCGCAGCCAACAAAGAGCTGAAGGAGAGTATCAGGAGATGGAATCAAGAGAAGATAGCTGAGACG
CTCCACCAGCAAGACATCAAATGGGAATTCAACCCCCCAGCAGGTTCCCACTTTGGAGGGAGTATGGGAGCGTCACATCAGAACCATAAGGAAAATCATTTTTTCCTTACTCAAAGAACAACCACTTCGCATGTCAGATGAAGCACTCCGAACGCTGTTCTGTGAAGTGGAAGCCATAATTAACAGCCGGCCAATCACCAGGCTGTCAGAAGATCCGAA
TGATCTAGAGGCATTAACACCCAACAACCTGTTAATGCTGAAACCAAAACAGGCCCTACCACCTGGCTTGTTTCAGAAGAATGACAGCTACGTACGCCGTCGATGGAGACAAGTGCAGTACCTGGCAGATATATTCTGGAAGAGATGGATGAAAGAATATCTCCCGTTGCTACAAGAGAGACAAGAAGTGGCTCGAGATCAGAAGAAACCTCAACGTTG
GAGACGTGGTGTTGATCGTCGACAACACAGCGCCGAGGAACTCCTGGTCCATTGGCAGAATCACCAAAGTGATGGAAGACAAGAAGGGACTCGTCAGGTTTGCAGAAGTCAAGACCAGAACGACTACTCTACGACGTCCGGTGGATAAGTTGTGCGTGCTCCTGGAGGCCCATGCCATTGAAGATTCGTCTGAAGGACACAAGAACAGCTAAGAACTTT
AATCTTTTTTATTAGTGATATATGAACAGTATATTATTCATGTTTGTTTAGTTAGATTGAATTTTGACCAGTTTAGAATATTGGATTTATAATTGTTCCGAGTCACATCTACACAATTAGGGGACCGGAATGTTACGGTCAAAATCAACATCCCCTTATTTGGCCATAGTTCAAATTCAAAGTCATGAGTCACCATCAACTTCCGTCCTTCAATGTTTA
CCTTTTGTATTACGCATTGCGCACATGTACCCATTCCAAGCATGCTTCTAGACACTTTAAGTTAGACCCTCAGAGAGTGGACATGTCTCTGAAAGCGACCGGCTATAGTTCTCGCCAAGATTGCTTGTAAAGACCATTTTGATGAAACAGCTTTGAACCTGTTGATGCGATAGTCTTTACAAACTCCACACAAAATCGGTTAGCCATTCAGTTGATTCT
TTTGTGACTTTAAAGCAGACAGTCTTGAAGGAACATTGAATAGCCTTATTTCATTGTGAGACTCTAAGAGCACGTGCGCCTAACAAAGGCCAAGGACTCATTGAAGTATGCGAATAATGGACACATTCAACGGACTCTGATTACGTTTATAGTACAACTTTTTGAGACCTTTTAGTTAAGTAGTTTTAGTGATTTTATTCTTATGCAACTTTGAGTTA
GTGATTGTTTCAAAGCAGCTAATTGTTAAGTCAAGGCTGCTTTTATGCAAATGAGATCTTGTCAACGAAAGTAAACGTTCAGTCACCGTTAGAGGGCGTTTACTGAGGTGGGTCACGTGACCAGTCCACCTATTATGACGTTGTGACCCCTGAGTCATCTTAGAAGGGTATCACTTCCTGTTAGTTATTTCCTGTTTCTAATTTCCTGTGTCATAGAGT
GTATTTACTTCGTGGTCAAAGAAAGTATTTACTTCCTGTGTCATAGTGTGAACGTAGCTTAGTTCTCACTTCCTGTATCCTTTTGTTTGATTTGTTTTCAATTAATCCTTCCCAAACGTTCTCAACGTTTGTAGTACAACATTCTACTCAGAGCATTTGTTGCTCTCAGTCTGGAAGACATGCTTATATTCGAATAAACCACGTGAAACTTAGATCGGA
TATCTGCCTCGTCTTATTTAAGAGGGAACGCATTACAGAATATCAGCTGTAGACGACTATTGCATAGTAGTTGCAGACAAATAATTTTGAGTCTTAAGGAATTTGTTTGCGCGCTACATCCCTCATCAAGAATACTGGTCCAGTGATCCAAAACTTTGAGTAGCTGTAATGTGGTCGATCTGCCACTTATGAACCATACTGTTTGTTACTGAATAAGTC
ATGTTTGTCCGTGTGGTACAATTTTTCTCCGAATAAGTTTTCCATAGTCTTACAGATAATACTGGTCAAGCTGATGGGTCTGTAGTTGGACGGAGATTTTTTGTTTCCCTTCTTAAAAATCGCTGAAATCTGTGCTTTTTTCATGCTTTTGGTACAACACCAGAGAAGAATGATTTGTTGAAGATAATAGAGAGTGGCGTATGGATAATGTCGGCTACT
TCTTTCAAGATTTTAGGATGAACTCCATCAGGGCCAGATGATTTCGTCACATCAAGTTGTAATAGGAGATTCCTAACTTCTGATTTAGTAAAAGGACTGCCACTGGACATGACATCAACATGACTACCGTTGAGATCTGGAATATTAGAGTTACTATCATCTTCTGAAGTGAAGACACTGCTGAAGAAGTCCGCTAAAAACTGCTTTCCCCTCATCACT
ACTTGCAGTAAGCAGGTGACCATCATCAATAATCTGGAGATCAGAAATTCCAGATCTGACTTTCCTCTTTTAATTAACATATTGCCAAAAGTTCTTACAGTTAGTTTTAATAGATTCCGCTATTCTTTTTTCTCGTGCTCTCCTGTCATTCTTGACAGTGTTTTTGAGTTATTTCTAGACTTACAGTATTCACGGTATTTAACAAGATCCTTAGATTCC
ATATATCTTTTCAAAAGTCTGTGTTTCCTCTTTAAACACCTCACTGTTACTTGTTTTAGTGGAGGCTGATCATTATAAGTATCATGTCGGAATGTTTTCACAACATACCTTTTCATTGCAATGTTCCACATTTCCATGAAGAAGTATATGCTTCTTCAGTAGTTTTGTTCAGCAGTGTGTGCTCCCAGTCTATAGACAGCTCATTTTGAATTCCCAACA
GATCAGCTTTGGAGTAGTTGTAACGTGTGGTAGAAGTGGAGACAATTTTGGCGTAACATTTGACATCAAGGCACAGCACACAGTGGATACTGGATCCAAGAGAGGACTGATCATGCACAATATTAGTGATGCAGCTGTCGTCATTAACAATTACTAAGTCTAAAATGGATGGCTCTTGGTTAATTCTGGTCCTAGTGGGCTTAGATATCGTCTGATAAA
CATAACAATCCCGAAAATACTCAAGAAACATATCACTGCTCTTGCATTTAGCAGTCGAAGACGACCAGTCAATATCTGGGTAATTAAAATCCCCTGTAATAATTAGATGAGTGTAATTGGTGTTGTCAATATAGACTGACTTAAACATATCACAAAGGTCGCGACGATTGTTCTCCGAAGATGAAGGACTTCTGTAAACACAGCAAAATAGGAGATTTT
CACAATCATGTGAACCAACTTTAATCCACACACATTCCTCAAAGTCTGTGTGGATTTTCGCAAGTTGTGCGTTCAATTTCTTTTTGTCATAAATCACAATTCCACGTCTAGCTTTGCTGGAAAAGTCATTTGAAATAAGTCATAATCTTCAATGCTAAGTTCAGATGTGTTGACCGTGTAGCGCATATTCTTGGGTAATACTTCACTGATGGCAATAAT
ATCAGGTGCTGCTCCTTCTGCATTGATAATGTAGTGTTGTAAGAGCTCATCAAATTTATTTGGTAAACTGTCAGCATTAGTATATATACAATATACGTTATGTCCACGAAAACTATGTACATTCCTATTTACAATATTTTTCCTCTCGTTAGATTGTATCCTTTCCTTTCACTTTGATACTTTTCCTTTCCAAGGTGGTCCTCTCACCAGGAACGGAAG
TCCCCGACGCACAGCGTAACATGTCAATATAGAAATCGATTAACACTACGATGAGTCGCAGAGAGTATCGACCAATCAGAGAGCAGCATTCATCAAGCCACGAACTTTCACGTATGGCTTCGAGATTTCGCGTGATCTAGGGTGTTTACTTCAGTAACTCTACAATCTACTAAATACTAAGGGGTCTCAACTGTTCTACACATGCGCAAACGCCATCTT
GAAGAGTGGCTGATGATCATGTCGAAAACCGGAGGTCGAAAGACGTAAGTATCGCCGAGATATCGTAATCTGAATCTCACGTTGCAATTTTTCACTCGATTATGTGTATCAAAATTATGCGGATACAATTTTTGGATTTTCTGTCAAACTTGGTTTTTCCTGTTGAAAGTTTGGGTTTTTCTCAGTCCGTAACGACATTGTTTACCGAAGGAACATGTG
CTGTACGCTTGGACTTTCTTCAAATGGCTTGGACTTTTAGATCTATACCCATCCAACATGTGTGATTGAGTGTAATGGCCGTTATATTTAAAATGTTCCTGTGATGGTAGCTAAGTCGGGGAAATGATCGTTTTCTGTTTAATATATTTTTTAAGGTTTGAATCAGCGTGTTACGATATTTAGAATTCCGAGCAGCATCCTAGAGGGTCAATTTGGGTG
TACAACGCATGGTTCATGCGGTAGCAAGCTAACCCAGGCCATATGGCACCAGGTACAGGTTGACTAACGAAGTGTTTTGAGGTTAGCATAGTATGAGTACGTCAGATTCATATGTCATCGGCATAGAAGGTCGCTTGTGTAAGATAACACCATATTTTTAGGCTGTCATTCATTTCATTTATTATTTTGCAGTGGGTTGCAGTCAGCAATGAATCATTT
ACAAGTTTGTTGCACTGTTATGCAATGCAATGCCTTGACCTGATTCAGTATTTCTTGCTTTGGTTCCTGCGGTATGAAGTAGAAAATTAGGCTGGCCACGGTGTCCTTGGAATTAATGTTTTACCACCAAGGTAAGAATAAAATTCTATTGGATTTATTGTTTACACATATATAAGGATGAAAGGTACAGACCATTGAACTGATTGGTACTGTACTTGC
CAAATTGATATGCTTAAATAATATTTTAAATATGTTAATTGTTTTCAGAGCGGA*ATGACGTCAGCTTTTCAACACAGAGAGTACCTTAAGAGGGTGTGTCGTGTATGTGGTAGAACCCTGAAGACCACATCTTACAGCAACTCAGGACATTACAGACAGGGAGAAGTGTGCATCCCGTACAGTGAACTTTTAGAAGAGTGCTTTAAAGTAAATGTTCAA*
*AATGATGATCCTTTTGTACATCCGCCTAGATTTGGTTCATGTTGTCGTAGAGTTCTTTTAAGTATTCTGTTGCACAGAACCTGGGTAAACATTACTATACCCATATCAGTGTTTGTAACTGGAAACATCATTGTAAGTATGAAAACCAAACTTGTCCTCTTTGCAAACGTTTTTCTCAAAATGGTTCTGGTAGACCTAAAAAACTTAAAGGGGTCGCT*
*TAAATTAAATACAAATGAAAATAATGAAACTGACAAGAGTCCTGTTGCAGACAATTTTGCTCATATACTAAATGACCACTGTTATACCACCCCAGCTTCCTGTAAGTTTGCCCAGTCTGTGCAATCAGACCAAAAGACAGATACATATGAATTTGTACAGAGGAAAATATTCAGACTGTACCCATGACAAGTCGATTTCCAACAGATTTATGTGTAACT*
*GAACCACATCCAGAATTCATTTGTAGAATGTGTTTAGGAGTCCTAGACACACCACTTCTAACACCGTGTACTCACACTTTCTGCGCAGGCTGCATTAAGAACTGGTTAAGTCTGTGTCATTTTTGTCCTCTCTGTAAACAACCTATGGTGTCATCAGATTTAAAAGAACCATATAGAGGATTTGTAGAAATTCTTAATGATACTAGAATGAAATGTAAT*
*TGTCCAGAAGCATTTTCAAGCAAAAAAGGTGTCAAGCTTAGTGATTGGCTTCAGCACACTTTAAATTGTGGAAACGATTTTGAAATTGATGAATGGTTAGAAAAAGTTTTGTCATTGTGGAACGAAATTTCTCTCACTCAAGTTGATTCACATTTAAATAGTACTGAACATTTGCCTGAGACAGATAAGTCTAGTCTGACAACAAGCCATGGCCATCCA*
*GACATTGCTGCAGGTAGCACTGTGGAAGACCTAAAACTTCACTGTTGTGTCTCCAACAACAAGGACAAAATTTAAGATTGAAAAATATCAAAAAACAAATAGACGTGTTTGCCAGAGAGAAAGGGGAAGATCTCAGGGTAGTGTATTTTGGCCTTTTGCAAAAACTATCTGAGAGCCAACAAGCAAGGTAGTTTAGCTGAACAAGTGAAGTTGCTATAT*
*AATGGGGAATCTTGTCAAATGTCACCTCAGGAATGTCTAGCCTTGCGTGTAAATACTTTACCAGTTGTAATCAGTACAAGAAAATTTACTCGGCTACAAATGAGCTTAGTGGAGTTAAAATCTTCCAACCTCTCATCAATGTTGTGAAAGCTGAAGAAAGTTTCTACCAGGGTCTGTAAAGTTCTACTTAACCCATCTTTGAAAACTGTTTCTCCCAT*
*TTGGCAAGGTACAGAGCATGTTGATGCTCTGTCAGGACTCAGTCGCGATATACCACTTCTAGAGTCACCTGATGTTTATGGGGCAAGGTATAGATATGACAAAGCAGTTGCTATGGCTTTGAAAGATATTGATAATGAAATCTCAGAAGGCCTCAGTAAATTAGAAGGGGTGGATACAAATCAACTTGTTCTAACAGCCACTGTCAAAGATGGCAGTGA*
*TGGTCTTGGCGATGTGAAAGAAGTCAGATCTCAGGGGCTTGTCGTTCCAAACAAGGCCTTACGGTTTTCATTTGTAATTTTAGAAGTCACTGTCAATGTAGATGGCAAGATTATTACTGTATTTGAAGAAGATGCTCCCAACTCAGAATTATGCACACGCCCTCTCTTGGTAGCTCTGGCAGATGAAAATGACAGAAAAGGGTTATGCATGACACATAT*
*ACCAATAATGATAGAGAGAGAGGAGATTGAGAAAAGTACCTTGTTTGTAGAAATTGAAGAGGGAAAACAAAGAATCTACAAAGTTGTATTAAAGGGACTATGTATGATGAAAAATTGCAGAGAAGTGGTGAGGGAATGTTAGCGGCAGGGTCATCTTGGTTTTGTACACTTTGTGAGAAAGGAAGACTTGATCCCATATCATCAGTCTTGCCAATAACA*
*AGAACACATGAAAACAATATGGCTTTGTATTCAAAGTATGTTGAAAATGCAAATCATATGAAGCAGGCAGATCTAGTGAAAGAAGTCAAAGGCATTGTTAGAAAACCATTTATAATGTCAGAGCCTTGTGTTGATGCCACGCATGCTGAAATTCACTGGGGAGAGAAAAACTATGATATGTATGTACGTGAAATATCAGGTATAATGTGTTGGGGAAG*
*CCATCCAATGATGATGATGCAAAAAAAGTGAAATCAACGAAAAGAATGTTGGATACCGAATTACAGCTGCAATGTGGCTTAAAGAGAGAGCATGGCTTTATGGTTGATGGAAATTATGCACGTGATTTGGTCAAATCAGAAACAGTTGATGTGGTTTGCTCTTTAATTCAGAGTGAAGCAAGAAAATCTGTTGTTCGTGAATATGTTTCCTACTATCGA*
*GAATTGCGATCAATATACAGGAAACACAATGCTGACTTAGATGATGCCGTGAAATTTAAAACCATAGCTGCAAATTTTACAAATTGCTACATGAGAAATTTCCTATTTGCATCCACTCTCTAACTATCAACACAAGGTATTAGATCACATTCCACAGTTGATCGAGCAATATGGCTCTGTTGGTAAATTTGCCTCTGAAGGCAATGAAAAAGGCAATAA*
*ACTTTTCAGAAGATTTCGTAAATTTAATGCACGTAAGTCTCAAATTCATGAAATGCCAGACATTCTGAAATTTCATTGGCTATATACAATGAAAAGTCTTCAGATTATGGTAAACTGGGAAAAAACTAGATCACTAGTCTGTAGCAAATGTGGTTGTGCAGGTCACAATCAAAGGACATGCATTACATAA*TTGTGCCCAAGGCCATGATATGAAATCAA
AGATTAGATAATGACCATGCATGTTTTATGTATCTGAATCTGTAAGTTTGAAATTAAGTAGCATCTGTATGTATCAGACTACAGAGTCCAAACTTCAGACACAAAGAGTAGGCTGGCCATACACTCTTATTTACTTGAAAAGTGTTTCTGTTCCATATACTGACTGTACTGTCAGGAGTCAATTCTAGCTTCTATGCTTCTATACCTAGATAGTTAAAG
TGTATTATATTTTGATGTACATAGATGAATTCCACAAGAAAACATCAAATATAAGAAAGCTGAGTTTGTGTTAAATATTTGTATGTGTTTTAATCTTACCTGCCCACACTTTTGAAGGGACTCAAACAAAATAATTTGATTACTGTGGCCAAAATGTCAAAAGTTTGAGTAACTGGCCATGAATATGAGATGAGAAGCTTAAGGTTTGCAATAGAATTT
TGTACATATATTCATGAAAGATTGTAACAAGTTGTAAAATGAACTCTCAGTAAAGGTTTCATTCAATTCATGCACATGCTACATGTACATGCTATGATTTCTTGTGCAGTTGTCATGTTCTCA*GAAATTACAATCAGAACATGTAAATTTGATGTTACTCTTATTTTCCAGTTGTCGTGAAGAAAATCGTTTGCCAGAGCATTTAATGTGAAAAGCTCT*
*AACACATTGTGTACATCGCACTGTTGAAAAGTAACTCTTATCAGTCACGCTACAATTAATGTTAGCACATGTTGTGAAACATGAAGGGCATTCAAAGTTTATACAGTCCAGCTTTGTTCTATTTAAGTGATCTAATTCAACACATTTGTCATGGTACCACCTTCCACAGAATACAGTTTGATTGTTCATGATGTAAGTTCGCTCACATTTCAGCCAATT*
*GAACTCTGCTTTATCACTTGCTGGTAAACTAGCAAACTTGCAGCCATCCAACATACATTCCCATGTTTCATTCGTCACTGTCATCTGATGTATTGTCTATCTTTCCTGCTCTGCTGCCCTCTTCTGACTCATTCCTGTCTGTGCTAACTTTGACCTCAGCTGCCTCACTTATTCCGTTCTGATCCTCAGTCTCTGAGCCTATACTTTCCTGATCTGTCA*
*TGTCTATGCTTATCCTTTCCTGCTCTTCCTCCGCTATGCTTACTCTGTCCTCATCTGTTGACCTTTCCTCCTCATCCCCCAACTGACTGTTATTACTCTGTGTCCTGGTTTCAGAGCTTGTCCCTTCATTTTGAAAACTGAATTTTATCATTTGCTTTTTTACTGTATCACAAACAGCAATTGAGCTGAAGAATGGAAGACTGTAGTATTACTACTTAA*
*GGTATCATAGTGTGCTGTTTCTATGAAATCTATGGCATCACAAGGTGACTTGTAAACTGTGTTGTTACTCAATATGTAAACAGTGTTACTACATTTGAATGAGACTGGGTTAGGAATACCAATAAATGGTACAGATGTACAGACCATTGAATTCAGGTCAATTACATCAACATCGGGTGGATGTCTTACATTGTTACCTATCACATAAATTCCACCAAC*
*AATATACATTTTACTGCTATTACGAAAAACTGAATGGTAGGCTCTTGGAGTGATCGCTTGTAAACTAATACTTGATAATTTCATAGATTGTAAATTAAGAACAGAAAATATGTTTGAAGCCATGGACACATTGTAAAAAGCATGACTGAGTTTTTGTGTATATTGTAAGCCTCCATACAGATAAAGAGAATCTCCAAACTTAATTACACTGTGACCAAA*
*AGCATGAATTTCCTCACCACTATATTGTCCTTTAGACCATGTTAATGAAATTTTCCCCCTTGGATCAACAAAGACAGAAAGCTGCCACAATTTTGTATTGTGCATTTCTGAGTCAGTCAATCCACCAAGGAGAACATAATGGTTTGGTTCAACTAAACACAAATTTGAGCCATGTACATGTGGAGGTAATTTGACTGAGAAGAATAATTTTTGATTTCT*
*TGTATTGTATCATCTTTGTCTATTACAAGCCAGCCTGTTACTGTTGTCTTAGACTTGACCCAGACCAAAACTCTGTCACCTAAAGAAATAACTGGATTTGTACATCCAGCAGTTGCTAGGCAAGTTGCCAGGATGTTTTTGGTGATTTGTATTT*CTGAAATAAAAAAGATGATGACAAAAGTTGTTGTATTAGAGAGTGAAGCTTATCTGCATGCATGT
ATGGTGAGCAGGTCATTTTATTTTGTCAGTGTTGAAATATGGCACTGAAATATAAATTTGTTAATTTCATATTTTAGTATATGGATATCTTTGTAGTTCAAAAAAATGCTCAAGACAGAAAGTGAGATGTCTTTAGTCGTGACATTTATTAATATTGTTCACTGTACATTTCTCTACCCTTTGCAACCATACAATTGTTGAACACCTATCCTGTATATT

CAAACAAACAACTTAAAGGCAAATCTGATATCTTTGGGGCAGCAAAATACATGTTCTCTTGTCTTTTGCATTTGAGTAACCTCCTTATATAAATATGATCGAAAGAGATTGTCAACATGGAGCAGCAAATCCGTTTCTTTAACCTAGACTGTAAATCAATACAAAATGGGTGCACATGTAGCTAGGGGAAAAGGTAATGCACAGATAGAGCCCCATTGA
TGACTGCTGGAAATGTCATTTGGATGTCTTGCTGGTGTACATGCAACTGAAAAAAATAATTTTAGCCTGTGGGCCGTACAATAGCTGCACTTCTACTTCGCTCACTGTAATTTGAATCTTCGTATATAGTGTACATGGGGATCGACTTTCATAGTATGCTTGTTGGATCGAGGGACCGTGGCTTGTGTTACAGTTGAGAACGATCACATCGAGAATTTT
CTTTTTCGAATGGGCGATTACAAGTGCAAATTTTTGGAAGTTAGCTACCATATAAGAACTTTGTTGTTTTAACTTCAAGAATAATCAATCTAAAACACGAACAGAATGTTCTGTGACTTTTGAAGCTACATTGAACTGACGACCTACCGGGACTATCGAGCATGGCCGCTGCCATCTTGGATATTGTTTACGACTCACTGAGCATGCTCAGATCAAAAA
TATTTCCCACAACCCATTGCGGACTTTCTACATTGTAACATAGGAGATGTTACGGTGTGCGACGACTTGTCGTTAGATTTCTTGCCTCCATCATCAACTTTTTGTTCAGTTCTTCTCTTGTCATGTCATGGGTAACATGAATTTTTTGAAGAGACTATGTGAAGGGACCGTCCCAAAGTTTTGGAAGACATTTCATGAAATGCTTTTTTGCAGCAACAT
CAGTCATGCTGACCTTAATTGGTCTGGGTCTGACAGAGTCTGCTTCAGATGAATTATTGATTTTCCTATTCGGACAACATTCTGTATATGTGTACAATCTGCTTCAATAACATCAGCTAATTCACGAAGTACATTAGTATCCGCCATTTTTCTTCCAGTTGGGTCAGGTTTATCTGCTCAGGAACACCATAAACAATCATATTCTTCTCTCTGTCGTT
ACTCTCTCTCTCAGTTCCTGTACTTTCTTTTTGATAGACTCCTCAATGTTTCCTAATGATTTTGTAGTTATAAGTCCTTCAGCTTCAGTGTGCTCCTTAATGCTATCACTTGTCCTCGTCGACTTTTGACTTCATACTGTTTTCAAGAGTAGTAATTCTATCTTCCATTGATTTCAAAAAAGCTGTACTTGGCTATCAGTTCTTATGTTTTTCATCACC
TTCTCTTCGCAAGTGCAACAGAACCAATGAAGTGCAGGATTCTTGAACATCTCATAGTCATCTACTGACAGTTTCAAACATTCTCGGCAAAAGTACAGTATTCACACTGAACAATTTCTGCACTGTCGTCTGAAAATTCTTCTTTACAAACTTCACAAATCCATGCCTCCTGGTTCTGGCGCATATGACTCGGACTTGATATAGCTTTCTGCCTTTTGG
TAAGAGAGTTTGAAGTGGAATCTGTGTTTTATCACAGATGTGTTTTGTGATTACACCTAGCTATCTGCCTTCCTTACATCTTCACTGTCATCCCCTTAGCCAATCGGCTCTTCTACCCATGACTGTTATCAATCCTCCTTCCCAGCCAATCAGCCACGTTTGCACACGACACTACATAATCAAGCCTTTGTCTCATCGCTCCCATTCATTCATTCATTC
ATTCATTCATTCAGCCGCTGGGTTCATGCCATACCAGCTCAGCCTATTACACTATGTCCGGAATCGTTTACGATCCCTTCCTTCGAACAACCATGCCGGCCAAAGGACGTCCAAAGACAGCCAAGAAATCTGAGTGTCCGAATGCTCAGCCTGCTTTGAGACCCCGTGCTCAGCCCTACACAAGACCGGCCACTGACCTTTGTCCCGCTCCGCTGCCTG
TGCACGTTGGATCTAGCTGCGAATATGGAGCACGCCCGCGGTGTACTAACGTATTATTACTCACAATCCTTCTGATCGGCCTCGGGGAATGACAGTCCTGGCATTCAAACCGAGTTCAGAACCAGGTCTCCTCCTAAGTTCCATTGTCTGCAATGGTTCCTTCTGCCAGCATTTCGTCGGGCAGCGTCCCAAGCCAACATTCCAGAAGATAGTTATGTA
ACACTTCCATAAACATAGCTGCAAGACTCGGTCAACACGATATCCAATATCCTGCAACAGTAACTTCGACCAGCGGCCACTAAGTCACCTATGAACTCTCTGACGTCACTTCCCGAAGTTCCACTCCGGAACGTAAACAACACAATCCAAGATGGCGGCGCCCTACGTTCGAACGACGTCGATGTCATCCCCTCCTCATCGTCACGCGTCATCCAACAC
CTTGCTACAGACCTGCGGAACGCCTTCAGAAAGCCTACCACATATCGAAATGGTGTCTCCAGCGATTCGTCGCGATATCCTGGCAGGAAAGGATGGGAATCTTGCCTACTCATTCCCGGCTATGGTTCAAGCGACGGACTTCTGCTGCTTATCCACACCTTCAAGCCGACGACCTGCATATTTCCATGCGTACATGAATGAACATACATGCCACTGCCA
GGCAAAAGACGACGCTCACGAAGGCTCCATCGACATGGGCACGTCCTCGTGTTCAAACCATAAGTCACCGGACTCATTACTTTGCTATAAATATCTGGCATCTTCCAGTTCGTCAAGCGTTTGTACCAATTTGAAACGTACTACCGCATGTCGGAACAAGCTGCAGGCCGACTACTGCCCAAAGACGTGGGACTTTGGCACATCTTCGAGTTCAACCAT
AAAGTATCGGACTCTTTTATTACTCTGCCTATGTCCGACATTTCTAGCTCGTCTGTACGTTGGAAAGTTGCAGGCTGACTACGAGTAACCATAGACTGACTTTGCTTGGCCATATACCGCCCATGGCGATGTTTTCCTGGCCTATGTCGAGCCCGGTTTTTCTGTCACGTCCGACCGCTCAATCCGACCCATCGTACCAATCCAGCGCCAGCTCCGCAT
CTGCAAGAACTTCATAAGCATTATTCAGATGACTGCGGATCAACAGGACACAACAGCCAGACGCCGTCAAGAAGCGAACGTAACGACGGCCAGAGTGTATACCACATTGACTCCAGACTCGGTCGACCTGCCTGTTTTGGTGGCGGGGCCAGAAACCTCATCAAACCTGATGTCATCAGTTCCTATCACGTACATCGTATCACGTCCAGCAAGAATCGA
CAAGAGGATCAGTTGACTGACCGACGACTATATCAGTTTCCGTTTCAGATTAGTAATTAGTCGTGCGTAGCCCGTCATAAGACAGCTTGCTGCACAGGGAAAACAACGGACCTTGACCCTGTGACGTCAACAAACTCAGTCTTAGGACCAAAGATGGCGGCGCCCATGGTGTCAAGTGGCCCCATCGTCATCCCAACTCTAGCTTTGCTGGAAAGTCATTTGAA
GGCGACTATTTAAGTAACATCTGCATGATTTCAAGCATAATAACGAGTCCTTCCGACCGGCAGTTCACAACGATCTACACACGGCCTCACTTCCTGCTTGACTAGCATTTCCTGTTTGACTGAAAGCGAAACAAGGTACAGTTTCAATTCAAGACTCAATCGAGTCAAGACTCTGTTTACGCTGCTCAACCAACAATGACTAGCATTTCTCCGAGCATG
TAAATATGCGGCAGCTCAAGGCTAGCCCTACCTGGTGAACTTTCAGACTGTTAAGCGAAGAGTTCGATGCGTATCAACATGACACTCCTGCGTTTGTGGACAATCAAGTCTGACAAGCAAATAAATTCAGAGCATACATATGCGGAAGCCGAAGGCTAACTCGCTGTAAAGCATCCTATCGGTGGCCTATCGGTGGCCTACTTAACAGTCGTCTCACCA
ATCGGACGTAACCTGCTCCCAGCTACCTATTGGATTTTACAGACCAGTTTCACCTGTCAGTTAATGTGTCACCAAGTGCAGCTGCTGGATGTCACCCGATGATTTCTCCATACTTCTACCTTGGATGACGCTGCGGTTCCACAAGAATGGAACACAAACTACAGCTGGAGATCGAGAACGACCGGTCATTCTATATCGCCTTTCAGACTGATGTTTGC
CATCTCGTTGGCATGTTCTGTCTGCTGGGGTGGACTTGGCAATGAATCTTGCCTCCTTCTTTTTTTCTTATGCTACGATAAGGCCATGACTTACCTTTTACGTTGAAATTACTACTTATTAGTAGTTCAGTTTTGCTAGCATTCGCCTACTTGTCGCTCGTTTATTTACCTGTTTGCAGCTCATGTTACCTGGCTCTTCGGTTTGCATGCATTATCG
TGCCCCCCTTTTTTAGCATGTCGACTTGGCCATCCTGCACAAGCGTTTTCCCAGCTTTGCATGTATTGCGCAGTCTGCAGTCTGGCCTTACATGCGGCCCACGCAATTTGCTCTTCATACAGCGTACCGATGTTTTTCACGTTTCTTTTTCCTTTCATTTTAGATGCACCGTTTCAAACTACTGTGTCACTCCCTGGAAGCCGAACCCTGCAGCTGCAC
TTGTCGGCAATCTCACAAGCATCCCAGCGAAAACATACGAAAACAAAAACATTTGCTAGACCCATGGATACCTTACCACAAACCGCATTACGGCATCACGTCAGCACTTTGACTTTGTGGATTGTACAGTTCACATCAGTGTGACTGGAGTAACATGCTTTCTTTCAGAACACTTTCACTTTCAAAGCAAAGGACTACATACTTCATCCTTTGCCATTT
AGCAGTATTACAGTTTTTAGTAATGTCTCATAAATGCCACTTTTTTCCACATCAGTTTCCTCATTTCGAAAGAACTGGCATTTAACGCTGTCATGGACTTCACACTTCGTTTATGCAATGCTTTGCTTTGCAGCCAGCAACATGCCATTGTAGAGGCTTTGCTTCGCATGCTCGGACATTTTGCCAGTTAACAGCGACGTTCCGCTTGGCCTGATCCGTTCACTGCTCTACAGGGA
AATCCCAGACCACCTTTTAGGTAGATGGTCTAGCAACTGCCACCGACTTACATGTACATTAAAGAACATCTGAAATTTTCCCTGCGTCACATTTTTCAACGGCTAGCCGACCACCTTTTAGGTAGACGGTCTAGCAACTGCTACCGACTTTACATGTACATAAGAACGTCAGAAATTTCCCTGCGTCACACTTTTCAGCGACGTTCCACTTTGCCTGAT
CCGTTGACTGCTGCTCTACATGGAAATCCCGGACCACCTTTTAGGTAGACGGTCTAGCAACTGCTACCGACGTACATGTACATCAGAACATTAGAAATTTGC

>PflTRAGL-A_29_v2
TACTGGTCCAGTGATCCAAACTTGAGTAGCTGTAATGTGGTCGATCTGCCACTTATGAACCATACTGTTTGTTACTGAATAAGTCATGTTTGTCCGTGTGGTACAATTTTTCTCCGAATAAGTTTTCCATAGTCTTACAGATAATACTGGTCAAGCTGATGGGTCTGTAGTTGGACGGAGATTTTTTGTTTCCCTTCTTAAAAATCGCTGAAATCTG
TGCTTTTTCATGCTTTTGGTACAACACCAGAGAAGAATGATTTGTTGAAGATAATAGAAAGTGGCGTGTAAATAATGTCGGCTACTTCTTTCAAGATTTTAGGATGAACTCCATCAGGGCCAGATGATTTCGTCACATCAAGTTGTAATAGGAGATTCCTAACTTCTGATTTAGTAAAAGGACTGCCACTGGACATGACATCAACATGACTACCGTTG
AGATCTGGAATATTAGAGTTACTATCATCTTCTGAAGTGAAGACACTGCTGAAGAAGTCCGCTAAAAACTTGCTTTCTCCTCATCACTACTTGCAGTAAGCAGGTGACCATCAATAATCTGGAGCTCAGAAATTCCCGATCTGACTTTCCTCTTTGAATTAACATATTGCCAAAAGTTCTTACAGTTAGTTTTAATAGATTCCGCTATCTTTTTTCTC
GTGCTCTCCTGTCATTCTTGACAATGTTTTTGAGTTATTTCTAGACTTACAGTATTCACGGTATTTAACAAGATCCTTAGATTCAATATCTTTTCAAAAGTCTGTGTTTCTTTAACACCTCACTGTTACTTGTTTTAGTGGAGGCTGATCATTATAAGTATCATGTCGGAATGTTTTCACAACATACCTTTTCATTGCAATGTTCCACATTTC
CATGAAGAAGTATTTGCTTCTTCAGTAGTTTTGTCCAGCAGTATGTGCTCCCAGTCTATAGACAGCTCATTTTGAATTCCCAACAGATCAGCTTTGGAGTAGTTGTAACGTGTGGTAGAAGTGGAGACAATTTTGGTGTAACATTTGACATCAAGGCACAGCACACAGTGGTCACTGAATCCAAGAAAGGACTATCATGCACAATATTAGTGATGCAGC
TGTCGTCATTAACATTACTAAGTCTAAAATGGATGGCTCTTGGTTAATTCTGGTCCTAGTGGGCTTAGATATCGTCTGATAAACATAACAATCCCGAAAATACTCAAGAAACATATCACTGCTCTTGCATTTAGCAGTCGAAGACGACCAGTCAATATCTGGGTAATTAAAATCCCCTGTAATAATTAGATGAGTGTAATTGGTGTTGTCAATATAGA
CTGACTTAAACATATCACAAAGGTGGCGATGATTGTCCTCCGAAGATGAAGGACTTCTGTAAACACAGCAAAATAGGAGATTTTCACAATCATGTGAACCAACTTTAATCCACACACATTCCTCAAAGTCTGTGTGGATTTTCGCAAGTTGTGCGTTCAATTTCTTTTTGTCATAAATCACAATTCCACGTCTAGCTTTTGCTGGAAAAGTCATTTGAAA
TAAGTCATAATCTTCAATGCTAAGTTCAGATGTGTTGACCGTGTAGCGCATATTCTTGGGTAATACTTCACTGATGGCAATAATATCAGGTGCTGCTCCTTCTGCATTGATAATGTAGTGTTGTAAGAGCTCATCAAATTTATTTGGTAAACTGTCAGCATTAGTATATATACAATATACGTTATGTCCACGAAAACTATGTACATTCCTATTTACAAT
ATTTTTCCTCTCGTTAGATTGTATCCTTTCCTTTCACTTTGATACTTTTCCTTTCCAAGGTGGTCCTCTCACCAGGAACGGAAGTCCCCGATGCACAGCGTAACATGTCAATATAGAAATCGATTAACACTACGATGAGTCGCAGAGAGTATCGACCAATCAGACAGCAGCATTCATCAAGCCACGAACTTTCACGTATGGCTTCGAGATTTCGCGTGA
TCTAGGGTGTTTACTTCAGTAACTTCACAATCTACTAAATACTAAGGGGTCTCAACTGTTCTACACATGCGCAAACGCCATCTTGAAGAGTGGCTGATGATCATGTCGAAAACCGGAGGTCGAAAGACGTAAGTATCGCCGAGATATCTGAATCTGAATCTCACGTTGCAATTTTTCACTCGATTATGTGTATCAAAATTATCGGATACAATTTGG
ATTTCTGTCAAACTGGTTTTTCCTGTTGAAAGTTTGGGTTTTTCTCAGTCCGTAACGACATTGTTTACCGAAGGAACATGTGCTGTACGCTTGGACTTTCTTTCAAATGGCTTGGACTTTTAGATCTATACCCATCCAACATGTGTGATTGAGTGTAATGGCCGTTATATTTAAAAATGTTCCTGTGATGGTAGCTAAGTCGGGGAAATGATCGTTTT
CTGTTTAATGTATTTTTTAAGGTTTGAATCAGCGTGTTACGATATTTAGAATTCTGAGCAGCATCCTAGAGGGTCAATTTGGGTGTACAACGCATGGTTCATGCGGTAGCAAGCTAACCCAGGCCATATGGCACCAGGTACAGGTTGACTAACGAAGTGTTTTGAGGTTAGCATAGTATGAGTACGTCAGATTCATATGTCATCGGCATAGAAGGTCGC
TTGTGTAAGATAACACCATATTTTTAGGCTGTCATTCATTTCATTTATTTATTTTGCAGTGGGTTGCAGTCAGCAATGAATCATTTACAGTTTTGTTGCACTGTTATGCAATGCAATGCTTTGCTTTGGTTCCTCGCGGTATGAAGTAGAAAATTAGGCTGGCCACGGTGTCCTTGGAATTAATGTTTACCACCAAGGT
AAGAATAAAATTCTATTGGATTTATTGTTTACACATATATAAGGATGAAAGGTACAGACCATTGAACTGATTGGTACTGTACTTGCCAAATTGATATGCTTAAATAATATTTTAAATATGTTAATTGTTTCTAGAGCGGAATGACGTCAGCTTTTCAACACAGAGAGTACCTTAAGAGGGTGTGTCGTGTATGTGGTAGAACCCTGAAGACCACATCTT
ACAGCAACTCAGGACATTACAGACAGGGAGAAGTGTGCATCCCGTACAGTGAACTTTTAGAAGAGTGCTTTAAAGTAAATGTTCAAAATGATGATCCTTTTGTACATCCGCCTAGATTTGGTTCATGTTGTCGTAGAGTTCTTTTAAAGTATTCTGTTGCACAGAACCTGGGTAAACATTACTATATACCCATATCAGTGTTTGTAACTGGAAACATCATT
GTAAGTATGAAAACCAAACTTGTCCTCTTTGCAAACGTTTTTCTCAAAATGGTTCTGGTAGACCTAAAAAACTTAAAAGGGGTCGCTTAAATTAAATACAAATGAAAATAATGAAACTGACAAGAGTCCTGTTGCAGACAATTTTGCTCATATACTAAATGACCACTGTTATACCACCCCAGCTTCCTGTAAGTTTGCCCAGTCTGTGCAATCAGACCA
AAAGACAGATACATATGAATTTGTACAGAGGAAAATATTCAGACTGTACCCATGACAAGTCGATTTCCAACAGATTTATGTGTAACTGAACCACATCCAGAATTCATTTGTAGAATGTGTTTAGGAGTCCTAGACACACCACTTCTAACACCGTGTACTCACACTTTCTGCGCAGGCTGCATTAAGAACTGGTTAAGTCTGTGTCATTTTGTCCTCTC
TGTAAACAACCTATGGTGTCATCAGATTTAAAAGAACCATATAGAGGATTTGTAGAAATTCTTAATGATACTAGAATGAAATGTAATTGTCCAGAAGCATTTTCAAGCAAAAAAGGTGTCAAGCTTAGTGATTGGCTTCAGCACACTTTAAATTGTGGAAACGATTTTGAAATTGATGAATGGTTAGAAAAAGTTTTGTCATTGTGGAACGAAATTTCT
CTCACTCAAGTTGATTCACATTTAAATAGTACTGAACATTTGCCTGAGCAGATAAGTCCAGTCTGACAACAAGCCATGGCCATCAAGACATTGCTGCAGGTAGCACTGTGGAAGACCTAAAACTTCACTGTTGTGTCTCCAACAACAAGGACAAAATTAAGATTGAAAAATATCAAAAAACAAATAGACGTGTTTGCCAGAGAGAAAGGGGAAGATC
TCAGGGTAGTGTATTTTGGCCTTTTGCAAAAACTATCTGAGAGCCAACAAGCAAGGTAGTTTAGCTGAACAAGTGAAGTTGCTATATAATGGGGAATCTTGTCAAATGTCACCTCAGGAATGTCTAGCCTTGCGTGTAAATACTTTTACCAGTTGTAATCAGTACAAGAAAATTTACTCGGCTACAAATGAGCTTAGTGGAGTTAAAATCTTCCAACCT
CTCATCAATGTTGTGAAAGCTGAAGAAAGTTTTCTACCAGGGTCTGTAAAGTTCTACTTTAACCCATCTTTGAAAACTGTTTCTCCCATTTGGCAAGGTACAGAGCATGTTGATGCTCTGTCAGGACTCAGTCGCGATATACCACTTCTAGAGTCACCTGATGTTTATGGGGCAAGGTATAGATATGACAAAGCAGTTGCTATGGCTTTGAAAGATATT
GATAATGAAATCTCAGAAGGCTCAGTAAATTAGAAGGGGTGGATATACAAATCAACTTGTTCTAACAGCCACTGTCAAAGATGGCAGTGATGGTCTTGGCGATGTGAAAGAAGTCAGATCTCAGGGGCTTGTCGTTCCAAACAAGGCCTTACGGTTTTCATTTGTAATTTAGAAGTCACTGTCAATGATAGATGGCAAGATTATTACTGTATTTGAAGAA
AATGCTCCCAACTCAGAATTATGCACACGCCCTCTCTTGGTAGCTCTGGCAGATGAAAATGACAGAAAAGGGTTAATGCATGACACATATACCAATAATGATAGAGAGAGAGGAGATTGAGAAAAGTACCTTGTTTGTAGAAATTGAAGAGGGAAAACAAAGAATCTACAAAGTTGTATTTAAAGGGACTATGTATGATGAAAAATTGCAGAGAAGTGGT
GAGGGAATGTTAGCGGCAGGGTCATCTTGGTTTTGTACACTTTGTGAGAAAGGAAGACTTGATCCCATATCATCAGTCTTGCCAATAACAAGAACACATGAAAACAATATGGCTTTGTATTCAAAGTATGTTGAAAATGCAAATCATATGAAGCAGGCAGATCTAGTGAAAGAAGTCAAAGGCATTGTTAGAAAACCATTTATAATGTCAGAGCCTTGT
GTTGATGCCACGCATGCTGAAATCACTGGGGAGAGAAAAACTATGATATGTATGTACGTGAAATATCAGGTATAATGTGTTGGGGGAAGCCATCCAATGATGATGATGCAAAAAAAGTGAAATCAACGAAAAGAATGTTGGATACCGAATTACAGCTGAATGGCTTAAAGAGAGAGACGGCTTTATGGTTGATGGAAATTATGCACGTGATTG
GTCAAATCAGAAACAGTGATGTGGTTGCTCTTTAATTCAGAGTGAAGCAAGAAAATCTGTTGTTCGTGAATATGTTTCCTACTATCGAGAATTGCGATCAATATACAGGAAACACAATGCTGCTACTTAGATGATGCCGTGAAATTTAAAACCATAGCTGCAAATTTTTACAAATTGCTACATGAGAAATTTCCTTATTTGCATCCACTCTCTAACTAT
CAACACAAGGTATTAGATCACATTCCACAGTTGATCGAGCAATATGGCTCTGTTGGTAAATTTGCCTCTGAAGGCAATGAAAAAGGCAATAAACTTTTCAGAAGATTTCGTAAATTTAATGCACGTAAGTCTCAAATTCATGAAATGCCAGACATTCTGAAATTTCATTGGCTATATACAATGAAAAGTCTTCAGACTATGGTAAACTGGGAAAAAACT
AGATCACTAGTCTGTAGCAAATGTGGTTGTGCTGGTCACAATCAAAGGACATGCATTACATAATTGTGCCCAAGGTCATGATATGAAATCAAAGATTAGATAATGACCATGCATGTTTTATGTATCTGAATCTGTAAGTTTGAAATTAAGTAGCATCTGTATGTATCAGACTACAGAGTCCAAACTTCAGACACAAAGAGTAGGCTGGCCATACACTCT
TATTTACTTGAAAAGTGTTTCTGTTCCATATACTGACTGTACTGTCAGGAGTCAATTCTAGCTTCTATGCTTCTATACCTAGATAGTTAAAGTGTATTATATTTGATGTACATAGATGAATTCCACAAGAAAACATCAAATATAAGAAAGCTGAGTTTGTGTTAAATATTTGTATGTGTTTTAATCTTACCTGCCCACACTTTTGAAGGGACTCAAAC
AAAATAATTTGATTACTTTGGCCAAAATGTCAAAAGTTTGAGTAACTGGCCATGAATATGAGATGAGAAGCGTAAGGTTTGCAATAGAATTTTGTACATATATTCATGAAAGATTGTAACAAGTTGTAAAATGAACTCTCAGTAAAGGTTTCATTCAATTCATGCACATGCTACATGTACATGCTATGATTTCTTGTGCAGTTGTCATGTTCTCAGAAA
TTACAATCAGAACATGTAAATTTGATGTTACTCTTATTTTCCAGTTGTCGTGAAGAAAATCGTTTGCCAGAGCATTTAATGTGAAAAGCTCTAACACATTGTGTACATCGCACTGTTGAAAAGTAACTCTTATCAGTCACGCTACAATAATGTTAGCACATGTTGTGAAACATGAAGGGCATTCAAAGTTTATACAGTCAGCTTTGTTTTGTTTTTGAAG
TGATCTAATTCAACACATTTGTCATGGTACCACCTTCCACAGAATACAGTTTGATTGTTCATGATGTAAGTTCGCTCACATTTCAGCCAATTGAACTCTGCTTTATCACTTGCTGGTAAACTAGCAAACTTGCAGCCATCCAACATACATTCCCATGTTTCATTTGTTACTGTCATCTTATGTATTGTCTATCTTTCCTGCTCTGCTGCCCTCTTCTGA
CTCATTCCTGTCTGTGCTAACTTTGACCTCAGCTGCCTCACTTATTCCGTTCTGATCCTCAGTCTCTGAGCCTATACTTTCCTGATCTGTCATGTCTATGCTTATCCTTTCCTGCTCTTCCTCCGCTATGCTTACTCTGTCCTCATCTGTTGACCTTTCCTCCTCATCCCCCAACTGACTGTTATTACTCTGTGTCCTGGTTTCAGAGCTTGTCCCTTC
ATTTTGAAAACTGAATTTTATCATTTGCTTTTTTACTGTATCACAAACAGCAATTGAGCTGAAGAATGGAAGACTGTAGTATTTACTACTTAAGGTATCATAGTGTGCTGTTTCTATGAAATCTATGGCATCACAAGGTGACTTGTAAACTGTGTTGTTACTCAATATGTAAACAGTGTTACTACATTTGAATGAGACTGGGTTAGGAATACCAATAAA
TGGTACAGATGTACAGACCATTGAATTCAGGTCAATTACATCAACATCGGGTGGATGTCTTACATTGTTACCTATCACATAAATTCCACCAACAATATACATTTTACTGCTATTACGAAAAACTGAATGGTAGGCTCTTGGAGTGATCGCTTGTAAACTAATACTTGATAATTTCATAGATTGTAAATTAAGAACAGAAAATATGTTTGAAGCCATGGA
CACATTGTAAAAAGCATGACTGAGTTTTTGTGTATATTGTAAGCCTCCATACAGATAAAGAGAATCTCCAAACTTAATTACACTGTGACCAAAAGCATGAATTCCTCACCACTATATTGTCCTTGAGACCATGTTAATGAAATTTTCCCCCTTGGATCAACAAAGACAGAAAGCTGCCACAATTTTGTATTGTGCATTTCTGAGTCAGTCAATCCACC
AAGGAGAACATAATGGTTTGGTTCAACTAAACACAAATTTGAGCCATGTACATGTGGAGGGTAATTTGACTGAGATGAATAATTTTGATTTCTTGATTTGATATCATCTTTGTCTATTTACAAGCCAGCCTGTTACTGTTGTCTTAGACTTGACCCAGACCAAACTCTGTCACCTAAAGAAATAACTGGATTTGTACATCCAGCAGTTGCTAGGCAGT
TGCCAGGATGTTTTGGTGATTTGTATTTCTGAAATAAAAAGATGATGACAAAAGTTGTTTATTAGAGAGTGAAGCTTATCTGCATGCATGTATGGTGAGCAGGTCATTTTATTTTGTCAGTGTTGAAATATGGCACTGAAATATAAATTTGTTAATTTCATATTTTAGTATATGGATATCTTTGTAGTTCAAAAAAATGCTCAGACAGAAAGTGA
GATGTCTTTAGTCGTGACATTTATTAATATTGTTCACTGTACATTTCTCTACCCTTTGCAACCATACAATTGTTGAACACCTATCCTGTATATTCAAACAAACAACTTAAAGGCAAATCTGATATCTTTGGGGCAGCAAAATACATGTTCTCTTGTCTTTTGCATTTGAGTAACCTCCTTATATAAATATGATCGAAAGAGATTGTCAACATGGAGCAG
CAAATCCGTTTCTTTAACCTAGACTGTAAATCAATACAAAATGGGTGCACATGTAGCTAGGGGAAAAGGTAATGCACAGATAGAGCCCCATTGATGACTGCTGGAAATGTCATTTGGATGTCTTGCTGGTGTACATGCAACTGAAAAAAATAATTTTAGCCTGTGGGCCGTACAATAGCTGCACTTCTACTTCGCTCACTGTAATTTGAATCTTCGTAT
ATAGTGTACATGGGGATCGACTTTCATAGTATGCTTGTTGGATCGAGGGACCGTGGCTTGTGTTACAGTTGAGAACGATCACATCGAGAATTTTCTTTTTCGAATGGGCGATTACAAGTGCAAATTTTTGGAAGTTAGCTACCATATAAGAACTTTGTTGTTTTAACTTCAAGAATAATCAATCTAAAACACGAACAGAATGTTCTGTGACTTTTGAAG
CTACATTGAACTGACGACCTACCGGGACTATCGAGCATGGCGGCTGCATCTTGGATATTGTTTACGGCTCACTGAGCATGCTCAGATCAAAAATATTTCCCACAACCCATTGCGGACTTTCTACATTGTAAACATAGGAGATGTTACGGTGTGAACCTGCGACGACTTGTCGTTAAGATTTCTTGCCTCCATCATCAACTTTTTGTTCAGTTCTCGCT
CTTATCTTGTCATGTCATGGGTAACATAAATTTTTTGAAGAGACTATGTGAAGGGACCGTCCCAAAGTTTTGGAAGACATTTCATGAAATGCTTTTTTGCAGCAACATCAGTCATGCTGACCTTAATTGGTCTGGGTCTGACAGAGTCTGCTTCAGATGAATTATTGATTTTCCTATTCGGACAACATTCTGTATATGTGTACAATCTGCTTCAATAA
CATCAGCTAATTCACGAAGTACATTAGTATCCGCCATTTTTCTTCCAGTTGGGTCAGGTTTATCTGCTCAGGAACACCATAAACAATCATATTCTTCTCTCTGTCGTTACTCTCTCTCTCAGTTCCTGTACTTTCTTTTTGATAGACTCCTCAATGTTTCCTAATGATTTTGTAGTTATAAGTCCTTCAGCTTCAGTGTGCTCCCTTAATGCTATCACTT
TGTCCTCGTCGACTTTTGACTTCATACTATTTTCAAGACTAGTAATTCTATCTTCCATTGATTTCAAAAAAGCTGTACTTGGCTATCAGTTCTTATGTTTTTCATCACCTTCTCTTCACAAGTGCTACAGAACCAATGAAGTGCAGGATTCTTGAACATCTCATAGTCATCTGCTGACAGTTTCAAACATTCTCGGCAAAGTACTTCAGCCAGTATTC
ACACTGAACAATTTCTGCACTGTCGTCTGAAAATTCTTCTTTACAAACTTCACAAATCCATGCCTCCTGGTTCTGGCGCATATGACTCGGACTTGATATAGCTTTCTGCCTTTTGGTAAGAGAGTTTGAAGTGGAATCTGTGTTTTATCACAGATGTGTTTTGTGATTACACCTAGCTATCTGCCTTCCTTACATCTTCACTGTCATCCCCTTAGCCAA
TCGGCTCTTCTACCCATGACTGTTATCAATCCTCCTTCCCAGCCAATCAGCCACGTTTGCACACGACACTACATAATCAAGCCTTTGTCTCATCGCTCCCATTCATTCATTCATTCATTCATTCATTCAGCCGCTGGGTTCATGCCATACCAGCTCAGCCTATTACACTATGTCCGGAATCGTTTACGATCCCTTCCTTCGAACAACCATGCCGGCCAA
AGGACGTCCAAAGACAGCCAAGAAATCTGAGTGTCCGAATGCTCAGCCTGCTTTGAGACCCCGTGCTCAGCCCTACACAAGACCGGCCACTGACCTTTGTCCCGCTCCGCTGCCAGTGCACGTTGGATCTAGCTGCGAATATGGAGCACGCCCGCGGTGTACTAACGTATTATTACTCACAATCCTTCTGATCGGCCTCGGGGAATGACAGTCCTGGCA
TTCAAACCGAGTTCAGAACCAGGTCTCCTCCTAAGTTCCATTGTCTGCAATGGTTCCTTCTGCCAGCATTTCGTCGGGCAGCGTCCCAAGCCAACATTCCAGAAGATAGTTATGTAACACTTCCATAAACATAGCTGCAAGACTCGGTCAACACGATATCCAATATCCTGCAACAGCAACTTCGACCAGCGGCCACTAAGTCACCTATGAACTCTCTGA
CGTCACTTCCCGAAGTTCCACTCCGGAACGTAAACAACACAATCCAAGATGGCGGCGCCCTCGATGTCATCCCCTCCTCATCGTCACGCGTCATCCAACACCTTGCTACAGACCTGCGGAACGGCTTCAGAAAGCCTACCACATATCGAAATGGTGTCTCCAGCGATTCGTCGCGATATCCTGGCAGGAAAGGATGGGAATCTTGCCTACTCATTCCCG
GCTATGGTTCAAGCGACGGACTTCTGCTGCTTATCCACACCTTCAAGCCGA

### PflTRAGL-A_25

| Genomic loci | RAG1L status | RAG2L status . |
| --- | --- | --- |
| PflTRAGL-A_25 | *pseudogenic (frameshifts)* | *pseudogenic (frameshifts)* |

#### FASTA DNA

```
>PflTRAGL-A_25
GTTCAACTATGTTGAACATGCTGTAGGGAGTAGAACAAGAACACAGTTGTATAAACTGAACAAAAATATTGTCAAAATAATTAATCAAAGTTAATACAGGTACTGCCTTATTACTGCGGTGAAACTGTAGGTGCTTACCGGCGGTAACAGAGATAGCTCTAACTCTAATGTAGTTGAAAAAGAGTTTTGGTCAAATGACGCTCATGACAGTGAAATATA
CTTGTACATAAGATCAGACCAGAGAGCAACACATGATATAGAAGACCAGGAAACTTGAAAGTTATCAGGGATTTTTGAATTCTGGCATATGAGAATAATCAAATATTAAAGAAAAAGATGGGGTCACCATGCTTGAATTTCTAAATTTTCACATATTCATCAAAAGGTACACAAAATTTCGAACAGTAGAGACACTAATTTAAATAAAAAAATGTGTGCC
TGAATGGAATTATTTCTAGTCTACCAAATTTGCCATTATTGGTATTACATCCCAAATTTTAAGATTAAACGTTTTTTATGCAATATTTTAATCTTCCAGTATTTTAAATTTAGAACAGCATCTAAGTTATATATGTCTCGTGCTTTTGAAAAAAAAATTGCAAACTTTGGCAAAGTGTGGCCAGGACTACTCTCTCATGATTGTCATCTTGTGTGCTCT
TCAGCAGGTGCCATTGACCTGGCTAGAGTTGGATTAACTCAAGCCGGCTTAGACTGATAATCCAAAAAATGCATTGATTCGTTTAAAACGAATCAATTTAAGAACTTGCCTTATAGGAATGTGGTCCTACAAACTGATAATAACAGGTAGTGTCGAATATCTGAGAGTAGAACCGCCCAATACATGTCTATTTTTTCTTTGCGTGCGTCACTAATAAGT
ATGCTGCTATATGATAATTTAAGGGGGACTTGACAAATATTGATCATATATTTTTATATAAATTTGTGAACGCAGCCTTGGTAGCTATGCTCACTATGAAGGAAGTTTCTAGACAGAAGGCCATCACAGCGTAACTTGTCAATATAGAAATCGATTAACACTACGATGCGTCCCAGAGAGTATCGACCAATCAGAGAGCAGCATTCATCAAGCCACGAA
CTTTCACGTATGGCTTCGAGATTTCGCGTGATCTAGGGTGTTTACTTCAGAAACTCTACAATCTACTAAACACTAAGGGGTCTCAACTGTTTTACACATGCGCAAACGCCATCTCAAAGAGTGGCTGATGATCATGTTGAAAACCGGAGGTCGAAAGACGTAGGTATCGCCGAGATATCGTAATCTGAATCTCGCATTGCAATTTTTCACTCCATTATG
TGTATCAAAATATGCGGATACAATTTTTGGATTTTCTGTCAAACTTGGTTTGTCCTGTTGAAAGTTTGCGTTTTTCTCAGTCCATAACGACATTGTTTACCGAAGGAACATATGCTGTACGCTTGGACTTTCTTCAAATGGCTTGGACTTTTAGATCAATACCCATCCAACATGTGTGATTGAGTGTAATGGCCGTTACATTTTAAAATGTTCCTGTGA
TGGTAGCTAGGTCAGGGAAATGATCATTTTCTGTTTAATATATTTTTAAGGTTTGAATGACCGTGTTACGATATTTTGAATTCTGAGCAGCATCCTAGAGAGTCAATTTGGGTGTACAAGGTATGGTTCATGCGGCGGTATTAGCTAGCTAACCCAGGCCATATGGCACCACGTCTAGGTTGACTAACGAAATGTTTTGAGGTTAGCATAGTATGAGTA
CGTCAGATTCATATGTCATCGGCATAGAAGGTCGCTTGTGTAAGATAAGACCATATTTTTAGGCTGTCATTCATTTCATTTATTTATTTGCAGTTGGTTGCAGTCAGCAATGAATCATTTACAAGTTTGTTGCACTGTTATGCAATGCAATGCCTTTACCTGATTCAGTATTTCTTGCTTGGGTTCCTGCGGTATGAAGTAGAAAATTAGGCTGGCCAT
GGTGTCCTGGGAATTAATGTTTTACCACCAAAGTAAGTATAAAATTCTATTGGATTTATTGTTTACACATATATAAGGATGAAAGGTACAGACATTGTACTGATTGGTACTGTACTTGCCAAATTGATATGCATAAATAATATTTAAATATGTTAATTGTTTTCAGAGCCGAATGACATCAGTCTTTCAACACAGAGAGTACCTTAAGAGGGTGTGTCG
TGTATGTGGTAGAACCCTGAAGACCACTTCTTACAGCAGCTCAGGACATTACAGACAGGGAGAAGTGTGCATCCCATACAGTGAACTTTTAGAAGAATGCTTTAAAGTAAATGTTCGAAATGATGACCCTTTTGTACATCCGCCTAGATTTGGTTCATGTTGTCGTAGAGTTTTTTAAAGTATCCTGTTGCACAGAACCTGGGTAAACATGACTATACC
CATATCAGCGTTAGTAACTGGAAACATCATTGTAAGTATGAAACCAAACTTGTCCTCTTTGCAAACGTTTTTCTCAAAATGGTTCTGGTAGACATAAACAACTTAAAAGGGGTCGATTTAAATTAAATACAAATGAAAATAATGAAACTGACAAGAGTCCTGTTGCAGACAATTTTGCTCATATACCAATGACCACTGTTATACCACCCGAGCTTCCTG
TAAGTTTGCCCAGTCTGTGCAATCAGACCAAAAGACAGAGACATATGAATTCTGTACAGAGGAAAATATTCAGACTGTACCCATGACAAGTCGATTTCCAACAGATTAATGTGTAACTGAATCACATCCAGAATTCATTTGTAGAATGTGTTTAGGAGTCCTAGACACGTACACCACTTCTAACACCGTGTACTCACACTTTCTGTGCAGCGCTGCATT
AAGAACTGGTTAAGTCTGTGTCATTTTTGTCCTCTCTGTAAACAACCTATGGTGTCATCAGATTTAAAAGAACCATATAGAGGATTTGTAGAAATTCTTAGTGATACTAGAAGCTTAGTAATTGGCTTCAGCACACTTTAAATTGTGGAAAAGATTTTGACATTGATGAATGGTTAGAAAAAGTTTTGTCATTGTGGAACGAAATTTCTCTCACACAAG
TTGATTCACATTTAAATAGTACTGAACATTTGCCTGAGACAGATAAGTCAAATCTGACAACAAGCCATGGCCATCCAGACATTGCTGCAGGTAGCACTGTTGGAAGACCTAAAACTTCACTGTTGTGTCTCCAACAACAAGGACAAAATTTAAGATTGAAAAATATCAAAAAACAAATAGACGTGTTTGCCAGAGAGAAAGGGGAAGATCTCAGGGTAG
TATATTTTGGCCTTTTGCAAAACTATCTGAGAGCCAACAAGCAAGGTAGTTTAGCTGAACAAGTGAAGTTGCTATATAATTGGGAATCTTGTCAAATGTCACCTAAGGAATGTCTAACCTTGCGTGTAAATACTTTTACCAGTTGTAATCAGTACAAGAAAATTTACTCAGCTACAAATGAGCTTAGTGGAGTTAAAATCTTCCAACCTCTCATCAATG
TTGTGAAAGCCGAAGAAAGTTTTCTACTAGGGTCTGTAAAGTTCTACTTTAACCCATCTTTGAAAACTGTTTCTCCCATTTGGCATGGTACAGAGCATTTTGATGCTCTGTCAGGACTCAGTCGCGATATACCACTTCTAGAATCACCTGATGTTTATGGGGCAAGGTATAGATATGACAAAGCAGTTGCTATGGCTTTGAAATATATTGATAATGAAA
TCTCAGAAGGCCTCAGTAAATTAGAAGGGGTGGATACAAATGAACTTGTTCTAACAGCCACTATCAAAGATGGCAGTGATGGTCTTGGAGTTGTGAAAGATGTCAGATCTCAGGGCTTGTCGTTCCAAACAAGGCCTTGCGGTTTTCATTTGTAATTTTAGAAGTCACTGTCAATGTAGATGGCAAGATTATTACTGCTTTTGAAGAAGGTTCTCCCAA
CTCAGAATTATGCACACGCCCTCTCTTGTAGCTCTGGCAGATGAAAATGACAGAAAAGGGTTATGCATGACACATATACCAATAATGATAAAAAGAGAGGAGATGGAGAAAGTACCTTGTTTGTAGAAATTGAAGAGGGAAAACAAAGAATCTACAAAGTTGTATTTAAAGGGACTATGTATGATGAAAAATTGCAGAGAAGTGGTGAGGGAATGTTAG
CGGCAGGGTCATCTTGGTTTTGTACACTTTGTGAGAAAGGAAGACTTGATCCCATATCATCAGTCTTGCCAATAACAAGAACACATGAAAACAATATGGCTTTGTATTCAAAGTATGTTGAAAATGCAAATCATATGAAGCAGGCAGATCTAGTGAAAGAAGTCAAAGGCATTGTTAGAAAACCATTTATAATGTCAGAGCCTTGTGTTGATGCCACGC
ATGCTGAAATTCACTGGGGAGAGAAAAACTATGATATGTATGTACGTGAAATATCAGGGATAATGTGTTGGAGGAACCCATCCAATGATGATGATGCAAAAAAGTTCAATCAATGAAAAGAATGTTGGATACTAAATTACAGCTGCAATGTGGCTTAAAGAGAGAGCATGGCTTTATGGTTGATGGAAACAAGTCATCGTTGATGACACAGTCCCCACT
TGTTAATGGGTACTTTGATTACATGTCCTAAGAGGATAGAGTCCTCTTCATTAATTGAGACATGCCCAAGTTTTGGCTAAGGACACGAAAAAATTGTAACAAAATGGCTGCCATGCAGCCATATTGGATCATATCAAGAAACAAATTGACATACATATGTATGACATAGATTAATGTCCTTGTACCAACTTTGAATAAAATCGGTTAAGATATGCCTTA
GAGTATTATGGCCCTCGACAGGAAAAAATCGTAATAAAATGGCCGCAAGGCAGCCATATTGGATCGTATCACATAACAAATTGACATGCATATGTATGACATTAGTCAATCTCCTTGTACCAACTTTGAATAAAATCCGTTGAAACATGTCTGAGTTATTGCTCTGTACATGAAAATATCGTAATAAAATGGCTGCCTAGCGGCCATATTGGATCGTAT
CACATAACAAATTGACGTACATATGTAAGACATAGGTCAATGTCCTTGTACCAACTTTGAATAAAATCGATTGAGATATGCCTGAGTTATGCCTCTGCATATGAAAAAATCGTAATAAAATGGCCACACAGCAGCCATATTGGATTCTATCACAAAACAAATTGACATGCATATCTATGACATTGGTCAATGTCCTTGTACCAACTTTGAATAAAATCG
GTTAAAACATGTCTGAGTTATGGCACTGTATATGAAAATATTGTAATAAAATGGCCGCCTGGCGGCCATATTGGATCGTATCACAAAACAAATCGACATGCATATCTATGACATTGGTCAATGTCCTTGTACCAACGTTTGATAAAATCAGTTGAAACATGTCTGAGTTATGGCTCTGTACATGAAAAAATCGTAATAAAATGGCCGCCTGGCGGCCAT
ATTGGATTGTATCACAAAACAAATTGATGTGCATATCTATGACATTGGTCAATGTCCTTGTACCAACTTTGAATAAAATCTGTTGAAACATGTCTGAGTTATGGCTCTGTACATGAAAAAATCGTAATAAAATGGCCGCCTGGCGGCCATATTGGATCGTATCACAAAACAAATCGACGTGCATCTGTATGACATATGAAGTAATCCTTGTACCAAGTT
TGAATGAAATCGCTTCTTGCATCTCTGAGATATCTGCGTGAACGGACGGACGCACACACGCACGCACGCACGCACGGACGCACACACGCACGGACATGACCAAACCTATAAGTCCCCCCGGACGGTGTCCGTGGGGACTAATTATGCACGTGATTGGTCAAATCAGAAACAGTTGATGTGGTTTGCTCTTTAATTCAGAGTGAAGCAAGAAAATCTGTT
GTTCGTGAATATGTTTCCTACTATCGAGAATTGCGATCAATATACAGGAAACACAATGCTGACTTAGATGATGCAGTGAAATTTAAAACCATTGCTGCAAATTTTACAAATTGCTACATGAGAAATTTCCTTATTTGCATCCACTCTCTAACTAGCAACACAAGGTATTAGATCATATTCCACAGTTGATCGAGCAATATGGCTCTGTTGGTAAATTT
GCCTCTGAAGGCAATGAAAAAGGCAATAAACTTTTCAGAAGATTTTGTAAATTTAATGCACGCAAGTCTCAAATTCATGAAATGCTAGACATTCTGAAATTTCATTGGCTATATACAATGAAAAGTCTTCAGACTCTAGTAAACTGGGAAAAAACTAGATCACTAGTCTGTAGCAAATGTGGTTGTGCTGGTCACAATCAAAGGACATGCATTACATAA
TTGTGCCCAGGCTCATGATATGAAATCAAAGATGAGACCATGATCATGCATGTTTTATGTATCTGAATCTGTAAGTTTGAAATTAAGTAGCATCTGTATGTATCAGACTACAGAGTCCAAACTTCAGACACAAAGAGTAGGCTGGCCATACACTCTTATTTACTTAAAAAGTGTTTCTGTTCCATATACTGACTGTACTGTCAGGAGTCAATTCTAGCT
TCTATGCTAGTCTATACCTAGATGGTTAAAGTGTATTATATTTTGATGTACATAGATGAATTCCACAAGAAATCATCAAATACAAGAAAGCTGAGTTTGTGTTAAATATTTGTATGTGTTTTAATCTTACCTGCCCACACTTTTGAAGGGACTCAAACAAAATCATTTGATTACTTTGGCAGAAATGTCAAAAGTTTGAGTTATACTGGCCATAAAAAA
TGAGATGAGAAGCGTAAAGTTTGCAATATAATTTTGTACATATATTCATGAAAGATTGTAACAAGTTGTAAAATGAACTCTCAGTAAAGGTTTCATTCAATTCATGCACATGCTACATGTACATGCTATGATTTCTTGTGCAGTTGTCATGTTCTCAGAAATTACAATCAGAACATGTAAATTTGATGTTACTCTTATTTTCCAGTTGTCGTGAAGAAA
ATCGTTTGCCAGAGCATTTAATGTGAAAAGCTCGAACACATTGTGTACATCGCACTGTTGAAAAGTAACTCTTATCAGTCACGCTACAATTAATGTTAGCACATGTTGTGAAACATGAAAGGCATTCAAAGTTTATACAGTCCAGCTTTGTTCTATTTAAGTGACCTTATTCAACACATTTGTAATGGTACCAACTTCAACAGAATACAGTTTGATTGT
TCATGATGTAAGTTCGATCATATTTCAGCCAGTTGAACTTGGCTTGTTCAGTAGCTGCTAAATTGGCAAATGTGCAGCCATCCAACATACATTCCCATGATTCCTCCTCACTATCATCTGCTGCCCTCTTCTGACTCATTCCTGTCTGTGCTAACTTTGACCTCAGCTGCCTCACTTATTCCGTTCTGATCCTCAGTCTCTGAGCCTATACTTTCCTGA
TCTGTCATGTCTATGCTTATTCTTTCCTGCTCTTCCTCCGCTATGCTTACTTTGTCCTTATCTGTTGACCTTTCCTCCTCATCCCCCAACTGACTGTTATTACTCTGTGTCCTGGTTTCAGAGCTTGTCCCTTCATTTTGAAAACTGAATTTTATCATTTGCTTTTTTACTGTATGACAAACAGCAATTGAGGTGAAGAATGGAAGACTGTAGTATTTA
CTACTTAAGGTATCATAGTGTGCTGTTTCTATGAAATCTATGGCATCACAAGGTGACTTGTAAACTGTGTTGTTACTCAATATTTAAACAGTGTGATTACATTTGAATGAGACTGGGTTAGGAATACCCATAAATGGTACAGATGTACAGACCATTGAATTCAGGTCAATTACGTCAACATCGGGTGGATGTCTTACATTGTTACCTATCACATAAATT
CCACCAACAATATACATTTACTGCTATTTAGAAAAACTGAATGGTAGGCTCTTGGAGTGGTAGCTTGTAAACTAATACTTGATAATTTCATAGATTGTAAATTAAGAACAGAAATATGTTTGAAGCCATGAACACATTGTAAAAAGCATGACTGAGTTTTTGTAGATTATCATACATACAGATAAAGAGAATCTGCAAACTTTATTACATTGTGACC
AAAAGCATGAATTTCCTCACCACTATATTGTCCTTGAGACCATGTTAATGAAATTTCCCCCCCCCCCCTGGATCAACAAAGACAGAAAGCTGCCACAATTTTGTATTGAGCATTTCTGAGTCAGTCAATCCACCAAGGAGAACATAATGGTTTGGTTCAACTAAACACAAATTTGATCCATGTACATGTGGAGGGTAATTTGACTGAGATGAATAATT
TTTGATTTCTTGTATTGTATCATCTTTGTCTATTACAAGCCAGCCTGTTACTGTTGTCTTAGACTTAACCTAGACCAAAACTCTGTCACCTAAAGAAATAACTGGATTTGTGCATCCAGCAGTTGCTAGGCAAGTTGCCAGGATGTTTTTAGTGATTTGTATTTCTGAAATAAAAAAGATAATGACAAAAGTTGTTGTATTAGAGAGTGAAGCTTATCT
GCATGCATGTATGGTGAGCAGGTCATTTTATTTTGTCAGTGTTGAAATGTGGCACTTAAATAATAATAATAATAATAATAATAAGGATATTTATTGTGCGCCTATCTCGCAGAGCTCTAGGCGCATGGCTACAAAAGAACTCTCATCAGGTGTGAGATTTAAAAAGGTGTGTTTTAAGAGTAGAACGAAATGATGCTAAAGATTGAGTATGACGGAGAT
TTGCAGGTAAACTATTCCAAGTCGTAGGACCAGTATAGGAAAATGATCTTTCGCCGAAGCTTTTCGTGACAGCACGAGGTACTTTGAGTACACGAGTATCAAAGGAAGAACGCAACTGTCTTGAGGGACAGTACTGTTGAAGAAGGTCTGTAAGGTATTGCGGCCCAGATCCAATAAATGAATTGAAACAGATTACTGAAAGTTTGTATTGTATGCGAG
ATGTAATTGGTAACCAATGTAGCGAACACAAAAGAGGTTGGACGGAGTCCGATTTCTTGACCTTATACACTAGTCGTGCAGCAGCATTTTGCACTCTCTGCAACTTTTGGAGTAAATATTTGGGACAACCAGCTAAAAGAGAGTTACAATAGTCCAGGCGCGAAAGTACAAGTGCGCAGATTAAAGTTTTGGTGGCTTCCACAGTGAGTAAGTGACGAA
TTGAACTAATATGTCGGAGGTGGATGTAGGCGACTCTACAAATATTTTGAACATGAAGCGACATATTCAAGTGACAGTCAAGTGTGACACCAAGGTTTTTCACATTTCTAACAAATTGAATATCAGAATTCCCGACACGCATAGAGGTAGGGGTTAATTTTGGAGATGATCTTTGAGATATGACCACCATTGCTTCAGTTTTGTTGTCGTTCAACTGAA
GTTTATTGTGGGTCATCCAAGTTTTGACGTCACTGATACAATTTTGCACAGTATTGATTGTCACTTGAATATTGTCAATTGTGCTGACTTTATAAAGTTGGTTATCGTCAGCAAAAGCATGGTGCTGCACTCCGTGATTTCTGACAATTTCGAAGAGTGGTTCTGTGTACATGACAAACAAGACGGGCCCAAGTACAGAACCTTGAGGAACTCCAAACG
AGAGAATGGACGAGTCTGATCTCACACCATTGATAGTAACAGTTTGCGTTCTACCAGAAAGATAGGACTCAAACCAGGACAGCGCAGATTCAGTTATTCCAAAAGTGTGCTGAAGTCTGTGAAGGAGGATGCTGTGGTCTATAGTATCGAAAGCAGCTGAAAGATCTAAAAGGGTCAAAATGCTGACTTTGCCATTATCAAGAGCATTCAAGAGATCAT
TAAATATTCTCAAGAGAGCAGTTTCAGTGCTATGTTTAGGTCTATAGGCAGACTGAATGGGGCTGAGGAGATTATTGGAGCTGATGTGGTCATTTAATTGCATGAGAATTATTCTTTCTGTTATCTTGGACAAGAATGGAAGATTAGAAACAGGGCGGTAGTTTTTAGATCATTGGGGTCTAACGATGGCTTTTGAGGAGAGGTGTTATAAGAGCTG
TCTTGAAAATTGATGGAAATGTTGCAGAAGTGATGGAGTCATTGATAATCTTAGTAATGTGCGAAAGTAAAGTGTCTAAACACTCTACTAACAAACTAGTGGGCATGACATCAAGATCACATGTTTTTGTTGCAGAATTAAGTGTGGTACGTTTGACTACATCCTCGGAAACTGGTTTGAATTGATTCAGGGGTCGTCCATGGAAGCAGCGTTCAGAGT
AGCTAAGTGTCTGTGGATTGATGCCCTCAAGATTATCTCTGATGGATTTTATTTTACCATTAAAGAAATCCAGAAATCGCTGTGGTAATTCAGATGTAGAAAATATGTTGGGAGAATTAGGGGTTTATTTTTGCCAAGTAACTGATTAGCAACACTGAACAACTGCTTGCTTGAACGACTCTCACTGATCTTGCTGCTATAGTAATTGGTTTTAGCCAG
GTGAACAAGTTCAGTTACACGCGACTTTTCATTGCTGTAGATTTGTTTGTGAATTGCCAATCCTGTTTTTAGCCATTGTCGCTCTGCACGCCGTCGTTGTTGCTTTGCCGCTCTGAGATCATCCCGCACAGCATTATACCAAGAAGAACTTCTCTGTGAAGAAAGTCTTTGTGGTCTCGATGGTGCGTGTTTGTCCATGATTAACTGCAATTGGGAGTT
GAGAGAAACCGCTGAAGGATGTTCAAGATGTAACAGAGCATTATGAAGGTCTTCTTTCAAGGCTGGGCGATTTATCGTACGAATGTTACGTGTCATCTTGATTGTGGTTGATGAGCACGGAAGCGACAAGTTCAACTGGCTTATAACACACAGATGATCGGATGAAATATAAATTTGTTAATTTCATATTTTAATATGGATATCTTTGTAGTTCCAAAA
ATGCTCAAGACAGAAAGTGAGATGTCTTTAGTCGTGACATTTATTAATTTTGTTCACTGTACATTTCTCTACCCTTTGCAACCATACAATTGTTGAACACCTGTCCTGTATTCAAACAAACAACTTGAAGGCAAATTTGATATTTTGGGGGCAGCAAAATACATGTGCTCTTGTCTTTTGCATTTGAGTAACCTCCTTATATAAATATGATCGAAAGAG
ATTGTCAACATGGAGCAGCAAATCGTTTCTTTATCCTAGACTGTAAATCAATAAAAATGGGTGCACATGTAGCTAGGGGAACAGGTAATGCATAGATAGAGCCCCATTGATGACTGCTGGAAATGTCATTTGGATGTCTTGCTGGTGTACATGCAACTGAAAAAATCATTTTAGCCTGTGGGCCGTACAATAGCTGCACTTCTTCGCTCACTCTAATTT
GAATCTTCGTATATAGTGTACCTGGGGATCGACTTTCATAGTATGCTTGTTGGAGGGACAGTGGCTTGTGTTACAGTTGAGAACGATCACATCAAGAATTTCCTTTTTCGAATGGGCGATTACAAGTGCAAATCTTTGGAAGTTAGCTACGATATAAGAACTTTGTTGTTTTAACTTCAAGAATAATCAATCTAAAACACGAACAGAATGTTATGTGAC
TTTTGAAGCTACATTGAACTGACGACCTACCGGGACTATCGAGCATGGCGGCTGCCATCTTGAATATTGTTTACGGCTCACTGAGCATGCTCAGATCAAAAATATTTCCCACAAACCATTGCGGACTTTCTACATTGTAAACATAGGAGATGTAACGGTATGGGCCAGTGTTCCAATCGTTCTGTTTAATAAGTAAAGTCCGAGACATATCTCTCGGCC
ACAGTGATCGGTGTCTACAGTCCTGTAAGATATACTTACAACCTACTCATTGGCTGGACCCTTTCTCTTAGGTCTCACTACTGTGTAATGTTCAATTCACGGCACTGTGATTGGATACCACATGTCCCTGTGTTAACCACGCTTTGTAATGTTTTAGTTCTCTCTAGTTTTTAGAATTCATTTTCGTTCTTACGGTGTTTTTAAAGCTGAGTTGTTTTA
ACCTTGGTTTTAAAGTATCCTAGCATCAAACATTTCGGAAGAAGACGTCATCTGTCGTTCCAGAGACTGTAACTTTCTGCTTCTTTTCAGAGTTTCTGACGCACTTTGTGATCAGTTCAACATTGCAGCGTGTACTTTCCTATTCCGTCATAGCCTTTGTTCTTCATTGTAAAAAACTTCCAAACCCCACACATGCAAAGCTGGACGGAAAAATCCTAA
GCCAGTATACCGTATTTCGACCATCGTCCATACAGATCGGACTTCCAGGCTACTAAAGAAAAGTTCTCGAACAAGTATGCTTCAGCAGATATGTGTTCCAAATATTGCATACGGTGAGCTAAAAGCAATTGAAACTGTCGTTACAAATCATACCATTCAACCAGCCGCTAAGAAGAGAAAAATAAGTGTTATTGAAAATTTCTCCACATGTTTTCAAAG
TTGCAACTGAATGCTGCCCTTTGGAATGCACATTCAATGACTAGGAAAATTGGCATCATTGCATCAACTCTGATCGAAGATAATCTCGATATCTTAGTCATCACAAAACAAGGCTCAAGAATCAAACCGACCCTTTAATCGCCGATTTTCATTCAGCGGTTAAGTGGATACAATGTTCACCACCAGTCTCGCTCAAGAAGAAGAGGCGGCGGTTTCGCT
GTGATCGCTCACTCCAATTTCGCGTGTTTCAGAGAAGCAATCGGTTCATTAAATTCCTTTGAAT
```

### Ophiothrix spiculata

#### RAGL-A

##### OspRAGL-A.1

| Genomic loci | Scaffold ID | RAG1L status | RAG2L status |
|---|---|---|---|
| OspRAGL-A.1 | JXSR01S003992.1:1-9300 | potentially functional (Obs*) | potentially functional (1 CDS) |

***Obs.** The RAG1L-A locus on the WGS scaffold JXSR01S003992.1 is intrerupted by 2 contig merging area, each flanked by duplicated segments of <100bp. To recover the complete sequence, region spanning the conting merging areas were amplified and sequenced as described in the Methods section.

```
             1                                                                                                                                                                                                     200
JXSR01S003992 GTTCCTGTTTTAGGGACAAAGCAAATTGTGAACAGCACAATCATTTTACCAGTTTATACAAATGTAATTACTAATCATCTCAAAAACTACAAGACAGATCAAGAATTTTTTTGCATCAAAACATGCGCAACCCCTAACTTGACGATTACATGGAACAACTTTATTTGGGTTATGTTGTATTTTTGCGTGAAATCATGGGT
         Seq1 GTTCCTGTTTTAGGGACAAAGCAAATTGCGAACAGCACAATCATTTTACCAGTTTATACAAATGTAAT-GCTAATTATCTCAAAAACTACAAGACAGATCAAGAATTTTTTTGCATCAAAACATGCGCAACCCCTAACTTGACGATTACATGGAACAACTTTATTTGGGTTATGTTGTATTTTTGCGTGAAATCATGGGT
         Seq2 GTTCCTGTTTTAGGGACAAAGCAAATTGCGAACAGCACAATCATTTTACCAGTTTATACAAATGTAAT-GCTAATTATCTCAAAAACTACAAGACAGATCAAGAATTTTTTTGCATCAAAACATGCGCAACCCCTAACTTGACGATTACATGGAACAACTTTATTTGGGTTATGTTGTATTTTTGCGTGAAATCATGGGT
         Seq3 GTTCCTGTTTTAGGGACAAAGCAAATTGTGAACAGCACAATCATTTTACCAGTTTATACAAATGTAAT-GCTAATTATCTCAAAAACTACAAGACAGATCAAGAATTTTTTTGCATCAAAACATGCGCAACCCCTAACTTGACGATTACATGGAACAACTTTATTTGGGTTATGTTGTATTTTTGCGTGAAATCATGGGT

             201                                                                                                                                                                                                   400
                                                                                                                                                                                                      RAG1L-A
                                                                                                                                                                                 M  D  Q  H  V  S  H  L  A  K  C  C  R  V  C  A
JXSR01S003992 CCATATGGACCCCCCCCCCC-ATTACATGTAAGTGTTAATTTTTGAATTAAAAAATGTTTGTTTGTTTTTGTTTTTAAAAGTGTTGCAAACAAAATCAAAAACAAATATTCTAATTTTGATTTCTGGATTGTTCCTTGCAGGCAGTGATCTCAACATGGACCAGCACGTCAGCCACTTAGCGAAATGCTGTCGAGTATGTG
         Seq1 CCATATGGACCCCCCCCCCC-ATTACATGTAAGTGTTAATTTTTGAATTAAAACATGTTTGTTTGTTTTTGTTTTTAAAAGTGTTGCAAACAAAATCAAAAACAAATATTCTAATTTTTATTTCTGGATTGTTCCTTGCAGGCAGTGATCTCAACATGGACCAGCACGTCAGCCACTTAGCGAAATGCTGTCGAGTATGTG
         Seq2 CCATATGGACCCCCCCCCCC-ATTACATGTAAGTGTTAATTTTTGAATTAAAACATGTTTGTTTGTTTTTGTTTTTAAAAGTGTTGCAAACAAAATCAAAAACAAATATTCTAATTTTTATTTCTGGATTGTTCCTTGCAGGCAGTGATCTCAACATGGACCAGCACGTCAGCCACTTAGCGAAATGCTGTCGAGTATGTG
         Seq3 CCATATGGACCCCCCCCCCCCATTACATGTAAGTGTTAATTTTTGAATTAAAACATGTTTGTTTGTTTTTGTTTTTAAAAGTGTTGCAAACAAAATCAAAAACAAATATTCTAATTTTTATTTCTGGATTGTTCCTTGCAGGCAGTGATCTCAACATGGACCAGCACGTCAGCCACTTAGCGAAATGCTGTCGAGTATGTG

             401                                                                                                                                                                                                   600
               I  M  F  C  R  I  P  K  G  G  K  G  Y  E  C  L  G  L  D  P  V  S  K  I  P  W  R  E  L  L  L  E  C  F  E  V  D  P  A  N  D  D  A  N  I  H  P  K  K  F  C  C  K  C  K  A  I  I  K  R  F  C  D  A  K  V  .
JXSR01S003992 CAATTATGTTTTGTAGAATTCCAAAAGGCGGCAAAGGCTACGAATGTCTTGGACTTGACCCAGTATCTAAAATACCATGGAGAGAATTATTGCTAGAGTGCTTTGAAGTAGACCCAGCAAATGATGACGCAAATATCCATCCCAAGAAATTTTGCTGCAAATGTAAAGCAATTATAAAACGATTTTGTGATGCTAAAGTA
         Seq1 CAATTATGTTTTGTAGAATTCCAAAAGGCGGCAAAGGCTACGAATGTCTTGGACTTGACCCAGTATCTAAAATACCATGGAGAGAATTACTGCTAGAGTGCTTTCAAGTAGACCCAGCAAATGATGACGCAAATATCCATCCCAAGAGATTTTGCTGCAAATGTAAAGCAATTATAAAACGATTTTGTGATGCTAAAGTA
         Seq2 CAATTATGTTTTGTAGAATTCCAAAAGGCGGCAAAGGCTACGAATGTCTTGGACTTGACCCAGTATCTAAAATACCATGGAGAGAATTACTGCTAGAGTGCTTTCAAGTAGACCCAGCAAATGATGACGCAAATATCCATCCCAAGAGATTTTGCTGCAAATGTAAAGCAATTATAAAACGATTTTGTGATGCTAAAGTA
         Seq3 CAATTATGTTTTGTAGAATTCCAAAAGGCGGCAAAGGCTACGAATGTCTTGGACTTGACCCAGTATCTAAAATACCATGGAGAGAATTACTGCTAGAGTGCTTTCAAGTAGACCCAGCAAATGATGACGCAAATATCCATCCCAAGAGATTTTGCTGCAAATGTAAAGCAATTATAAAACGATTTTGTGATGCTAAAGTA

             601                                                                                                                                                                                                   800
             N  H  R  H  F  S  H  S  T  E  V  V  H  W  E  A  H  C  D  E  C  K  V  C  L  R  F  L  G  K  S  R  G  R  H  V  K  K  H  R  K  C  G  R  P  K  A  S  C  R  N  E  Q  I  E  E  P  F  I  Q  T  S  E  N  L  N  D
JXSR01S003992 AATCATCGACATTTTTCTCACTCCACAGAAGTAGTACATTGGGAAGCACACTGTGATGAATGTAAGGTTTGTTTACGCTTCTTAGGAAAATCTAGAGGACGGCATGTTAAGAAACATCGAAAGTGTGGACGTCCTAAAGCTTCATGTAGAAATGAGCAAATAGAAGAACCATTTATTCAAACATCAGAAAACTTAAATGA
         Seq1 AATCATCGACATTTTTCTTACTCCACAGAAGTAGTACATTGGGAATCACACTGTGATGAATGTAAGGTTTGTTTACGCTTCTCAGGAAAATCTAGAGGACGGCATGTTAAGAAACATCGAAAGTGTGGACGTCCTAAAGCTTCATGTAGAAATGAGCAAATAGAAGATCAATTTATTCAAACATCAGAAAACTTAAATGA
         Seq2 AATCATCGACATTTTTCTTACTCCACAGAAGTAGTACATTGGGAATCACACTGTGATGAATGTAAGGTTTGTTTACGCTTCTCAGGAAAATCTAGAGGACGGCATGTTAAGAAACATCGAAAGTGTGGACGTCCTAAAGCTTCATGTAGAAATGAGCAAATAGAAGATCAATTTATTCAAACATCAGAAAACTTAAATGA
         Seq3 AATCATCGACATTTTTCTTACTCCACAGAAGTAGTACATTGGGAATCACACTGTGATGAATGTAAGGTCTGTTTACGCTTCTCAGGAAAATCTAGAGGACGGCATGTTAAGAAACATCGAAAGTGTGGACGTCCTAAAGCTTCATGTAGAAATGAGCAAATAGAAGATCAATTTATTCAAACATCAGAAAACTTAAATGA

             801                                                                                                                                                                                                  1000
              H  S  Y  V  S  L  A  A  Q  F  I  G  K  R  Q  P  S  S  T  A  N  S  T  D  P  D  L  S  S  D  V  G  L  S  D  S  C  I  S  G  L  T  F  N  D  S  S  L  S  V  C  V  N  L  S  T  H  D  Q  N  N  S  L  C  G  N  S
JXSR01S003992 TCACAGTTATGTGTCTTTAGCAGCTCAGTTTATTGGAAAACGACAGCCAAGTTCTACGGCAAATTCAACTGACCCGGATTTATCTTCAGATGTAGGCCTTAGCGACTCATGTATATCAGGCCTCACGTTTAATGATAGTTCTTTGAGTGTATGTGTGAACTTATCAACTCATGATCAAAACAACTCGCTATGTGGCAACT
         Seq1 TCACAGTTATGTGTCTTTAGCAGCTCAGTTTATTGGAAAACGACAGCCAAGTTCTACGGCAAATTCAACTGACCCGGATTTATCTTCAGATGTAGGCCTTAGCGACTCATGTATATCAGGCCTCACGTTTAATGATAGTTCTTTGAGTGTATGTGTGAACTTATCAACTCATGATCAAAACAACTCGCTATGTGGCAACT
         Seq2 TCACAGTTATGTGTCTTTAGCAGCTCAGTTTATTGGAAAACGACAGCCAAGTTCTACGGCAAATTCAACTGACCCGGATTTATCTTCAGATGTAGGCCTTAGCGACTCATGTATATCAGGCCTCACGTTTAATGATAGTTCTTTGAGTGTATGTGTGAACTTATCAACTCATGATCAAAACAACTCGCTATGTGGCAACT
         Seq3 TCACAGTTATGTGTCTTTAGCAGCTCAGTTTATTGGAAAACGATAGCCAAGTTCTACGGCAAATTCAACTGACCCGGATTTATCTTCAGATGTAGGCCTTAGCGACTCATGTATATCAGGCCTCACGTTTAATGATAGTTCTTTGAGTGTATGTGTGAACTTATCAACTCATGATCAAAACAACTCGCTATGTGGCAACT

             1001                                                                                                                                                                                                 1200
               Y  V  E  S  N  S  D  S  G  L  D  S  S  N  N  T  S  E  F  S  Q  P  L  A  A  S  T  P  C  R  R  G  R  P  P  L  P  L  T  S  L  C  Q  N  S  Q  N  Q  R  L  K  P  I  K  T  V  I  K  Q  H  A  E  H  F  H  E
JXSR01S003992 CCTATGTTGAGAGCAACTCTGACTCTGGATTGGACTCTAGTAATAATACATCTGAATTCAGTCAACCTTTAGCAGCCTCAACACCATGTCGAAGAGGCAGACCTCCACTTCCGCTAACCAGCTTGTGTCAAAACTCACAAAATCAGAGACTGAAGCCTATAAAGACAGTAATCAAACAACATGCAGAGCATTTTCATGAA
         Seq1 CCTATGTTGAGAGCAACTCTGACTCTGGATTGGACTCTAGTAATAATACATCTGAATTCAGTCAACCTTTAGCAGCCTCAACACCATGTCGAAGAGGCAGACCTCCACTTCCGCTAACCAGCTTGTGTCAAAACTCACAAAATCAGAGACTGAAGCCTATAAAGACAGTAATCAAACAACATGCAGAGCATTTTCATGAA
         Seq2 CCTATGTTGAGAGCAACTCTGACTCTGGATTGGACTCTAGTAATAATACATCTGAATTCAGTCAACCTTTAGCAGCCTCAACACCATGTCGAAGAGGCAGACCTCCACTTCCGCTAACCAGCTTGTGTCAAAACTCACAAAATCAGAGACTGAAGCCTATAAAGACAGTAATCAAACAACATGCAGAGCATTTTCATGAA
         Seq3 CCTATGTTGAGAGCAACTCTGACTCTGGATTGGACTCTAGTAATAATACATCTGAATTCAGTCAACCTTTAGCAGCCTCAACACCATGTCGAAGAGGCAGACCTCCACTTCCGCTAACCAGCTTGTGTCAAAACTCACAAAATCAGAGACTGAAGCCTATAAAGACAGTAATCAAACAACATGCAGAGCATTTTCATGAA

             1201                                                                                                                                                                                                 1400
             T  T  T  Q  V  Y  S  T  L  L  I  N  H  L  N  A  E  K  E  F  K  K  A  E  L  V  H  S  V  L  T  G  Q  Q  E  L  D  A  V  A  C  L  A  I  R  I  N  N  F  Q  S  V  N  K  Y  R  K  M  Q  Q  A  T  K  R  E  G  K
JXSR01S003992 ACCACCACACAGGTTTATAGCACATTGCTAATAAACCATTTGAATGCTGAAAAAGAATTCAAAAAAGCAGAACTTGTCCACAGTGTGTTGACAGGGCAACAGGAACTCGACGCTGTCGCGTGTCTTGCAATCAGGATCAATAACTTTCAGTCGGTGAACAAATACCGCAAAATGCAACAAGCGACAAAACGCGAAGGCAA
         Seq1 ACCACCACACAGGTTTATAGCACATTGCTAATAAACCACTTGAATGCTGAAAAAGAATTCAAAAAAGCAGAACTTGTCCACAGTGTGTTGACAGGGCAACAGGAACTCGACGCTGTCGCGTGTCTTGCAATCAGGATCAATAACTTTCAGTCGGTGAACAAATACCGCAAAATGCAACAAGCGACAAAACGCGAAGGCAA
         Seq2 ACCACCACACAGGTTTATAGCACATTGCTAATAAACCACTTGAATGCTGAAAAAGAATTCAAAAAAGCAGAACTTGTCCACAGTGTGTTGACAGGGCAACAGGAACTCGACGCTGTCGCGTGTCTTGCAATCAGGATCAATAACTTTCAGTCGGTGAACAAATACCGCAAAATGCAACAAGCGACAAAACGCGAAGGCAA
         Seq3 ACCACCACACAGGTTTATAGCACATTGCTAATAAACCACTTGAATGCTGAAAAAGAATTCAAAAAAGCAGAACTTGTCCACAGTGTGTTGACAGGGCAACAGGAACTCGACGCTGTCGCGTGTCTTGCAATCAGGATCAATAACTTTCAGTCGGTGAACAAATACCGCAAAATGCAACAAGCGACAAAACGCGAAGGCAA

             1401                                                                                                                                                                                                 1600
                                                   <---------------------Duplication at contig merging area---------------------------------->
             D  V  Y  Q  S  Y  F  A  I  K  K  A  E  R  L  F  L  P  G  S  L  E  N  C  I  F  T  V  T  P  P  L  T  H  P  P  S  G  G  S  S                       Contig merging area
JXSR01S003992 AGACGTCTATCAATCGTATTTTGCAATTAAAAAAGCGGAGAGACTTTTCTTGCCAGGATCGTTAGAGAACTGTATTTTCACAGTTACCCCACCTTTAACCCATCCTCCAAGCGGTGGGTCCTCNNNNNNNNNNNNNNNNNNNNNNNNNNNNNNNNNNNNNNNNNNNNNNNNNNNNNNNNNNNNNNNNNNNNNNNNNNNNNNN
         Seq1 AGACGTCTATCAATCGTATTTCGCAATTAAAAAAGCGGAGAGACTTTTCTTGCCAGGATCGTTAGAGAACTGTATTTTCACAGTTACCCCACCTTTAACCCATCCTCCAAGTGGTGGGTCCTC-------------------------------------------------------------------------------
         Seq2 AGACGTCTATCAATCGTATTTCGCAATTAAAAAAGCGGAGAGACTTTTCTTGCCAGGATCGTTAGAGAACTGTATTTTCACAGTTACCCCACCTTTAACCCATCCTCCAAGTGGTGGGTCCTC-------------------------------------------------------------------------------
         Seq3 AGACGTCTATCAATCGTATTTCGCAATTAAAAAAGCGGAGAGACTTTTCTTGCCAGGATCGTTAGAGAACTGTATTTTCACAGTTACCCCACCTTTAACCCATCCTCCAAGTGGTGGGTCCTC-------------------------------------------------------------------------------
```

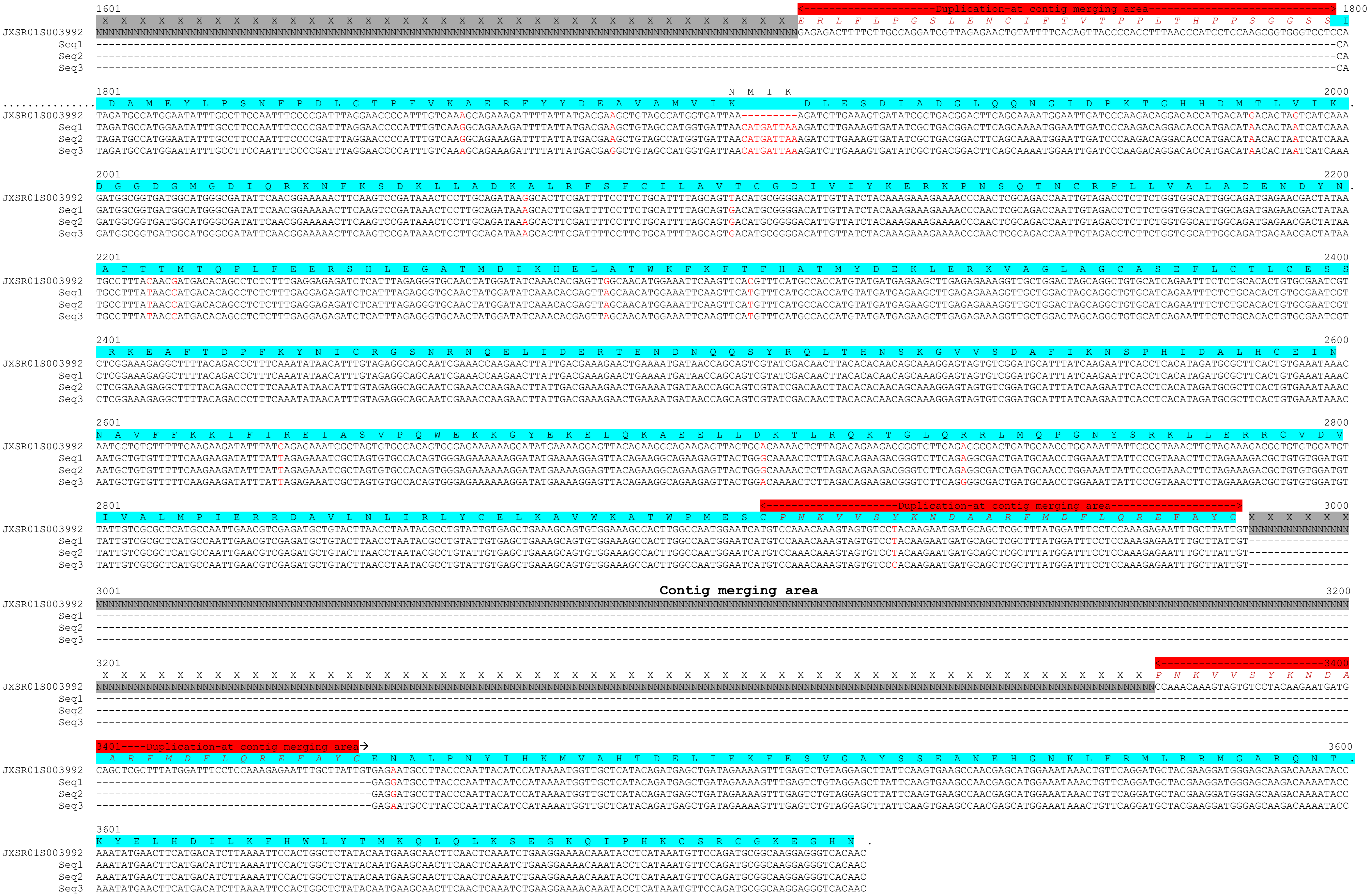

1601
1800
Duplication at contig merging area
JXSR01S003992
Seq1
Seq2
Seq3
1801
2000
2001
2200
2201
2400
2401
2600
2601
2800
2801
3000
3001
Contig merging area
3200
3201
3400
3401
3600
3601

#### FASTA DNA

```
>JXSR01S003992.1 Ophiothrix spiculata isolate Osp01 Scaffold13590, whole genome shotgun sequence
AATACATTACTGAAATAAAAATGTAAACAAGTTGTTTCGTTCCTGTTTTAGGGACAAAGCAAATTGTGAACAGCACAATCATTTTACCAGTTTATACAAATGTAATTACTAATCATCTCAAAAACTACAAGACAGATCAAGAATTTTTTTGCATCAAAACATGCGCAACCCCTAACTTGACGATTACATGGAACAACTTTATTGGGTATGTTGTATT
TTTGCGTGAAATCATGGGTCCATATGGACCCCCCCCCCATTACATGTAAGTGTTAATTTTTGAATTAAAAAATGTTTGTTTGTTTTTGTTTTTAAAAGTGTTGCAAACAAAATCAAAAACAAATATTCTAATTTTGATTTCTGGATTGTTCCTTGCAGGCAGTGATCTCAACATGGACCAGCACGTCAGCCACTTAGCGAAATGCTGTCGAGTATGTGC
AATTATGTTTTGTAGAATTCCAAAAGGCGGCAAAGGCTACGAATGTCTTGGACTTGACCCAGTATCTAAAATACCATGGAGAGAATTATTGCTAGAGTGCTTTGAAGTAGACCCAGCAAATGATGACGCAAATATCCATCCCAAGAAATTTTGCTGCAAATGTAAAGCAATTATAAAACGATTTTGTGATGCTAAAGTAAATCATCGACATTTTTCTCA
CTCCACAGAAGTAGTACATTGGGAAGCACACTGTGATGAATGTAAGGTTTGTTTACGCTTCTTAGGAAAATCTAGAGGACGGCATGTTAAGAAACATCGAAAGTGTGGACGTCCTAAAGCTTCATGTAGAAATGAGCAAATAGAAGAACCATTTATTCAAACATCAGAAAACTAAATGATCACAGTTATGTGTCTTTAGCAGCTCAGTTTATGGAAA
ACGACAGCCAAGTTCTACGGCAAATTCAACTGACCCGGATTTATCTTCAGATGTAGGCCTTAGCGACTCATGTATATCAGGCCTCACGTTTAATGATAGTTCTTTGAGTGTATGTGTGAACTTATCAACTCATGATCAAAACAACTCGCTATGTGGCAACTCCTATGTTGAGAGCAACTCTGACTCTGGATTGGACTCTAGTAATAATACATCTGAATT
CAGTCAACCTTTAGCAGCCTCAACACCATGTCGAAGAGGCAGACCTCCACTTCCGCTAACCAGCTTGTGTCAAAACTCACAAAATCAGAGACTGAAGCCTATAAAGACAGTAATCAAACAACATGCAGAGCATTTTCATGAAACCACCACACAGGTTTATAGCACATTGCTAATAAACCATTTGAATGCTGAAAAAGAATTCAAAAAAGCAGAACTTGT
CCACAGTGTGTTGACAGGGCAACAGGAACTCGACGCTGTCGCGTGTCTTGCAATCAGGATCAATAACTTTCAGTCGGTGAACAAATACCGCAAAATGCAACAAGCGACAAAACGCGAAGGCAAAGACGTCTATCAATCGTATTTTGCAATTAAAAAAGCGGAGAGACTTTTCTTGCCAGGATCGTTAGAGAACTGTATTTTCACAGTTACCCCACCTTT
AACCCATCCTCCAAGCGGTGGGTCCTCNNNNNNNNNNNNNNNNNNNNNNNNNNNNNNNNNNNNNNNNNNNNNNNNNNNNNNNNNNNNNNNNNNNNNNNNNNNNNNNNNNNNNNNNNNNNNNNNNNNNNNNNNNNNNNNNNNNNNNNNNNNNNNNNNNNNNNNNNNNNNNNNNNNNNNNNNNNNNNNNNNNNNNNNNNNNNNNNNNNNNNNNNNNNNNNNNNNNNNNNNNNNNGAG
AGACTTTTCTTGCCAGGATCGTTAGAGAACTGTATTTTCACAGTTACCCCACCTTTAACCCATCCTCCAAGCGGTGGGTCCTCCATAGATGCCATGGAATATTTGCCTTCCAATTTCCCCGATTTAGGAACCCCATTTGTCAAAGCAGAAAGATTTTATTATGACGAAGCTGTAGCCATGGTGATTAAAGATCTTGAAAGTGATATCGCTGACGGACTT
CAGCAAAATGGAATTGATCCCAAGACAGGACACCATGACATGACACTAGTCATCAAAGATGGCGGTGATGGCATGGGCGATATTCAACGGAAAAACTTCAAGTCCGATAAACTCCTTGCAGATAAGGCACTTCGATTTTCCTTCTGCATTTTAGCAGTTACATGCGGGGACATTGTTATCTACAAAGAAAGAAAACCCAACTCGCAGACCAATTGTAGA
CCTCTTCTGGTGGCATTGGCAGATGAGAACGACTATAATGCCTTTACAACGATGACACAGCCTCTCTTTGAGGAGAGATCTCATTTAGAGGGTGCAACTATGGATATCAAACACGAGTTGGCAACATGGAAATTCAAGTTCACGTTTCATGCCACCATGTATGATGAGAAGCTTGAGAGAAAGGTTGCTGGACTAGCAGGCTGTGCATCAGATTTCTC
TGCACACTGTGCGAATCGTCTCGGAAAGAGGCTTTTACAGACCCTTTCAAATATAACATTTGTAGAGGCAGCAATCGAAACCAAGAACTTATTGACGAAAGAACTGAAAATGATAACCAGCAGTCGTATCGACAACTTACACACAACAGCAAAGGAGTAGTGTCGGATGCATTTATCAAGAATTCACCTCACATAGATGCGCTTCACTGTGAAATAAAC
AATGCTGTGTTTTCAAGAAGATATTTATCAGAGAAATCGCTAGTGTGCCACAGTGGGAGAAAAAAGGATATGAAAAGGAGTTACAGAAGGCAGAAGAGTTACTGGACAAAACTCTTAGACAGAGACGGGTCTTCAGAGGCGACTGATGCAACCTGGAAATTATTCCCGTAACTTCTAGAAAGACGCTGTGTGGATGTTATTGTCGCGCTCATGCCA
ATTGAACGTCGAGATGCTGTACTTAACCTAATACGCCTGTATTGTGAGCTGAAAGCAGTGTGGAAAGCCACTTGGCCAATGGAATCATGTCCAAACAAAGTAGTGTCCTACAAGAATGATGCAGCTCGCTTTATGGATTTCCTCCAAAGAGAATTTGCTTATTGTNNNNNNNNNNNNNNNNNNNNNNNNNNNNNNNNNNNNNNNNNNNNNNNNNNNNNNNNNNNNN
NNNNNNNNNNNNNNNNNNNNNNNNNNNNNNNNNNNNNNNNNNNNNNNNNNNNNNNNNNNNNNNNNNNNNNNNNNNNNNNNNNNNNNNNNNNNNNNNNNNNNNNNNNNNNNNNNNNNNNNNNNNNNNNNNNNNNNNNNNNNNNNNNNNNNNNNNNNNNNNNNNNNNNNNNNNNNNNNNNNNNNNNNNNNNNNNNNNNNNNNNNNNNNNNNNNNNNNNNNNNNNNNNNNNN
NNNNNNNNNNNNNNNNNNNNNNNNNNNNNNNNNNNNNNNNNNNNNNNNNNNNNNNNNNNNNNNNNNNNNNNNNNNNNNNNNNNNNNNNNNNNNNNNNNNNNNNNNNNNNNNNNNNNNNNNNNNNNNNNNNNNNNNNNNNNNNNCCAAACAAAGTAGTGTCCTACAAGAATGATGCAGCTCGCTTTATGGATTTCCTCCAAAGAGAATTTGCTTATTGTGAGAATGCCTTACCCAATTACATCCATAAAAT
GGTTGCTCATACAGATGAGCTGATAGAAAGTTTGAGTCTGTAGGAGCTTATTCAAGTGAAGCCAACGAGCATGGAAATAAACTGTTCAGGATGCTACGAAGGATGGGAGCAAGACAAAATACCAAATATGAACTTCATGACATCTTAAAATTCCACTGGCTCTATACAATGAAGCAACTTCAACTCAAATCTGAAGGAAAACAAATACCTCATAAATG
TTCCAGATGCGGCAAGGAGGGTCACAACAAAAGACGATGCCCAGATTATATTGCGAACAGAGAAGCCTTTTAATATTGTACATTTTGTACATGTAAATATGATCAGTTTGGATGGACAGACTTTAATGTACACCTAGAGTTTAAAAGTCTTAAATATTGGTCTACATGTATGTGAATGATAATCATGTTAACCTAATTTGTGCGATACTTTGGAAAAGA
TGAAACATATTCCAAAAATCAAAATCATAATTAAAGCATTTCATTTTTGTCTACTGATTGTGAAGTTAACTTTTCATGCAAAGTGTCTAATAGAGTTGGTTCTATGTGATAATCATGCAAGTGATGGAATGAAATATAATAGACTTACATGTAAACATAATAGTTTTGGTATTACGACGTTTTGTTTATTAACAATTAATAATATACAAGCTATCATT
AACCTCATTTGAGGCTTTGAAAAAGTTTAGTCTTAAACAACAAAATAATTGCAAACACTAGCTATGTAGACTAACTGAATTAAAATCATTTCAGTTTGTGTGTTTTGGTTGTGAAATCATGAAATTTTCAGGTGAAATGTACACATTGTTAATGTTAACATGTCAAACTTAATACAAACTAAAACAGTTAAATTTAAGTGATATCATCAGCCCAACCTG
ACTTTGGAAAGACTTGGTTAAGTCTACAAGTATTTTAACAAGATTAAAAGGTACTATTGTATACTAAGATGGACTAGGACTTAACTAAAGGTATTTTAGTATTTGTCTTCATGTTGAATAATTTCATTATATGAATTGTGCAGAATTGTCAGTTATGAGGACTTTGTTCAAGTCTACATGATAGTATACTGAACATGAATGGAATATGTGCGATAGAC
TCAAACAAAGACTGCAGATGTTTACTTGTTTNNNNNNNNNNNNNNNNNNNNNNNNNNNNNNNNNNNNNNNNNNNNNNNNNNNNNNNNNNNNNNNNNNNNNNNNNNNNNNNNNNNNNNNNNNNNNNNNNNNNNNNNNNNNNNNNNNNNNNNNNNNNNNNNNNNNNNNNNNNNNNNNNNNNNNNNNNNNNNNNNNNNNNNNNNNNNNNNNNNNNNNNNNNNNNN
NNNNNNNNNNNNNNNNNNNNNNNNNNNNNNNNNNNNNNNNNNNNNNNNNNNNNNNNNNNNNNNNNNNNNNNNNNNNNNNNNNNNNNNNNNNNNNNNNNNNNNNNNNNNNNNNNNNNNNNNNNNNNNNNNNNNNNNNNNNNNNNNNNNNNNNNNNNNNNNNNNNNNNNNNNNNNNNNNNNNNNNNNNNNNNNNNNNNNNNNNNNNNNNNNNNNNNNNNNNNNN
NNNNNNNNNNNNNNNNNNNNNNNNNNNNNNNNNNNNNNNNNNNNNNNNNNNNNNNNNNNNNNNNNNNNNNNNNNNNNNNNNNNNNNNNNNNNNNNNNNNNNNNNNNNNNNNNNNNNNNNNNNNNNNNNNNNNNNNNNNNNNNNNNNNATTTCTGATACTTCAGCATGTTACCTTTTATTACAGACAATAAAATA
CTACAAAACAAAACTTTTAAACTTATTTATAGGTAAAATTAAAAGACGATTTTGACATTTACAGTTAAAAAAACACGTTGCTGTTGGCGGGAAAATACGGGCATAGATACGCGTCCTATTTTTTGCGTTTAGGTTGCGTGTTTTGATGTGCGTTCCTTTTTGCGCGGCAACAAGTACTGCGTGCGTGGGTGAACGCGTTGCGTTATGTTCTTGCATGA
GCACATTTGTTAAAATCCCGCAATAAACTTGTCAATATTGTAANNNNNNNNNNNNNNNNNNNNNNNNNNNNNNNNNNNNNNNNNNNNNNNNNATAAACTTGTCAATATTGTAAACAAAGACCAAATAATTGAACATTGGTATTTTAGATTTCTTTCAATATTTACAAAATAAGGAGCTTAAAATGGGATCTGAAAGTTGAGCATAATACTTACCAATA
ATATCAATTACATAATAAGTTTTCATGTCCGATACCCAAGGATTAGACCTCAAAATTCATGGGTTTTAGGTCAAAATGTGCAATTTCTTCATTAAACATGGGATTTCTCCTTTTCAGATTTGATGAAAAACATCATTTTTTCGGGAAACGTAACTAAAAAAACTAAAAAGTACAAGAAAAGCTTTTTCAACCATTATGTACAGTATATCAAAATCTAG
GATCGCGTTTCTCATTTTTAATATAAGGTCCTAAAATCAAAGATATTCATTAGTTTCTATCAAAATTGATGCAGATAAATGTCGGAAACGGCACCTTTAACATTGTTTTCATTATCTATCGAAAAATTCGGCTTCATTTGAAAAAATAAAAAATGCGATCCTAGATAATTTTGTGCAGTTTCCAAAAAACGATATGTGAAAAAATGGCATTAACAATT
TTTTTAGTTACGTTTCCCGAAAAATGTTCAAGCGAAATCACTGTTTTTTGCTATTTTTTCCAAATATGGTCATTTTTGAACCAAACCAATAGCATAAAGAGATTCCTTGTGAAATTTCCTTTCTAGAAATGTATACTTTTCTATACTGTAACTTAATATTTTTAGACTTATAAGCCAAAAAAGAAAGTTAAAATAAAAGCCTTAACAGGCCCCTTAAGA
AGGTACAGCAGTATTCAAATAACGTATAGCTTATTGTGCCATATACTGTATAGCCATCGAATAGTGTATACTTCTATGTGACTGCAGCCTTAAAACAATGCAAGTGATACTTGGCTTTAAATTTGTACGTACAAATGTGTTAATTGTCACCTTGATATAAAAACAGTATTCAAATTCTATTCTTGATGAAATTTTATTTTGGTAACTTTTTTGTGAAT
AAAAAACAGAAATAAAAGACAAGTAGTAGTGATTTTTCTTGCTTTTAATTAAGTTGTATTCTGCTGAATTAGGATTGATTGTTATGTTTTCAGCTTCTTATTGCCTATCAATCTTCCATGCTTTATTAAATGCCAACACCCAGTTTTGTTTTGAAAACAGTCAATTTCCAAGCATCTAAGAAAGTAGTAAATTTTCTGACTACTTTGGGTTGCCGT
GACCTTTTTAAGGTACCGATCCCGGAAATAAAGCAAAAAAATTTCACTACTTCTGTTTTTCGCATCCAATATTGCCATATTTTTTAAGCTTACACCAGGAAGATTAATAGGAACAATACTTTTGTCCGAAACTTAATTATATTCTTCCTTAAAATTTTACCCACATATGCCCCAGTAAAATGAAGCCAAGGTGACGAAAAAGAAGACCCTGTTTTATGG
GCGGGCGGACCTTTCCAAGTAGGGTCCTAAGGCAAATTTTTTCTAAATAATTTTTTTTGAATAAAATTACCCAGAAAAAAAAATGGGAGAAAAAATCTTATTAAGAAATTTCTTTTAACAATTTTTAACCAAAATCATTGAATTTTTAGGGTCGGTCGGGTCCGTAAAACAGGGTTGTTTTTTTCGTCACCTAAAATGAATCAAATTTCACACATTGCAT
ATTCAGGAATTTTTCACTTTATTTTAATTTCATTCACACAAAACAGACACTGTACTTTTGAATCTGATTCTATAGATAGTCGTGCAATGATGGATTTGACATAAGTTAAGAAGTGCTTACACCAAAATCATGAGCTAAGACTGAAAATTGGCTAAATAAAGCCAGGAAAAGAATCTGTTTAACACTGTTTCAAAAACGTGTTGCGGGAGTTGTTGCGCC
AATGACGCGCAACACTGATCACAACTAATTGTAATAATTAAAACACTGATTATGTACATTGTATGGAATCAGTTGCAATGTATCCCTTAATGCAATTTTCGACTTTTTGAGTCACAAAATACCCATCGGTGGAAATATAGGAATTTTTGTAGGAATTTGTTTATTTGATAACACAATTTTCTGGGTACTAGTCTTGTTTCAAATAGACCATTCCTATAA
TTCACATATTTTTGTAGCTGCACTCGGTGCACGGAGGGATATTGGCACTCATAATGAGGCCAAGTATGATATCAGGTGATGCCCGATATGACTAATTCTTTACGTCACTGTGCATCACTTGATGTCATACTTGGCCTCGTTATAAATTCAATACACCGGCGATGGCAAGAACATGCCAATTATAGGAATGGTGTATAAAAATAAACAGCTAGTTTTATT
GGAGCATAGATTTTTAAGTGTCTTAACAAGCAATGCAATGTCCAGTAGCAGCAGCACAATTGACGTGATTCCACATGTAGCATAACTTACATTGAATCCATTTCAACTTAGATTGTTTAACTTGACTAAGTTCATTGTATTTACAGTGTTCGTTATTACAAAATAGTTCGCTATCATTCTTCATCGTCGCTACTACTGTCAGAATCAGAGAGTCCAAACT
CCTGCGTCAACTCAGTTTCACTGCAAACGTCACTCTCTAAAGCTATGGACTCTGTCGTCTCAGCATAATCATCCACTTCTGCATCTTGCTGTCTATCATTACCAGACTCGGTGTCGGGTTGTAACTCTAGCTCAGGTTCTGACTCTGGTTCGTCTTTGCTCAGACGCAGTCATGTGTGTAGTGGCGGTCAATGTCCAGATTGATTTGTCCTCACACCTAA
AAAATATTCAGAGTTCTTAGTCCATCACGAATAGCATAGCATGGGTCAAATGTATTTGACAAACAGTCACTTTCAATTTGCAAATGTCCTTGAAGTAGAGATGGGATATAAATGCGTCGGATCAAGGACGATGGCATGGCCAACGCAGCAGGAGACGGTGCCTCAACGCCTCCGAGAATGTATAAGTAATCGCCAAACTGTTCTGCAGTATGTGACGATA
GGAACAAGTTGGCATCAAACTTTTAACGATAGTGCTACGTAAATTCCCAGTCAGTTCAATTAAATGAAGCTGTCCTTCACGAACCATAGAGTTGTTCTCTAATTTGCAGCCACCAATTATAACCAAACGGTTCTGTCCCAGATATGTAGCTGAATGATACGCAATGTTAGGCATATCTACAAGCCGAGTTCTCCACTCACGATCATTTAAGTTATATG
CATACATGTTATTGTTCACATGAACAAATTGGCCATCAGAAAAGTCCAAGCCACCATATACAATGGCTGTCTCTCCACACTTGGTTAGAGTATGCCCAAATCTTGGTGTCGGTGTGTAGCCGGTCTGGTTAACACACTGCCAGTGAAAAAATGATCTTCCTTCAAACTGTCCACAAACAAAATCATCAGTTGCATTTTGTCCAGATCTGGCCCATTGTC
TTCCTCCGAAGATGTAAACACGATTCAGGTGTTCAAACGACGCACTGCCAACAGTTGGTCTAATTAGCCTGGTTTTTAGTCTTCCAGATGTTGTGAAATGAAATCGTTGCCGTCCGCTACTGCCACTGTTATAAGATAGTGGAAATCATCGTGGAAGTTCAGTCCTCCAAAAAACATCTCTCGTCCGTTTGCCTTCCAGCAGACCCCTCGAGAAGAAG
GGAATCTATCATGTGAATCACAAACGTATAACTTCTCAAAAACACAATCGGTAAAAACCGAAGCCATATTTAATGTAAAAATCGAGCTGAATTGACCGAGATCGTAACACAACACCACTCTTGTCGTATTCGCTATTGTTTGTTTACATCCGGTGAGCGATGTGATTTAACGGTTGTCAAGTAGCAACCTCGTCCAAACAACAATTTTTGTATACATTT
ATTTCAGTTATCCTTTTAAAAAATAAGCAATTTCATGTTTTTATAAAAGTAAAATGAAAACACATATTTTGTGATATTATATGAAGATTTAATATAGTTATTATGATTTTTGAAAATAATACTGGAAATGAATAAACTAATTTTGTGAACGAGGCGCAAACATCGCGAAGATTCAAGATGGAGGATTTTTCACGGACACATACGGGAATAGTTACA
GTGTGCATGTGCGCCCTATTAGGCTATCTAAATGAAAGTTTGCAAAATCTGGAAGTGATTCAAAAATGAATCACTGTGCACTTTTAAGTGTTAAAGGTGACCTAGCACCGAAATGGTGTCGGACATGAGTGAGCTGATATCAGCAGTGTTGGACACAATTTCAAAGTATTTATCCTGTGCTTTTAGGGTACTCTCTGGCATACTTATCGTGCGCTTTAA
TACAACGCGATCTTACACGTGCGTCATTTTTATATTCACTTATGCATGAATATAAACGCATAAGACCATGCGCATTACTCATGACAGTGCAGGGCTCAAAGCGCACAGTGCTTAAGGTATTAGTCATTGCTAATAAAATATCAATTTTTTTGCTTAATTTTTCTTAATATCTCAAAAAGTAAGTCGGTGACCATATTTTTTTATTACGATATTTTAAAG
AGCTATATAAGGACCAAAACATATGTGAGAAAAAGTGAAAAAATAGAGCCGTTTTCAATTATAAGCATTTTTGTCTAATTATTTTGATTTTACCATAAAAGTCAATGGTAAAAATACTGGTTCTCGGAACCACACGATCCCCCTGGCCGACGCGATCAGAAAAATGGAAAGTTTCGGGAAATTTGACATCATATGACCTTTGACCTTCAGATGTGACC
TTTGACCTCATCAACATTTGAGGGTTCTTATATTATATGCACAAACCAAGTTTGGTTCCAATCAGACTTCAACTTCTCAAATTAGGCCAAATTGACATTTTCAGCCTATTTTACAACTTGACCTCAAATGACCTNNNNNNNNNNNNNNNNNNNNNNNNNNNNNNNNNNNNNNNNNNNNNNNNNNNNNNNNNNNNNNNNNNNNN
[...]
```

#### FASTA protein

```
>OspRAG1L-A.1
MDQHVSHLAKCCRVCAIMFCRIPKGGKGYECLGLDPVSKIPWRELLLECFEVDPANDDANIHPKKFCCKCKAIIKRFCDAKVNHRHFSHSTEVVHWEAHCDECKVCLRFLGKSRGRHVKKHRKCGRPKASCRNEQIEEPFIQTSENLNDHSYVSLAAQFIGKRQPSSTANSTDPDLSSDVGLSDSCISGLTFNDSSLSVCVNLSTHDQNNSLCGNSYVE
SNSDSGLDSSNNTSEFSQPLAASTPCRRGRPPLPLTSLCQNSQNQRLKPIKTVIKQHAEHFHETTTQVYSTLLINHLNAEKEFKKAELVHSVLTGQQELDAVACLAIRINNFQSVNKYRKMQQATKREGKDVYQSYFAIKKAERLFLPGSLENCIFTVTPPLTHPPSGGSSIDAMEYLPSNFPDLGTPFVKAERFYYDEAVAMVIKDLESDIADGLQQN
GIDPKTGHHDMTLVIKDGGDGMGDIQRKNFKSDKLLADKALRFSFCILAVTCGDIVIYKERKPNSQTNCRPLLVALADENDYNAFTTMTQPLFEERSHLEGATMDIKHELATWKFKFTFHATMYDEKLERKVAGLAGCASEFLCTLCESSRKEAFTDPFKYNICRGSNRNQELIDERTENDNQQSYRQLTHNSKGVVSDAFIKNSPHIDALHCEINNAV
FFKKIFIREIASVPQWEKKGYEKELQKAEELLDKTLRQKTGLQRRLMQPGNYSRKLLERRCVDVIVALMPIERRDAVLNLIRLYCELKAVWKATWPMESCPNKVVSYKNDAARFMDFLQREFAYCENALPNYIHKMVAHTDELIEKFESVGAYSSEANEHGNKLFRMLRRMGARQNTKYELHDILKFHWLYTMKQLQLKSEGKQIPHKCSRCGKEGHNK
RRCPDYIANREAF

>OspRAG2L-A.1
MASVFTDCVFEKLYVCDSHDRFPSSRGVCWKANGREMFFGGLNFHDDFPLSYNSGSSGRQRFHFTTSGRLKTRLIRPTVGSASFEHLNRVYIFGGRQWARSGQNATDDFVCGQFEGRSFFHWQCVNQTGYTPTPRFGHTLTKCGETAIVYGGLDFSDGQFVHVNNNMYAYNLNDREWRTRLVDMPNIAYHSATYLGQNRLVIIGGCKLENNSMVREGQL
HLIELTGNLRSTIVKSFDANLFLSSHTAEQFGDYLYILGGVEAPSPAALAMPSSLIRRIYIPSLLQGHLQIESDCLSNTFDPCYAIRDGLRTLNIFRCEDKSIWTLTATTHMTASEQDEPESEPELELQPDTESGNDRQQDAEVDDYAETTESIALESDVCSETELTQEFGLSDSDSSSDDEDDSELFCNNEHCKYNELSQVKQSKLKWIQCKLCYMWN
HVNCAAATGHCIAC
```

### OspRAGL-A.2

| Genomic loci | RAG1L status | RAG2L status . |
|---|---|---|
| OspRAGL-A.2 | not found | potentially functional (1 CDS) |

**Scaffold: JXSR01300694.1**

#### FASTA DNA

```
>JXSR01300694.1 Ophiothrix spiculata isolate Osp01 contig_300694, whole genome shotgun sequence
[…]
ATGAGATACGTACATTGTGCATCAAATGTTGTCACAACTTTTCAATAATCGTATGAAATGTCATAAATTTGGTATGATTTTGAAGATAATGTATTAGCCTATAAAATCAATTGAAAATATTAGCCAAATAAAGGTATTAATTAACTCTAATCAACGATTACCAGGAAGAAGTTAGGTATGGATGAAGTGAATGAGATGAAACGCAAATGGGCCTCACCT
CAAAAAGGTTGACCTTATGTAATATGTAATATGTTTATTGTTTTACTTACATATAATCGATTTATGTCGCTTTTCGGACTTTAATGTGATCAATCACAGAGTAGATCAATCCCTAATTTTACACACACCACTGGTCAAGAGGGGGTCTGTATATATACCCAACATTTAAAAGGCGCTTTGTGCATAGTTCGATTAATAGAACCCTTGTTTTTTTCGGGT
GACACAATCCAACATCATGACATAACTAAACAAGTCACACAATAACGAGTCCCGTAGACTTCCGTGAGAAAGTCGGCCATTTTGATATTCGCCACTTTGCAACCTCGTGCGCATTCATAATTTTACTGAGTCATTTTGTATCATAATAAATCTGTATATAGTGTTGGTAATGTTTATCAATATATTTTATTTATTATTCAATAATATTTGTTAACACTG
CAAGTCATTTTAACATTTCAAAATATCATAACTTTTCTTTATGTAAACATTATTTTGACCATGGATGTCGTGTCAATCAGAGGTCATCCACCTTAGCAACGGAGATAGTTAAAATCCATCACTCACAGGAAGGTGTGTTTATTCTCTCGTTCCGGTATATTTCTTTCAAAAGCTATCATAAAAAGTGTCCTTTTTTGTGTTTCTTCGTGATATGGCCAA
CGTTTTTTCGGATTGCAACCTGGAGAAAGTTTATGTTAGTGGAGCCAACGAACCATTTCCATCTCCTCGTGGTGTATGCTGGAGTTTAGCCGATAAATTTCTGTTTTACGGCGGCTGTAATTTTGATTCGGATTTTCCGCTAATCTACAACGTTAATGCACGCCAAGGTACAAGACTAAAATCTTCATCAAGTGGAAGAACAAAGACTCGACTGATAAG
GCCAAGTTTAGGGAGTTCTTGCATTGAGCACGAATCGCGAGTGTATGTATTTGGTGGGCAAAATTGGAAGAGATTTGGACAACATGCTAGTGATGATTTCGTTGTTGGTCAATTCAAGGGATCGGCTTTTTGCTGGGAGACTGTAAACCAACGTGGTTATACACCATCACCCAGATTTGGACATACATTTACCAAATGCAGAAATTTGGGTGTTGTGTA
CGGTGGATTAGACTTAGTGGATTCAAAATTTATTCACGTAAACAACAATTTCTATTTTTTCAATCTCATTGATAAGAGTTGGAGCACTCGACCAGCACCATCTGGCTTGACTGATTTAGCACACCATACATGTACATATGTAGGACAAAATACATTTATAATTATTGGTGGGTGTACCCTGCGTGGAAACTATATGCTTCGAACAGGAGGAGTTCATGT
CATTAACTTCAACGGAACAACTAATCTTCGTTTGTCATCCTATGAATCAGAACTGCACGTCTCTAGTCATTCAGCAGAGCAGGTTGGAGACTTCTTGTACATATTTGGAGGGTTTCGAGCTGATAGTGAAAGGGGGATTGCATTACCATCCCGTCATATTCATCGAATCCATTTGCCATCAATTTTGAATGGAAAACTAGTTATTGAAACTCATGCCTT
GGCTGACCATTTTTCACAATGTTATGTAATGAAACGAGGTCTCCGGACATTCCATGTGTTTAGACTTGAAGATCACTCAGTCTGGCAGTTTACTGCTGCCACATTCTCTGATGATCTTGTTGTTAACGATGAACAAATAAACGGTGATCAAGAAGTAGAAGATGACATGATTGTGGTCATGGCCGTTACTCCGGCACACAATGATACACCAGATGATGT
GACTATGACTGAACCTGTGCCTGCTGATGATGACAACGATAATGATTTTGTTATACAAGCAGATGGATCTGATCCGGGATCTGAATCGGAATCAGAGTCGAGTGACGAAGAAAGTGAATTATTTTGTAATAATGAAAACTGTAAATATGCCAATTACAATAAGGAAAAGGCTTCAAAATTAAAGTGGCTCCAGTGTGGTAACTGTTGTTTGTGGAACCA
CGAATCTTGCTCCGGAACAGATGGGACATGTCTTCAGTGCTATTAATTAAATAATGAACTGCATGTAGTGTATCTCTAATGAAACATTGTACAAACAAGACATTCCAAAGTTCATATAATCAGGAAATATCTGCCTATAAAAAAATGTTGTTTTAATTTGCCATTTACCGACCGCAATTTCGTGTCAAAATAAATTGATAAAAGTCATCTTTTGAAATT
GACTTTTGGCAAGCGACTTGATACGACTTGAACATGTTCACACTAGTTATAAACAAGAAAAAATAAATCGATGATGACCTAAGTGAAATCGATGATGACTTTTTGCATTGTGTGAATGGGGTTAATAATTGCATGAGTGAACAAGAGGTCCTGTGGGGAAACCCAAATTTTCCCGGACCGGACCGAAACCGGTCGGGGATTTTTTTGAGCTATTTTCGG
GTTTGGTACGGTCCGGGAAAAATATGATCGGGTCGGGGAAATTTAACCTAGAAAGCCACTTTACTACCTACCCCGAAAAAAGGATTAATAGAACCCTTGGTTTTTTTTCGGGTAACACAAACCAACATAACTAAACAAGTCCAGTAATCCAGAAAGCCACTTTACTACCTACCCCGAAAAACAATTTTCGGGAAGGGCGGTCAATCTGCCATGTAATCA
GTCTGCTATGTATAATTTTGACTTTCTAGTAAAGTGGTATGATGTGATAGTGGATTTATATTGTGGATAGTAACACAGAAAAAATCAAGGTCCTAGGATCGGACTACTTTGTGCATGGGGGTAATGTTGAAACCCAGGTTATTAAAGCAGGGATAGACCTATTACTGGCAAAATCTCTTTTATGCTCAAATTATGTAATTATCAAATAAACCACATAAT
TGTGCACACTTCCAGAAGCGGATTTTTATGCTGAGCATGATGGCCATGTCCTTTTTTTACCAAAATGGTAAATGTTTGCCCCAAACTATTCACTGAAAATTTTGTTTCATCATTTGACAAATAATAAGTTATCATGAAAATGGTATTCACAAATTCAAAAAGAATTTCTTAAATTTCAAAGAAGATACATAATATTTATAATGTATAAAACATAATTTG
TTAAATATTTTGTTAAATAGATTCATATGGCAATGTAAATTTAGGTAGGGTAAACAAGGATATGCTATGAGGTAA
[…]
```

#### FASTA protein

```
>OspRAG2L-A.2
MANVFSDCNLEKVYVSGANEPFPSPRGVCWSLADKFLFYGGCNFDSDFPLIYNVNARQGTRLKSSSSGRTKTRLIRPSLGSSCIEHESRVYVFGGQNWKRFGQHASDDFVVGQFKGSAFCWETVNQRGYTPSPRFGHTFTKCRNLGVVYGGLDLVDSKFIHVNNNFYFFNLIDKSWSTRPAPSGLTDLAHHTCTYVGQNTFIIIGGCTLRGNYMLRTGG
VHVINFNGTTNLRLSSYESELHVSSHSAEQVGDFLYIFGGFRADSERGIALPSRHIHRIHLPSILNGKLVIETHALADHFSQCYVMKRGLRTFHVFRLEDHSVWQFTAATFSDDLVVNDEQINGDQEVEDDMIVVMAVTPAHNDTPDDVTMTEPVPADDDNDNDFVIQADGSDPGSESESESSDEESELFCNNENCKYANYNKEKASKLKWLQCGNCCL
WNHESCSGTDGTCLQCY
```

### OspRAGL-A.3

| Genomic loci | RAG1L status | RAG2L status |
| --- | --- | --- |
| OspRAGL-A.3 | not found | potentially functional (1 CDS) |

**Scaffold: JXSR01S006948.1**

#### FASTA DNA

```
>JXSR01S006948.1 Ophiothrix spiculata Scaffold16250, whole genome
[…]
GCATAATTTAATTGACTTATATGTGTTACAGATAAACAGCTGTAGTATAGAATAGGACCAGGATGAAATTTCAGTTGAGTATGATATAAACAAGAGTAACCAAAATTAGATCAGTAAACTGAATTGTCGGATGATTCAATAACCTTTGCGAGAAAATGATTTGAAACGTTGAGTGTGATTTATTGTCTTCCCATTGTGACACAGGTGAAAATATACCGT
AGTTTTAAAAATATAAATAATTGGTGTTTTTCTGTTTCTCCATGCACTTCAATTAGTTGTATGATGCTAACTCCGTTAATACAGATGGTGTGAGTTTTGGATGATACAATTGCATGAAGATTTATATTTTCAAAAAATCGCTTATTACTGTTAAATGTCGTTGATAGTTGAGACATAATAGGCGCAATGTAAGCAAAGAATAAGAATGTAAACAAATGT
GTGGCGAGTCAACATTAGGAATTTTGACCGATTGGTCACACAAAATGACATCATTCTTGGGGAGCCAAGATAAGAGAACCCCAAAAGACTTAAACAAGGTAATGTACGAGTATATACAGTAGGAATATACGATTACACACAGCTTTACTCTTACACATGTAATAATATTAAACAGTTTAATAAAAACTTCTATATATTTCATTGGCCCGTATTCCAGAA
AGATGTTTAGCTAAACCATTGTTTAACTGAGCATCACACAGCGTAACACACCAACACCGCGTACGTAAATAAATCGCCATCTTGGTTTCCAATCCCAGATTGCACCGCGGCCGATCCTCGTATTTTAAAGCAAATTTATAGTTTGTTTTGAAACATGTTTTACACGATTTTTTGGTCAAATCAATATCTTAAAAAATATAATTATATCTTAATTATATT
TATTTAACAATACAATGACTATTTGATAATTTTAAGTAATTAAAATTATCACATAAAATTAAATTTATCGGACATGGCCGATAGGTCGCGAAGCCGGGTCATCACTAAGTAAATAAACAAATCTATCGCCTCAGATCAGATAGTACAAGTCAGCTGTTGTTTTGATCGCACTCGCATGAAATCAATTGCGATCCACAATGTCAACAATTTTTAGAAACT
GTACTTTATCTAAAGAATATGTTGCAGGCCAAAATGAAACATTTCCATCTCCACAAGGTGTTTGCTGGAGTAGAGGGGATAATACTTTATTTTACGGCGGAGCGAATTTTAATCATGATTTCCCGATCATGTATAAGCCTGGATGTGCGAGTGGACAACGATTTGTTGTCACTACCTCCTCCAGGTTGAAGACCAAGATTCAGCGGGGAACTGTAGCTG
CAGCAGCAGTATGTAGACAGGATAAAGTTTTAATTTTTGGTGGTCGCCAGTGGAAACGGCCTGAACAGCCTTCAACATCATCTTTTCTACAGGGGAATTTCCAAGGACAATATTTTGCCTGGGACTTTGTCAATCAATTTGGATGTACACCATCGCCACGATTTGGACACAGTTTTGTTTCATGTACGAACTTTGAATTTCTTTATGGTGGATTAAGCA
CAGAGGATTTTTATTTTCATCATGCTAATACTAATGTTTTCACATATTCCACTTTGAATAAATGGAACACTAAACCATCAGAACTACCTCCCCTTGCTTACCACACTGCAACCTACATTGGGCATAGTACAGTGGCCATCATTGGAGGTGTGCAACTACACACTGCTACAAATAGAATGGTAAGACCTGGACATATCCATGTTGTGAGGTTCTCCGCAA
GTGGATCAGAGACTAAAGCTATAACATTGGATGCCAGACTTTTTGTTTCTGGACATTCCTCTGTGCAACTCGGCGACTTCATTTATGTTTTTGGTGGTTATCGGGCAACGTCAAAAGATGGCGATGCTGAGCCGTCACAGAGTGTTAGTCGCATCCATATTCCATCCCTTTTAAATAGAAATCTAAACATTGAACATGACACAGCCCCAAGTTCGTTTA
CACCAAGCTATCTGTATGTCAGTAAGCAAGGCCTGCGGGAAGTCAGTATCTTCAATGTAGAAGAACGGTCGATTTGGAAATTTTCAGGACAGAGACAAGCAATTGATCAATACTCTGTTATTGATCGGGAGGATATTGCGGACATAGCAACTGATGAAGCAGCAAATGCCCCAACGGAAGAAAATATGGCAAATGTTCAAATTGAAGCTGCAACATGTA
ATCAACAGGCAGATTCCGACAGGGATTCGGACATTGATTATGACTTGCCTCTTGACGATTCTGATACATCAGATTCAGAAAGTGACGATGATGAACCTTTGTTTTGTAATAATGACTCTTGTGAATATTCAAAATTAATCACTACAGGACAGAAGAAGCTTACTTGGGTTCATTGCTTTAACTGTTTGGACTGGTTCCATGAGCGTTGCTTATCAGGAA
TGATACGCATCATTGCAGCTTTCAAGCAGAACAGTCCAATGAGTCGTGCTGTTGACGAGAAATAAAACCTTGATTTAGCTAAACAACTTTCTGGAATACGGGCCAATTTAGTTGACTAAGATTTTAACATAATTTTAAATAATAGTATATCCATCCAAAAGTACAGTATATGTTTCATCATCATGAAAGTGTCGATATTTGTTTCTATTTGGAGCATGA
ACCGTATTTTACTCCCAATCGACAGAAGGGGGATTTTGAGGAGTAAAAATAGAAAGGATGTCAACCTGCTGTTCTAAGGATGTCTGTTTAACCCCTGCAAAGATGCAAATACCATATCTTTCAAACATCCGCTGTGCTACTGTGATGCATTAGCTTAGCTCAAATGAAATGCATGCTTTTGTTAATTACAAGTAAACTAAACACCTTTGTTGTTAGTGA
GATAGTTTACTAAAGATTTGTACTCAGGTATTGAATGCAGCTAATATTGATTATAACTATATCTTGGGTGTTATTGACATGTTTTCTTGACCTACTCAATCGATTTCCTTCCTTGTCATTCTTTCGTGTACCTTGGTCAATCACGGGGAAATTGCATCGATTCTTTATTGTAGGCAGGTGGCAATCAAGGGCACTGCCTTAGTTTAGTTGATGAACTGA
AGGGATGTCTTCACATTTTAATGACAAGTAAAACAAATCAGAGAATTGAATGCGCCAGAATAAGAACATTCCAAGTGTAATAGCTCCCTTCTGTTAAGTTCTGTAAATATAAATGCATACAAATCTCTGAAAATGCTTTCAAATTACTAAAAAGCATTTCCATGTCTTTATCCCAATCAAAGTGTCACACTTGACCTTGTAGTTGAATTTTTTAGACAT
TGTTAGATAATTTTCTAGCATATAGTTGCAGAGAGGAGCTTATGGGGTTTTCTTTTAATGGCTGTTTAAATATAGCAACTATGTACTATAACAATTATTAAGTAGCAGCACAGAATACGGTAGAGCAATCATTATTGTCTTTATTTTTATTTTATATTTTCCAAAAATATCTTCACATGCACCTTTGTTTTAAACAATTCAGTTATTCAAACCACCGCG
AACGTAATTCTGCACAAATTGTGTTCAGCTGTCGTGTGCACAAATGAAGGGCGTCCAACACGAGTC
[…]
```

#### FASTA protein

```
>OspRAG2L-A.3
MSTIFRNCTLSKEYVAGQNETFPSPQGVCWSRGDNTLFYGGANFNHDFPIMYKPGCASGQRFVVTTSSRLKTKIQRGTVAAAAVCRQDKVLIFGGRQWKRPEQPSTSSFLQGNFQGQYFAWDFVNQFGCTPSPRFGHSFVSCTNFEFLYGGLSTEDFYFHHANTNVFTYSTLNKWNTKPSELPPLAYHTATYIGHSTVAIIGGVQLHTATNRMVRPGHI
HVVRFSASGSETKAITLDARLFVSGHSSVQLGDFIYVFGGYRATSKDGDAEPSQSVSRIHIPSLLNRNLNIEHDTAPSSFTPSYLYVSKQGLREVSIFNVEERSIWKFSGQRQAIDQYSVIDREDIADIATDEAANAPTEENMANVQIEAATCNQQADSDRDSDIDYDLPLDDSDTSDSESDDDEPLFCNNDSCEYSKLITTGQKKLTWVHCFNCLDWF
HERCLSGMIRIIAAFKQNSPMSRAVDEK
```

### RAGL-B

#### OspRAGL-B.1

| Genomic loci | RAG1L status | RAG2L status . |
| --- | --- | --- |
| OspRAGL-B.1 | frameshift / potential splicing | incomplete (Kelch domain only - 3 CDSs) |

```
>JXSR01S003628.1:5000-12500 Ophiothrix spiculata isolate Osp01 Scaffold13262, whole genome shotgun sequence
TGTATAAGCAGCTATTGTAATAAGGCCAGTCATGATAATGAAAAAAGTGGTCTGTATCAGGTTCCACCATTTGCTTCACTTGTTAAAATGCATATATGTACTTTTGAAATGTTTAGCCATGTTTTTTCTTTTCTTTGCACATATTAAAAATAACAGGTAGTGATACTTAGAAAAACTTTCCTTTTGTGTTGGGTTGGCGAACCCTGTGGACAACTCTCC
AAACATGATAGTGTGTCAATTTTTAATTTGCAGATCATGGCAGAGAGCAAGTTGGCACCAATCCAGGTTCACATGGAAAGGCTTATGATGGTTTGCCGTGTCTGTTGGAATGGTACATTGAGAATGTATTCCAAAAAAAGACCTAAAGTACTTGTTACCAGCTTTGCAAATGAAATCAACCGTATTTATAACTATGATATTACCTCGGATAAGAAGGAA
ATTCACCCCAAGAAAATGTGTTGGTCTTGTGCTGGTAAATTTCGACGAAGCAAATCTGTCCAAAAACTGAATCTACCCGAACTTGCAAAGTTTTCACCCCATTCAAGTGACTGCAAATTATGTTTCAAGCAGCCTGTTCAGCCAGAGTCATGTGGAGAGAGAAATGCAGAGGAAGGGTCCCCTCCTGAGATGGATACAGCCTCGGTTTGCACACCATGT
GACTCATCTACACCATGTGTTGTTGAAATGGACACTACCTCAGATGCTTCACCAAATCAGTGTGGAAAGAAAGATACAGTTGCAACAGAGGAAGGGTCCCCTCCTAAGGTACCTAAAGCAAGTGAGATGGATATGGCCTCGGCTTGCACACTAAGTGACTCATCATCTACTCCATGTGTTGGTGAAATGGACACTACCTCAGATGCTTCACCAAATCAG
TATGGAAAGAAACGTACAGTTGCAACAGAGGAAGGGTCCCCTCCTAAGGTACCTAAAACAAGTGAGATGGATATGGCCTCGGCTTGCACACCAAGTGACTCATCTACACCATGTGTTGTTGAAATGGACACTACCTCAGATGCTTCACCAAATCAATCATTTACACCATGTGATGCTGATACCGTCACACAAATTACAACTACAACTAGAGTGCTCTTC
CAAGAGTCACAAGATGTCCAAGTAATCCCACCATTAGATCCACATAGATGTGTAGTCCCTGATGTGGCTCTCGTACTCAAATGTACTTGTTGTCAACTGATTCCTTGTCATCCAACTGTGATGCACTGTGGAGATATTGCATGTGAAAAATGTGTCCGCAGAATGTTTTCCCTTGGCATATCAGCATGTCCATCATGTGGCATTGCATTTGAAGATGAT
GACTTTACGTCGATGGAGGGTAATGCACTCAAACTTCTTCAATCTCTACCCTTTAAATGCAAGTATGCTTCTAAAGGATGTGCCGAAAACAAAACCAGGGCAAATTTGTTAGAACATGAGAATAATTGCCCCTTCCAACATGCCCCACTCAGAGCGTTGCAACAGCAGCATCGTGGTGCTTATAAAAAAGAGCCGCTCTTTACTGTTCATGAGTCTACA
CGTAAAAGGCGTACACAGGAGGCATATGATATGCTAAAGCAATGGGCATCAAATAGAATAGAGGATCTTGGAGACACTCTTGCCTATCTCCTAACGCGAGAATGGACGTATAATGAAGATGATCGTCTAGGTCCTTTCCTCAAATTACTTGATGATGACATTTTGACAGAAAAACTGAGTGTAGATGAATGCCTTGCACTTAGAATTACATCCCTTCAA
AGTAAAGCTAAATATGAATCAAATTATATGTTTAACAAACAATATTTAAGCCAATCTCCGTATCAACCGCCACAACGCCTGGCATCAAGAGAATACACATACATGCCAGCCACAACACGTTATTTCATAAAGAGTCCCAACTCCTATTACTACCACTCACCAGTGAAACACAACTTGTCAGGCCAACTTCACAACACTGAAACAGGAAATACAGTTGAA
CCATTAAATGTTTTAAGTGATTTCACACTGTATGAAATGCCCACCCCTAATTTAAAAGGTGTGCGTTGGCGATACTGGGATGCTCTAGCTAAAACCCTGCAGGAATTAGAAAGTGAAATCAAAGCTGACCTTAGATTAGGATCCTGAGGAAGAAGAATCCACACTAATAGTTACAGTAAAGGATGGGGCAGACGGTTTAGGTGATGTTTCCGTTTATAG
GGACAGAAATAACCGTTCCCTACCCGACAAGGCCTTTCGATTTGCCTTTACTGTCATTGGTATTGACATAGAAATACGAGGTGAACGCAAACATGTATACATTAATGAACATCCTAATTCTGTGCAATGCGCAAGGCCTCTTCTCGAGGCTATTGCCGATGAGAATAATCATGGATCGTGCAGAGTGCTGATGCAACCTATCGAAGAAGAGCGTGAATT
CATAAAGGATAAGATTCTAAAAGTGGAATGTGATGGACAATGGCGCAATTTCTCGCTCAAATTTTTTACAAGCATGGTAGACGAGAAGTTCAGTCGAGCAGAATCAGGGTTAAAGGGAAGTGGCTCAAAATTCATGTGTACTCTATGTAAAGCTACACGGGAGACATGTAAGACCAATCTTGGGACTTTTGAAATTGACAGGACACATAGAGAGACTTG
CGAAAAGGCAACATTGATTGAAATAAATCCTGACAAACTAACTCAAAATGAACTGGAAAAAATTGCAATGGGTGTGAAATCAAAACCTCTCCTCCTATCTGAACCTGCTGATCAAGGCATAGATGCCACTCATGCAGATATAAATTTGGGTATTTTTTTTCGAAAATTACTTATTCGGGAAATAGCCCAAAAGCCTGAGTGGGAGGCTAAAGGAGATGT
AAAACACATTTTGGAAGATGCTGAGGTCAAATTTAATGCGCATCTAAAGCAACACATAGGTATAAATCCAACACTCATGATGCCTGGAAACTACGCACGGCAGCTGTTTGCTGAACAAAACGAAGACATTATATTGGAGTTGGTTGAAGATGTTAATAGAAAGGAGTTACTTAGAAAAGTGTTCTCACTCTTCCGTGATGTGCGCACTGTTTACCGTGC
ATCAACACCACTTGTTACTTATCCAGATCTCACATGTAGAGTCAAAAATGTAGCACTTCAAATGGGACACATACTGAAGAATGAATTTGCATATGCATCTTGGCCAAATTATCTTCACAAAATAATTGAACACACACAGGAATTAATTGAATGTGGACCTGGTACTATTGGTGCTCTCAGTGGGGAGGGTAATGAAGCTGCAAACAAATTGTTCCGGAA
ATTGAGACTAAATCTATCTCGAACGAACAACACCTACCTAGGGTTAAGAGACGTTCTATGGGTCCATTGGCTCTATACAAGTCCGCGGCTTCAAGCAATTGGCTCAATTGTCAGAAAAAAAGAGAAAATGCCCCAATTGCCTTGAAGTGGGCCATTACCAAAATACTTGTAGAAGGCCCAGGGTTACTCTTAATGATTAAGAAAAATCAAATGACTACAT
CTGTATTAAGTGTTCGAAGTTTGTTGTTAACCGTATGGCTACGAATAGAAAATTACAAAGGCAATCTTCAGTTTCTGAAATTGATAAGTAGTCAGTTAATTCCATGATACTAATTAGTATTTGGCTTTGTCGAAATGTTTTTGTTTGTATATGCAAAAAAGTGCTTTGCAGTACCCCCACATATCATGATGCAATTATCATCCAATTTGAGTATGCTAT
GACCAGCTGTTTTGAAGATTTGATCTGGGGCTTCAACAACATGGATCTGGGTTTCTGGAGCAGATACATCTATCAGAAACATACACTTTGATATCTCTGGGACTTTCACTTTGGTCATTGTTTGCTCATTTTGAACAGTTCCATATAAAGACAACAAGTCAACTTGCTTATATTGGAAGCCTCCATATACACATAGCAAGTTCTCTTTCAACTCTATTG
TGTCATGGTATGTTAGTTGAATTGGTATTGATTGGCATACACGCAATGACGATAAGTGAAACTGCCTGTTTGAAAGGTTGTCAATGTGTAAAATGGTAATGTCATCAAATGAGAGTATGCTGACTTGGTCTCTTGTGCCTCCAATAATGTAGATCTTACTATTATGAAGTGTCGCAGTGTGAAATGTTCTTGGTTGCACTTCAGGGATAATCAATATCT
CCCAATTATAATGCGAGTCAGATACATCTAGCAAATAAAATTTGCCATCTTCACAGCTTGTTTTGAAAGGTGTTGGTATACAGCTTGACTGATTCTTTCTGGATAATCCACCATGAAGAATGAACTGATTTTGCCCAATCCTGGTCAAAGTATGGCCAAATCTTGCTGCAAGTAGGGGAAGAAATACAAAAGTTAACTTTCAGATACATGATAGCATAT
ATTGCTGTAAGAGAGGGAGCCCGTTATATAGACCACTCCATAACGAGGCCATATATGGTGTCACATTATGCACTGTCACTAAATGATTACATATGTAATTAATCGTGGTACATAATGCCATTCTCTGGCAATTAGAATAGCCACACTATTCTCGGTGGTTGTGTTTGAGAGTCACCTTCCTTGTGTCAGTTGAGATCATAGCGCTGATTTGAATGAGTT
GAAAACTAGTACTCAATATTTACAGATCAAATGTATCAACCTGTTTTTTCCCTGCTTTTTCTTTATTTCTGTCTTTCTTTCTTTCCATTTTTAATGCATGGATTGTGTTGCACCATTTCTGACATTATGACAAATTTCTTCTTTCTGTCAATCATATAACTCTTGACCATCTTATTGGTTGTAAGAACCAATGACCTTACCTGAACATGTCATATGATC
AGTGTTCCTACTTAGACCCCAAAGGTCCAAGTTTGTAGGTCAAAGGTCTTTGTAAATGCTCCTAACGATATGAAATGTGATGATCTTATGTCCTTTAGCACATTCTGAGCAATACTAGTATTAAAACATTATAAAAATCATTCAGGTCACACCAAAAGGTGATCAAAGGTCTAATTTTGAATTAAAAAGAATTTAAAAGCCAATACCACCTCTGACAAG
TTCAAGGTAGCAGAATTATTGTACTTGCAAGAATATATTGACTAAAATTACTACTACTAAACACACAGATGACGTAAATAAAAGTTGATGGTGAGCCCATAGGCCGACCATTCCTTTGTTGAAGGGTCATGATAAAAATCACATTTAATTACCTTGTGGTACACTGCCCTGTTGTACCTTCAGTGAAATGGACCCATCCTCTGATGGCTGGATTGTGGT
ACAATCAAATGTTAGTCTGGCAGACCGTGATAGTGTCTCTCTCAGCACAACTAAGTTGGATGTTGGCTGTAAATATACCAAGTCCTGCCCTCCCCAAATAAGGAGGGAGACTTTGCCTTCCTCTTCATCAAGTGTCACCATGGTGGCGCTGTGTAGCGGCCTTAAAGTACTTCCTGCAAATAATTTGAACTATATGTGGGCCGCGGCGGCAAAACGATG
CAGTTGTCGGAGACCCACATATAATACATACAATATTCTTGGAGTGGAGTATGATATATGATTGTTAAGGTGCAAGACCTCACCGGAAGCCCTCCCAATGTGAATAGGATCAAACGAATACCATATGCTCCAGGACTTTTCAATGTGATTTGTCTTAATTGTATCCCGGGAATTGGATGAAAGAAGTAACAGACATGAACAGCTTTGCTTCGAGGCGTA
AAACTGTAGCACCACCCCCACCCATCCTATATCCAGTATATCTGTGCAGGGCTTTATGCGCTTATCCTGTGCTATTGAAATTCTTGATATGGGTATCATTTTAACCATTTTTACTCCTGCAGATATTTATAGTACACAGTACCCGCCTGAATTCAGTGTCTTAGTGATCAGTCCACTTATCCTTAACACTTAAGTGACCAAGGGGTGGGGGGAGAGGTT
CCTAGGGGCCCTACCACAGGTTTTCGCGCGTAGCTCGAAAAGGCGACATTTTTTGACATCGCCGCTCCATGTAATTGTCAAACTTAACGCAGCAGGTCCCTCGGTGGCAAAAAAATTCAAATCGACCCAGTTGTTTTCGTGCTAATTAGCATAAAGTGATTTTTCGAGGGCGCCGTTCAAAATTGTTTTGCCCATAAAATCAAAACCGAATAACTTTTT
TCTATTTTTATTTTTGCAATGTAATTGTACTTGCAATGTGCAAATGTGAGCAAAATTTCGTGTGAAAATTCCTTCGGGAACACAGTTTTTGAAATTTGTTGGGGAAATTTTTTGACGCAACACAAAAATAAAAATTGGGCCGCCATTTTGAAACGGTTCATCCGATTTTCAATTTTTTTTTCTGTATTTTGGTCCTTTTTACATGCTTTATTATTGACC
AGACTTTCAATGTAAAATTTATACTGGAAGGATGTTTTTTAAGGGGGGTCCCTAGGAACCCCCCCCTTGGGCACCAACGGGTGGGAAATACCTTGGTCACTTAAGTGTTAAAGCAGTGACCTAACCATGAAGGTCGTTTGATTATATTCAAATATAAAATAGTGTATGCAGGGAATGGGATGTAGACCCACCATGCCCACGGTCAAATTGAAATGAAAT
TGGCACTGTGCTGTCTTCCAGATTATTTTGATTTTTGATTTTACTTCCACATATTTGGGAATACTTCTATAATGAGGTTTTTGTTCCTGAGAATTAGGAATTTTAGTACTACATTGAAGCATTGAAGGCGGAAATGTACCTTTTGCAGAAAACTGTTTGGAGAGGAGAACCATTTTACACTGATCATGCTGCTTTCAGAGAAAACTTTTTGGAGAGGAA
AACAATTGTACACTAATCATGCTGATTACATGAATTGATTTGATAAGTATAAGATGTGTGTACTGGGGATTGTCTGTGATAGTTCTTATAAAATTCCATGGTCAAATTAGAGTAACTTTTGTAATCTTTGGAGGTGATTGGTCTGTATGAAGGAACTTGAATGAAAATAGTGGTTAATTTGAATTTGTTCTCCAAAACAATTTGAGACTTATGTGCCTG
AAAAATAACAAACATGTGCCCTTACATGCCCTTTTGTTCTATCACATTGTAAAGCTACATGTAACTAACTTTACAGCTCTTCAAGGCAGATAACATAAATGCCTAAAAGTCAAAATGAGTACTACATTATACATAGGTTTAAACATTGCTTAAGTTTTATTGGGGACAACAATAGTTTTTGCATGGTTTGGTCTGAACACTATCATTACCTTTTATGAG
GCACTCTTCAAAGCTTTTGAAGCTGATTTCATTTTGTGCACTCTCTACCTGTACTTTTACAACACGTGAGGCAAATGGCCATGTAGCTTCTTCTGTGTAGCGCCCACCTCCAAAAAATATGTACTCCTGGATCACTTCTGGAAATGTTAGCATGAACATATAATAATGCACTTATCACTTGTAACTCCGGCCCCCGGGTACCCGGGCACTAGCGGGGAA
CTCGTCATGCTTGCGCGGGGAAGTTGTGCATGCTGATGTCCCCGCCAGGCGGGGA
```

```
>OspRAG1L-B
MAESKLAPIQVHMERLMMVCRVCWNGTLRMYSKKRPKVLVTSFANEINRIYNYDITSDKKEIHPKKMCWSCAGKFRRSKSVQKLNLPELAKFSPHSSDCKLCFKQPVQPESCGERNAEEGSPPEMDTASVCTPCDSSTPCVVEMDTTSDASPNQCGKKDTVATEEGSPPKVPKASEMDMASACTLSDSSSTPCVGEMDTTSDASPNQYGKKRTVATEEG
SPPKVPKTSEMDMASACTPSDSSTPCVVEMDTTSDASPNQSFTPCDADTVTQITTTTRVLFQESQDVQVIPPLDPHRCVVPDVALVLKCTCCQLIPCHPTVMHCGDIACEKCVRRMFSLGISACPSCGIAFEDDDFTSMEGNALKLLQSLPFKCKYASKGCAENKTRANLLEHENNCPFQHAPLRALQQQHRGAYKKEPLFTVHESTRKRRTQEAYDML
KQWASNRIEDLGDTLAYLLTREWTYNEDDRLGPFLKLLDDDILTEKLSVDECLALRITSLQSKAKYESNYMFNKQYLSQSPYQPPQRLASREYTYMPATTRYFIKSPNSYYYHSPVKHNLSGQLHNTETGNTVEPLNVLSDFTLYEMPTPNLKGVRWRYWDALAKTLQELESEIKADLRLGS|EEEESTLIVTVKDGADGLGDVSVYRDRNNRSLPDKA
FRFAFTVIGIDIEIRGERKHVYINEHPNSVQCARPLLEAIADENNHGSCRVLMQPIEEEREFIKDKILKVECDGQWRNFSLKFFTSMVDEKFSRAESGLKGSGSKFMCTLCKATRETCKTNLGTFEIDRTHRETCEKATLIEINPDKLTQNELEKIAMGVKSKPLLLSEPADQGIDATHADINLGIFFRKLLIREIAQKPEWEAKGDVKHILEDAEVKF
NAHLKQHIGINPTLMMPGNYARQLFAEQNEDIILELVEDVNRKELLRKVFSLFRDVRTVYRASTPLVTYPDLTCRVKNVALQMGHILKNEFAYASWPNYLHKIIEHTQELIECGPGTIGALSGEGNEAANKLFRKLRLNLSRTNNTYLGLRDVLWVHWLYTSPRLQAIGSIVRKKRKCPNCLEVGHYQNTCRRPRVTLND
```

```
>OspRAG2L-B
MFMLTFPEVIQEYIFFGGGRYTEEATWPFASRVVKVQFKLFAGSTLRPLHSATMVTLDEEEGKVSLLIWGGQDLVYLQPTSNLVVLRETLSRSARLTFDCTTIQPSEDGSISLKVQQGSVPQARFGHTLTRIGQNQFILHGGLSRKNQSSCIPTPFKTSCEDGKFYLLDVSDSHYNWEILIIPEVQPRTFHTATLHNSKIYIIGGTRDQVSILSFDDIT
ILHIDNLSNRQFHLSSLRVCQSIPIQLTYHDTIELKENLLCVYGGFQYKQVDLLSLYGTVQNEQTMTKVKVPEISKCMFLIDVSAPETQIHVVEAPDQIFKTAGHSILKLDDNCIMICGGTAKHFFAYTNKNISTKPNTN
```

### *Ophioderma brevispina*

#### RAGL-A

##### **ObrRAGL-A.1**

| Genomic loci | RAG1L status | RAG2L status . |
|---|---|---|
| ObrRAGL-A.1 | potentially functional (1 CDS) | potentially functional (1CDS) |

```
>JAMKCH010037210.1:5000-15000 Ophioderma brevispina isolate Catalog1970-2016 scf_37218_, whole genome shotgun sequence
ATTAAAGTTCATAAGAAATTTGGTAGTAAACATTTGTTATTTAAGAGGTGATAATTTTTTAAAAACCCAAATAACAATTTTGGTATTGCAAAGGTTGCCAAATTTTGCACCTTTTCTGGCACTTATACTACAGTTTTAAATTTTGCCCAACCGACTTTTTCTCTATGCCTTTTTAGACTCAACGTAAGAAATTGAACAAATTTGGCATAATTTTGTTTT
GCAAGGTTTCTGAATTTTGCACCTTTCCTGGGCTTATTATATCCAATGACACATAACGAAGTGGGCTCGGTGCACCCATTGGATTACACGCTTTATTCCGTCAGGGTATAAAATGTATGATAAAAAAATAAAGCGTGTAAACCAATGGGTGCAGCGAGCCTGTTTTGTTTCGTTTCATCGAACAATATAAAAAGCCCTTTTCTGGAACTTTTTTTCATT
CAGGTTTCCAAATTTGTATACCATGATTTCAAATTTTGACCTGTCAAAAATTTTCTGCATGTTCTCGATCCCAGATAAGAAACTGAAAAATTTTGGCGCGGCAAATTTTTAACAAATTTGCTCAGAATTTGATTGTGTCATGACACAGAATCAGTTTCGGGGAAAAATTGATTCTGTTGCGGGTCTGCGACACAATCAAATTCTGAACAATTTGACGGT
TTCCTTATTTGGGATCCAGAACATGAATTAACATACATTGTATAACCTGAATTTTATGACAGTGATTTACTTTTTTATCCAATTTATACAGGTTCCGTGCTGTTAAGTCATGGATAATTTTCACCAAGAGAACCTTAAAAAATGTTGCCGTGTGTGTGGTACAGTGTATGAGAAAGTGGCCAAAGCAAAACGTGGCTATGACTGCAGTGGGATAGACAA
GAAGTCTGGCGTTCCATTTTCTGAACTCCTTCAACATGTGTTCAAGATATTTGTAGAACATGATGATCCTGATCTCCACCCAACAAAGTTCTGTCACCGGTGCTACCTTATTGTATCAAAATGGTGTGTTGCACAACAAAGGAAGCGTAAATACAACCCTGCTGCGAGATCATATGACTGGTTACCTCATGACGGCGTGTCTTGTACCTTTTGCAGAAG
ATTCCAAGGAAAAGGGAAAACATCCCAGGGGAAAATTCAATCCAAAAGAGTTAAGAATGTAGCTGGTGAGGAGACTCCGGAGGTCACTACTCCCTACAAAGGTGATCATACTTACGCAGATGTCCCAGCTCATGTGTTGTTTGTACGTGCAAGAAAAAGAAAGCGTGATGAAAAGTACGCTCCAAACAAGAAATTGATCTTATCAACAGATATTGATGA
CTTTCAGTCTGACTTCAACCTGTGCAGCCCGTTTAGTGCTGACCTGTGCCTAGCTGAACCCCTGCATCGTAGTTACCAGTGTGGAATTTGTGCCGGCATAGTAGATAAGCCACATAAAACCAAGTGCCTTCATGTATTTTGTGCTGATTGCATTAAGACATGGGTAGCTCGATTCACTTGTTGTGGATTGTGTGAAGAGATCCTCAGTTTGCGGGATAT
TAGCCCATGCGACAAAGTGTTTTGCCAGTTGATCCAAGATTTACCAATGAAATGCATAAATTATGGCTGTAAGGAAACTCTAACAGTTCAAACTTTTGAAGCCCACATCACTTTGTGCAAAGAACAGTTCACTGGCCTAATACATGTATCTCCTACAATTCAAGAGCCCACACAGAGCTGGATTGATGCAAGTTCTTCTGGCAGCAATTCATCTATGGA
CTGCCTATATACAGATGAACCAAGCACTACGGACTCCACTGATGATGAGTCGTCGGCAAGTTGTCTTGATGAATCGGTCTCGTCCACTTGCTCAACTGAAGATGAGACTTCGTCACTAGATGAGAGCTCCTGCTCAACTGCCTCATCATCTTCAAATAAGTCAAAACGTGGTAGGAGGCCGCAGCCCTTACTTGACTTGAAACAACCAGCTCAAAATAT
GAGGCTTAGAACTTTAAAGAAAGCCATACAAGAACATGCAGAGAAAAACAGTGAAGATATTAAAACTGTTTACTATGCTCTACTGTTAAATTACCTGAGATCTACTAAGCAGTGGCAGGAATTTTCAGAGGTGAAATGTGCACTTACTGGTATGAACTTCATGGGTAGGACAAGGTGTTTAGCAGTGAGAGTTTCAACGTTTATGTCGGTTACAAAATA
TCGACGTGTACAACAAGCAACAAACTATACATCAGGGAAACGAGTCTTCGAACCTTACAGTGCTTTGCTGAAAGAAGAGAAACTGTATCTGCCTGGGTCTAATGGATTAACATTTTCAGTTACTCCTCCCATTCATCATGTTCAGAAAAGGGAAGGCGAATCAATTGAAGCGTTTAGCAACATCCCCCAGAACTTTCCTGAGATTGGACTCCCACATGT
GAAAGCGGCACGCTATCATTATGATGAGGCCATAGCAACGTCCCTGGCTGATCTTGAGCTAGAAATTGAAGAAGGACTAAGATCTGTTGGGGAAGACCCAAACACATACTGGCAGCCAATAACTGTCCATATCAAAGATGGCTGTGACGGAATGGGGGAGGTGAAATGCAAGCAGTATAAAACAAGTAAAGCAAAACTTGCTGATAAAGCATTCAGATT
TAGTATGGCCATAATCAGAGTAACAATAGAGCAAAACAATTCAACAGTCACAATTTATGAGGAAAAGACTCCCAACTCTCAACTCAACTCAAGACCAGTGCTCATTGCCCTTGCTGATGAAAATGATTACCATGCAGTGACATCACTACTTGATCCTCTCTTGCAAGAAAGGAAACACCTGGAAAAGTCAGATTTACAGCTGCAAAGTCAAGCTGGACG
ATGCTGGAACTTCAAAACCAAATTGTATGGGACAATGTACGACGAAAAGTTGGAAAGAAAAAATTTCAGGTCTTGCTGGGTCAGCCTCCGAATTTTTATGTACAATGTGCGAAACAACAAGAAGTTCAGCATTTTCCCAGCCATTCAACTGCACAGAGGTAACACGAACAGCCCAACAAAATGCAGCCAATGTAGATAGACTCTACACAAATGATAGTAA
TTTGAGCTACAGCCAACTCCAGAAGGAATCAAAGGGTGTGAAAAGTGACAGTTTTGCCGATATTACACCTCACATTGATGCACTTCATTGTGAAATAAATAACGCACTATGGCTGAAGAAAGTATTTGTTCATGAAATTGCAGGACTTACCACGCACTGGTCCTACCGTGACAACACAAGCAAAAACAAGCTCAAGGATGCCGAAGAAAGGCTCAACGA
AGAGTTGCGTAAAGGTGTGGGTCTGCAGAAGCACTTAATGCAGCCGGGCAACTATTCCCGGCAACTAACAGGTCCTAAGGCTGTGAATATAATCGTTCAACTTATACCTGAAAATAGACGTGAAATTGTGCAGCAACTGGTGGATGAATTCAACTTAATGAAGGTTGTATGGAAAGCATCAAAACCAGGTGAAACCTGTCCAGACCTGCTGCGCCAGTA
CCCCGAGCATGCAAGATCTTTCATGCATATTTTGCAAACCCACTTTCGGTATACCAAAGATAATATGCCAAACTACTTGCACAAAGTAGTAGCCCATGTTCCTACACTCATTGAAATGTTTGGCTCTGTGGGTATATTTTCTAGTGAGCCGAATGAGCACGGGAACAAATTATTCCGTCTTTTTCGGAGGATGTGTGCACGTCAAAACAGTGACTGTGA
ATTAAAGGACATATTAAAATTTCACTGGTTGTATACATCTAGGCGGCTACAAAATCAGGTAAACAAAACACTCTGCCAGCAGGCGTGCTCTTTGTGTGGGGTGTTAGGGCATAAAAGTTCCACCTGCCAAACACATACAGGGGTGTTGTGAAAAAAAAAAAAAAAAAAAAAACATTGACCCCGGGCATTGACCTAACTGTTACCACGATGGAGTCAGAT
TTCACTGATCGTGCCAGTGGTAGAGGCAACGGGTGCCGGTCATAAAACACTTAACATGGTTGAGGAGTTGACTGATGAACTCATATGAAATTCGAAAATAAATGATGCGACGATGTACATGAAATAAAAAATTATAATTGATGGTGGTATTGGATCCAGTTTATTGGATACGATAAGGTGATTGAGTATGAATGAGATGAAGCATACTTTATCATTATT
GGCCTATACTGCGATCATATTATTATTATTTATTATTATTGTTATTATTATTATTAGTATACTGAAAAAACATACCATACAAACCAGAAATGCATTTTCAATCGTTCAGTTACATTGAAACGTTCTGGGAACGTTGCGGTTTCGTCTCGTCAAGTTGGTTGTTAGTTGGTTCTAATGAAACGTTCTGCGAACGTCGTAGTTTCGTCTTAATGAGATTCA
TATTTTTGGTGATTATCAAAGGTAGTAAATCCGTTCCACAAACGAAATTCGAATAACCAATTTGTAATACGAATGTACAGTCACGTTTTAGAAACGTGTTTACAACGTTTCTCAAAGACGGAACCAAAGTAGAACCTATTGGAAACGTTTCAGAAACGTTTCGTAACGTTTCGTGTTAGCTGGGGTGCTCTTTGTGTGGGGTGTTAGGGCATAAAAGTT
CCACCTGCCAAACACATACAGGGGTGTTGTGAAGAATGTGTTTTAATTGTAACATGTTGACTCATACATGTACATGTAGGCTAACCTTCACATATATTGTAGGAGTGGTTGAACAAATTGTTGGTACGTGTACGCACGCTCCGTGCACATGCCTTTGTGATAGAATTTCAGTGTAAAAAAGTTAACAAATATAACAAAGTTAAATCTGGTGGAGTGGTG
GCAGTGGAAGAGTATGTACAAACAAACTCAATGGGAATTTAATGTTCTGACACATACACTGTATTGTAATGCGTGTTAATTTAATACAGAATGTGCTTTATATATTAGCCCTTCCTGGTGAATGCTGACAATTTCAAATTTAATCGTAATTTACTTTGCAATTATAATCAGAATTACTGATTGGGTTAATAAAAGTCACAAACTCATCACACGTTCATG
TCTTATGGAAAATTAACAAAATTTTAAGGATTCATGTTTGTACGTGTTCATACATGTACCTACAGGGTATCAAAATTGCCAATTCTGTCTATTGCAGGCAAATTAATTGTCATGCATTTGAGCCATACATTGTACATGTACATGTATGTTGTACATGCAAAATGTATGTGTAGACGTGCATGTACATGTATTTACTGTTCATCAGTAATTCTGAATCAA
ATTTACGTGCACATAAAGTTGCTTTTACATGTAAACATGTATTTTCTAAAAACCAGTAATCAAGTTACTCATGTTGCACAATTTCTGGTAGTTGCTGGTGGGACGTCTAGAGATATTTTCATAATGCCCTGCATTGCCAGTGGGACGTGTTACTAGTATTGGGGTGAGAGTGACATGTTAGGTTAAACGGCTTAGTATATCCAATGACACGAAACAAAG
CGGGCTCAGTACACCCACTGGATTACACGCTTTATTGTCATACATTTTATACTACATGATGGAATAAAGCATGTAATCCAATGGGTGCACTGAGCTTGTTTTGTTTCGTCAAATACGTGTATAATAAGCCAGGTTAAACTGCCAAACCTCCAAATACATGTACCAATCAATATTTGTGCATGTTTGTAGCGCTAGTATATGTCTTCTTTCAACATACCC
TACAAGCAGGATTTGTTAGTGAATCATGACCATGGCGATTTAGGAGGCCCTTTCCCAGGAAATTTTTGTTTGGATGGAAATTGTGCCTTTTTGCGCACAAAATTTTTTTGAAAATTTGGGAGCTGGCTTCCACGGAAACATTTTTTTTTTCCCATGCCCTGGTCATGATATTCTGTCATACATGTGGAAAAAATGCTTACATATGTAGTCCGATTAATA
ATTTGTTGGACAGGAACAAACACAAGCGTTTGTGGCCTGCATGAGCCAGTGGAACATTGCACCTTGGTTTTTTTCAATGTGTAATTGGTCAAATAATATTTATTCAAATAACCTATGAAATGGTGTGCACAGAAATTTAGTGTAGCTTGTTGGGCTGACATGAGGTGTGTGTATCTCCTGTCCAAGGAATTTTTAATCCAACTGTACATGTACATTCAA
TTGAACAGGGGTAGAATTTCATAGCAAAGTGCGAGAATCAGTCTTGATTCTCAAAAATGAATTTTGCGGGAACCATCTAATTGGCTACTCTGTTGGAATCAACATGGATTCACGGAAATCAATTGAGAATCTCTGCGAGAATTGATTCTCGGGTTGTCGCCGAAATTCTAGCCCTGATTGAATGTGATTTTAAAACATTTTTACCAAAAAATGTGTTCT
CCATGAACTAATAGAGCTAAAAATATATAAATGATGCATCAAGTACATAGTATGTTGTTCAAAAATTTATTTTAATTCTTCAAACATCTACATCTATCAAATATCCATTTGAGTATGGAATAAACATTTCAGAGAAAATGAAAGTGAAATAAGCTACAGTTTAATTGGTGAATAAATGAAGGGAATAACTACTTGGTAAATGAAGTAATGCCCCTTTAA
TTGGCAAATTTAAAGAAATGGCTTTTGAAATGGTAAATGAAGAAATGCCCTTTAATTGACTTCATTAAATGAGTGTATATGGTGGTCCACGTGGACCCAAGATTTTGTTTTTCGCGTGCCATGTCTTAACCAGTAATTTTCACTCACATTTTTATCTTGGAGTTTTTCAATCTTTGGGCGAAATTCTCAAGAGTGAAATGTATAGAAGCCAAAATATGG
CGACCAAGGACAAGGTTGTTGATCGGATATGTTTACATTCTTGCTATGATTTATTATGAAAGTTTTTCCCACTCAATTGACCCGTATCCCATATTGATGGAATAGAAAGCATTTTCCCGAAGGCGACGAAAAAAAAAAAATGTTTTACGGGTGGGCTGACCTTTCCAGTATGGTCGGTCGGGCAATTTTTTAAACCTCTGACGGTTAAAATTGGCCTAAA
ATAGCCCCGAAATCTTGAGAAAAAGTTAAAAATTTCAACAAACATATTTTTTGAAAAATTAGGCAAGAAATGGTAAATTTTTGCCAAGAACGGCTGATTTTACATGATTTTGGCAAAACTAAAAAAAAGTAATCTTCCAAAATGCCAAATCTGGGTCGGTCGGGCCCGTAAAACATGTTTTTTTTTTGTCGCCTAATTACTCAATTTTAAAAAAAAAAA
GCTGCAGAATTTTGGACACCTCCATACAGTACACTGGCACCTGTATGTATTCAGTATGAGTTGTATTAATCGGCCGTAATTACACGTAGCTAGTAATCGGCTTATGTTTTTGTGCAACAAAACAAAATGAAACAGGCTCAGTGCACACATTGGTTAATTACATGATTCATGTACAAGGTGTACAGTACATGTGTGTACAAGACTGAATTGTTTAAGGTT
GTCGAATTTGGAAACCTGTATGCATACCTTTGTCCTCAAAACTAGTCCTACAAAATTGATAACATTCCAAAAGTTGATTATCTTCGATGTATTTCTGTTGCAGTTTTGGCTTATTAAGAATAAGTTTGAGAATTAAATTTGGTTTTGCTGAAAATTTTTAATTTTTTTTTTTAAATCAGGCCAAGGATATAACTGAAGCATCCATCTTTGGAGGCCCCT
TTCATAGAAGTACAGTAGAACATAAACAAATTATGAAAAATAAACAATTACAAAAATATGATTACATGAGAAAAAACAATTACTGACACATCTTGTGATAAAGCGCTGATTCAAGCTAAGCCCAATTTGTTAGCAGGATTTGATGAATTAAGTGAAACATGTACTTATGTATATATTATTACAAATCTTAACAGTGTTTGTATACTTGTACACGTATGT
TTTTGTTTTACAGCTGGACTGCATGTACACGTACAAATGTACAGTATGTGAAAATAAAGAATTATCTAAATTAAAAATATAGACACTTTGAATATCAGATTGTTTTTTTAAGTTGTTTGTAGCATGTGAACATGTGCTCGTAAGTTCCTTTTACCACGAGTTTGGTGGTGATGGCTCTTGTGATTTAGGTTGTATGATATAACATGTGCGCCGTCTTT
GCATAATACACGCTAATAAATGCTGTTGTAAATTATAAACAAGAATGCAGGGAGAGATAAATTGCGATGTCACTACCAAACTCGAGGTGAGAACTTAAGTCCGCATTTACACTAATTTTACAGCCACCATTCCCAGTTTAGCTTCAACTTTAATTCTTCCCATGGTTCATTCTATTAGGCCCCCAAAAGTGCCTCAGAACTTTAAAAATGAACATTTTA
ATATATTTTCTTAGTTTTTGTGAATATTCCAAAATGTAAAATTTTGAAGCAGTTTAAGAAGTGACCTAATTTTAAAAAAAAAATTGCGTGGAGATTTTACCTGAACTCTGAGACACTTTTTCTTATTCGGCCTTACATACACTCTAAGCATACATTATTCTGTATACACCGTTCGTGAAACCAGTCAGCACAAGAATTGCATTGCAGCCATTTCAGTTT
CTTTTGTTTGTTTAGAGTTAAATTGTTCAAGGTGCAGTCTTCAGCATTACAAAAAAGTGGTTCATCCTCGCTCTCACTTTCAGAATCTGATGACGAAGCTTCCAAGTCTTCACCTACTTGTACCAACTCTATACCACAGTCTAAATTGTCATCATCCATGTTATCATTATTCAGAGTTGATGCTAAATTGTCCATAGCACATGTAGTGGGTTGCATGTC
CTCCTCTACTACATGTAGTTCATCTTCAGGCTCGGTCATGTTGTCATGCATGTCCAATTCCAAATCTGGTTGGGCGACTGCTTGGAACTTCCAAACAGACTTGTCCTCTGTATTAAATAGTAAGGTTTCTCGAATTCCAGTTCTGATTGCATAAACTGACTGGAACATTACTGGTATGGTCTCACTTTCGATGACGACATCTCCATTTCGAAGGGACGG
TATATGTATACGGCGAATTTTATCTGATGGTACAACATTGGCATTTGCAAAAGGTGCCTCAAATCCACCAAAGAGATAAAGATGGTCGCCCAGTTCAAGAGTAGTGTGTCCTGAGATGTGAAGTAATTCTGACTTCAAGTCAGAGATAGCAATTCCTTTATGATGAAGTTTAATATCAAATAACTTTCCTGGTCTAACCATACGATTTCCATCAAGCTG
CAATCCACCAATAATAATAGCAGTGTCCTGGCCTATATACGAACATGTATGGTATGCCAGTGGTGGTAGAGAAGTAGGTACAGTCGACCAAGAATTGTTAATAAATCTGAAGATATTTGTGTTGACATGTCGAAATGTGCAATCCTCGTTTGTGAGTCCTCCGTATACAAACGCAAAATTGCGGCAAGATGTGATACTATGTCCAAATCGTGGCGATGG
CGAGCAGCCTACGTGATTCACAGTGTCCCATGAAAAATAGTTTCCTGTGAACTTTCCTGTATAAAAGTCGTCAGTCGAAGGTTGCCCAGGTATTTTCCAGTTCCTTCCACCGAATACATAGACTTGCCCTTCTCTACAAAATGAAGCCTGGCCGACACTCTCTCTCGGAATTTTAGTCTTTAGTCGTGACGATGTTGCACTCACGAAACGTCTACCGGA
CGCATTGGCTTTGTACACTAAAGGAAAGACATCTTTGTAGTTAGAACCACCAAAAAATAAAACCTGATCTGCACCTCTAATCCACGATATTCCATGCGGTGATGGGAAGTGTTCGTCTTGACCGGCCGAATATATTTTTTTCAGAACAGAATTCCCAAAAATGGTCGCCATGTGTATCGATGTACGTGAATATTGATTGAAAACAGCTGATGTTGTTTA
CGTATGACAACCGGCGCGACCCGATCTGACCTGCCCTGGCGATCACTTAGCCAAGAACGCCATTCCTATATGAACTAAATAAAACTATTTTAAAATAACAAACCATTATGTTTACCATAAAATAAATATTTTACTACGTATGCAA
```

```
>ObrRAG1L-A.1
MDNFHQENLKKCCRVCGTVYEKVAKAKRGYDCSGIDKKSGVPFSELLQHVFKIFVEHDDPDLHPTKFCHRCYLIVSKWCVAQQRKRKYNPAARSYDWLPHDGVSCTFCRRFQGKGKTSQGKIQSKRVKNVAGEETPEVTTPYKGDHTYADVPAHVLFVRARKRKRDEKYAPNKKLILSTDIDDFQSDFNLCSPFSADLCLAEPLHRSYQCGICAGIVDK
PHKTKCLHVFCADCIKTWVARFTCCGLCEEILSLRDISPCDKVFCQLIQDLPMKCINYGCKETLTVQTFEAHITLCKEQFTGLIHVSPTIQEPTQSWIDASSSGSNSSMDCLYTDEPSTTDSTDDESSASCLDESVSSTCSTEDETSSLDESSCSTASSSSNKSKRGRRPQPLLDLKQPAQNMRLRTLKKAIQEHAEKNSEDIKTVYYALLLNYLRSTK
QWQEFSEVKCALTGMNFMGRTRCLAVRVSTFMSVTKYRRVQQATNYTSGKRVFEPYSALLKEEKLYLPGSNGLTFSVTPPIHHVQKREGESIEAFSNIPQNFPEIGLPHVKAARYHYDEAIATSLADLELEIEEGLRSVGEDPNTYWQPITVHIKDGCDGMGEVKCKQYKTSKAKLADKAFRFSMAIIRVTIEQNNSTVTIYEEKTPNSQLNSRPVLIA
LADENDYHAVTSLLDPLLQERKHLEKSDLQLQSQAGRCWNFKTKLYGTMYDEKLERKISGLAGSASEFLCTMCETTRSSAFSQPFNCTEVTRTAQQNAANVDRLYTNDSNLSYSQLQKESKGVKSDSFADITPHIDALHCEINNALWLKKVFVHEIAGLTTHWSYRDNTSKNKLKDAEERLNEELRKGVGLQKHLMQPGNYSRQLTGPKAVNIIVQLIP
ENRREIVQQLVDEFNLMKVVWKASKPGETCPDLLRQYPEHARSFMHILQTHFRYTKDNMPNYLHKVVAHVPTLIEMFGSVGIFSSEPNEHGNKLFRLFRRMCARQNSDCELKDILKFHWLYTSRRLQNQVNKTLCQQACSLCGVLGHKSSTCQTHTGVL
>ObrRAG2L-A.1
MATIFGNSVLKKIYSAGQDEHFPSPHGISWIRGADQVLFFGGSNYKDVFPLVYKANASGRRFVSATSSRLKTKIPRESVGQASFCREGQVYVFGGRNWKIPGQPSTDDFYTGKFTGNYFSWDTVNHVGCSPSPRFGHSITSCRNFAFVYGGLTNEDCTFRHVNTNIFRFINNSWSTVPTSLPPLAYHTCSYIGQDTAIIGGLQLDGNRMVRPGKLFDI
KLHHKGIAISDLKSELLHISGHTTLELGDHLYLFGGFEAPFANANVVPSDKIRRIHIPSLRNGDVVIESETIPVMFQSVYAIRTGIRETLLFNTEDKSVWKFQAVAQPDLELDMHDNMTEPEDELHVVEEDMQPTTCAMDNLASTLNNDNMDDDNLDCGIELVQVGEDLEASSSDSESESEDEPLFCNAEDCTLNNLTLNKQKKLKWLQCNSCADWFHE
RCIQNNVCLECM
```

### ObrRAGL-A.2

| Genomic loci | RAG1L status | RAG2L status . |
|---|---|---|
| ObrRAGL-A.2 | complete, 1 CDS, 1 frameshift | complete, 1 CDS, 1 frameshift |

>JAMKCH010010543.1:14000-27800 Ophioderma brevispina isolate Catalog1970-2016 scf_10551_, whole genome shotgun sequence
AGAAAATACAAAATTGCGAATTTCTTCTAGAGCAGATCAAATTTGCACTATTGGGCCAACAGTTCAGATAATATGAAAATCTCAATTTCTTTGAGGTAAATTAGGGGATGAGGCTGTGGGTCGTGAATAATAACCCTTATGAGTGAATGCGTCAGTATGTGACGCGGCCGACGATATGTACATGTAGTATCAGGGCTAGAATTTCGGCGAGAATCGATT
CTCGCAAAGATTCTCGTACTTTGATTTGCGGGAATCAATGTTGATTCTCGTAAACTTTAGAATTTAACACAGTAGCGAATTAGATGATTCCCGCAAAATTCATCTTTGAGAATCAAGACCGATTCTCGCACTCTGCTACAAAATTCTACCCCTGAGTATCCAGTTCATTTTTGATACTTCTAGGTTTGTACTACAACTGCTCAAAATGCCTACAATTTT
ATAAGGCTCCTTCATATTAAACGAGACACTGAAACTTCGCTATATATTCATGTTCATATTCAGTGCAAGGTTGCAGAATTCGGCAGTCTTGCAGCACTTGCAGCATGTCTCAATTCCTTGAAACCCATGGAAATTTGGTTTTGTTTATAGGGATTTTTTAAAGGAAATGTGTTGCTACATTACTTTTTATTGTATTGCAGTGTTAATAAGTGCAAATTA
CTAGAAAATTACCTTGTTAATTTCACAAATTTCCAATGTATAGTTCAGTTGATTAATAGTGCACTACAATTAAAAATAATGTTGGAGCACATTTTCTTTAATTAAAGTTATCAAACTAGCATTTCCTGTGGTTTTAAGAAATTGATGTGCTGGAAGACTGCCAAATTCCGCAACCTTGCACTGAATATTTCTGATTTATTTTCATAAATTATAGCGGAC
AGTTAGTTGGTATAAGAGGAATCATATCCCTACCTGTTGTATTTATAATATGTGTAAAACATTTTCACATTTTCTTCTTGTTTTTGCTTTCAGGCAACTTTTGCACTATGTCTTCCTTGGATAAACACATTAATCATCTACAAAAGTGTTGCAGAGTATGTGGTGTGTTATTCTCAAAGATTCCACGTGGTGGCAGTGGTTATGAGTGTACAGGAAGAG
ACGAGAAGTCAGGGGTGCTGTGGAGTGACTTACTTCGCGCGTGCTTCAAGTTCGACCCCCTTTATGATCATCCACAAGTTCATCCGAGTAAGTTCTGCCTTCGATGTAGGCAAGTTGTATCAAGGTGGTGTGCAGCTCAAGTAGAAAATGCAGACTTCAAGCATTCCACTGAAGTGGTAGTATGGAGGCCACATGATGATGTCTTTGAGTGTGCTTTT
GCTATCGGTTTTCTGGGAAGGCACCGCCAGTCAGACGAGCCAGAAGACATAAGAAGTGTGGTAGGCGAAAAGCACGCACTGAGGAATGTGAAGCGGTGATACATGTACAAACATCAGCCAGCTTAAATGATCACTGCTATGCCACTATTTCTCCACCGGCTCAGTTTGTATTGAAAAGGAAACGTGATAAGGCTGATAGGTTATCAACAAAACAGCGCC
TTCTAGATCTTGACATGTTAGATCACAAAAGTCAATTCAAAGACTCAACACTTTGCAGCATGTATGATGCAGATCTGTGTGTAGCAACACCAATCCATGACACTTATATATGCAAAGTTTGTTATGGCATATGTGATGAACCGTACGTAACACCGTGTGGTCATGTTTGGGTGTAAAGCATGCTTGGAAAGTGGACAGCAACTTTCACTGCCTGCCCAG
ATTGCCAAGAGGCAGTCAGCTTGCGCAACCTCAGGAAATGCGACCCACTCTTCACAAAACTGTTGCTTGATTTACCAATGAAATGTCCCAACAACTGTAAAAAAATGTTGTTAATAAGGTCCTTTGAAGCCCATTTAAGAGTTTGTGAAACTGGTAGTGAAAGCAGTACAGAATTAACAGATGACAGCTCTTGTTCAGATAGTGTGACTGGGGAGAGTA
CCGGTAGTGAAGATGCTGAAAGCACCGAGACAATCAGCTTGGCATCTTCAGCGTCCTTGGACAGTACCAACGGTAGCAGCACTGATGAGGACACTACTGTTGGTATACCATTGGCCACATCTACACCTAGAACCAGTACACGTACATGTAGACGGGGTAGGCCACCACAGCCATTAGTGACCCTAAATCAGTCTGCACAGAATGCGAGACTGAAAACCA
TCAAGAAAGCTATAGCACATCATGCTGCTCATTACGAAGAAAACCTGCACACAGTGTACTACACGTTACTTATGAATCATTTAAGAGCATCCAGGGAATATTCAAAAGCTGATGCAATTAGTGACATTTCACCAATAAGACAGAGCTCGATGCAAGAACTTGTTTGGCAATACGCATCACCACATTCCAATCGGTACAGAAGTACCGCAAAATGCAGC
AAGTTACCAAAGATGTTGGTAAGAATATCTTTCAGTCGTATTTTGCCATGCGGGAAGCAGAGAGGGACTTCCTACCTGGATCAACGGGTATAAGCTACTCTGTCAAGCCACCGATGGAGCATGAATGTAGCAATGGAGTGTCCATAGATGCAATGGGGCACTTGCCATCAAATTTTGCTGACCTGGAACTTCACACATCAAAGCTGAAAGGTACAGAT
ATGACGAGGCAATTGCAACTGCTATCTGGGACTTGCAAGCAAAGATATCCAGTGGCTTGAAACAGGCTGGAAAAGATCCCAGTGTTGGCACACAATATGTCCATCCTTATTAAAGATGGAGGTGATGGTATGGGAGAAGTTAAGCGGAAAAACTATAAATCTGAACAGCAACTCGCTGATAAAGCCTTGCGCTATTCTTTTTGTGTAATGGAAGTCA
AGAGTGGTGACGAAACATCTACCGTGAAGAAAAACCTAACTCTCAACTCAACTGTCGTCCCCTACTGATCGCATTAGCAGATGAAAATGACTACACTAGCAGTTACATCACAACAATGATGGCACCTTTCTTCAGGAACGGAACCACTTGTCAGATGGGGAGCTTGTAATTACACATAGCGGAACGTGGAATTTCAAATGTCAATTTTTTGCAACTATGT
ATGACAGAAAAATTGGAAAGGAAGGCCACTGGGTTGGCAGGCAGCTCATCTGAATATTTATGTACACTGTGCGAATCTACACGGCAAGAGGCCTATAGCGACCCATTCAAATACCAAATTACAAGGAGTGATCAAAGGAACCAGGAACTAATTGATGAAAGAACCAACAATGACAACAAGGAAACATATCGGACACTAACACATAACTAAAGGGAGTGA
AGGCAGATTCCTTCATTCGAAACACGCCGCATATAGATACCCTACATTGTGAGATTAACTCCGCAATATGGTTGAAAAAAATATATGTGCGAGAAATTGCAGGAGTGAAGCAATGGGCAAAGAAGGCCATGAAAAGGAGTTAAGTCATGCTGAGGAGATGCTTGACAGTAAGTTAAGAAAGGATGTGGATTGTTGAGGCAGCTGATGCAGCCTGGGA
ATTATTCTCGGAGACTGCTCTCTAATAAATGCACAGCTGTTGTGATGTCTTTAATACAGGAAGAGCGTAGACCAGTTGTTCAGCAACTTATAGAGCAGTATCGCCAACTGAAGAGAGTTTGGAAATCTACACAGCCAATAGAATCATGTCCATTGGATGTCATTGCATACAACCAGAATGCAGCCGTGTTTATGGACACTATCAGAACACATTTTGATT
ATGTACAAGCCAACTTGCCAAACTATGTACATAAATTTGTAGCTCATGTTCCACAGCTGATAAATATGTATGGTTCTGTTGGAGCATTTTCCAGCGAAGCCAATGAACATGGAAACAAGCT[redacted]TTCCGCTTGTTAAGAAGAATGTGTTCACGGCAGAACACCAAATTTGAGTTGGCAGATATTTTGAAATATCACTGGCTTTATACAATGAAGACTTTGC
AAGACTTATCAAGTGCCAACAGAGTCCTTCAGAAATGTTCAAAGTGCAATGAGGAGGGTCACAAGCGCAGTACATGTTTTAATGACTCAGAGTAGATAAAGCTGGATTTCAGTTCAATTTAAAGGTGCAATGAGAGATTTCACAATCAGGTGTCCATAAAAAATTGAAAGTACCGATTTTGCCCTGATGTAAGGCTTTCATTAAAAATATCCCCATA
TGAAAGGATTAAGCTGAGAACTGTTTCTTTGGTACAAAATTGTGAAATTTCTAATTTCGTATAGGTATAGGAGCAAATACAAAATTCAGCCTCACCTTTTTTTTAAGTCTCTTTTAAGATGTTCTGTAGCCTAATACAAGGTAGAAATCTATTGAGTGTCTAATCTATATGAAATTACATATTATGACACAATATTTGGCTGCCTCAAAATCTCCA
TTGCGCCTTTAACTAAATTTAAATTGGTTTCATCACATACAGTGTAATTACATGTATCCAGTAGAACCTTATTCTTGTTGCCAAAGAAAGTGAAAATGTGCAATATTGAAAAAAGTATGCCAATATGTTTGTATTTGCCAAAGTCCATTTCAACTCCTTACAGTGTGTATGTACAAATCAGCACTTGTGCAAGCTCTGTTCAATAGTATGGTATATAGT
ATTATGCAACTCCTTTTGTCGTTAGCAATATTGGGAGCACGCATTCAACTTTAATGTAACAAAAACCAAACTCCCTAAATACCACTCTTTTCAGGGGCCATTTTATATCAACTCTTGACTGTAAAAAAATTGAAATTGTGTTTTCCTCATTCTTCACCAATGAGGTTTGTGGAGAAATCATATGGTACGTGTTTATTATGCAACTCTTTTTTGTCATTA
GTTATTTGTGAACACGCATTCAACTTAACATGACAAAAACAAAATCCCCAAAATAACTCCTTTAATTTTCAAGGCGCCATTTCTAAATTATCTATTCTGGCTATAACATTTTTTTGAATATATGTGTGTCCTCCACATTCTTTACCCATGAGGTTCACAGAACAAATCTAAAACATAGGCAAATGTTTTATTTTTGGGTTAAAATTTGAACAAACAT
TCTTCCCTGTAAGAAAGAATTTTGAGAATGTGATGACATTTCTATTGAATTTTGCACCCGTGAAGCTCAAAACTTGTGAATTTTGGTGTTATAGACCTTGTAATTGTGTGTGCGTGTGTTTGTATTAGACCATTTTTTAATCTGGTCTGTAAGAAAAATTAAATGTGGATGTACTCGGTGGGCACAATCAGAAGAATGTAATACAGTGTATATGCAA
TACATGTACACCATTTACAATGGTTGTGAATTTAGTTGTAAATGCAAATTATTTAAACATTATTGGTTACATTGGTTTAATATTGTAATAACATATATTAAGCCATATCATTATGTTAGTCTTATATTATTACAATATTAAACCAAAATATAACCAATAACATTATAAACTGCATACAATTCACAACCAAAATACAATGTATGATGTTGAATACTGATC
TTGCTCATCAGTGGGTAAGGCCATATAGCTGTCTAAATTCTACCCATTACATTCAACAATCAATTGGCAAATTTTAGTAAAAAATGCAATAGTTTTTTACTTGTTTTGGACCATTTTAGCACTTCTATTCATGGTCTTTAAGGAAATGTGATAAAGGAAATGAAGTAAAAATGTGCACATCTACATGTATCTATAAGGAACAAGATTGAAAATTAATT
ACATGCCTGTTGAATTCTGCACCCATGACATCCAGGATATGTCCCAAAAATTTGGCAGTTTTATTGCTTTGGACCATTTTTAATTTTGAGCACATCTAATCCAGTCTGTAAGGAGATGTGATGTGCATGGTGAATTTTGCACCCATGACGTCCAGGATAAAAGTCATGTCCGTTTGAATTCTGCACCAATTATATATACATGTAGATCATTTTAGTA
TTATAGAGCACTATTGTTTTATATGTTTGGATTATTTTAAGCACATCTATTCTGGTGTGTGAAGAAAAAGATAGGAAATGTGATGGCATGTTTGTTGAATTCTACACCCATGATGTCCAGAATCATGTACCAAAAATTTGCAAAGTTTTAGGGGTGTTTCCGTAAGGTGACCCACTTCGTTCTGACACACGAATTTGGAAACCTCAACGGATAAACC
ACACACTTTGTGATTTATTAGGTTCCAAATTCAAAGTGTGCCAGAAAGAAGTGGGTCACCTTACAGAAGCATCAAGTTTATATAGTTGCTGTCACAAAAGAACGGCCCTCTCTCGACTGCGTATACGCTCGCGATCAGATAATGAGCGATATGAGCGGGCTCATTTACGCAACATTCTCGGAATTCGAACGCGCAGTAAGCCAATCTGATCAACGCGC
TGTTGTTATGGCTATGACCTGATTCAAAAAGCCCATCGGACTCGCACTTACGTGTTTACGCACGATATGCTCCATTTTGAGAAGTGAGGGCCGTTCTTTTGTGACAGCTACTACATGTATAGTGTAATATGGAACATTGATTATTTGTTTGGGACCAATTTTTAGCACACCTATATTCAGGTCTGCAAGATGGGATGTCATATCTGTTGAATTCTACAC
CCATGACATTCCTTACCCCTGTCATGTCATATGTCTGTTGAATTATGCACCAATGAAGTTTACATATGTAGAAAGCAAATCAGAATTGTGCACATTTTAGTATCATACGAAGCACTAGCTGTTTTATATCTGCTGGACACTTTACAGCACAGTCACTGCCATTATGTCTCGGGCAAGATCGGGGATTGTGCAACGTGATGTGCATCAACCCGGGAGCAC
AAATAAGTTTTTGGCAACTGACGCGATTTGTAATTGGTCAAAATGTGCACGCCTATACTACGATTTAATTTGCATAATCTATTGTGTGCGTACAGCTTGCCTGAGAAATTCTGGTAGTGACTGTACATGAAGGAAAAACATTGGAATTATGATGCCATGTCTGTTTAATTCTGCACCCATGACATTGAACCAAAACATTGCAGTATCATATGTAATGAT
GTTGTGTGTGTGTGGCCCATTTTTAACACATCTGGTATGGTCTTTAAGAAAAAGAATGTAATATTTGTGAAATTCTGCACCCATGGCAACCAGGATCATGTACCAAATATTTTGCAGAGTTCTAGTGCTACATGTATTTGGAACAATGCACTGTTTGTTTTGTTTTGGACTATTTCTCACCCATGACATCCTGCCCCCCCCCCCAAAAAAAAGAGAATAG
TACCATGTGAAAGTTTGTAAAACATACTGATGATAATTATGTGTTTTGGCCCATTTGTAACAAACTTCTGGTCTGTGAGAAAAAAATGTCATGTCTGTTGAATTATGCACCATTGAAGTTCTGGGACACAGTACAAAATCCCTCTGCCATGGAAAGTGCTACACCGCTTTAGTTCCTACATGTATAAAACCAGAATTTGAGCATTATTTGTGTAGTT
TTATGGCAGATATTTGTCCTAGAGATTTGCTTTTGTTATCATTCATGTGTATTCTTTAAATTTCATATTTAAAAAATGGACGAGAAAGCGTGTTGGGGTATGATACAGCGAAATGTGAACTGCCACAAAACCCGATTCTGTGGAACTCAAATTTGAAAATGCACCCTGTCTTTTCTAACTCCTAAGTTGCAAGACACCGACAATGCCAATAGGTTCA
AAAGAAGATATTTTAAAAACCATGAATTTAAAATTCTCTCTTGCAAATTCATTGGCAGGCAGTGAGGCAGGGGCAAAGATATGACGGATACAAGAGGGTGCATAAAGCACATGTTTCAATAGGCAGAACTTTTTGTACTGTGCCCCCTTCAGAAACCAAATAACTGCACATTTTAGTATTATATAGAGCTGTGTTTTTATGTATTTTAGGCACTTTTTCG
CACTACAATGTATGTATTCTGGTCTGATACATGTAAATGTAAGGGAAAACTTTGGAAATAAATGTATGATGCCATGTCTGTTGAATTCTACACCCATGACATCCAGTATTAAGTAAAAAAAGTCAATTTTTAGTGCTATTTAGACATACATGTAGTAGCATAGTTGTTTTTGTTTATTGTTTTACGTACATGCACTACAATCCCTGCCAGAATTATTCA
GGCAAGGTTGGGGTATGCGCGCGTGACTTGCGTCAACCTGGAAGCACAAATAAGGTCTTGACAACCGCCATGAGATGGGTATTGCTGAGTGACCGAAAAGCTTATGCTTGTATCAAGATTTAATTTGCATGATTCATTGTGCACGTGCGGCTTGCCTGAGAAATTATAACAGCGACTGTAGTACTTAAAATGGAAGAAAGACTGTCAAGTTGAATATAATT
ATTCAGGACAAATTTTAAGTTGAGTGTGGCAATTTCAGCTTTTATAACACCAATTATTTAAGTACAACATTCCCCAATACAACAAATACAGTATGTGCTTGTAACTTAATAATAGCTTTTATGATCAGCACAATCAAAAGAGAAATACATTATTAACAAAACACAATGTCATGGTGGATACTACATCTAGATCACATGGCTAAAATATTTGGTTTAC
TGTACTTTGCAAAACGAAATGCCCATTGGATTACACGCTTTATTCCTTTGTGGACGGAATAAAGCGTGTAAAGCAATGGGCGTTTCGTTTCACAAAATACAGTAAGCCAAAATATTCTGGCTTATTCCTATAGCCCCATTATTTAATGAAGGGATAAAGTTTGTACTACATGTACGTGGTCGTACAGCTGATGAGCCAATTTAACTGGTACATTTGT
ACAATTTCTAATTTTGTACATGGATGTATTGATGCATATTTGCCATTTTCTTAACTTTGATTTTAACTCACAAAACAACATCAGTCAGCCTGGCCATCAAGATTGAGAATTTTATCACCAAATCCTGAATTTCACACTTGGTCTTTGTTGTATTTAACTTATAAAAGGTAGATATATTTTCACAAGTATATTTTTTATTTATTTATTTTACCTTGTAT
TTAGCAATACAAGTTCTTGTACAGTGTAGTGCACTTTTTTAGAAATAAAGCATTAAAAAACAAGTACTTAACAATGCCTTTTCTGAACTTTATTGACAGTGCATGTATGTACATGTACATGTATGTGTGTATGTGTAGAATTGCAAATTACATTACAATAGTGCTTTACTGAGCAGAAGTTCGATCAAATAAGCAATCAGTCAGTTTTCGAAGACCA
ATGCAATACATTGTACAAGCCCCAACGCTCCTCTCATAATTACATATAATTGTCATTGCCGTGCCTAACCACCAAATGCAGTGCCACAGGGGGGATATCTGGGTTTTTGTGGGGTGTGGCTATTTTGTGGCCCACTGACACACATGACCCTGATGAATTCAGTAGACTTTACGGAATAAAGTGGACTAATTCCGTAGAATTCGTGTAGAATTCGTAGGG
TCGTGTGTGTCAGTGGGCCACAAAGTGACCACACCCGTTTAAAGGCGTTTTTATGCTAACATACGAGGAATTGTACTACAGGAAGCCAAAATACCATGGTGGAAGCAGTGATATTTTGTGTGTATGTACTTGGAGTGTTGGGCTGGATAAGCAAATAAGCAAATCTTCGAGTTGAAAAGTTCTCTATTTTATACAATAATTGTCAGATTTTTTTTC
TCAATAGCACAAAGCTAAATTTAAAGGGGCATTTGTGATTCTTGTACATGTAGCCTCATCCTCCTGACTTGACATGACAAATGCAATTTTACTAGTTACTACAGTCAGTGAACCTGCCATTTACATGCATGCATAGACACGTATGTGTAGTGTTCCTTCAGACTGATGTACATGTCTGTAAACAGAGTCAAGGTGGCCTACAGCTACATGTACAGGGCA
TCATGACACAAACACTTGCTCATTACATAATTACCATCACCAATGCACCATAATTTTTTTTTTACTTGGTAAAACAAATAAAATATATACAGGTACTAATTAAACTCACGGCTCGTAACATCAAAACGACGTTCAAAATCTACATGTACTTCATGTTGGGAAATGAGCTTTTCTCAATATTTTATTTCATCTGGGGTGATTCACCAAGACTTGTGAT
GCATCGTAAACCCACACGTCAATGCATAGCAATTCAAACAATAGGGATTTATAACAGCAACTTGTGGTGCAGGTGAGCCTGCTAATTACGATGGGTGTATTGCTGTGTTGAAATCTTTGTGAAATCCACTGTACCAGGCTAAATGTATTTTGTGCAGTGAAACAACACCATCTTGGTGCAGCCATTGGTTATACGCTTCATTCCGTCATGTACAAGG
TGCACGATGGAACAAAGCATGTTTTGTTTCGTTGCACAAAATACAGGCAACCCATGCCATGCGCCCATGTACCATAAGTAAATGGATGTCGTTGAGATTGCAAAATGCACCAAATCAGAACACGTACGGAAAGAACTTGCAGGCATGCTGTGTGTGTAAAAAAATGCATAGTACAGTACAATGCTGAGCATTGTTTCTATACACCGAGCATGAGTGCA
ATTAGTTTTTCTATGAAGTGATCAAGCTAAGTTCAGATTGACAAGAGCTAAACGAGCATCACTTTTACGGCTGATTTTGATATTTAATATTCACAGTCAATAATATGGTAAATATGAACAGGGTTCATTCAGGAATTTGGGTATGAGCATCGTTTATGCCAGAGGGGTTCACACCCAAAATCCTATTTATAAACAAACCCCTGGCATGCATACCTGTAC
ATGTGAAATGATTAAGCTGAGAATCATTTACAAAAGTTAAATTTCTTAATGTAGGGGCAAATACTAAATTCAGACTGTTTTATATTGAAATTAAGAAAATCAAAACTAGTATTTTTTTTCACTTTCAAGGCTTATTATATTCGACGAAACGAAACAGGCTCAGTGCAGCCCATTGGATTACACGCTTTATTCCGTCAGTGTAAAATGTACGATATACAGAA
TAAAGCGTGTAATCCAATGGGGACACTGAGCCTGTTTCGTTTCGTCAAATATAATAAGCCACTTTCAAGGTATTTCTGTAGCCAAATACAAGGTAAGAAATCTGTTTAGGGCTTTAATTATGATTATCTATATGAAATCACATAAAACATAATTTATACATGTATGGGTGCCCCAAAATCTCTCGCTGCACCTTTAAGGCCCATGAAAATCAGTTCCGA
GTTTATCGTTACTTATTCTTGCAAAATGGAAATGCGCCGAAGAAATTTTCACAAAAGGCAGGGCCGCAGGAAAGCCTGGTTTTGCTTCAAAGTTATCAAAATGAGTTTTCAATTGCTAACCTGGTGAATACGACAATATTTTGAAAATAATTATTTTGGCAATCTTGAAATTTTGACGCGGGTAGTTCTAGTCTCAAAAAGCGATGCGGGTGGGTAACG
ATAAACTCAGAATGGACTTTCATGTGCCTAAACTGTATCAATTACACGTAACAGGAGATACATTTCCCTTTGCTCCAATGCAAGAGGCATGGTTCCACATAAAACATGACACACATTGAACCCACCTTTAATTTGAAGTAATGTCTTTGGAAGAACCAACATATTTACATTTATCATAGTTACAAAACAGTTCATCGTCGTCATCATCGTCATCATCA
CTGGATTCTAGTTCATCGTGGCACTGCTGTCCCACATCCATCTCTAGATCACTGCTCATGCTCTCAGTCTCATAGTCATACTCATCGTCGTCAGAAATGGGTTCTACGGGGTGTACGGCATTGTCGTCTTCTACTGTTTCTTGTGACGTGGGTGCTTCAAGCTCCGGGACATGCCTTGCTGTAACGACAGTAAATTCCCAAACGGAATTATCATGGTGT
CTAAACACATTCATTGTTCGGAGTCCTCGCTTAAGAACATAGCATGGTTGAAATGTGTGTGTGATAGTGTCGGATTCAATTTGTACAATGCCTTTGAAAATAGATGGCAAATGTAGTCTGTGTACAGAATGAGATGGTGTGGCATACTCAGTTGGAGATTCTGATCTCAGTCCACCGAAGATATAAATAAAATCCCCCCACCTGCTCTGCTGAGTGACTC
GATACATATAGTTCGACTCATACGACTTAACTTGGAAGTTTTCTGGATGTGCAGTACCATTAAAGTGAACAACATGGATCCCACCTTCGCGAATCATGGCTATTCCCGGGAGGCTGCACCCACCAATAAAAATCAATGTGTTCTTTGGCCAACATAGGTGCATGTGTGGGCTAAATTTGGGATTTGCGATGGGCGCGTCCTCCAAGTAGAATCAAGT
ATTGAAAAAAAGTACAGAGTTTTGTTAA[redacted]CTGGTGAAATGAGGAGTTCACACAGTCAAGTCCTCCATACACAACACCTAGTTGACCACATTTGGTAAAAGTATGTCCAAACCTTGGAGATGGTACATATCCCCGCTGTTCCACAGTTTCCCAATTAAAAGATGCCCCTTTGAAGCAACCGGTTACAAAATCATCTGTACCATTCTGACCCCATTTCATC
CAATTTTGTCCTCCAAATACGTACACACGACCCTCGTGTTCAATACAGGCATTGCCAACACTGGGTCTTATCAGCCTTGTTTTGGTTCTCCCAGATGTCAAGCACTTGAATCGGCTTCCTTGCCTTGCATTTATGTTGTAAGAAATTGGGAAATCGTCGTCGAAGTTACACCCGCCGAAAAATAACAATTGTCGGTTGCATTCCAGCATACACCGCGA
GGAGCTGGAAACTTCTCGTTAGACCGCTCACGTATATTTCTTCAAAGAACAGTCAGAAAACACCGAAGTCATGTCACGAAAATAAGACTGAAATAATATACAGACTCACGTCTGAAAATAAGACTGCTTCGAAAGTGTAATCGTATCGTAATCGTTACGATTGTGTCACACTGTGAGTGATGCGATTTAACAATTCAACACGACACGTCGTTGCCA
AGCAGATGACCCGTCGGGTCACTCTATCACGTTCGAGTTTTAGAAATCCTAATACGTTTTTATTATTATTACAAAGTACGACATAGCAAAGATTATTTTTAAGGTTTTTAGCAATAGTAATTTATTGTAGTGATGTTAGGGGATTTTTGTGGGTTTTAATAATTATTATCAAGGTGTTAACGATGTTAATTGTAGGAGGCGCCCAGCCCCGGTAA
TTTCAAACTGGCGATTTTTTTACGGCGGTCTATGGGTATAGTTTCAGTGTATGGATGTTGGTATCCCGGGGGTGGTATTTTATTTTGAAAGGAAACCAAAGAGATTTAGCCAGATTTGCAGGTACCATATTATATTGGCTCAAATACGGTACAGACCACTCCGTAAATAATAACATAGATTTTCTTACTTATTTTGGATTTATAGAAATATCTAGA
AGTGAAAACAAAGGCAAATTTAAATTGCTTCGTTTCACTATAAAACATATGTGATTCATTTGTATTGGAGTAAATTTTTTAATCATAAATATAAACCTAGGGAAAATTACAGTTTTGTACAAAACAATTTAAGTTTAATCTTCGATATAGATGTGGAATATATTTGATGATAAGCATGCAGCAGTGTAAAAAAAAATATGCGAAATCTCTTGGTGCCC
GCACCTAAGTACCTAGAGTGCCTTAGCTCAATTTCTGAATTGATATGCTTTTCGAAAATAAAAGATAGGAAATCCGCGAGTTTAAAGGCATTGTCCGTGATCCTCTTAGAAATGGATTTTACTTTTGGTGCTGGAAAATATCTTGCAGAAGAGCTGTTAACATGTTCAACATCCCAGCTGTCGCCATGCACCCCATGCCGGGGGGGGGCATCACATAT
GAGGAA

#### Pseudogenised loci in Ophioderma brevispina

| Genomic scaffold | RAG1L | RAG2L | RAG1 status | RAG2 status . |
|---|---|---|---|---|
| JAMKCH010062600.1 | 72021-70062 | 61231-62612 | incomplete, missing N-ter | complete, 1 CDS, 1 frameshift |
| JAMKCH010009780.1 | 52461-55347 | 61714-59224 | pseudogenised | pseudogenised |
| JAMKCH010016220.1 | 56026-56922 | not found | incomplete, at scaffold margin | not found |

### Marthasterias glacialis

#### RAGL-A

##### MglRAGL-A.1

| Genomic loci | RAG1L status | RAG2L status . |
|---|---|---|
| MglRAGL-A.1 | complete (1 CDS) | incomplete, N-ter not found |

```
>CAJVRS010001272.1:29300-39700 Marthasterias glacialis genome assembly, contig: hap_tig00003862_156783_342000, whole genome shotgun sequence
CTGATTTTAAGGTAAATTTTCCTTTTATTTTTCCCAAGTTTCTATGATGGTTTTTGTGAAACCTACGATGTTGTTTGTTAAACAACACGGTGTACGAACGACTGGTTATGTTTGTATGTGCCGACATTTTCCCGTATTTGACGACTTTTATGTCAAAAAGGCCCATTTGACAGTGGATTTCATGAATTTCGTCATTCGGTTACAGGCACGTGATGGTTC
GTGTCACTGTTTATACAATACGATCAAAAGCCTGATCGGGACGCTAGTTTTTCTTTATTAAAATAATATTAAAAACATCAGCCCTGATCAGGCCAGGCCAGGGGGCCAGGGCACATCATTACAATGGCATAGTCAACACGTTAGTCTTAAATTCATGAGAATAGTATGTTAACTAAACGCTAAGGCCCCCTACCAAAAAACACTAATAATGATAATAAT
AAGTAATTATTCTGTTAAAATTTGGAATTGATTCCTTTTAGTTTAAACATTACCTTTTCTGTCTCTGATTTGTCATTTTTTTTTAAAGAAAATAAAGTCCGAAATTGTAAATCATATGGTCTTAAAAATTAAATTCCAGGTATCTCTTTACACAGTATAAACAACAATGTTGGTGAGGAACCATACTCTGATTTGTCTATTAGTTGAATTGTTTTTT
GGGGCAGTGGATATTTTGCCAAATCCGGTTATAATTTAACAAGCGTTAGTTTCATGAATGCTGCCCCATCGCAAAGTCGCCCCCTGACAGAGTCACTCCCGACTGTGTCTGGGGGCAACTTTGTGAGAGGCGACTTTGCGAGGAGGAGACATGACCGACACCCAGTTTCATGTATTGATGATGATGGACTTATACTTTAATCTTCCAGGATTTGTTTAT
ATACATTGACGATCTTCAGTCACATGTCAGTCTGATCTTGATTGGTTGTTAATGCTTTTCCTATTATTATTCCATCTTTACAGGAAGTGGCTAAGTATTTAAGTGCCATGAATCACATTGATCTGCAGCATCATCAGAGCAACCTAGAAAAATGCTGTCGGGTGTGTGGTGTCAAATACAACAGAGTTCACAAAGCCAAGGCAGGTTACACCTGCAAAG
GAGTGGACAGGAAATCTGGTTTACCATTTGCCGAGTTGTTACAACATGTATTTAACATTTTCATAGAGTACGATCACCCAGATATACATCCAAGCAAGTTTTGTCACAAATGCTACCTGGTTATATCTAGATGGTGTGTTGCACAAGACAGACAAAGACATTATAATCCCTCAACTAAACTATATGTGTGGATGGCCATGCTGACACTTGCTCATTTT
GTGCACGATTTGCAGTTATTGGAACATCGTCTAAAGGAAAGCACCCACCAGGCAAACCTAAAAAGGGAAACCCATGGTCAAATGCTACACATTTAAGTTGTTCACCCTTGTCAACACCACCAGCAGACCATGATTATGCTGAAGTGCCGATACACACACGTTTTGTTCAAGCACGAAAGAGAAAACGCGGAGATGACAAAGATGATATGCCCTCAGAAA
AGAAAAGAGTGCTTTCCACTGAACCGGATGAATTGGATAACAGTCTAGAATTGTGCAGTCTTTATAGCAGAGAACTTTGTCTGGCAGCTCCACTTCATGCATTATTTCAATGCAATATTTGTTCAGGTATTGTTGACATACCTTGCAAAACTTCATGCAATCATGTTTTCTGTTCAGGTTGTATAAAACGGTGGGTTCAAAGATTTCATTGCTGTGAAC
AATGTGATGAGTTGTTGAGTTTAAAGAATTTATCGCCATGTGACCAGCCATTCTGTGAACTTGTGCTAGATATCCCCATGAAATGTCTAACTCCTTGCAATGAACAGTTAACAGTTCGGACTTTTAAGGAGCACATTAGATCATGTAATTTCATTATGCCAGATGAACCTCAAGCAAGGGAAGATACATGTACGTCACTTAATGATAGTTTTCTTGGAA
GCTGTACAGATTCATCATCACTAGATGATAGTGTATTGTCAACATCATCAACCACATCACGAAGGGGGAGACCTCCAAAGCCCCTAATGAGCCTTAACCATACAGCACAAAATGCACGTCTTAAGACATTAAAGGAAGCTATTAAAGAGCATGCTATGAATTACAATGAAGATTTAAGGACAGTCTATTATGCATTTTTGTTGAATCATTTAAGGAGCA
ATAAAAGTTGGAAAGAACATGGGGAGGTGACATTGGGGTTCACAGGAGATTCCCACCTCAGCTCAACCAAGTGCTTGGCCATCAAGGTTTCTACTTTCATGTCAGTTACTAAGTACAGGAGAGTGCAACAAGTTACAAACTCTATGGCACAAAGAAAAAATTTCCAGCCATATTCAGCCCTAATTAAAGCTGAAAAGTCATTTCTGCCTGGCTCAGTTA
AATACCATTTGTCTCCACCAATGGTTCGACCTGACAAAAACAGTGCATCAGTAGGTGAATATTCTTATGTGCCACAAAATTTTCCAGATTTAAGACTTCCTTTTCTGAAATCTGAAAGGTACTGGTATGATGACGCATTGGCTGTGGCATTGACAGACTTGGAGGATGAAATCTGCAATGGACTTATGCAAGCAGGTGAAGATCCACAGAAATTTCTG
GCCTTTTTACTGTTAATGTCAAGGATGGAGGTGATGGCATGGGGGATGTCAAGAGAAAACAATATCAGTCCAAGGAAAAAGTGACAGACAAAGCAATTAGATACAGTTTTTGCATCTTGAAAATCACCATTGATCAAGGAGGACGTGCTGTAAGTGTGTTTGAGGAGAGTTCACCGAACTCTCAGCTCAATCACGCCCCCTTCTTATAGCTCTGGCAG
ATGAGAATGACTATTTTGCATTTACATGCCTTTTTTGCCCACTCATTCGAGAAAGAGAACACCTGCAAAACGCAAACTTACAGTTGACAAATAAATTCGACTCATTATGGATTTTTGGCTTCCAACTTCACAGCACTATGTATGATGAGAAATTAGAAAGGAAAGTTGCGGGCCTTGCCGGGTCATCCTCTGAGTTCTTGTGCACAATATGTGAAGCAA
CCAGACATGATGCCTTCTCAGACCCATTTAGCAGTGCAACTATCAGAAGTCCTCATCAAAACTTCATCAACTTGGACAAAAAAATTATCCAATGAATCTGATCTGCCTTACAACCAGCTTGTTCGGGAGGCCAAAGGAATTAAATCCGACAGCTTTTTTGATATGACCCCACACATTGACTCATTGCACTGTGAAATTAATAATGCCCTCTGGTTCAAGA
AAGTTTTGTTAGGGAAATCGCTAACCCTACAGAAAAGAAATGGAGTTATTCAAATGAAGACAAACATGACAAAGAAAAATTACAGGTTGCAGAGGAAGAGTTAAATGTCAAACTGCGGCAGGGCATTGGACTTCAAAAACACCTGATGCAACCAGGAAATTATCCAAGACTTTTGCTGAGTGAGAAATCAATGAAAATTGTTTTGGCTCTTATCCTG
AAGGTCGTCAAGGAGTGGTACAGCAGCTAGTTCATGAATATACCCTGCTAAAGCGAGTTTGGAAAGCAACAAATCCGAATGACTCTTGTAAAGAATTACTAGAAAATTACCCAGGAAATGCAAGGACATTTATGAGAACTCTACATACAAGCTTTGGTTATACTGCAGGCAACATGCCAAACTATTTACACAAAGTCATAGCTCATGTTCCTGAACTTA
TTTCAAAGTATGGTTCTGTTGGTGCTTATTCAAGTGAACCCAATGAGCATGGTAACAAACTTTTTCGAATGCTCCGCAAGATGTGCTCCAGACAAATGACAGACTTTGAGCTAAAAGACATATTGAAATTTCACTGGCTCTACACTATTAAATCTCTGCAGCGAGAGGTTGATCAGGTATCTACTCAGCAGAAATGCTCAGTTTGTGGGACATTTGGCC
ATAAGTGTAGTACATGCCCGAACAACAGCGAGCTGTAAAATCGTGTTTATAAAATAGACCTTTACCATGGTGCAGCCATCTTGATTTCCTCCCATTCTTATAAATGTAACCAAACCGAGGCTAAAAAGACAAATAGTCTGGTTTTTGTTTATAAAATGTGCATTAGCATTCGTTTCTGTTATTGGATACTGGGAACTAGACCAAGATGGTGGCAGCATG
ATAAAGGTCTATTTAATAACAATTTAGAGTTGTTGTCTATAGTTTATAAATGTACATTGTACATGCATATAACAGATTCTGTAGGTACCATCAGGGAAAAGAATCACTTGGCAACAAGTACATGTAGATTTTTTTGTTTAAAAGTTTTTACCAAGTCATAAGTAGATCTTTCTTATGGGAAAAACTTTGTGTGCTAGTTGTAATTGCTGTAAACAATCA
AAACAGACTTTAAAGAGACGACATTGTCACGAACAACATGGCATGGTAACATGGAGTGTTAGTTTGAGATAAACTATTTGGATATTGCTATAGCTGCTGCATAAACTGTGTTCATCTTAAAATCGGATCCTCATTCCATGCTCTGTTGAATATCTGCTCTGAACATCATAATATGTCTCTTTAAATGCGTCAATTTTGATACATGTACTTCATGGCAG
TTTGCATGTGGAAAGCACTAAAAAAATAGCCCCTGTGGTGTTTAAGACTTCTTCGGGTACATTTACTTGTATGTCAAATTGTAAGCCCTCAAAAGGTCTGAACTCTGGTTTGTGGTCATCATACAGTCTGCTTCAAAATTGATGTGGTGACTCACTTTTGGGGGTATTTTTGTTGTGTACAAACAGGGAAATTCATGATACAATTATCCCGTGTGTAT
GTGGGGACCAAACTTCACCACGCACAATTAAGAGAGGCCCAACAGAACCTTCATCCCCTACATTTTGTGATAAAGACAGAATTCCAATCAATCCAGCAATTTTGGCTGCAATTAGACATGAGAAAAACATATACTGAGAAACCACAGATATGGAGGGGACTTAATACATTTAAGATGCATTGACCATTTACATTTACACATGCGTGTTGGCTTCTCAGC
TTAAGTCAGGTCTGCTATTCACCCAGTTTAATTTTTGCATAATATTGATTCACTTTTGTTTTTATGGACATGCATATGTGCAATGTAAATGTGGAAATTTATAATATATTTTTGTTAATAAACTTCAAGGAAAAGTGTGAAAGTTAATTTCATGTTTTGCAATTTATTATTTACTTCAACTTGTCATAGAAATCTTTATATACATGTATGTTTAACA
AATAAAGGATGTTCGTGAACATAGTAAACAGATGAAATTTCAAATGTACATTTAATTGAATGCAGTTTCACATAAAATTTTATAAATGCCAACTACCACAATGATGCTATAGTATGTATACAAATACTCACTGCTTCGAGTGCTATAGCAGGCGCGGATCCGGGAGGGGGGGGGGGTCAACCCCCTGCTCGGGCCCAAAAATAAAAGAAACATAACAG
CGATTCTTTAATCAAACATTAAAAAAGAAGAGAATCACCCCCCCCCCCCCCGCAAAATTTCAGGATCCGCGCCTGTACAGTTACAACTACGATTCGGAGTGATCCCTCAAGTGCAAAGGGAAATTAAAATGCATCTTGGGTGTAATAGTTTTAACAATAGCACAAGTAAAAACAGTCAATATATTTTTTATAACACCTCATGAACAGATTGTGATATA
TACATTGTTTTTATGCAGTCGTCTTTAAGATCTTTTTGAAATGAATTGCGAAAATTTTCTCGCAAGAGGTCTAGAATGCACTGTACATAATCTAAAAAAGGCTACAGATTATGTTTTAACCTTTCACGTGATACACTAAAATAGACGTCTGGGTACCAAGCAGCTTCGGGTATAACACTGTGCATAGAAAGTTGCAATTCCATGCGAGCTTTCTCGCAT
AAAATCACAGGATAGCATGTTATAATAAGAAATATCACAGGCTTGCATATGCATGAGTGGATCCTACAAAAAATAGTAAACATAAGAGATTAATACTAGACAAAAGAAAGGTCTGACCCCCCCCCCCCCCCCCCCCCCAAACAAAAGCAAGAAAATTTTGATATTAAGTATAAATGCGATATTACCCGCGCACTTGTGGAATGCAGAAAAAAGTATAGC
GCTTCTGCATCACATCCTTCAGCCTCGTTGGGGTCCGTGGTTGAACATAGGACGCAGTAAGCGCTATATTTCTAGTAATCCACTCACGCTTGTGTGAAACTATGAATACCTCATGTACATGTATTCATGACATGAATTCTCACTGGTCCATTCACCAAAAGTGTTCACGCACCGGGTAAAAATACACATACTTAAGTGCATTGTAGACTGTTGCATAC
ATGTATGTGTAGCAGTGTACTCATGCGCTGCGCCACGCACACAAGAACATCGTAATTGCTACATACATATGAAAATATGGCGGCACATGTAAATACAGTCTGCTTTTTAATGCCGTACTAGAGATTATTGTGGGCAACTGAATATGTATGAGGTATAATAAAACAAACAAGACATGGTTTATTTCAGATGGACAGTCGATATTAGCTCCCCTCGGTATT
GGAAGTTGACAAAATAGTTTGTCAACTTCCAATACATGGGGAGCTAATATCAACTGTCCACCCTCAAACCATGTTTAATAAATACATATTAACTTTGTTGTATATGTTTGATTGTACGCTTGGCTGGATACATAGTAATTAGTAGAGTATGCTTATTTTATAGACAGTTTATAGATTCTGATTGGTCAAAACCTTATCACGTGGGGTTAATACACATTA
AGTTAAAACCCTTCGCTGTTAGCTGGCGCAGTATTTGACTGTACCGTTTTTAAAAATGGTGCGTTCCCTATTAACGGTACGGTCAAAAATCTTGAAATTTTACAAGTTCCTCAATGGAACATAGCAGTATAACCCTATAGCCTTCAAAATTAGCTGGGTGGTGAGTCTTAGACATACCGGCACCTTTAACCTCGACTTCCCAAAGGATACAATGCAAT
TTGAGCTCCAAGGTCTAAGATCTTAGCCGCAAACATGCAATCACTCATCCTGTCAGCGACACCCTTAGACCAGATGTTTTATTGTAAACAGTCTAGCTGTGGTTTCAGGCCAAGACAATAAAAAAAAACTTGTTTCTTAGTGTACTCATTTTGCTTGTGATAGAGTGCGAACGTTATTTTGTTTCACGATAGTCTGGTAGGCAGTGGAAACATAATTTT
ATTTTGTAGCGACTGCATGAGCTCACCCACGTATACTGGCACCACGCACACATGATTTTTTTTTTTTGCCTTTTGGGGGGACTCTGGGATTTATTGGACCTTTCTGAGTCTATTGATACCATATTAGAGGGGGTAGTGAGCAGAAAAATGCATTTTACACGCAAACTTTAACGGTACAGTCATTTACTGTACGATCCTACATCCGCGTTATTCTAAT
GTCGCCCTATGATTAACAGGCCAGTGATATAAAGACTGGCTTTGGGACTAGCGAATAAATGGCCCCTAAACCAGCACACATGTACATTGTGTACAACTAGTGTGTGCGCCTTGAGCTGTATCTTAAATTCATGCATCTTGGTTTCATTGGCACGCTGAAAGACTTGGAAAATGTGGCCCTTAAATGAGTACAACGAATGTAGGCAGTTATCAACAAA
GCAAAATTTACAAATTTTGATCGTTTAACCTTCTTTTTTTCATGAAAAGGCACTTAAATAACCCTCCTACTGATAGACCATTTTAACCCATTCGATATTCCAACTCTCATATAGCAGTTGTACACACCATTCAAACCATTTGAACAGTGTTATAAGCAAGTTGTTCCACCAATTGAAACCATTTGGGCCAATGGTTTCGAGTGGTCGCTCAACTCTCC
AGGAACAGGCACCAAGAGTAGGAGTTTTAATGGGAATAACAGTGTGAGGACCCGGTTCTTTAACCACCCAATAGTTGGGGTTGTTCCGCTCATGAGATAATGACGTTTGCATAATGCCAACAACAATTCAACCATCATTTGGTAAACAATTTTTGAATTTTGTTTCAAATTTAAAACAATAATGTCCAACCATCAA
CGAAACTATCTACTTGTTGATGACAACGAGTTTGGAACATTTCGGATATTTTTGGAAAAGGAGGAATTCTTCCTTTTTGGTCACGTGACTTTGGGTGATACCAGACCCTCAATTTGAAAGAGACTACATGTAGAATTTGGGGTCTTGGATTGGAAGCAGCAGACTATACCACGCAGTGAAGTCCGGGTTCTCACACTGTTATTCCCTAATTATAATTT
GTTTAATTTAGCAATGTCATACAGATGGTTATTACTTTACTGTATTACTTCGAAAATTAATAATGTCTGCCACTCTATTCAAAGTAAAAAAGAAAGAAATAATAACACAGTCAACAACATGCGGCACACTTCCGATAGTCTTTCATACACAGTGCATGAAACCAGTCTTGGCATAAAGGGCATTGTACCCATTTAGTACCCTTCGCTGACTTTGGGTA
TTCAAGGTACAGTTGTCTGCATTACAGAAGAGTGGCTCATCCTCATCTGATTCGGAGTCTGATGGTTCACTGTGTTCCTGATTCTGTGAAGTTACACATACTTCAGAGCTCAAAGCAGATTCAGAAGTCCAGACAGATGATGGCAAGTGGAACATATGATTCAACAGCATCGTCTGGTACATCAATTTGTTGCTCATGAAACTCTAGATCTGGAGAG
CATGTGGCTGCTGGATATCTGGACAAAGATTAGGTTCAACTGGTTGATCCTGAAATTTGTTGACTTGGGGATCTTGTGACTGATGGGTATCAGGATCTGCTAAGTTATCACATGGGTTTGATGAAGTAGATGGCCGAGAAATTAACTGCCATACAGCATTATTGTCAAAGATGACTGCTTCACGTACACCTATCTTTGCAACACCCACACATGTACCA
GAAAATTCCGTAGAAATGTTATGTGACTCAATAGAGATTTACCATTTTCAGTGACGGAATATGAAGACGGGTGATAGATCTTGCAGGTTCTACGTCGCCTTCAATGGAGGATGCTGCATATCCACCAATGATGTAGATGTAATCCCCGAGTTCACATGCATGGTGACCAGAGATATACAAATTAGAATCATATGAAGCAATGCCAACAGACTGATCG
TTAATAAAGTTCAAAAGATGTATCTGTCCCGGCCTAATCATGCGTTGATTTTAATTTAGTGTCCACCGATAATGACTGCATTGATTTGGCACCGATGTACGTATGATGCATCCACCTGAATAGAATAAAATTAAAACACTTTATTGACTACATGAAATACTGTAAAATGTTAACTCCTACCATATGGTTTCAAATACAGCAACTTTTCTGATAAAT
TAAATTTTGTACAACTACTTCGCTAAAAGTAACACTTTTTCATTCATTAAACACGTGATCAGACAGTTATTCACAAACAGCAACATACAATAGAGTTATATCGAATAACTTTAATTGCACATCCCAGAACATGATCAAAGCATGTTTTACTTTTAAACAAAGACCTACATGCTTACAAAAAAATTAAACACGTTTGGTGTTCAACAGGGTAAATGTAAT
AAAGTAGTTTCAGCTCAAAACACTAGCGCTGAATCTATAATGTTTCAAATTGAGATACTTTCAAAAGAAATAGTTTTTGTGAACAGGACAAAAAAAATTGAAAAGTTTTGAATCTGTAGTATTACCAGATCTAAATTCTGATTAAAAATGTTGTTTTAAAAAGTTGTTAATTGCCAGAATTATACTGTATAGAATGACTCAACTTACCACATTCTGAC
TGATGTAGAGTGTCCATCACTGGTGATTTGATGGTAGTTAAAAAAATACCTGAAGCACGAAAACTCATTGTTTTCTCTTTTATCTCCTTCTCAGCTCCTGATATCAGTTCTCCATCATATATAAATAAAAAATCACCCTTTTTAATGTCCTCTGTTGCAAAGACACCTCTGTCTGTAGATAAAATTATGAAATTAGTTACTTTTAATAACATTCATACATG
TATTTAAAAATTGTAATTTCTAAAAACATATCATATTGCATGAATTGTTTTGAATTTAAAAAACAACCTATAATTGTGAATCCATTGTTACTGCATGTACTGCTTGTTTCAGACTGCTGTTCTAAATGCAGCATATGTTCCAGACTAATGCATGTTAACGTAAAAGCAAAATAATAATAATAATAATAGTAATAATACATGCTTTTCCTTGATAAAG
GGGCAAAGGGCTCATGATAAAGGCACTTGGAATGTTTTACATGTTAAAGTTACATGTAGCAAGACATTTGACTTTGATTAAAATCAAAGAAATACATAATGCAAAACA
```

```
>MglRAG1L-A.1
MNHIDLQHHQSNLEKCCRVCGVKYNRVHKAKAGYTCKGVDRKSGLPFAELLQHVFNIFIEYDHPDIHPSKFCHKCYLVISRWCVAQDRQRHYNPSTKLYVWNGHADTCSFCARFAVIGTSSKGKHPPGKPKKGNPWSNATHLSCSPLSTPPADHDYAEVPIHTRFVQARKRKRGDDKDDMPSEKKRVLSTEPDELDNSLELCSLYSRELCLAAPLHALF
QCNICSGIVDIPCKTSCNHVFCSGCIKRWVQRFHCCEQCDELLSLKNLSPCDQPFCELVLDIPMKCLTPCNEQLTVRTFKEHIRSCNFIMPDEPQAREDTCTSLNDSFLGSCTDSSSLDDSVLSTSSTTSRRGRPPKPLMSLNHTAQNARLKTLKEAIKEHAMNYNEDLRTVYYALLLNHLRSNKSWKEHGEVTLGFTGDSHLSSTKCLAIKVSTFMSV
TKYRRVQQVTNSMAQRKIFQPYSALIKAEKSFLPGSVKYHLSPPMVRPDKNSASVGEYSYVPQNFPDLRLPFLKSERYWYDDALAVALTDLEDEICNGLMQAGEDPQKFSGLFTVNVKDGGDGMGDVKRKQYQSKEKVTDKAIRYSFCILKITIDQGGRAVSVFEESSPNSQLNSRPLLIALADENDYFAFTCLFCPLIREREHLQNANLQLTNKFDSL
WIFGFQLHSTMYDEKLERKVAGLAGSSSEFLCTICEATRHDAFSDPFSSATIRSPHQNFINLDKKLSNESDLPYNQLVREAKGIKSDSFFDMTPHIDSLHCEINNALWFKKVFVREIANPTEKKWSYSNEDKHDKEKLQVAEEELNVKLRQGIGLQKHLMQPGNYSRLLLSEKSMKIVLALIPEGRQGVVQQLVHEYTLLKRVWKATNPNDSCKELLEN
YPGNARTFMRTLHTSFGYTAGNMPNYLHKVIAHVPELISKYGSVGAYSSEPNEHGNKLFRMLRKMCSRQMTDFELKDILKFHWLYTIKSLQREVDQVSTQQKCSVCGTFGHKCSTCPNNSEL
```

```
>MglRAG2L-A.1_inc
MHTCTYIGANNAVIIGGLQLKNQRMIRPGQIHLLNFINDQSVGIASYDSNLYISGHHACELGDYIYIIGGYAASSIEGDVEPARSITRLHIPSLKNGKISIESHNISTEFSGTCVGVAKIGVREAVIFDNNAVWQLISRPSTSSNPCDNLADPDTHQSQDPQVNKFQDQPVEPNLCPDIQQPHSSPDLEFHEQQIDVPDDAVESYVPLAPSSVWTSESA
LSSEVCVTSQNQEHSEPSDSESDEDEPLFCNADNCTLNTQSQRRVLKWVQCPLCQDWFHALCMKDYRKCAACC
```

### MglRAGL-A.2

| Genomic loci | RAG1L status | RAG2L status . |
|---|---|---|
| MglRAGL-A.2 | complete, 1 CDS, 1 frameshift | incomplete, N-ter not found |

>CAJVRS010001383.1:4059750-4070000 Marthasterias glacialis genome assembly, contig: H_29, whole genome shotgun sequence. Reverse complement
CCTACGATGTTGTTTGTTAAACAACACAGTGTACGAACGACTGGTTATGTTTGTATGTGCCGACATTTTCCCGTATTTGACGACTTTTATGTCAAAAAGGCCCATTTGACAGTGGATTTCATGAATTTCGTCATTCGGTTACAGGCACGTGATGGTTCGTGTCACTGTGTATACAATACGATCAAAAGCCTGATCGGGACGCTAGTTTTTCTTTATTAA
AATAATATTAAAAACATCAGCCCTGATCAGGCCAGGCCAGGGGGCCAGGGCACATCATTACAATGGCATAGTCAACACGTTAGTCTTAAATTCATGAGAATAGTATGTTAACTAAACGCTAAGGCCCCCTACCAAAAAACACTAATAATGATAATAATAAGTAATTATTCTGTTAAAATTTGGAATTGATTCCTTTTAGTTTTAAACATTACCTTTTCT
GTCTCTGATTTGTCATTTTTTTTTAAAGAAAATAAAGTCCGAAATTGTAAATCATATGGTCTTAAAAATTAAATTCCAGGTATCTCTTTACACAGTATAAACAACAATGTTGGTGAGGAACCATACTCTGATTTGTCTATTAGTTGAATTGTTTTTTTGGGGCAGTGGATATTTTGCCAAATCCGGTTATAATTTAACAAGCGTTAGTTTCATGAATGC
TGCCCCATCGCAAAGTCGCCCCCTGACAGAGTCACTCCCGACTGTGTCGGGGGGCAACTTTGTGAGAGGCGACTTTGCGAGGAGGAGACATGACCGACACCCAGTTTCATGTATTGATGATGATGGACTTATACTTTAATCTTCCAGGATTTGTTTATATACATTGACGATCTTCAGTCACATGTCAGTCTTATCTTGATTGGTTGTTAATGCTTTTCC
TATTATTATTCCATCTTTACAGGAAGTGGCTAAGTATTTAAGTGCC*ATGAATCACATTGATCTGCAGCATCATCAGAGCAACCTAGAAAAATGCTGTCGGGTGTGTGGTGTCAAATACAACAGAGTTCACAAAGCCAAGGCAGGTTACACCTGCAAAGGAGTGGACAGGAAATCTGGTGTACCATTTGCCGAGTTGTTACAACATGTATTTAACATTTT*
*CATAGAGTACGATCACCCAGATATACATCCAAGCAAGTTTTGTCACAAATGCTACCTGGTTATATCTAGATGGTGTGTTGCACAAGACAGACAAAGACATTATAATCCCTCAACTAAACTATATGTGTGGAATGGCCATGCTGACACTTGCTCATTTTGTGCACGATTTGCAGTTATTGGAACATCGTCTAAAGGAAAGCACCCACCAGGCAAACCTAA*
*AAAGGGAAACCCATGGTCAAATGCTACACATTTAAGTTGTTCACCCTTGTCAACACCACCAGCAGACCATGATTATGCTGAAGTGCCGATACACACACGTTTTGTTCAAGCACGAAAGAGAAAACGCGGAGATGACAAAGATGATATGCCCTCAGAAAAGAAAAGAGTGCTTTCCACTGAACCGGATGAATTGGATAACAGTCTAGAATTGTGCAGTCT*
*TTATAGCAGAGAACTTTGTCTGGCAGCTCCACTTCATGCATTATTTCAATGCAATATTTGTTCAGGTATTGTTGACATACCTTGCAAAACTTCATGCAATCATGTTTTCTGTTCAAGTTGTATAAAACGGTGGGTTCAAAGATTTCATTGCTGTGAACAATGTGATGAGTTGTTGAGTTTAAAGAATTTAAATTTG ACCAGCCATTCTGTGAACTTGT*
*GCTAGATATCCCC*ATGAAATGTTTAACTCCTTCTTGCAATGAACAGTTAACAGTTCGGACTTTTAAGGAGCACATTAGATCATGTAATTTCATTATGCCAGATGAACCTCAAGCAAGGGAAGATACATGTACGTCACTTAATGATAGTTTTCTTGGAAGCTGTACAGATTCATCATCACTAGATGATAGTGTATTGTCAACATCATCAACCACATCACG
AAGGGGGAGACCTCCAAAGCCCCTAATGAGCCTTAACCATACAGCACAAAATGCACGTCTTAAGACATTAAAGGAAGCTATTAAAGAGCATGCTATGAATTACAATGAAGATTTAAGGACAGTCTATTATGCACTTTTGTTGAATCATTTAAGGAGCAATAAAAGTTGGAAAGAACATGGGGAGGTGACATTGGGGTTCACAGGAGATTCCCACCTCAG
CTCAACCAAGTGCTTGGCCATCAAGGTTTCTACTTTCATGTCAGTTACTAAGTACAGGAGAGTGCAACAAGTTACAAACTCTATGGCACAAAGAAAAATTTTCCAGCCATATTCAGCCCTAATTAAAGCTGAAAAGTCATTTCTGCCTGGCTCAGTTAAATACCATTTGTCTCCACCAATGGTTCGACCTGACAAAAACAGTGCATCAGTAGGTGAATA
TTCTTATGTGCCACAAAATTTTCCAGATTTAAGACTTCCTTTTCTGAAATCTGAAAGGTACTGGTATGATGACGCATTGGCTGTGGCATTGACAGACTTGGAGGATGAAATCTGCAATGGACTTATGCAAGCAGGTGAAGATCCACAGAAATTTTCTGGCCTTTTTACTGTTCATGTCAAGGATGGAGGTGATGGCATGGGGGATGTCAAGAGAAAACA
ATATCAGTCCAAGGAAAAAGTGACAGACAAAGCAATTAGATACAGTTTTTGCATCTTGAAAATCACCATTGATCAAGGAGGACGTGCTGTAAGTGTGTTTGAGGAGAGTTCACCGAACTCTCAGCTCAATTCACGCCACCTTCTTATAGCTCTGGCAGATGAGAATGACTATTTTGCATTTACATGCCTTCTTTGCCCACTCATTCGAGAAAGAGAACA
CCTGCAAAACGCAAACTTACAGTTGACAAATAAATTCGACTCATTATGGATTTTTGGCTTCCAACTTCACAGCACTATGTATGATGAGAAATTAGAAAGGAAAGTTGCGGGCCTTGCCGGGTCATCCTCTGAGTTCTTGCACAATATGTGAAGCAACCAGACATGATGCCTTCTCAGACCCATTTAGCAGTGCAACTATCAGAAGTCCTCATCAAAA
CTTCATCAACTTGGACAAAAAATTATCCAATGAATCTGATCTGCCTTACAACCAGCTGGTTCGGGAGGCCAAAGGAATTAAATCCGACAGCTTTTTGATATGACCCCACACATTGACTCATTGCACTGTGAAATTAATAATGCCCTCTGGTTCAAGAAAGTTTTTGTAGGGAAATCGCTAACCCTACAGAAAGAAATGGAGTTATTCAAATGAAGA
CAAACATGACAAAGAAAAATTACAGGTTGCAGAGGAAGAGCTAAATGTCAAACTGCGGCAGGGCATTGGACTTCAAAAACACCTGATGCAACCAGGAAATTATTCAAGACTTTTGCTGAGTGAGAAATCAATGAAAATTGTTTTGGCTCTTATTCCTGAAGGTCGTCAAGGAGTGGTACAGCAGCTAGTTCATGAATATACCCTGCTAAAGCGAGTTTG
GAAAGCAACAAATCCGAATGACTCTTGTAAAGAATTACTAGAAAATTACCCAGGAAATGCAAGGACATTTATGAGAACTCTACATACAAGCTTTGGTTATACTGCAGGCAACATGCCAAACTATTCTACACAAAGTCATAGCTCATGTTCCTGAACTATTTCAAAGTATAGGTTCTGTTGGTTCTAGGAGCCATTCAAGTGAACCCAATGAGCATGGTAACAAACT
TTTTCGAATGCTCCGCAAGATGTGCTCCAGACAAATGACAGACTTTGAGCTAAAAGACATATTGAAATTTCACTGGCTCTACACTATTAAATCTCTGCAGCGAGAGGTTGATCAGGTATCTACTCAGCAGAAATGCTCAGTTTGTGGGACATTTGGCCATAAGTGTAGTACATGCCCGAACAACAGCGAGCTGTAAAATCGTGTTTATAAAATAGACCT
TTACCATGGTGCAGCCATCTTGATTTCTCCCCATTCTTATAAATGTAACCAAACCGAGGCTAAAAAGACAAATAGTCTGGTTTTTGTTTATAAAATGTGCATTAGCATTCGTTTCTGTTATTGGATACTGGGAACTAGACCAAGATGGTGGCAGCATGATAAAGGTCTATTTGATAACAATTTAGAGTTGTGTCTATAGTTATAAATGTACATTGTA
CATGCATATAACAGATTCTGTAGGTACCATCAGGGAAAAGATCACTTGGCAACAAGTACATGTAGATTTTTTTGTTTAAAAGTTTTTACCAAGTCATAAGTAGATCTTTCTTATGGGAAAAACTTTGTGTGCTAGTTGTAATTGCTGTAAACAATCAAAACAGACTTTAAAGAGACGACATTGCACGAACAACATGGTAACATGGAGTGTA
GTTTGAGATAAACTATTTGGATATTGCTATAGCTGCTGCATAAACTGTGTTCATCTTAAAATCGGATCCTCATTCCATGCTCTGTTGAATATCTGCTCTGAACATCATAATATTGTCTTTTAAATGCGTCAATTTTGATACATGTACTTCATGGCAGTTTCATGTGGAAAGCACTAAAAAAAAAATAGCCCCTGTGGTGTTTAAGACTTCTTCGGGTA
CATTTACTTGTATGTCAAATTGTAAGCCCTGAAAAGGTCTGAACTCTGGTTTGTGGTCATCATACAGTCTGCTTCAAAATTGATGTGGTGACTCACTTTTGGGGGTATTTTTGTTGTGTACAAACAGGGAAATTCATGATACAATTATCCCGTGTGTATGTGGGGACCAAACTTCACCACGCACAATTAAGAGAGGCCCAACAGAACCTTCATCCCCTA
CATTTTGTGATAAAGACAGAATTCCAATCAATCCAGCAATTTTGGCTGCAATTAGACATGAGAAAACACATACTGAGAAACCACAGATATGGAGGGGACTAATACATTTAAGATGCATTGACCATTTACATTTACACATGCGTGTTGGCTTCTCAGCTTAAGTCAGGTCTGCTATTTCACCCAGTTTTAATTTTTGCATAATATTGATTCACTTTTG
TTTTTATGGACATGCATATGTGCAATGTAAATGTGGAAATTTATATTATATTTTTGTTAATAAACTTCAAGGAAAAGTGTGAAAGTTAATTTCATGTTTTGCAATTTATTATTTACTTCAACTTGTCATAGAAATCTTTATATACATGTATGTTTAACAAATAAAGGATGTTCGTGAACATAGTAAACAGATGAAATTTCAAATGTACATTTAATTGAA
TGCAGTTTCACATAAAATTTTATAAATGCCAACTACCACAATGATGCTATAGTATGTATACAAATACTCACTGCTTCGAGTGCTATAGCAGGCGCGGATCCGGGAGGGGGGGGGTCAACCCCCTGCTCGGGCCCAAAAATAAAAAGAAACATAACAGCGATTCTTTAATAAAACATTAAAAAAAGAAGAGAATCACCCCCCCCCCCCCCCCGCAAAAT
TTCAGGATCCGCGCCTGTACAGTTACAACTACGATTCGGAGTGATCCCTCAAGTGCAAAGGGAAATTAAAATGCATCTTGGGTGTAATAGTTTTAACAATAGCACAAGTAAAAACAGTCAATATATTTTTTATAACACCTCATGAACAGATTGTGATATATACATTGTTTTATTGCAGTCGTCTTTAAGATCTTTTTGAAATGAATTGCGAAAATTTCA
TCGCAAGAGGTCTGTAGAATGCACTGTACATAATCTAAAAAAGGCTACAGATTATGTTTTAACCTTTCACGTGATACACTAAAATAGACGTCTGGGTACCAAGCAGCTTCGGGTATAACACTGTGCATAGAAAGTTGCAATTCCATGCGAGCTTTCTCGCATAAAATCACAGGATAGCATGTTATAATAAGAAATATCACAGGCTTGCATATGCATGAG
TGGATCCTACAAAAAATAGTAAACATAAGAGATTAATACTAGACAAAAAGAAAGGTCTGACCCCCCCCCCCCCCAAACAAAAGGCAAGAAAATTTTGATATTAAGTATAAATGCGATATTACCCGCGCACTTGTGGAATGCAGAAAAAAAGTATAGCGCTTCTGCATCACATCCTTCAGCCTCGTTGGGGTCCGTGGTTGAACATAGGACGCAGTAAGCG
CTATATTTTCTAGTAATCCACTCACGCTTGTGTGAAACTATGAATACCTCATGTACATGTATTCATGACATGAATTCTCACTGGTCCATTCACCAAAAGTGTTCACGCACCGGGTAAAAATACACATACTTAAGTGCATTGTAGACTGTTGCATACATGTATGTGTAGCAGTGTACTCATGCGCTGCGCACACAAGAACATCGTAATTGCTACA
TACATATGAAAATATGGCGGCACATGTAAATACAGTCTGCTTTTTAATGCCGTACTAGAGATTATTGTGGGCAACTGATTATGTATGAGGTATAATAAAACAAACAAGACATGGTTTATTTCAGATGGACAGTCGATATTAGCTCCCCTCGGTATTGGAAGTTGACAAAATAGTTTGTCAACTTCCAATACATGGGGAGCTAATATCAACTGTCCACCC
TCAAACCATGTTTAATAAATACATATTAACTTTGTTGTTTATGTTTGATTGTACGCTTGGCTGGATACATAGTAATTAGTAGAGTATGCTTATTTTATAGACAGTTTATAGATTCTGATTGGTCAAAACCTTATCACGTGGGGTTAATACACATTAAGTTAAAACCCTTCGCTGTTAGCTGGCGCAGTATTTGACTGTACCGTTTTTTAAAAATGGCGC
GTTCCCTATTAACGGTACGGTCAAAATCTTGAAATTTTACAAGTTCCTCAATGGAACATAGCAGTATAACCCTATAGCCTTCAAAATTAGCTGGGTGGTGAGTCTTAGACATACCGGCACCTTTAACCTCGACTTCCCAAAGGATACAATGCAATTTGATAAACTCCAAGGTCTAAGATCTTAGCCGCAAACATGCAATCACTCATCCTGTCAGCGAC
ACCCTTAGACCAGATGTTTTATTGTAAACAGTCTAGCTGTGGTTTCAGGCCAAGACAATAAAAAAAAACTTGTTTCTTTAGTGTACTCATTTTGCTTGTGATAGAGTGCGAACGTTATTTTGTTTCACGATAGTCTGGTAGGCAGTGGAAACATAATTTTATTTTTGTAGCGACTGCATGAGCTCACCCACGTATACTGGCACTACGCACACATTGATTT
TTTTTTTGCCTTTTGGGGGACTCTGGGTATTTATTGGACCTTTCTGAGTCTATTGATACCATATTAGAGGGGGTAGTGAGCAGAAAAATGCATTTTACACGCAAACTTTAACGGTACAGTCATTTACTGTACGATCCTACATCCGCGTTATTCTAATGTCACCCTATGATTAACAGGCCAGTGATATAAAGACTGGCTTTGGGAACTAGCGAATAAA
TGGCCCTAAACCAGCACACATGTACATTGTGTACAACTAGTGTGTGCGCCTTGAGCTGTATCTTAAATTCATGCATCTTGGTTTCATTGGCACGCTGAAAGACTTGGTAAAATGTGGCCCTAAATGAGTACAACGAATGTAGGCAGTTATCAACAAAGCAAAATTTACAAATTTTGATCGTTTTAACCTTCTTTTTTCATGAAAGGCACTTAAAT
AACCCTCCTACTGATAGACCATTTTAACCCATTCAATATTCCAACTCTCATATAGCAGTTGTACACACCATTCAAACCATTTGAACAGTGTTATAAGCAAGTTGTTCCACCAATTGAAACCATTTGGGCCAATGGTTTCGAGTGGTCGCTCAACTCTCCAGGAACAGGCACCAAGAGTAGGAAGTTTTTAATGGGAATAACAGTGTGAGGACCCGGTCT
TCACTGCGTGGTAATGACCTTGTTAACCACCCAATAGTTGGGGTTGTTCCGCTCATGAGATAATGACGTTTGCATAATGCCAACAACAATTCAACCATCATTTGGTAAACAATTTTTGAATTTTGTTTCAAATTTAAAACAATAATGTCCAACCATCAACGAAACTATCTACTTGTTGATGACAACGAGTTTGGAACATTTCGGATATTTTTGGAAAAG
GAGGAATTCTTCCTTTTTGGTCACGTGACTTTGGGTGATACCAGACCCTCAATTTGAAAGAGACTACATGTAGAATTTGGGGTTCTTGGATTGGAAGCAGCAGACTATACCACGCAGTGAAGTCCGGGTTCTCACACTGTATTCCCTAATTATAATTTGTTTAATTTAGCAATGTCATACAGATGGTTATTACTTTACTGTATTACTTCGAAAATTAA
TAATGTCTGCCACTCTATGCAAAGTAAAAAAGAAAGAAATAATAACACAGTCAACAACATGCGGCACACTTCCGATAGTCTTTCATACACAGTGCATGAAACCAGTCTTGGCATAAAGGGCATTTTACCCATTTTAGTACCCTTCGCTGACTTTGGGTATTCAAGGTACAGTTGTCTGCATTACAGAAGAGTGGCTCATCCTCATCTGATTCGGAGTCT
GATGGTTCACTGTGTTCCTGATTCTGTGAAGTTACACATACTTCAGAGCTCAAAGCAGATTCAGAAGTCCAGACAGATGATGGTGCAAGTGGAACATATGATTCAACAGCATCGTCTGGTACATCAATTTGTTGCTCGTGAAACTCTAGATCTGGAGAGCTATGTGGCTGCTGGATATCTGGACAAAGATTAGGTTCAACTGGTTGATCCTGAACTTTG
TTGACTTGGGGATCTTGTGACTGATGGGTATCAGGATCTGCTAAGTTATCACATGGGTTTGATGAAGTAGATGGCCGAGAAATTAACTGCCATACAGCATTATTGTCAAAGATGACTGCTTCACGTACACCTATCTTTGCAACACCCACACATGTACCAGAAAATGCCGTAGAAATGTTATGTGACTCAATAGAGATTTTACCATTTTCAGTGACGGA
ATATGAAGACGGGTGATAGATCTTTGCAGGTTCTACGTCGCCTTCAATGGAGGATGCTGCATATCCACCAATGATGTAGATGTAATCCCCGAGTTCACATGCATGGTGACCAGAGATATACAAATTAGAATCATATGAAGCAATGCCAACAGACTGATCGTTAATAAAGTTCAAAAGATGTATCTGTCCCGGCCTAATCATGCGTTGATTTTTAATTGT
AGTCCACCGATAATGACTGCATTGTTGGCACCGATGTACGTACATGTATGCATCCACTGAATAGAATAAAATTAAAACACTTTATTGACTACATGAAATACTGTAAAATGTTAACTCCTACCATATGGTTTCAAATACAGCAACTTTTCTGATAAATTAAATTTTGTACAACTACTTCGCTAAAAGTAACACTATTTCATTCATTAAACACGTGATC
AGACAGTTATTCACAAACAGCAACATACAATAGAGTTATATCATAATCAACTTTAATTGCACATCCCAGAACATGATCAAAGCATGTTTTACTTTAAACAAAGACCTACATGCTTACAAAAAAATTTAAACACGTTTGGTGTTCAACAGGGTAAATGTAATAAAGTAGTTTCAGCTCAAAAACACTAGCGCTGAATCTATAATGTTTCAAATTGAGATACT
TTCAAAAGAAATAGTTTTTGTGAACAGGACAAAAAAAATTGAAAAGTTTTGAATCTGTAGTATTACCAGATCTAAATTCTGATTAAAAATGTTGTTTTAAAAAGTTGTTAATTGCCAGAATTATACTGTATAGAATGACTCAACTTACCACATTCTGACTGATGTAGAGTGTCCATCACTGGTGATTTGATGGTAGTAAAAAAATACCTGAAGCACGA
AAACTCATTGTTTTCTCTTTTATCTCCTTCAGCTCCTGATATCAGTTCTCCATCATATATAAATAAAAAATCACCCTTTTTAATGTCCTCTGTTGCAAAGACACCTCTGTCTGTGGATAAAATTATGAAATTAGTTACTTTTAATGATAACATTCATACATGTATTTAAAAATTGTAATTTTCTAAAAACATATATTGCATGAATTGTTTTGAATTTAA
AAAACAACCTATAATTGTGAATCCATTGTTACTGCATGTACTGCTTGTTTCAGACTGCTGTTCTAAATGCAGCATATGTTCCAGACTAATGCATGTTAACGTAAAAGCAAAATAATAATAATAACAATAATAATAATAATAGTAATAATACATGCTTTTCCTTTGATAAAGGGGCAAA

>MglRAG1L-A.2_inc
MKCLTPSCNEQLTVRTFKEHIRSCNFIMPDEPQAREDTCTSLNDSFLGSCTDSSSLDDSVLSTSSTTSRRGRPPKPLMSLNHTAQNARLKTLKEAIKEHAMNYNEDLRTVYYALLLNHLRSNKSWKEHGEVTLGFTGDSHLSSTKCLAIKVSTFMSVTKYRRVQQVTNSMAQRKIFQPYSALIKAEKSFLPGSVKYHLSPPMVRPDKNSASVGEYSYVP
QNFPDLRLPFLKSERYWYDDALAVALTDLEDEICNGLMQAGEDPQKFSGLFTVHVKDGGDGMGDVKRKQYQSKEKVTDKAIRYSFCILKITIDQGGRAVSVFEESSPNSQLNSRHLLIALADENDYFAFTCLLCPLIREREHLQNANLQLTNKFDSLWIFGFQLHSTMYDEKLERKVAGLAGSSSEFLCTICEATRHDAFSDPFSSATIRSPHQNFINL
DKKLSNESDLPYNQLVREAKGIKSDSFFDMTPHIDSLHCEINNALWFKKVFVREIANPTEKKWSYSNEDKHDKEKLQVAEEELNVKLRQGIGLQKHLMQPGNYSRLLLSEKSMKIVLALIPEGRQGVVQQLVHEYTLLKRVWKATNPNDSCKELLENYPGNARTFMRTLHTSFGYTAGNMPNYLHKVIAHVPELISKYGSVGAYSSEPNEHGNKLFRML
RKMCSRQMTDFELKDILKFHWLYTIKSLQREVDQVSTQQKCSVCGTFGHKCSTCPNNSEL

>MglRAG2L-A.2_inc
MHTCTYIGANNAVIIGGLQLKNQRMIRPGQIHLLNFINDQSVGIASYDSNLYISGHHACELGDYIYIIGGYAASSIEGDVEPARSITRLHIPSLKNGKISIESHNISTEFSGTCVGVAKIGVREAVIFDNNAVWQLISRPSTSSNPCDNLADPDTHQSQDPQVNKFQDQPVEPNLCPDIQQPHSSPDLEFHEQQIDVPDDAVESYVPLAPSSVWTSESA
LSSEVCVTSQNQEHSEPSDSESDEDEPLFCNADNCTLNTQSQRRVLKWVQCPLCQDWFHALCMKDYRKCAACC

### Zoroaster sp.

#### RAGL-A

##### Pseudogenised loci in Zoroaster sp. YZ-2022 isolate SQW42HX01:

| Genomic scaffold | RAG1L | RAG2L | RAG1 status | RAG2 status . |
|---|---|---|---|---|
| JAQQFT010000002.1 | 32200489-32197496 | 32191714-32193439 | pseudogenised | pseudogenised |
| JAQQFT010000002.1 | 18291459-18286232 | not found | pseudogenised | not found |
| JAQQFT010000001.1 | not found | 74095850-74097306 | not found | pseudogenised |
| JAQQFT010000001.1 | 68992521-68989034 | not found | pseudogenised | not found |
| JAQQFT010000001.1 | 102181225-102183331 | not found | pseudogenised | not found |
| JAQQFT010000022.1 | 3654173- 3652413 | not found | pseudogenised | not found |

### Ophionereis fasciata

#### RAGL-A

| Genomic scaffold | RAG1L | RAG2L | RAG1 status | RAG2 status . |
|---|---|---|---|---|
| CZLG013968190.1 | 1-1901 | not found | incomplete | not found |

#### RAGL-B

| Genomic scaffold | RAG1L | RAG2L | RAG1 status | RAG2 status . |
|---|---|---|---|---|
| CZLG010623336.1 | 1596-64 | not found | incomplete | not found |
