## Supplemental File S2 for "Insights into RAG evolution from the identification of “missing link” family A *RAGL* transposons"

### Supplementary File S2

#### A. RAG1/RAG1L Multiple Sequence Alignment

**Contacts:**

- RAG1-DNA interface
- RAG1-RAG1 interface
- RAG1-RAG2 same dimer
- RAG1-RAG2 opposite dimer

**Secondary structure (SS) :**

- Helix
- Beta sheet
- Coil

$SS_{cEM}$ – CryoEM structure;
$SS_{pred}$ – Predicted structure (consensus)

**Abbreviations:**

Mmu : *Mus musculus*
Dre: *Danio rerio*
Pfl: *Ptychodera flava*
Osp: *Ophiothix spiculata*
Obr: *Ophioderma brevispina*
Mgl: *Marthasterias glacialis*
Spu: *Strongylocentrotus purpuratus*
Bbe: *Branchiostoma belcheri*

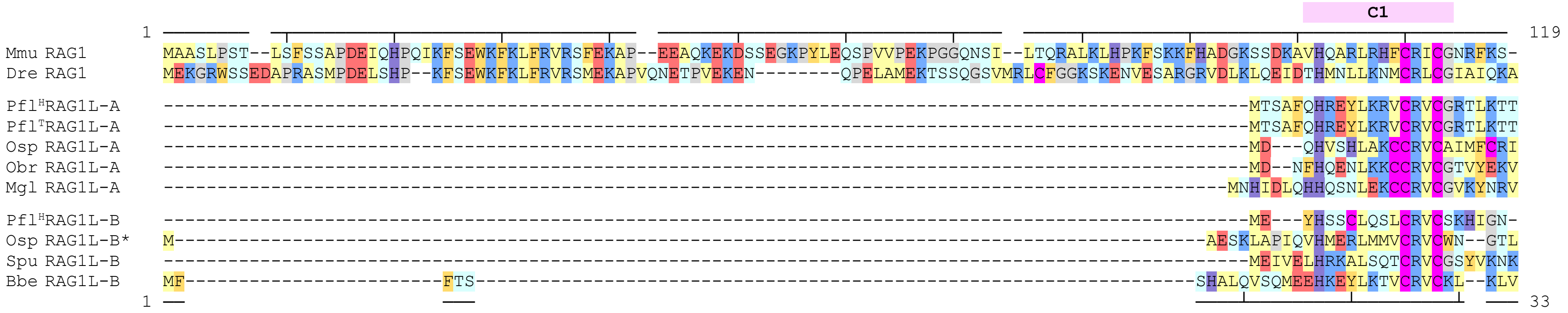


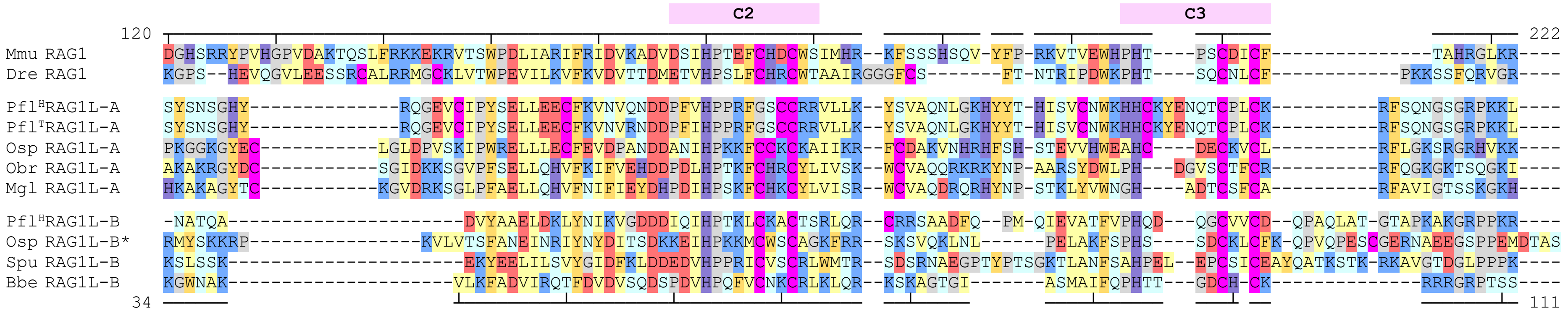

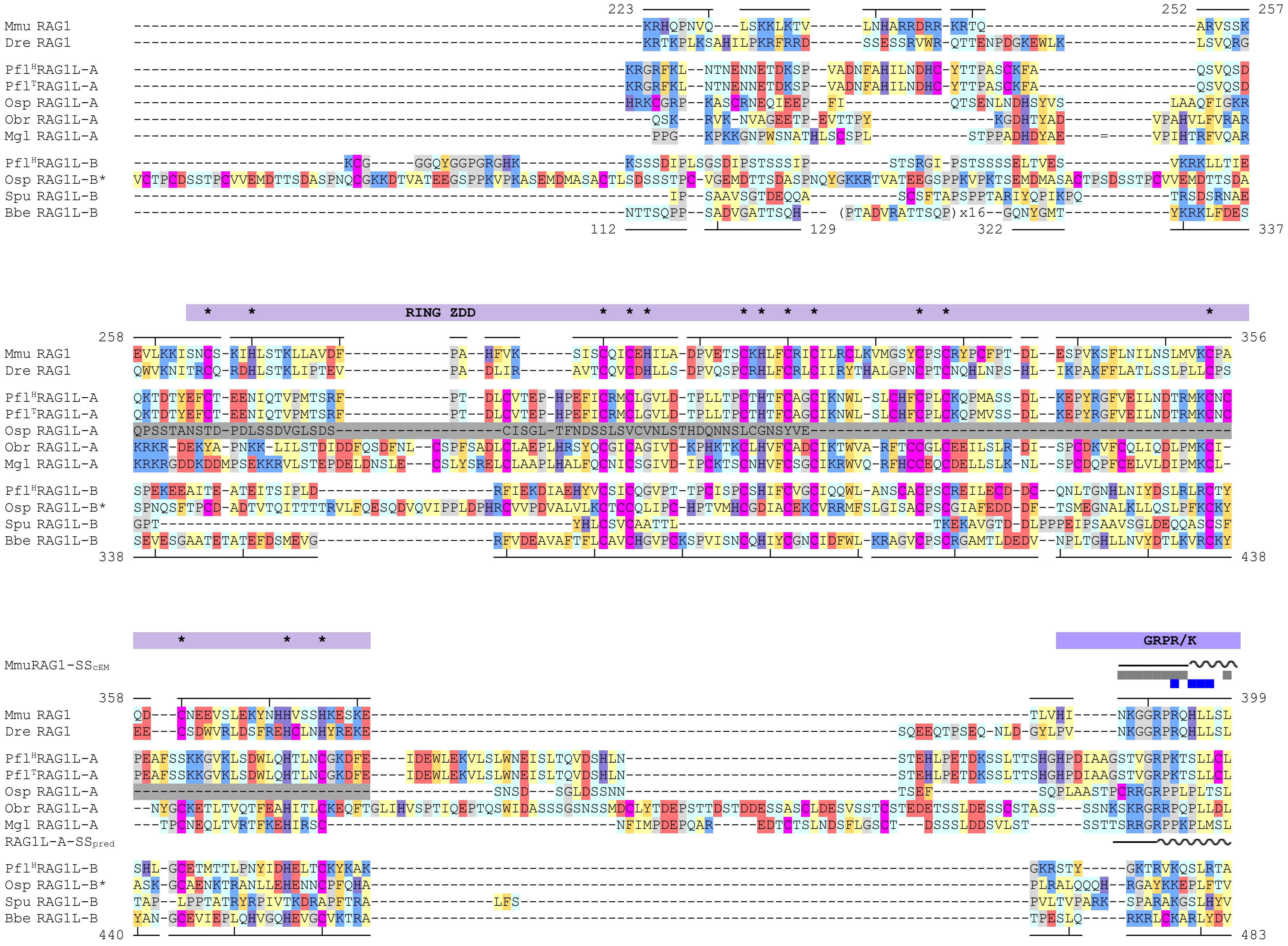

223
252
257
Mmu RAG1
Dre RAG1
Pfl[H]RAG1L-A
Pfl[T]RAG1L-A
Osp RAG1L-A
Obr RAG1L-A
Mgl RAG1L-A
Pfl[H]RAG1L-B
Osp RAG1L-B*
Spu RAG1L-B
Bbe RAG1L-B
(PTADVRATTSQP)x16
112
129
322
337
RING ZDD
258
356
338
438
GRPR/K
MmuRAG1-SScEM
358
399
RAG1L-A-SSpred
440
483

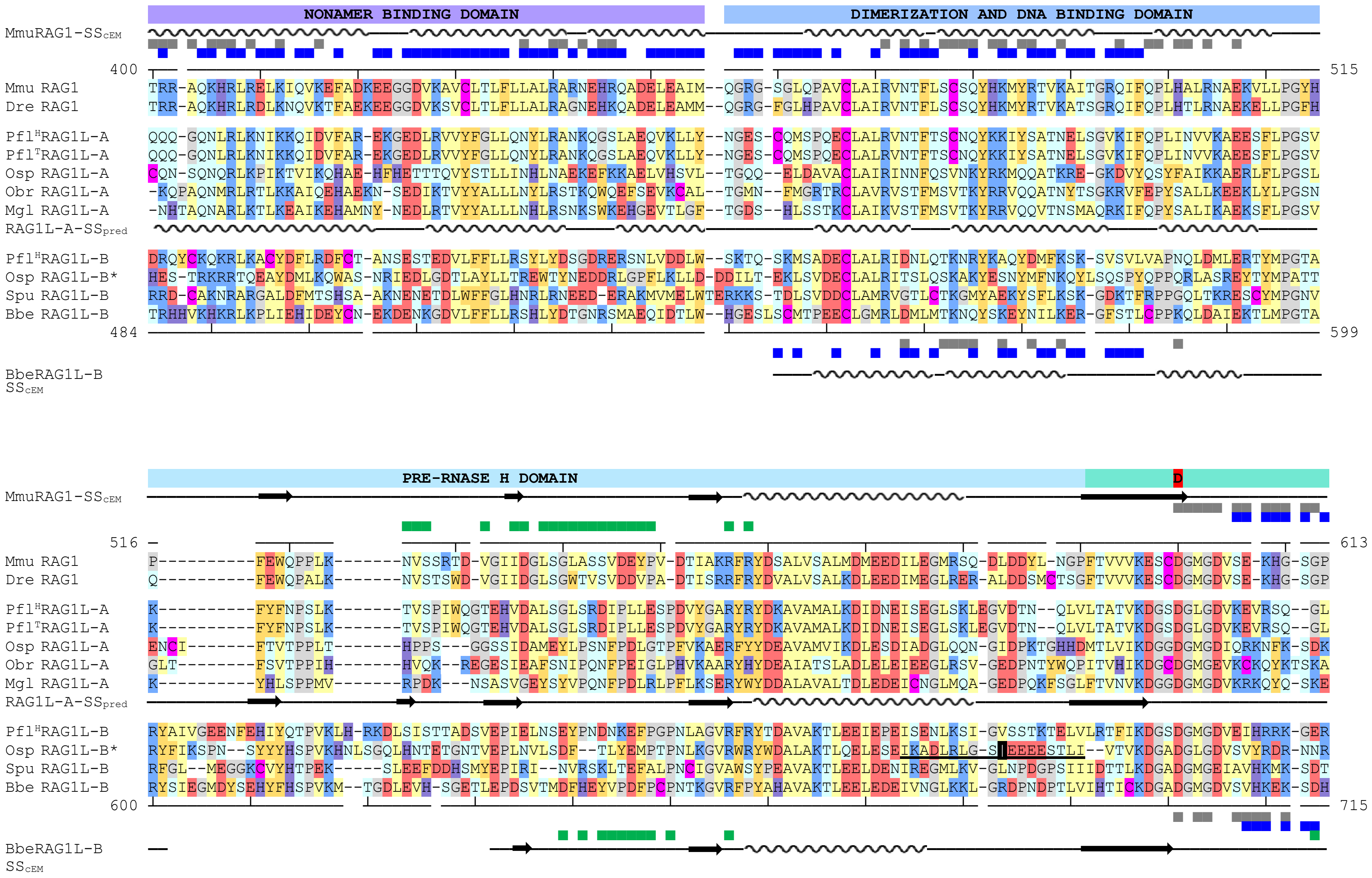

NONAMER BINDING DOMAIN
DIMERIZATION AND DNA BINDING DOMAIN
MmuRAG1-SScEM
400
515
Mmu RAG1 TRR-AQKHRLRELKIQVKEFADKEEGGDVKAVCLTLFLLALRARNEHRQADELEAIM--QGRG-SGLQPAVCLAIRVNTFLSCSQYHKMYRTVKAITGRQIFQPLHALRNAEKVLLPGYH
Dre RAG1 TRR-AQKHRLRDLKNQVKTFAEKEEGGDVKSVCLTLFLLALRAGNEHKQADELEAMM--QGRG-FGLHPAVCLAIRVNTFLSCSQYHKMYRTVKATSGRQIFQPLHTLRNAEKELLPGFH
Pfl^H RAG1L-A QQQ-GQNLRLKNIKKQIDVFAR-EKGEDLRVVYFGLLQNYLRANKQGSLAEQVKLLY--NGES-CQMSPQECLALRVNTFTSCNQYKKIYSATNELSGVKIFQPLINVVKAEESFLPGSV
Pfl^T RAG1L-A QQQ-GQNLRLKNIKKQIDVFAR-EKGEDLRVVYFGLLQNYLRANKQGSLAEQVKLLY--NGES-CQMSPQECLALRVNTFTSCNQYKKIYSATNELSGVKIFQPLINVVKAEESFLPGSV
Osp RAG1L-A CQN-SQNQRLKPIKTVIKQHAE-HFHETTTQVYSTLLINHLNAEKEFKKAELVHSVL--TGQQ--ELDAVACLAIRINNFQSVNKYRKMQQATKRE-GKDVYQSYFAIKKAERLFLPGSL
Obr RAG1L-A -KQPAQNMRLRTLKKAIQEHAEKN-SEDIKTVYYALLLNYLRSTKQWQEFSEVKCAL--TGMN--FMGRTRCLAVRVSTFMSVTKYRRVQQATNYTSGKRVFEPYSALLKEEKLYLPGSN
Mgl RAG1L-A -NHTAQNARLKTLKEAIKEHAMNY-NEDLRTVYYALLLNHLRSNKSWKEHGEVTLGF--TGDS--HLSSTKCLAIKVSTFMSVTKYRRVQQVTNSMAQRKIFQPYSALIKAEKSFLPGSV
RAG1L-A-SSpred
Pfl^H RAG1L-B DRQYCKQKRLKACYDFLRDFCT-ANSESTEDVLFFLLRSYLYDSGDRERSNLVDDLW--SKTQ-SKMSADECLALRIDNLQTKNRYKAQYDMFKSK-SVSVLVAPNQLDMLERTYMPGTA
Osp RAG1L-B* HES-TRKRRTQEAYDMLKQWAS-NRIEDLGDTLAYLLTREWTYNEDDRLGPFLKLLD-DDILT-EKLSVDECLALRITSLQSKAKYESNYMFNKQYLSQSPYQPPQRLASREYTYMPATT
Spu RAG1L-B RRD-CAKNRARGALDFMTSHSA-AKNENETDLWFFGLHNRLRNEED-ERAKMVMELWTERKKS-TDLSVDDCLAMRVGTLCTKGMYAEKYSFLKSK-GDKTFRPPGQLTKRESCYMPGNV
Bbe RAG1L-B TRHHVKHKRLKPLIEHIDEYCN-EKDENKGDVLFFLLRSHLYDTGNRSMAEQIDTLW--HGESLSCMTPEECLGMRLDMLMTKNQYSKEYNILKER-GFSTLCPPKQLDAIEKTLMPGTA
484
599
BbeRAG1L-B SScEM
PRE-RNASE H DOMAIN
D
MmuRAG1-SScEM
516
613
Mmu RAG1 P----------FEWQPPLK-------NVSSRTD-VGIIDGLSGLASSVDEYPV-DTIAKRFRYDSALVSALMDMEEDILEGMRSQ-DLDDYL-NGPFTVVVKESCDGMGDVSE-KHG-SGP
Dre RAG1 Q----------FEWQPALK-------NVSTSWD-VGIIDGLSGWTVSVDDVPA-DTISRRFRYDVALVSALKDLEEDIMEGLRER-ALDDSMCTSGFTVVVKESCDGMGDVSE-KHG-SGP
Pfl^H RAG1L-A K----------FYFNPSLK-------TVSPIWQGTEHVDALSGLSRDIPLLESPDVYGARYRYDKAVAMALKDIDNEISEGLSKLEGVDTN--QLVLTATVKDGSDGLGDVKEVRSQ--GL
Pfl^T RAG1L-A K----------FYFNPSLK-------TVSPIWQGTEHVDALSGLSRDIPLLESPDVYGARYRYDKAVAMALKDIDNEISEGLSKLEGVDTN--QLVLTATVKDGSDGLGDVKEVRSQ--GL
Osp RAG1L-A ENCI--------FTVTPPLT-------HPPS---GGSSIDAMEYLPSNFPDLGTPFVKAERFYYDEAVAMVIKDLESDIADGLQQN-GIDPKTGHHDMTLVIKDGGDGMGDIQRKNFK-SDK
Obr RAG1L-A GLT---------FSVTPPIH-------HVQK--REGESIEAFSNIPQNFPEIGLPHVKAARYHYDEAIATSLADLELEIEEGLRSV-GEDPNTYWQPITVHIKDGCDGMGEVKCKQYKTSKA
Mgl RAG1L-A K----------YHLSPPMV-------RPDK---NSASVGEYSYVPQNFPDLRLPFLKSERYWYDDALAVALTDLEDEICNGLMQA-GEDPQKFSGLFTVNVKDGGDGMGDVKRKQYQ-SKE
RAG1L-A-SSpred
Pfl^H RAG1L-B RYAIVGEENFEHIYQTPVKLH-RKDLSISTTADSVEPIELNSEYPNDNKEFPGPNLAGVRFRYTDAVAKTLEEIEPEISENLKSI-GVSSTKTELVLRTFIKDGSDGMGDVEIHRRK-GER
Osp RAG1L-B* RYFIKSPN--SYYYHSPVKHNLSGQLHNTETGNTVEPLNVLSDF--TLYEMPTPNLKGVRWRYWDALAKTLQELESEIKADLRLG-SIEEEESTLI--VTVKDGADGLGDVSVYRDR-NNR
Spu RAG1L-B RFGL--MEGGKCVYHTPEK-----SLEEFDDHSMYEPIRI--NVRSKLTEFALPNCIGVAWSYPEAVAKTLEELDENIREGMLKV-GLNPDGPSIIIDTTLKDGADGMGEIAVHKMK-SDT
Bbe RAG1L-B RYSIEGMDYSEHYFHSPVKM--TGDLEVH-SGETLEPDSVTMDFHEYVPDFPCPNTKGVRFPYAHAVAKTLEELEDEIVNGLKKL-GRDPNDPTLVIHTICKDGADGMGDVSVHKEK-SDH
600
715
BbeRAG1L-B SScEM

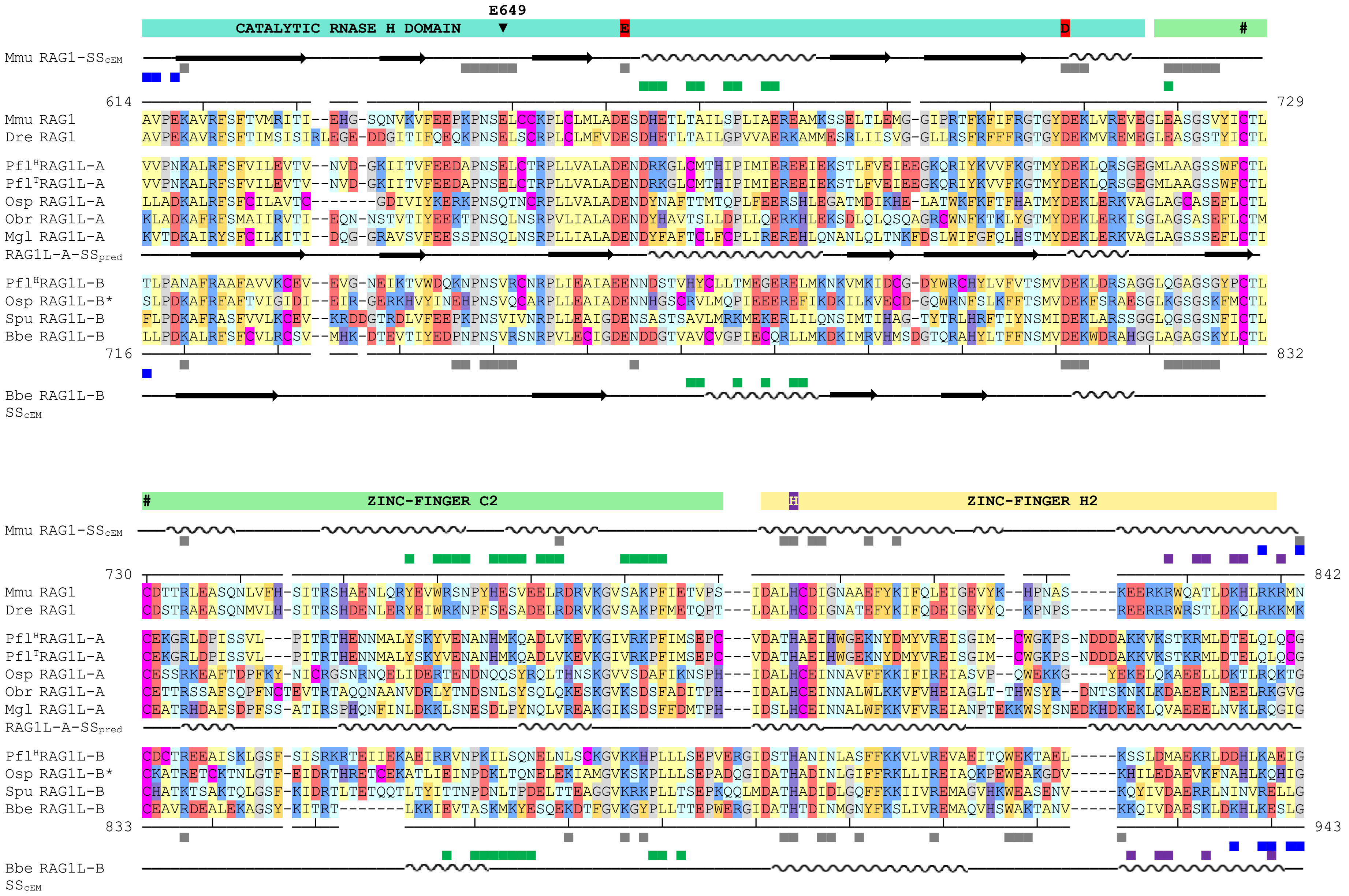

E649
CATALYTIC RNASE H DOMAIN
E
D
#
Mmu RAG1-SScEM
614
729
Mmu RAG1 AVPEKAVRFSFTVMRITI--EHG-SQNVKVFEEPKPNSELCCKPLCLMLADESDHETLTAILSPLIAERREAMKSSELTLEMG-GIPRTFKFIFRGTGYDEKLVREVEGLEASGSVYICTL
Dre RAG1 AVPEKAVRFSFTIMSISIRLEGE-DDGITIFQEQKPNSELSCRPLCLMFVDESDHETLTAILGPVVAERKAMMESRLIISVG-GLLRSFRFFFRGTGYDEKMVREMEGLEASGSTYICTL
Pfl^H RAG1L-A VVPNKALRFSFVILEVTV--NVD-GKIITVFEEDAPNSELCTRPLLVALADENDRKGLCMTHIPIMIEREEIEKSTLFVEIEEGKQRIYKVVFKGTMYDEKLQRSGEGMLAAGSSWFCTL
Pfl^T RAG1L-A VVPNKALRFSFVILEVTV--NVD-GKIITVFEEDAPNSELCTRPLLVALADENDRKGLCMTHIPIMIEREEIEKSTLFVEIEEGKQRIYKVVFKGTMYDEKLQRSGEGMLAAGSSWFCTL
Osp RAG1L-A LLADKALRFSFCILAVTC-------GDIVIYKERKPNSQTNCRPLLVALADENDYNAFTTMTQPLFEERSHLEGATMDIKHE-LATWKFKFTFHATMYDEKLERKVAGLAGCASEFLCTL
Obr RAG1L-A KLADKAFRFSMAIIRVTI--EQN-NSTVTIYEEKTPNSQLNSRPVLIALADENDYHAVTSLLDPLLQERKHLEKSDLQLQSQAGRCWNFKTKLYGTMYDEKLERKISGLAGSASEFLCTM
Mgl RAG1L-A KVTDKAIRYSFCILKITI--DQG-GRAVSVFEESSPNSQLNSRPLLIALADENDYFAFTCLFCPLIREREHLQNANLQLTNKFDSLWIFGFQLHSTMYDEKLERKVAGLAGSSSEFLCTI
RAG1L-A-SSpred
Pfl^H RAG1L-B TLPANAFRAAFAVVKCEV--EVG-NEIKTVWDQKNPNSVRCNRPLIEAIAEENNDSTVHYCLLTMEGERELMKNKVMKIDCG-DYWRCHYLVFVTSMVDEKLDRSAGGLQGAGSGYPCTL
Osp RAG1L-B* SLPDKAFRFAFTVIGIDI--EIR-GERKHVYINEHPNSVQCARPLLEAIADENNHGSCRVLMQPIEEEREFIKDKILKVECD-GQWRNFSLKFFTSMVDEKFSRAESGLKGSGSKFMCTL
Spu RAG1L-B FLPDKAFRASFVVLKCEV--KRDDGTRDLVFEEPKPNSVIVNRPLLEAIGDENSASTSAVLMRKMEKERLILQNSIMTIHAG-TYTRLHRFTIYNSMIDEKLARSSGGLQGSGSNFICTL
Bbe RAG1L-B LLPDKALRFSFCVLRCSV--MHK-DTEVTIYEDPNPNSVRSNRPVLECIGDENDDGTVAVCVGPIECQRLLMKDKIMRVHMSDGTQRAHYLTFFNSMVDEKWDRAHGGLAGAGSKYLCTL
716
832
Bbe RAG1L-B
SScEM
#
ZINC-FINGER C2
H
ZINC-FINGER H2
Mmu RAG1-SScEM
730
842
Mmu RAG1 CDTTRLEASQNLVFH-SITRSHAENLQRYEVWRSNPYHESVEELRDRVKGVSAKPFIETVPS---IDALHCDIGNAAEFYKIFQLEIGEVYK--HPNAS-----KEERKRWQATLDKHLRKRMN
Dre RAG1 CDSTRAEASQNMVLH-SITRSHDENLERYEIWRKNPFSESADELRDRVKGVSAKPFMETQPT---LDALHCDIGNATEFYKIFQDEIGEVYQ--KPNPS-----REERRRWRSTLDKQLRKKMK
Pfl^H RAG1L-A CEKGRLDPISSVL---PITRTHENNMALYSKYVENANHMKQADLVKEVKGIVRKPFIMSEPC---VDATHAEIHWGEKNYDMYVREISGIM--CWGKPS-NDDDAKKVKSTKRMLDTELQLQCG
Pfl^T RAG1L-A CEKGRLDPISSVL---PITRTHENNMALYSKYVENANHMKQADLVKEVKGIVRKPFIMSEPC---VDATHAEIHWGEKNYDMYVREISGIM--CWGKPS-NDDDAKKVKSTKRMLDTELQLQCG
Osp RAG1L-A CESSRKEAFTDPFKY-NICRGSNRNQELIDERTENDNQQSYRQLTHNSKGVVSDAFIKNSPH---IDALHCEINNAVFFKKIFIREIASVP--QWEKKG----YEKELQKAEELLDKTLRQKTG
Obr RAG1L-A CETTRSSAFSQPFNCTEVTRTAQQNAANVDRLYTNDSNLSYSQLQKESKGVKSDSFADITPH---IDALHCEINNALWLKKVFVHEIAGLT-THWSYR--DNTSKNKLKDAEERLNEELRKGVG
Mgl RAG1L-A CEATRHDAFSDPFSS-ATIRSPHQNFINLDKKLSNESDLPYNQLVREAKGIKSDSFFDMTPH---IDSLHCEINNALWFKKVFVREIANPTEKKWSYSNEDKHDKEKLQVAEEELNVKLRQGIG
RAG1L-A-SSpred
Pfl^H RAG1L-B CDCTREEAISKLGSF-SISRKRTEIIEKAEIRRVNPKILSQNELNLSCKGVKKHPLLLSEPVERGIDSTHANINLASFFKKVLVREVAEITQWEKTAEL-----KSSLDMAEKRLDDHLKAEIG
Osp RAG1L-B* CKATRETCKTNLGTF-EIDRTHRETCEKATLIEINPDKLTQNELEKIAMGVKSKPLLLSEPADQGIDATHADINLGIFFRKLLIREIAQKPEWEAKGDV-----KHILEDAEVKFNAHLKQHIG
Spu RAG1L-B CHATKTSAKTQLGSF-KIDRTLTETQQTLTYITTNPDNLTPDELTTEAGGVKRKPLLTSEPKQQLMDATHADIDLGQFFKKIIVREMAGVHKWEASENV-----KQYIVDAERRLNINVRELLG
Bbe RAG1L-B CEAVRDEALEKAGSY-KITRT-------LKKIEVTASKMKYESQEKDTFGVKGYPLLTTEPWERGIDATHTDINMGNYFKSLIVREMAQVHSWAKTANV-----KKQIVDAESKLDDHLKESLG
833
943
Bbe RAG1L-B
SScEM

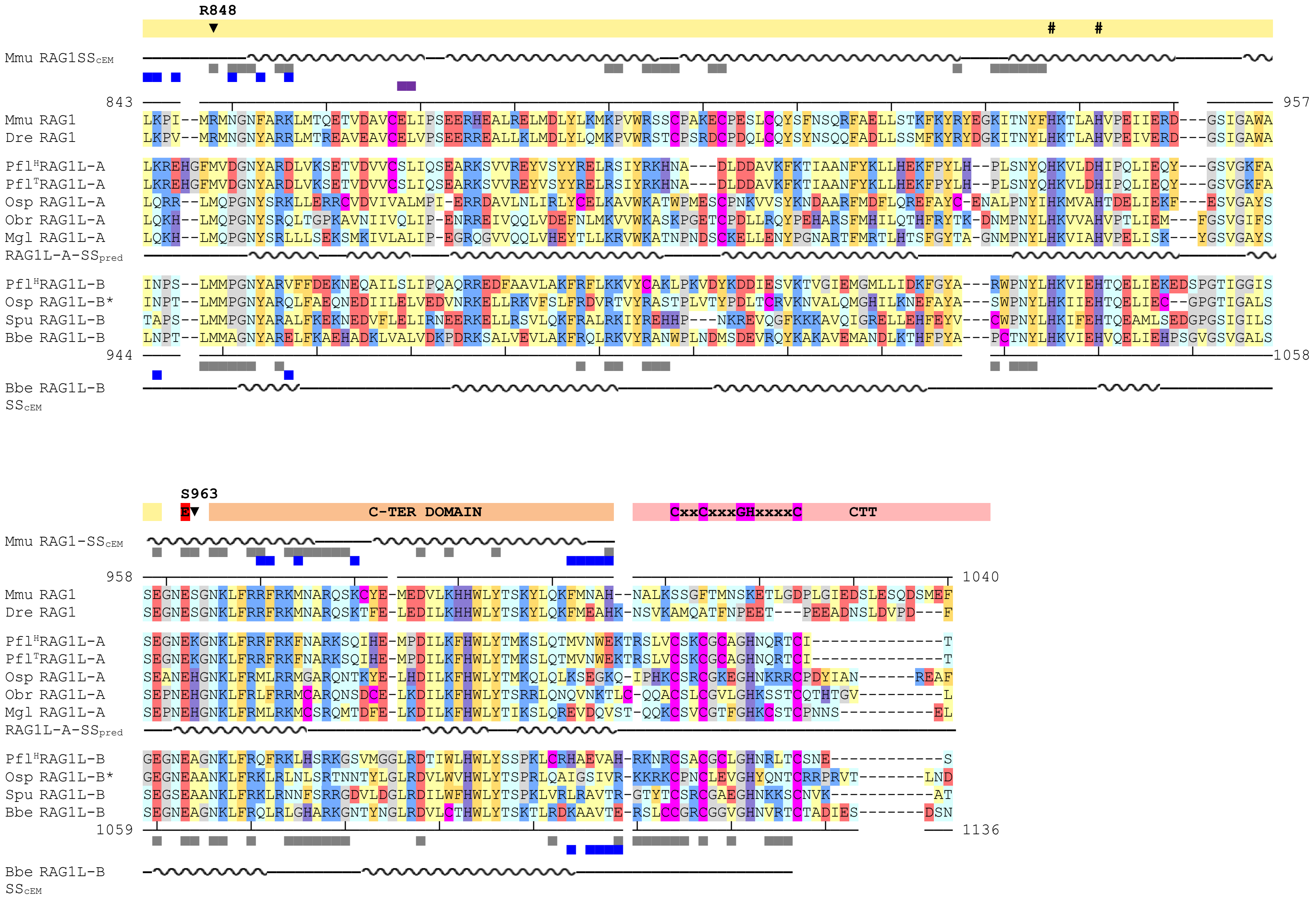


* OspRAG1L-B is less confident as it displays a potential splicing / frameshift (underlined in the sequence alignment)

Contacts and secondary structure is computed on the cryo-EM structures of mouse (5ze1) and *B.belcheri* (6b40)

### B. RAG2/RAG2L
#### Multiple Sequence Alignment

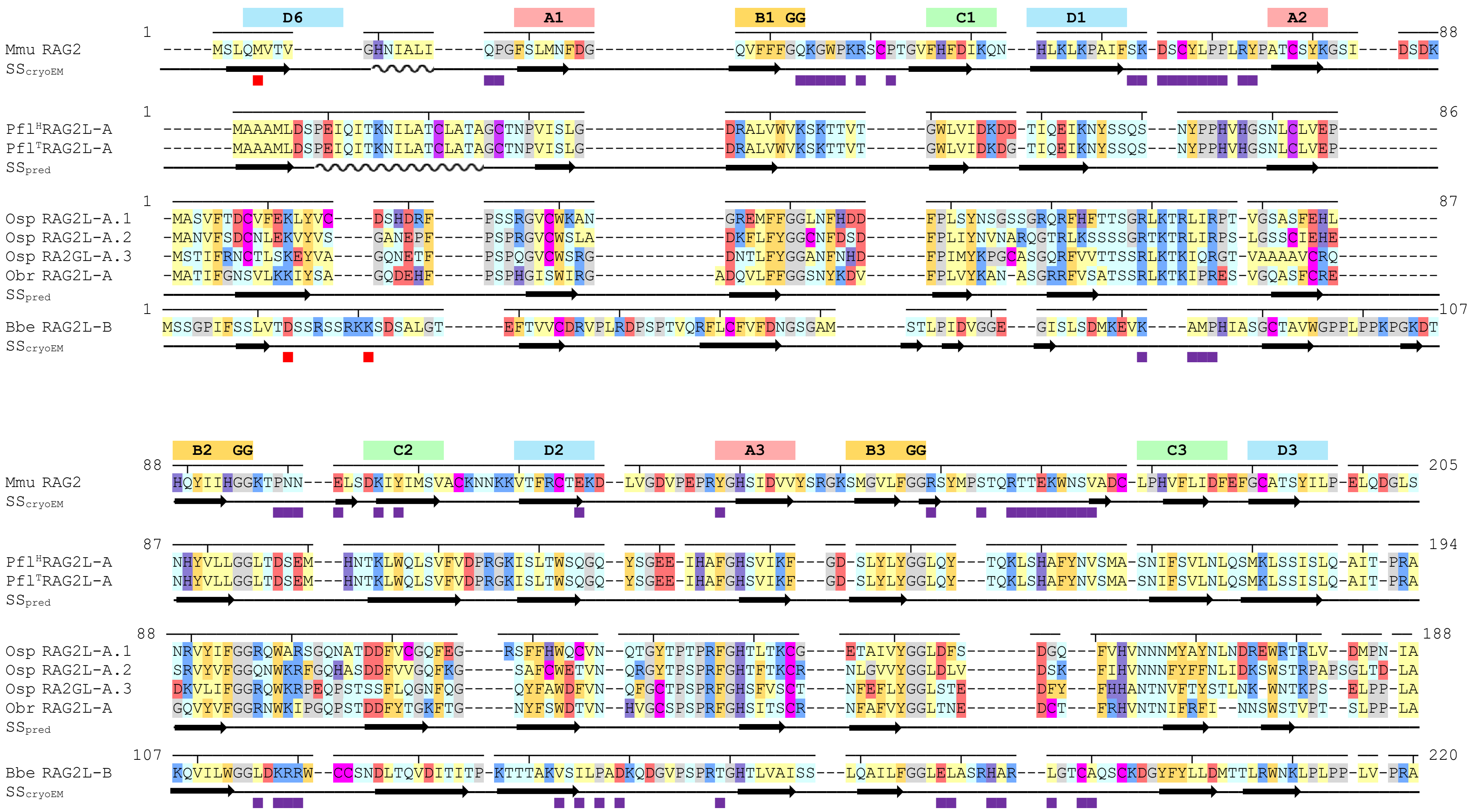

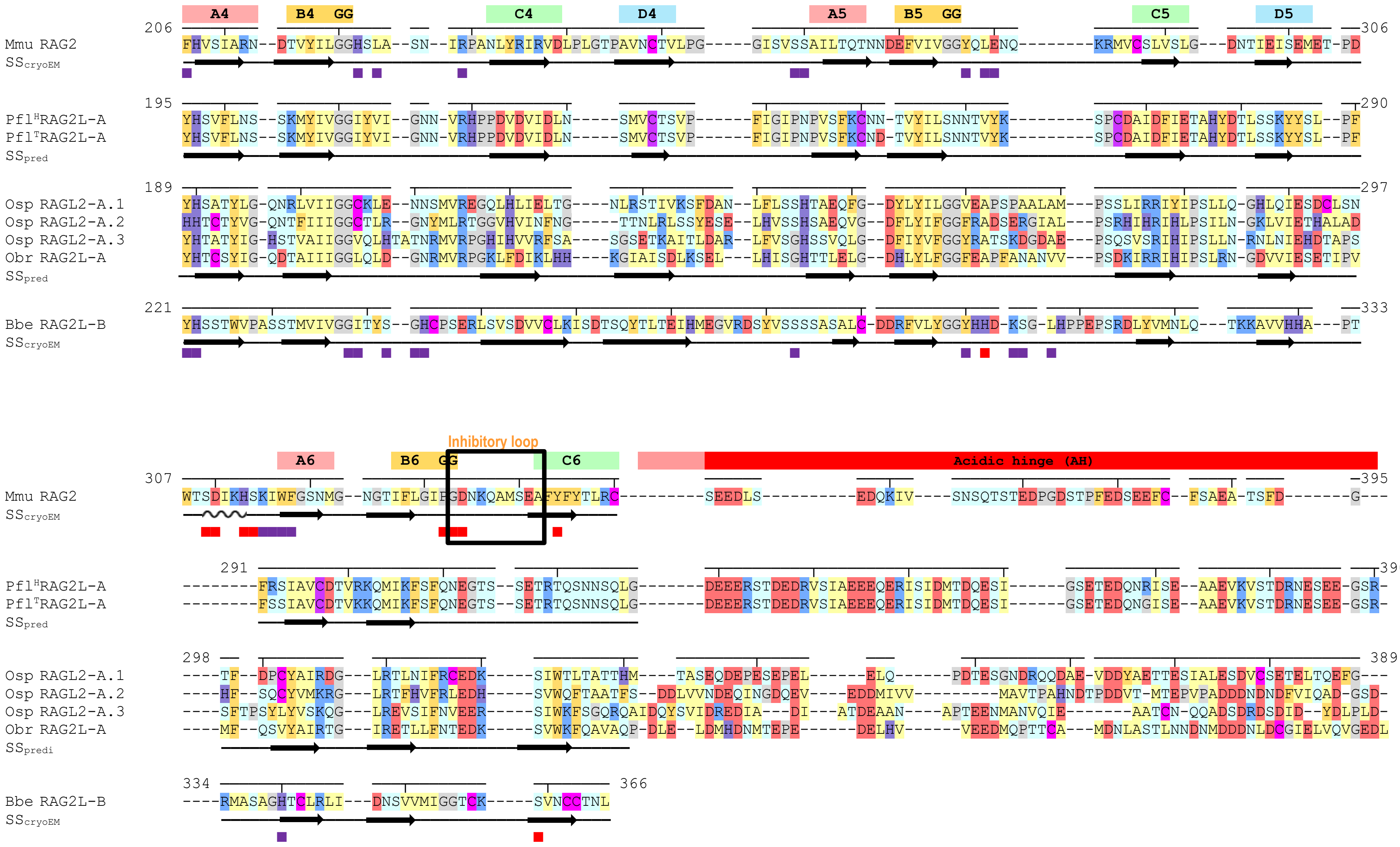

A4 B4 GG C4 D4 A5 B5 GG C5 D5
206 Mmu RAG2 FHVSIARN--DTVYILGGHSLA--SN--IRPANLYRIRVDLPLGTPAVNCTVLPG-----GISVSSAILTQTNNDEFVIVGGYQLENQ--------KRMVCSLVSLG---DNTIEISEMET-PD 306
SScryoEM
195 Pfl^H RAG2L-A YHSVFLNS--SKMYIVGGIYVI--GNN-VRHPPDVDVIDLN-----SMVCTSVP-------FIGIPNPVSFKCNN-TVYILSNNTVYK---------SPCDAIDFIETAHYDTLSSKYYSL--PF 290
Pfl^T RAG2L-A YHSVFLNS--SKMYIVGGIYVI--GNN-VRHPPDVDVIDLN-----SMVCTSVP-------FIGIPNPVSFKCND-TVYILSNNTVYK---------SPCDAIDFIETAHYDTLSSKYYSL--PF
SSpred
189 Osp RAGL2-A.1 YHSATYLG-QNRLVIIGGCKLE--NNSMVREGQLHLIELTG----NLRSTIVKSFDAN--LFLSSHTAEQFG--DYLYILGGVEAPSPAALAM---PSSLIRRIYIPSLLQ-GHLQIESDCLSN 297
Osp RAGL2-A.2 HHTCTYVG-QNTFIIIGGCTLR--GNYMLRTGGVHVINFNG-----TTNLRLSSYESE--LHVSSHSAEQVG--DFLYIFGGFRADSERGIAL---PSRHIHRIHLPSILN-GKLVIETHALAD
Osp RAGL2-A.3 YHTATYIG-HSTVAIIGGVQLHTATNRMVRPGHIHVVRFSA----SGSETKAITLDAR--LFVSGHSSVQLG--DFIYVFGGYRATSKDGDAE---PSQSVSRIHIPSLLN-RNLNIEHDTAPS
Obr RAG2L-A YHTCSYIG-QDTAIIIGGLQLD--GNRMVRPGKLFDIKLHH----KGIAISDLKSEL---LHISGHTTLELG--DHLYLFGGFEAPFANANVV---PSDKIRRIHIPSLRN-GDVVIESETIPV
SSpred
221 Bbe RAG2L-B YHSSTWVPASSTMVIVGGITYS--GHCPSERLSVSDVVCLKISDTSQYTLTEIHMEGVRDSYVSSSSASALC-DDRFVLYGGYHHD-KSG-LHPPEPSRDLYVMNLQ---TKKAVVHHA---PT 333
SScryoEM
Inhibitory loop
A6 B6 GG C6 Acidic hinge (AH)
307 Mmu RAG2 WTSDIKHSKIWFGSNMG--NGTIFLGIPGDNKQAMSEAFYFYTLRC---------SEEDLS----------EDQKIV----SNSQTSTEDPGDSTPFEDSEEFC--FSAEA-TSFD-------G--- 395
SScryoEM
291 Pfl^H RAG2L-A --------FRSIAVCDTVRKQMIKFSFQNEGTS--SETRTQSNNSQLG-------DEEERSTDEDRVSIAEEEQERISIDMTDQESI------GSETEDQNRISE--AAEVKVSTDRNESEE-GSR- 390
Pfl^T RAG2L-A --------FSSIAVCDTVKKQMIKFSFQNEGTS--SETRTQSNNSQLG-------DEEERSTDEDRVSIAEEEQERISIDMTDQESI------GSETEDQNGISE--AAEVKVSTDRNESEE-GSR-
SSpred
298 Osp RAGL2-A.1 ----TF--DPCYAIRDG---LRTLNIFRCEDK-----SIWTLTATTHM---TASEQDEPESEPEL------ELQ-------PDTESGNDRQQDAE-VDDYAETTESIALESDVCSETELTQEFG--- 389
Osp RAGL2-A.2 ----HF--SQCYVMKRG---LRTFHVFRLEDH-----SVWQFTAATFS--DDLVVNDEQINGDQEV----EDDMIVV---------MAVTPAHNDTPDDVT-MTEPVPADDDNDNDFVIQAD-GSD-
Osp RAGL2-A.3 ----SFTPSYLYVSKQG---LREVSIFNVEER-----SIWKFSGQRQAIDQYSVIDREDIA---DI---ATDEAAN----APTEENMANVQIE-------AATCN-QQADSDRDSDID--YDLPLD-
Obr RAG2L-A ----MF--QSVYAIRTG---IRETLLFNTEDK-----SVWKFQAVAQP-DLE--LDMHDNMTEPE------DELHV------VEEDMQPTTCA---MDNLASTLNNDNMDDDNLDCGIELVQVGEDL
SSpredi
334 Bbe RAG2L-B ----RMASAGHTCLRLI---DNSVVMIGGTCK-----SVNCCTNL 366
SScryoEM

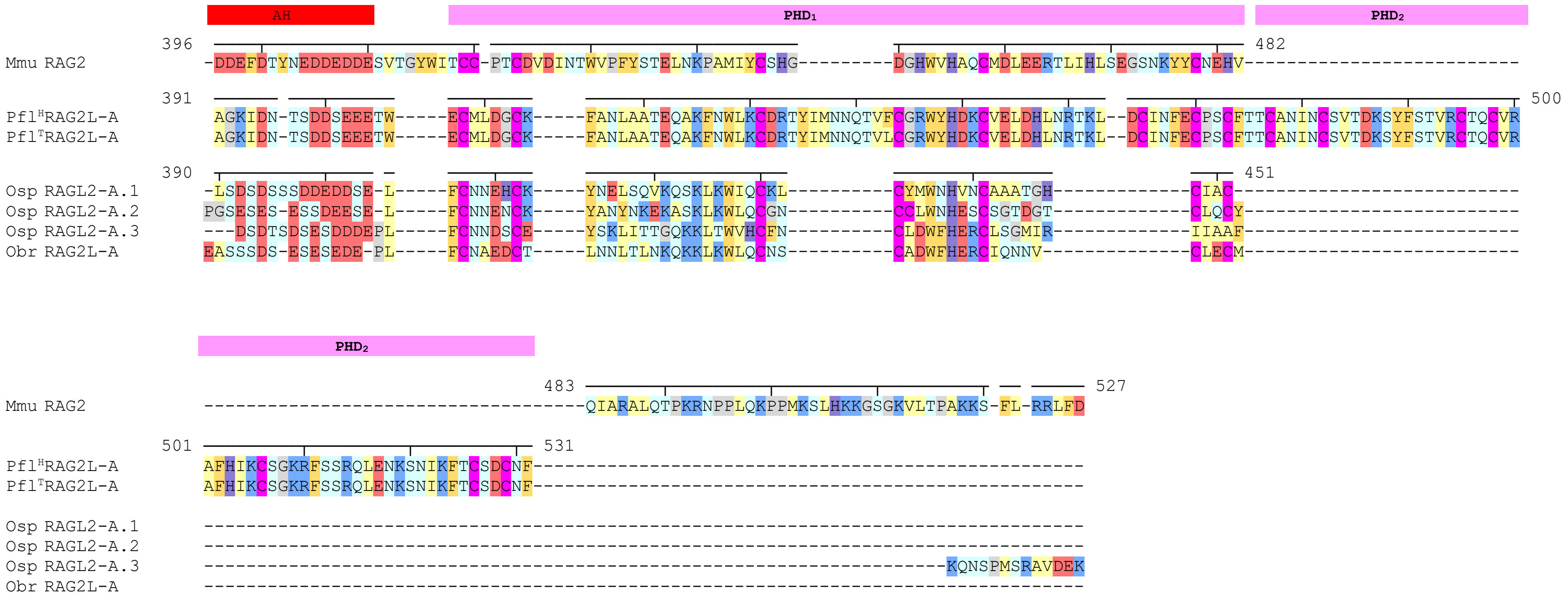
AH
PHD1
PHD2
396
Mmu RAG2 -DDEFDTYNEDDEDDESVTGYWITCC-PTCDVDINTWVPFYSTELNKPAMIYCSHG---------DGHWVHAQCMDLEERTLIHLSEGSNKYYCNEHV--------------------------
482
391
Pfl[H]RAG2L-A AGKIDN-TSDDSEEETW-----ECMLDGCK-----FANLAATEQAKFNWLKCDRTYIMNNQTVFCGRWYHDKCVELDHLNRTKL--DCINFECPSCFTTCANINCSVTDKSYFSTVRCTQCVR
Pfl[T]RAG2L-A AGKIDN-TSDDSEEETW-----ECMLDGCK-----FANLAATEQAKFNWLKCDRTYIMNNQTVLCGRWYHDKCVELDHLNRTKL--DCINFECPSCFTTCANINCSVTDKSYFSTVRCTQCVR
500
390
Osp RAGL2-A.1 -LSDSDSSSDDEDDSE-L-----FCNNEHCK-----YNELSQVKQSKLKWIQCKL----------CYMWNHVNCAAATGH-------------CIAC--------------------------
Osp RAGL2-A.2 PGSESES-ESSDEESE-L-----FCNNENCK-----YANYNKEKASKLKWLQCGN----------CCLWNHESCSGTDGT-------------CLQCY--------------------------
Osp RAGL2-A.3 ---DSDTSDSESDDDEPL-----FCNNDSCE-----YSKLITTGQKKLTWVHCFN----------CLDWFHERCLSGMIR-------------IIAAF--------------------------
Obr RAG2L-A EASSSDS-ESESEDE-PL-----FCNAEDCT-----LNNLTLNKQKKLKWLQCNS----------CADWFHERCIQNNV--------------CLECM--------------------------
451
PHD2
483
Mmu RAG2 -----------------------------------QIARALQTPKRNPPLQKPPMKSLHKKGSGKVLTPAKKS-FL-RRLFD
527
501
Pfl[H]RAG2L-A AFHIKCSGKRFSSRQLENKSNIKFTCSDCNF---------------------------------------------------
Pfl[T]RAG2L-A AFHIKCSGKRFSSRQLENKSNIKFTCSDCNF---------------------------------------------------
531
Osp RAGL2-A.1 ----------------------------------------------------------------------------------
Osp RAGL2-A.2 ----------------------------------------------------------------------------------
Osp RAGL2-A.3 ----------------------------------------------------------------------KQNSPMSRAVDEK
Obr RAG2L-A ----------------------------------------------------------------------------------
